# Predicting Relative Protein Abundance via Sequence-Based Information

**DOI:** 10.1101/2021.11.08.467260

**Authors:** Gregory M. Parkes, Robert M. Ewing, Mahesan Niranjan

## Abstract

Understanding the complex interactions between transcriptome and proteome is essential in uncovering cellular mechanisms both in health and disease contexts. The limited correlations between corresponding transcript and protein abundance suggest that regulatory processes tightly govern information flow surrounding transcription and translation, and beyond. In this study we adopt an approach which expands the feature scope that models the human proteome: we develop machine learning models that incorporate sequence-derived features (SDFs), sometimes in conjunction with corresponding mRNA levels. We develop a large resource of sequence-derived features which cover a significant proportion of the H. sapiens proteome, demonstrate which of these features are significant in prediction on multiple cell lines, and suggest insights into which biological processes can be explained using these features. We reveal that (a) SDFs are significantly better at protein abundance prediction across multiple cell lines both in steady-state and dynamic contexts, (b) that SDFs can cover the domain of translation with relative efficiency but struggle with cell-line specific pathways and (c) provide a resource which can be plugged into many subsequent protein-centric analyses.

## INTRODUCTION

Effective modelling across the ‘omics scales within a cellular environment plays a crucial role in understanding the principles that govern perturbations that lead to cancer or other diseases. The selection of alleles, RNA and protein relative abundance and epigenetic factors, all contribute significantly to the cell state and capacity to function. Further to this, features derived from DNA, RNA and protein primary sequences can contribute useful information in conjunction with measurements of expression levels to improve inference and control for steady-state variables. Recent technological advances in sequencing technology, for instance RNA-Seq (6) and Mass Spectrometry (MS) (7) have enabled the systemic analysis of gene expression and illuminated the limited predictability between transcriptomics and proteomics. A major limiting factor for many previous studies has been a lack of supportive data to coincide expression levels in the analysis across ‘omics scales.

Sequence-derived features (SDFs) are engineered features that are derived from the mRNA or protein of a respective gene. Early interest in SDFs came with the discovery of Kozak sequences (61), which are highly conserved short sequences which immediately precede the coding sequence. The translation of mRNAs is known to be influenced by a number of features associated with the 5’ untranslated region (UTR), such as the RNA secondary structure, the presence of ATG (methionine) codons, or upstream Open Reading Frames (uORFs). Further to this, the choice of codons in selecting an appropriate amino acid (given the codon has synonyms), also known as codon bias, has been demonstrated to correlate positively with mRNA level (13). Previous studies have highlighted the contribution of these codon/evolution-based features as predictive for protein abundance (13, 15, 51). In addition to translation initiation, elongation metrics have an impact on steady-state protein abundances, and are also related to the availability of tRNAs given certain codons. Likewise, mRNA processing and modifications such as polyadenylation have been shown to influence mRNA stability; the length of the poly-A tail correlates to the efficiency of the translation machinery. Numerous post-translational modifications (PTMs) are known to have an impact on protein abundance, in particular processes that target proteins for degradation, such as ubiquitination. They also promote functionalities such as cell signalling, protein folding and stability (16).

Due to the central dogma and reliability issues surrounding protein abundance measurement, mRNA expression has historically been taken as a reliable proxy for protein abundance (3, 5), although recent advances have added additional layers of complexity and weakened this assumption. This is due to complex post-transcriptional and post-translational interactions which modulate protein expression in a non-simple fashion. Previous studies in *S. cerevisiae* models have attempted to resolve this by building more complex statistical models which factor in SDFs, which in combination with mRNA expression, help to bridge the gap in relation to protein abundance (13, 15). Other authors have considered a vast selection of modelling techniques, such as Bayesian (11), coupled-mixture modelling (12) and outlier detection (13) to investigate transcriptome-proteome measurements. We have previously built upon this idea by exploiting outlier detection theory within a dynamic *H. sapiens* cell cycle context (8), integrating mRNA and translation rates (51) with selected SDFs. We found that even using a multi-’omics expression approach (mRNA, translation) without SDFs to model protein abundance produced correlations lower than expected (r=0.67) in a landmark cell cycle study (8). We found that including an additional 15 SDFs significantly improved corrected correlations against protein abundance, but that our feature scope was limited and had room for expansion. Whilst previously our focus was on the utility of PTMs to specifically uncover post-translational regulation processes, we think there is scope for a general-purpose role for SDFs.

## MATERIALS AND METHODS

Further details for the descriptions, plots and code for methodologies and preprocessing are described, implemented and plotted in the Jupyter notebooks provided in the Supplementary Data 3 (notebooks) and 4 (scripts).

### Data Retrieval

Human HeLa cell cycle data was taken from Aviner et al. (8) with triplicative expression measurements for mRNA, translation and protein. Microarray data is taken from the Gene Expression Omnibus (GSE26922), parsed using the GEOparse package. Protein levels for HeLa are pre-normalized using intensity-based absolute quantification (iBAQ) (35). U2OS, A431 and U251MG cell line expression data for mRNA and protein expression is taken from Lundberg et al. (55), whose RNA expression is estimated using RNA-seq, whereby RPKM values (58) were calculated for each RNA (see NCBI short-read archive with accession number SRA012517). Daoy cell line expression data and sequence-derived features are taken from Vogel et al. (2), whose cell line is cultured, collected and described previously (57). Gene expression values are estimated using Robust Multi-Array (RMA) analysis (34). Protein expression for all cell lines is estimated using MS/MS (Aviner, Lundberg) or LC-MS/MS (Vogel). Human mRNA transcripts and variants were taken from NCBI Entrez direct (36) or directly from NCBI FTP server, and loaded via the Biopython package (37) using the Python 3 programming language. Human Amino acid sequences were taken directly from Uniprot/Swissprot, selecting Proteome UP000005640. HGNC data (24) was taken from ‘genenames.org’ for the purpose of joining together RNA and protein datasets and sequence features. Post-translational modification (PTM) features were taken from PhosphoSitePlus (52) and counted. Gene Ontology terms were taken from the Gene Ontology Consortium (59), where labels to other related gene information were obtained via the Biomart portal (60).

### Data Preprocessing

HeLa Microarray data taken from Aviner et al. is mapped to GPL data, duplicates are eliminated and averaged across replicates. Timesteps 2h, 8h and 12h are mapped to S, G2/M and G1 of the cell cycle phases respectively in the HeLa protein dataset. RNA and protein expression sets from Lundberg et al (55) and Vogel et al. (2), whereby expression sets were preprocessed with 1) *log*_2_-transformation if not normally distributed, 2) missing values dropped 3) mRNA and protein expressions were merged together using a database intersection operations; achieved by utilizing a foreign key table as provided by Biomart, linking Refseq/Ensembl IDs to mRNA and Uniprot/Swissprot IDs to protein. Unique gene names (by HGNC) were mapped to NCBI accession numbers, whereby mRNA transcripts were split into CDS (coding sequence), 5’UTR and 3’UTR regions. Only mRNA transcripts that were curated were kept, which contained the NM/P label prefix (we eliminated all experimental/predicted sequences). Further to this, we check that all CDS sequences begin with ATG (methionine).

### Feature Extraction and Normalization

The number of exons, sequence-tagged sites (STS), misc features, regulatory regions and poly-adenylated tails in the mRNA transcript are counted. Protein sites, regions, molecular weight and more in the amino acid sequence are counted. We also count mono and di-nucleotide frequency for mRNA, CDS, 5’UTR and 3’UTR transcripts. Amino acid frequencies are calculated for the corresponding amino acid transcripts. Codon Adaptation Index (CAI) and Nc (effective number of codons) are calculated using CAIcal server (39) on CDS sequences, in conjunction with the Human Codon Usage table as reference. tRNA Adaptation Index (tAI) values are calculated using stAIcalc (41), with reference tRNA gene copy numbers from GtRNAdb (42) for hg19. CUB (relative codon usage) is calculated following the method shown by Roymondal et al. (43). Gibbs Free folding energy Δ*G* in 5’UTR regions, a proxy for mRNA secondary structure prediction, is calculated using the RNAstructure EnsembleEnergy algorithm (44), using window sizes of 10, 20, 30, 40, 60, and 100. We used ExPASy’s ProtParam (40) module, as a part of Biopython (37), to determine Aromaticity, Hydropathy (GRAVY), protein secondary structure and Isoelectric point (pI) features, among others. PEST regions for amino acid transcripts are calculated using the Emboss suite of the European Bioinformatics Institute (EBI) using a window size of 10. Text-based features associated with the metadata from Refseq/Uniprot are counted and normalized by the length of the corresponding mRNA/amino-acid transcript. For a full list and description of the features used, see S1 Table.

### Determination of correlations between heterogeneous feature types

To compute correlations *r_ij_* between all pairs of SDFs, we define criteria for algorithm selection as:

- features 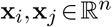: Spearman-rank method (*r_s_*)
- features 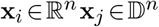: Point-biserial method (*r_bs_*)
- features 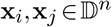: Spearman-rank method (*r_s_*)

where 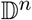 defines a dichotomous vector:

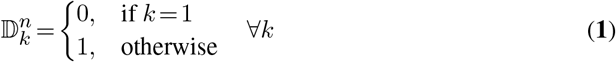

Here we denote *r*(*x, y*) as generic correlation form (be it Spearman-rank or point-biserial). Partial correlations (*r*(*s_i_, p_i_|m_i_*) as shown in Fig 3) are correlations between a SDF *s_i_* for gene *i* and the corresponding protein abundance *p_i_*, fixing mRNA expression *m_i_* as a covariate. Partial correlations can be estimated as the Pearson correlation between the residuals of the covariate/input and covariate/target vectors resultant from linear models. SDF features are sorted according to the direction of the Central Dogma of Molecular Biology (i.e DNA to protein). Where counts of mono- and di-nucleotide bases are considered, G+C and A+T bases are grouped together.

### Feature and Model Selection

Linear models between mRNA and protein levels are calculated using Ordinary Least Squares with 5-fold cross validation. To create models we use the Sci-kit Learn (45) package in Python. SDF-to-protein models are pre-filtered using a *missing value filter* (where features are dropped if 30% or more of the values are missing) and *low-variance filter* (where features with a variance less than 10^-6^ are dropped). We consider an algorithm space of linear and non-linear architectures, including regularized linear models, tree, ensemble and support vector machines. All remaining continuous SDF features are normalized using Z-score transformation prior to fitting. Dimensionality Reduction techniques such as PCA, Factor Analysis and Stratified PCA (i.e PCA by groups) are considered and explored, using the same set-up previously described. The same models and parameters selected for the whole proteome modelling were used for Gene Ontology subsets (as shown in Fig. 5). Gene Ontology terms were filtered such that they had at least 50 different genes that were associated with each term, for statistical robustness.

## RESULTS AND DISCUSSION

Previous work by Parkes and Niranjan (51) suggested that using an expanded set of sequence-derived features (SDFs) would increase the search space and thus eliminate more of the steady-state effects within the modelling of multi-’omics expression. In this cause we extracted over 200 sequence-derived features (SDFs) from both mRNA and amino-acid sequence data, of which 112 were derived from the NCBI Refseq repository (23) of 50007 human mRNA curated transcripts, with the remaining 78 from 20395 Uniprot/Swissprot human amino-acid curated transcripts, representing a near-complete coverage of the human transcriptome/proteome. In terms of mRNA sequence data, features extracted are split into the primary gene regions; coding sequence (CDS), 5’UTR and 3’UTR, as well as the whole sequence (mRNA). A full list of the features are described here (S1 Table). In addition to this, we associate post-translational modification (PTM) data from PhosphoSitePlus (52), also derived from amino-acid sequences. See Methods for descriptions on each feature and how they were extracted/measured.

### Not All Features Are Made Equal: Dominance Of Count-Based Features

The majority of engineered features are a form of count feature measuring a biological phenomenon, whereby a majority are derived from the frequency of mono-, di- or tri-nucleotide occurrence. A breakdown of sequence-derived features by broad affiliation and source (Fig 1.) shows a majority of features derived from mRNA (55.3%), a third from amino acid (33.4%) and the remaining PTMs constituting 4% of the features. HGNC features (24) consist of identifiers only and is therefore not used in downstream analyses. The imbalance between mRNA/amino acid features is partly due to the richer GenBank format which stores larger amounts of meta information regarding the mRNA transcripts.

**Figure 1.**
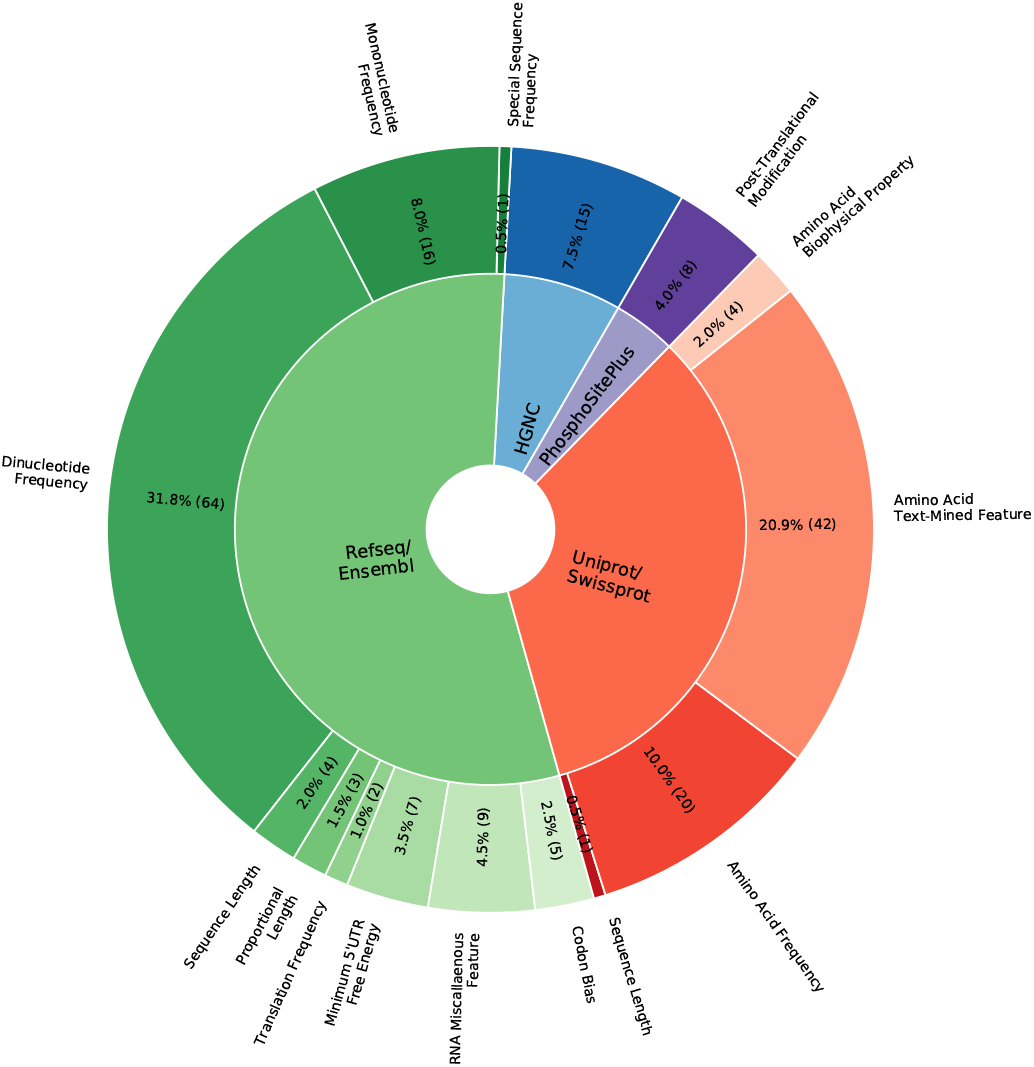
The Breakdown of Sequence-Derived Features and their sources. A pie chart detailing the number and proportion of features by database source (inner ring) and broad data-grouping (outer ring). mRNA/Ensembl/Refseq-like features (54%) are denoted in shades of green, amino-acid/Uniprot/Swissprot-like features (33%) are denoted in shades of red. Post-Translational Modification (4%) features are extracted separately (in purple). HGNC labels (7%) are not included in the machine-learning algorithms but are shown here for completeness (in blue).

The high prevalence of nucleotide frequency features in mRNA (71.9%) and minority amino acid frequency in proteins (29.9%) lead to a heavy length-based dependency, which we correct for by scaling by the appropriate length, or as a ratio to the expected frequency. The frequency of bases, and indeed the off-frequency of particular codons, otherwise known as *codon bias*, is known to correlate with post-transcriptional regulation, mRNA decay and influences translation (2, 25, 53, 54). Continuous feature groups include estimations of the Minimum Free Energy (MFE) of the 5’UTR region (using various window sizes) and amino acid biophysical properties, such as Isoelectric point (62) and GRAVY (63).

There are areas of SDF coverage that are poor, such as estimated modifications on the DNA template (proxies for DNA methylation, histone modification) or secondary structure predictions in non-5’UTR regions. In this work we decided to prioritise features closer to protein expression, meaning that extracting features from the DNA template was a lower priority. In addition to this, simulations to estimate mRNA secondary structure can be expensive and grows exponentially as the length of molecule increases; thus this was infeasible for this study.

### Many Sequence-Derived Features Exhibit Multicollinearity, But May Still Contribute To Understanding Expression Levels

As many of these SDFs are not entirely independent of each other, we would expect large levels of inter-correlation between features of the same type and source (Fig 2., see S1 Figure for *interactive version*, see Methods for details on calculation); particularly prevalent are the strong positive correlations between CG-containing mRNA base ratios, and likewise for AT-containing ratios. The strongest correlations are between mono/di-nucleotide ratios by mRNA, 3’UTR and CDS regions for each gene, as well as the minimum free energy (MFE) calculations in the 5’UTR region (as the window size is all that changes). Due to the need to eliminate length bias for count-based features, normalizing by length introduces intracorrelation between these features as a common factor. In addition to this, the majority of the spurious off-diagonal correlations that are high relate to cross-talk between the mRNA/Uniprot datasets having count features with the same description: for example the ‘signal peptide count’ (*r_bs_* = 0.69), ‘peptide’ (*r_bs_* = 0.5) and ‘transit-peptide’ (*r_bs_* = 0.63) count features had an equivalently named-feature in Uniprot. In general, the relationship between CDS and amino-acid derived features was very strong, this is largely because of the linear 3:1 mapping between codons and amino acids as determined by the genetic code; biophysical properties also correlated well with amino-acid/CDS frequency ratios. These findings are consistent to previous work done by Vogel et al. (2), albeit with a slightly more varied feature set; feature correlation overall tends to cluster by gene region and favours features that are biologically closer to it, as we successfully demonstrate (see S2 Figure). However, Vogel’s group did not consider features derived from text-based information or make use of meta information provided in these databases, as many of them were still under development at the time of research.

**Figure 2.**
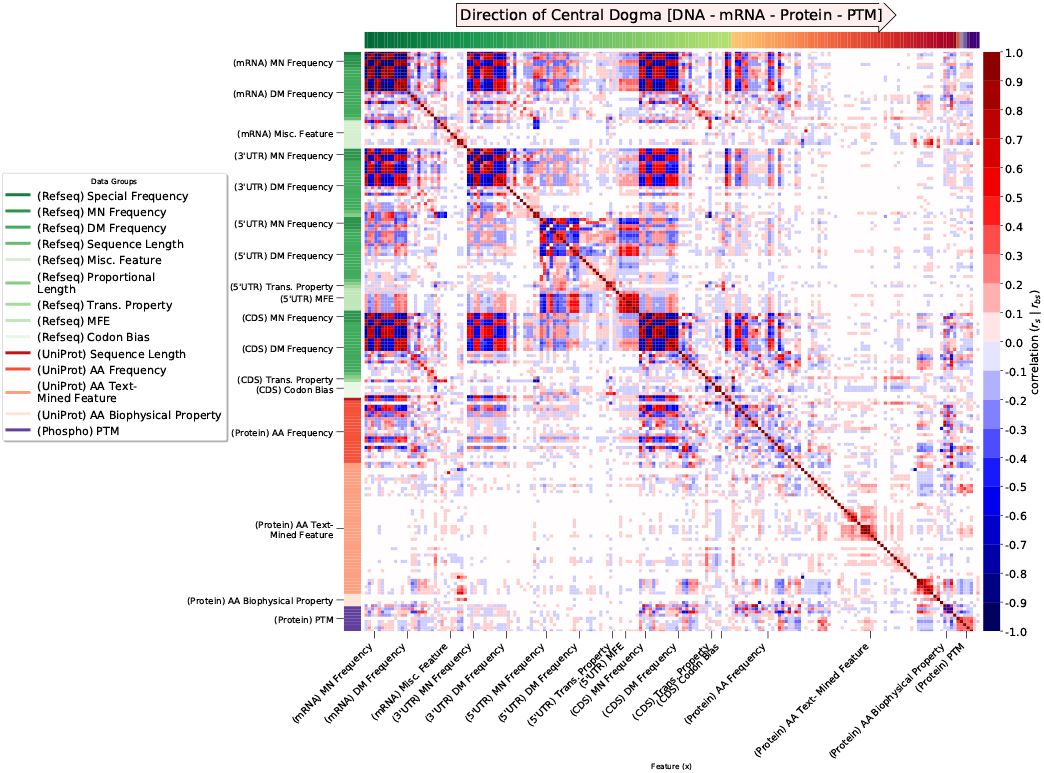
Large-scale interdependencies between SDFs reveal source dependency. Spearman-rank (*r_s_*)/Biserial correlation (*r_bs_*)-mixed correlation matrix between sequence-derived features (SDFs). See Methods for details on correlation method. Both axes indicate the direction of molecular biology (from DNA to post-protein). mRNA/RNA features are denoted in green shades, amino-acid features are denoted in red shades, PTM features are denoted in purple. Off-diagonal elements indicate a correlation (*r_s_*—*r_bs_*) between two features (red: positive, blue: negative).

The propensity of each feature (see S3 Figure.) to correlate with all of the others (calculated as the *ℓ*_1_-norm, see Methods) reveals that mRNA, CDS and 3’UTR features have high intracorrelation, particularly mono-/di-nucleotide base frequencies that consist of GC/AT-only bases. Of codon biases, which historically have been used extensively in previous studies as effective mRNA proxies, codon adaptation index (CAI) scores highly, but other metrics do not. In general, high ∥*r*∥ suggest redundancy, whereas low ∥*r*∥ may have a strictly non-linear relationship, or bare no resemblance to the problem domain. The most popular measures of correlation include Pearson and Spearman-rank correlation, but both of these methods depict either linear or monotonic relationships respectively, and struggle to model non-linear relationships. Therefore to account for this effect, we checked the Mutual Information (MI) between all SDFs (see Supplementary Figure 3.), which broadly depicts the same patterns in terms of how the features group, but due to the heterogeneity of data type it underestimates boolean-like features.

In application to the problem of protein prediction via mRNA proxy, the question now becomes whether these features can bare any relationship to experimental expression datasets, with the attributable noise that comes from spacetime measurements.

### The Experimental Procedure Of A Study Plays A Substantive Role In Discerning Relationships Between SDFs And Expression

The relationship between log-normalized protein and mRNA relative abundances is non-linear, but can be estimated using piece-wise linear functions (2) and is observed in many studies (3, 8, 13, 15, 51). However, a significant portion of SDFs are not normally distributed or cannot be easily transformed to be distributed as such; this presents a challenge in emulating the kind of information contained in the mRNA expression to be represented by SDFs. We calculate the monotonic relationship between each SDF and the corresponding mRNA, protein (left) and partial protein (right) expression for 5 different *H. sapiens* cell lines (*Daoy* (2), *A431, U2OS, U251MG* (55), *HeLa* (8)), corresponding to three different studies, to explore the capacity of SDFs as proxies (Fig 3.). We define ‘partial protein’ as the correlation between SDF-to-protein, controlling for mRNA level. Whereby ‘y=x’ represents features that correlate just as much with mRNA as protein abundance (meaning the feature is agnostic), in all cell lines we notice that post-translational modifications (PTM) correlate strongest with both mRNA and protein abundance; particularly Ubiquitination (*r* = 0.4, 0.6) which is associated with protein degradation. mRNA/protein length features are strongly negative with respect to protein level, particularly in HeLa/Daoy cell lines (*r* = −0.4, −0.6). The majority of changes in correlation when mRNA is controlled for is not substantial; most features are corrected for by Δ*r*< 0.2 (see S5-6 Figure.), and nearly always converging the correlation towards zero, rather than increasing it. Similar observations are made when the sequence length (mRNA/amino acid) is controlled for instead of mRNA level (see Supplementary Material 5.), particularly codon bias features are appropriately reduced in correlative power.

**Figure 3.**
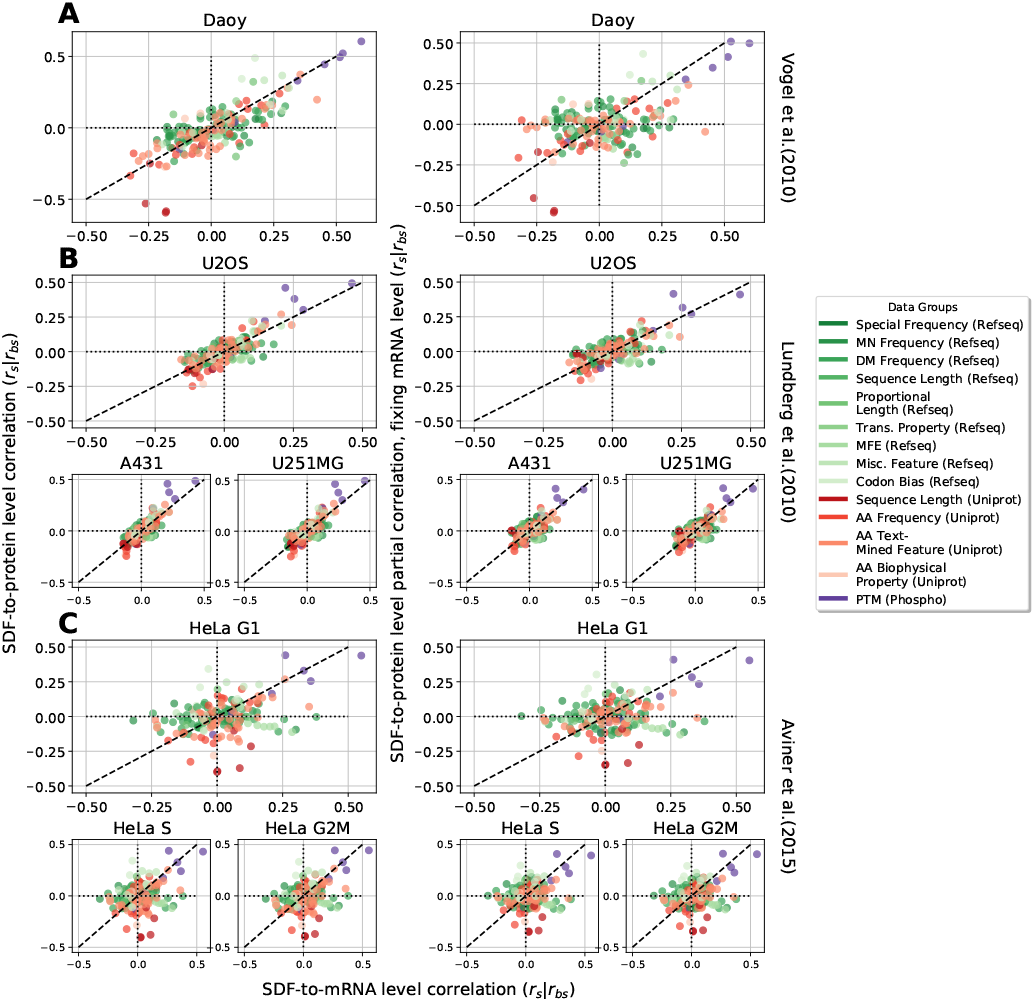
Variations in a priori experimental dataset assumptions considerably alter SDF impactfulness. Scatterplots of Spearman-rank (*r_s_*)/Biserial correlation (*r_bs_*)-mixed correlation between sequence-derived features (SDFs) and mRNA (x-axis), protein (y-axis, left) and partial correlations of protein fixing mRNA abundance (y-axis, right). From top; (A) Daoy (Vogel et al.), (B) U2OS;A431;U251MG (Lundberg et al.), and (C) HeLa G1;S;G2/M (Aviner et al.) from different mRNA-protein expression datasets. Each point represents a correlation between an SDF and an expression dataset. See Methods for details on correlation method. mRNA/RNA features are denoted in green shades, amino-acid features are denoted in red shades, PTM features are denoted in purple. All correlations have a *p*-value < 0.05.

One clear trend is the ‘flattening’ of the relationship for partial-correlations as the fixing of mRNA level has the tendency to drive partial-protein correlations to 0, and this is particularly noticeable in the Doay and HeLa cell lines. The ‘spread’ of correlated features is substantially higher in Aviner’s dataset than Lundberg; we suspect this is because HeLa is drawn from a cell cycle study where dynamic time-effects are not removed in the preprocessing pipeline, whereas Lundberg measurements are aggregated and more likely to conform to ‘steady-state’ protein levels. This loss of information could therefore lead to small correlation coefficients in these sets. Alternatively, differences in RNA sequencing technology (Aviner et al.; microarray, Lundberg et al.; RNA-Seq) are known to vary in variance and this may be reflected in the variance of correlation coefficients. These results are checked against the estimated mutual dependency (MI) conditioned against mRNA level or gene length, where we find similar loss in the relationship between SDFs and protein (see S5-6 Figure.), in particular we see that the mutual information reflects the same relationship either against mRNA or protein level, or both.

### Sequence-Derived Features Alone and In Conjunction with mRNA Expression Reliably Improve Protein Abundance Prediction

In order to overcome the issues surrounding the interpretation of variables affected by multicollinearity, there are a number of avenues to choose including: (a) Dimensionality Reduction, (b) Feature Selection and/or (c) Increased model complexity. We compared 13 different methods of feature selection and dimensionality reduction (see Supplementary Material notebooks), including more detailed analyses using Principle Component Analysis (PCA) and Factor Analysis (FA) techniques. Whilst dimensionality reduction techniques help to eliminate multicollinearity, the latent sub-spaces remove direct interpretation of the features of interest, and feature selection can sometimes be arbitrary when features are similar in selecting one of the close features.

Next, we developed machine learning models that incorporated our SDFs as input, to predict protein abundance levels as a target; to achieve this we performed model selection analysis to find the best technique and hyperparameter selection across linear and non-linear algorithm architectures. We found that across all 5 cell lines, tree-based models such as Gradient-boosted regression trees performed best, with the exception of the Daoy dataset, where ElasticNet performed best (see Supplementary Material Notebooks for details on the selection of parameters, graphs and more). To provide an effective comparison, we ran the same model using the SDFs provided by Vogel (2) across all cell lines (see Fig 4A, with 5-fold cross validation); in all cases these updated features outperform previous work and/or substantially reduce the error variance. Notably, there is a high degree of similarity between the cell line data sources (Aviner et al. (8), for HeLa, etc), indicating that the technique of mRNA/protein measurement and/or normalization methods may play a significant role in determining the efficacy of SDF feature use. The lack of root-mean squared error (RMSE) reduction in the HeLa lines may be due to the dynamic nature of cell cycle activities, whereby SDF features struggle to predict nonsteady state protein expression levels. Further to this, our SDF features have been calculated across the entire transcriptome, and are able to capture a much higher percentage of the proteome than previous feature sets (see Fig 4B); with the exception of the relatively small Daoy expression set, and assuming the hypothesis of ‘one gene to one protein’, we cover roughly 20-25% of the human proteome (around 4-5k proteins) in these example datasets.

**Figure 4.**
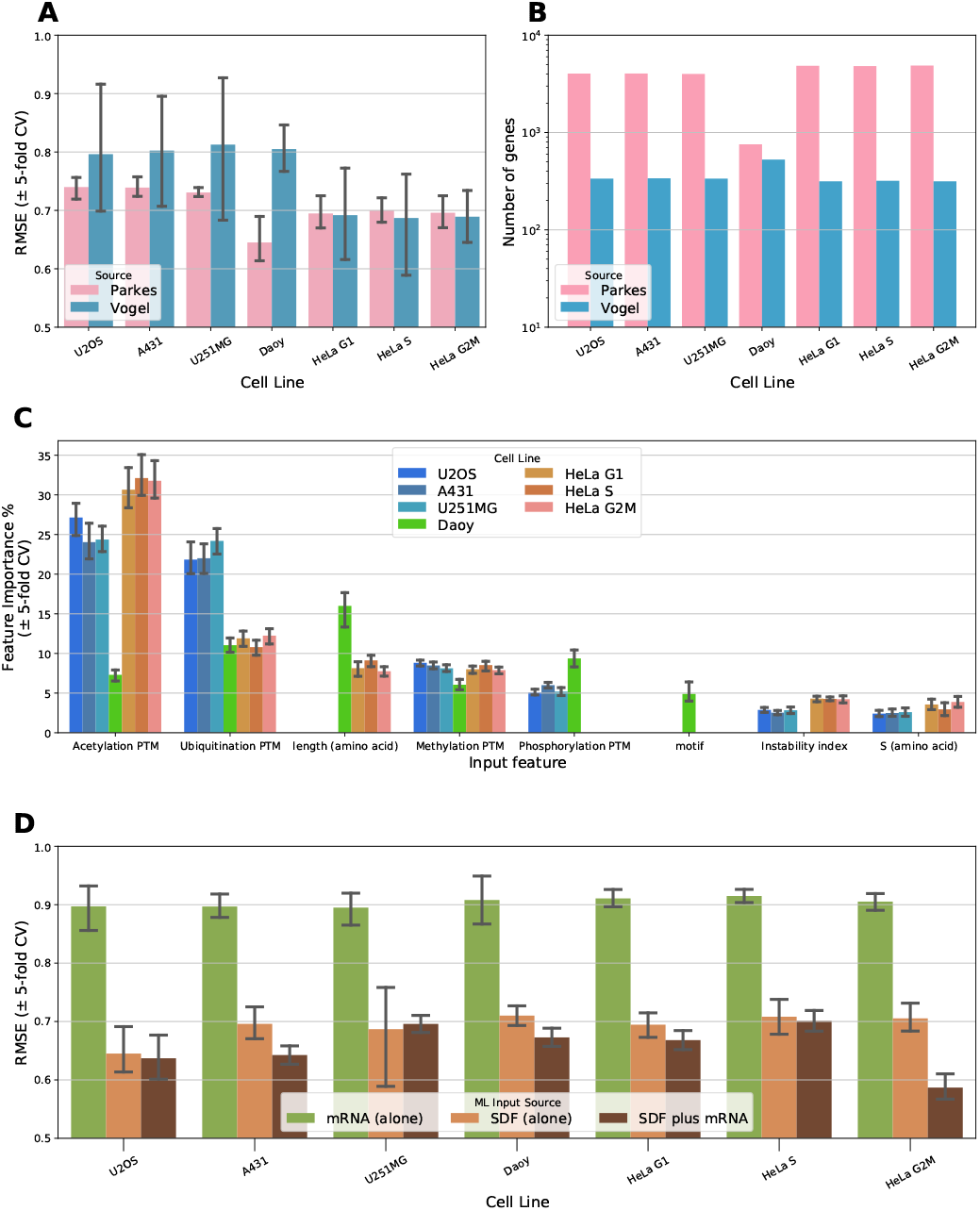
Substantial improvement in SDF inference capacity with respect to protein abundance to mRNA. Barplots of (A) RMSE estimations (± 5-fold CV) as a result of ML regression models across different cell lines for whole input sources, (B) Sample sizes for each resultant cell line ML model by input source, (C) Feature weights as a relative importance by the 6 most important features by cell line, (D) Whole genome RMSE estimations (± 5-fold CV) by cell line depending on input features (using just mRNA expression, just SDFs, or both). See Methods and Supplementary Material for details on model and feature preprocessing/selection.

To explore which of the features are most important, we took the best models by cell line and looked at the relative importance as weighted by the tree-models (see Fig 4C, with 5-fold out-of-sample cross validation), whereby a dominance of post-translational modification (PTM) features and length associated with the amino acid sequence appear dominant for all but the Daoy cell line. Both Acetylation and Ubiquitination are associated with protein stability, where Acetylation also deals with protein localization and synthesis, whereas Ubiquitination is also associated with cell cycle division and immune response reaction (56). Sequence length is consistent with previous studies (58) but relationships between expression and PTMs have seen less interest in the literature. Compared to models that just use mRNA expression levels (with Ordinary Least Squares), the inclusion of SDFs appears to reduce model error by about 25-30% across all cell lines (see Fig 4D, with 5-fold cross validation), with both mRNA levels and SDFs acting to improve RMSE slightly and to reduce model error variance.

To determine whether the sequence feature set from Vogel (2) covered a similar feature space to our own, we performed Canonical Correlation Analysis (CCA) between the two datasets (see S1 Figure). We discovered there was very limited overlap in the feature space, which we suspect is due to the time difference between the studies, and thus the evolution in the underlying genetic databases which form the basis on which SDFs are extracted. This remains true when we filter for just the subset of genes chosen in Vogel’s sequence feature set. It is wholly possible however that, as a number of features break inter-independency assumptions, that statistical insight from this check may be questionable.

### Sequence-Derived Features Aid Prediction By Capturing Information Regarding The Translation Process

Whilst we have begun to show that SDFs as a whole work across the human proteome, it remained unclear which functions and biological processes were benefiting the most from these features as input, and which processes were not being covered by this approach. Thus, using the same models (with hyper-parameters fitted), instead of using the entire expression dataset, we re-trained models whereby the out-of-sample test set consisted of all of the genes which shared a particular Gene Ontology (GO) term, and the training set consisted of every other gene not in this group. We filtered for GO terms which had at least 50 proteins associated with the term, to allow for statistical robustness. This process was repeated for models with different design matrices; containing just mRNA level, just SDF features, or both as input (see Fig 5A, see S5 Table for full GO analysis). Next, we selected the 10 GO term models which have the lowest (Fig 5B), highest (Fig 5C) and the most-improved (Fig 5D) average RMSE score across all cell lines. The lowest RMSE models, similarly to the global proteome model, reflect a 25-30% improvement in abundance prediction compared to the worst term models (see S5 Table for full GO analysis). In the vast majority of cases, significant improvement in prediction is achieved with the introduction of static SDFs into the input design matrix. Noticeably, the lowest and most-improved term sets have a good coverage of translation-situated terms, such as translation elongation and initiation, as would be expected by heavy usage of codon bias and frequency-based features, and such relationships have been observed in previous studies (2, 13, 14).

**Figure 5.**
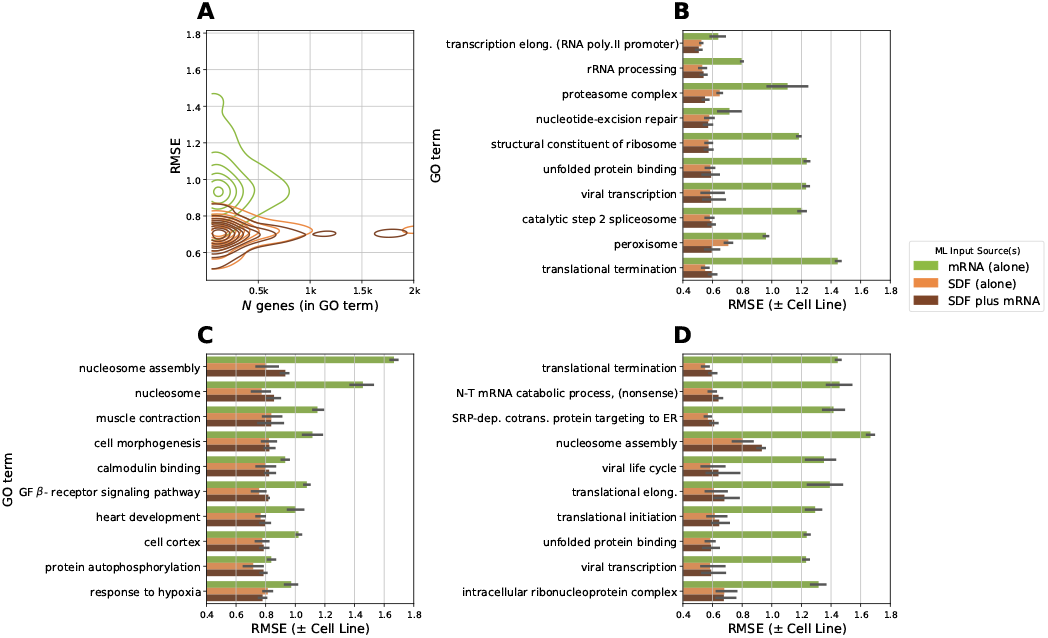
Representation of translation-oriented GO terms predominant in best SDF-based models. Out-of-sample root mean squared error (RMSE) scores of Gradient-boosted regression trees (GBRT) by Gene Ontology (GO) terms. (A) Bivariate kernel density estimation of the number of genes by RMSE for each GO term model by data input source. Barplots of the 10 (B) lowest, (C) highest and (D) most improved (difference between SDF plus mRNA and mRNA alone) RMSE-scoring GO term models by data input sources. Within GO; Biological Process, Cellular Component and Molecular Function terms are included. See Methods and Supplementary Material for details on model and feature preprocessing/selection.

Similarly. the selection of ribosome-oriented terms is expected, given the high correlation between mRNA and protein levels between ribosome-associated genes, which impacts on model performance. More surprising is the selection of protein localization (such as SRP-dependent co-translational protein targeting to ER), and mRNA/protein decay by translational termination and mRNA catabolic process terms. We did not include mRNA or protein halflife features by direct measurement as a part of the input features, so it is interesting that there are aspects of the SDFs that can predict these functions. Genes that underperform are associated with labile proteins that are associated with development (heart development, cell morphogenesis), complex signalling pathways such as response to hypoxia or cell-line specific functionalities not covered in the cell lines we modelled on, such as heart or muscle tissue (e.g muscle contraction).

## CONCLUSION

We present a large-scale analysis of the impact of SDFs on comprehensive prediction of protein abundance and function across a number of significant *H. sapiens* cell lines, both in a steady-state and dynamic (cell cycle) context. We demonstrate that, whilst introducing new features can present challenges in terms of model robustness, the contribution of these extracted sequence-based features can comprehensively cover many aspects of translation 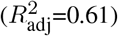 to inform probable protein abundance output, especially when mRNA abundance is also factored. This study goes further than previous analyses in retaining a relatively high coverage of the entire proteome, with these features having the ability to be applied to any expression dataset scenario. These features are provided with this study and can be used as a resource in application to a wide range of protein-centric problems, including protein abundance prediction as demonstrated here. However, outstanding questions remain as to what other latent variables SDFs may contribute to in terms of biological understanding (still an active area of discussion), and the fact remains that approximately one-third of proteins cannot be explained by steady-state features which hinder further model accuracy. Potential methods to overcome this without time-dependent measurement could include the integration of protein-protein interaction (PPI) pathways, and/or measured estimates of protein and mRNA half-life which would be the subject of further investigation.

## Supporting information

Supplementary Figures (html)

## DATA AVAILABILITY

All processed files are submitted as supplementary material, and are available on our Github: https://www.github.com/gregparkes/sdf_paper. Other sources (such as sequence-derived features) can be originally obtained from their respective open-access databases. See the Materials and methods section for further details.

## ACKNOWLEDGEMENTS

The proofreading from Xin Du and Frixos Papadopoulos is gratefully acknowledged.

## SUPPLEMENTARY FIGURES

In order to increase transparency and accessibility with the large dataset, we provide a number of *interactive* figures corresponding to the main figures in this paper, in a ‘.html’ format with embedded standalone Javascript. In order to view them, open the files within any Web Browser (best results using Google Chrome or Firefox). Plots are made using Bokeh: https://bokeh.org.

**S1 Figure.**

**Large-scale interdependencies between SDFs reveal source dependency (Interactive)** Spearman-rank (*r_s_*)/Biserial correlation (*r_bs_*)-mixed correlation matrix between sequence-derived features (SDFs). See Methods for details on correlation method. Both axes indicate the direction of molecular biology (from DNA to post-protein). mRNA/RNA features are denoted in green shades, amino-acid features are denoted in red shades, PTM features are denoted in purple. Off-diagonal elements indicate a correlation (*r_s_—r_bs_*) between two features (red: positive, blue: negative). Hover over squares to reveal additional meta information including *n*, method used, and *p*-value. Tools to interact are on top-right corner of plot.

**S2 Figure.**

**Interdependencies between SDFs from Vogel et al. 2010 Dataset.** Spearman-rank (*r_s_*) correlation matrix (as heatmap) between SDFs as derived from Vogel et al. See Methods for details on correlation method. The data is not sorted in any particular way and is as presented in the order of the data as provided. Blue colors indicate negative correlation, red indicate positive correlation. Tools to interact are on top-right corner of plot.

**S3 Figure.**

**Magnitude of feature importance as determined by correlations.** Barplot of *ℓ*_1_-norm values for each row of correlations ∥*r_k_*∥. Features with extremely high *ℓ*_1_ may indicate heavy redundancy, whereas features with low *ℓ*_1_ may not be valuable as features, have a strictly nonlinear relationship or only come into play when other intercorrelations are removed. Tools to interact are on topright corner of plot.

**S4 Figure.**

**Correlation changes when partials are factored in, factoring mRNA.** Scatterplots of the change in correlation from partial-correlation to protein factoring mRNA to mRNA (x) against partial-correlation to protein factoring mRNA to protein (y). Large delta changes indicate a significant impact of mRNA factor in correcting the correlation, indicating potential feature redundancy, or features that need heavy correcting for. Useful in the knowledge of which features can aid in appropriate mRNA proxying for protein abundance.

**S5 Figure.**

**Correlation changes when partials are factored in, factoring Length.** Scatterplots of the change in correlation from partial-correlation to protein factoring Length to mRNA (x) against partial-correlation to protein factoring Length to protein (y). Large delta changes indicate a significant impact of Length factor in correcting the correlation, indicating potential feature redundancy, or features that need heavy correcting for. Useful in the knowledge of which features can aid in appropriate mRNA proxying for protein abundance.

## SUPPLEMENTARY DATA

**S1 Table.**

**Full description of feature names, groups and sources.** Includes code names, the groups to which they belong, and the data source from which they are derived.

**S2 Table.**

**Full Table of preprocessed sequence-derived features.** Includes labels from HGNC/Uniprot/Ensembl to merge with other datasets. Most features are preprocessed by corresponding mRNA/amino acid length.

**S3 Data**

**Archive of supporting code to generate sequence-derived features.** Includes code in Jupyter Notebook format. To read this format, we recommend you install Anaconda (for Python). See instructions for how to achieve this on our Github repository where these notebooks also exist.

**S4 Data**

**Code scripts associated with S3 Data.** These scripts, such as accessing BioMart information are not essential to running the notebooks, but cover groundwork on how to access meta information.

**S5 Table**

**Gene Ontology Analysis of genes with each GO term as modelled by cell line and input type.** Machine learning models are constructed by filtering for genes that match with the appropriate GO term, and models use protein abundance per gene as the response variable. Input types ‘SDF (alone)’ refer to all sequence-derived features, ‘mRNA (alone)’ uses only the corresponding mRNA abundance, and ‘SDF plus mRNA’ is a combination of both.

