## Supplementary material for "Predicting Relative Protein Abundance via Sequence-Based Information": S3 Data: NCBI-Access.ipynb

```
# %% [markdown]
## Accessing NCBI GenBank data

We have downloaded the entire **NCBI** repository for human mRNA transcripts. We have already obtained protein information from Uniprot and thus this does not interest us. 

For best use, run parts of this notebook in Linux Ubuntu as we regularly dip into command line/terminal using iPython Magicks.

We made use of the script (accessed 17 Apr 2019):

    wget ftp://ftp.ncbi.nlm.nih.gov/refseq/H_sapiens/mRNA_Prot/human.*.rna.gbff.gz
   
Which retrieved all the mRNA transcripts in genbank format to `../data/NCBI/`. We are interested in mRNAs that are curated, so we will ignore any beginning with **XM_** or **XR_**.

We shall be making use of **BIOPYTHON** package to process our collated GenBank files into coherent objects.

BIOPYTHON can be downloaded using PyPI or through conda, using the command: 
    
    pip install biopython

in your local terminal.

# %% [code]
from Bio import SeqIO
from Bio.SeqRecord import SeqRecord
from pathlib import Path

import numpy as np
import pandas as pd
import itertools as it
import os
import re
import sys
sys.path.insert(0, "../../scripts/")

from caching import cache, load_or_execute_csv
import misc_functions as misc

path = "../../data/NCBI/"

# %% [markdown]
### Extracting Gene Accessions from NCBI

Found in the `gene2accession.gz` file in `ftp.ncbi.nlm.nih.gov/gene/DATA/`.

We keep:

1. All Homo sapien (Human) records.
2. All REVIEWED or VALIDATED genes, no experimental ones.

# %% [code]
def download_and_preprocess_human_accession():
    url = "ftp://ftp.ncbi.nlm.nih.gov/gene/DATA/gene2accession.gz"
    # check if file is there first, if not download.
    if not os.path.isfile("gene2accession.gz"):
        # download file using bash
        _ = os.system("wget %s" % url)
    # now open this file and preprocess
    dchunks = pd.read_csv("gene2accession.gz", sep="\t", iterator=True, 
                          chunksize=200000, na_values=["-", "?"], low_memory=False)
    
    accessions = pd.concat([
        (x[(x["#tax_id"] == 9606) & (x["status"].isin(["REVIEWED","VALIDATED"]))]) for x in dchunks
    ])
    # move the accession file to it's proper directory.
    os.system("mv gene2accession.gz ../../data/NCBI/")
    # return
    return accessions

# %% [code]
gene_accession = load_or_execute_csv(path+"gene2accession.csv",
                                     download_and_preprocess_human_accession)

# %% [markdown]
### Extracting Gene Groups, Gene Info from NCBI

Found in the `gene_group.gz`, `gene_info.gz`, `gene_neighbors.gz` file in `ftp.ncbi.nlm.nih.gov/gene/DATA/`.

We keep:

1. All Homo sapien (Human) records.
2. All REVIEWED or VALIDATED genes, no experimental ones.

# %% [code]
def download_and_preprocess_human_genegroup():
    if not os.path.isfile("gene_group.gz"):
        # download file using bash
        _ = os.system("wget ftp://ftp.ncbi.nlm.nih.gov/gene/DATA/gene_group.gz")
    # now open this file and preprocess
    dchunks = pd.read_csv("gene_group.gz", sep="\t", iterator=True, 
                          chunksize=100000, na_values=["-", "?"], low_memory=False)
    groups = pd.concat([
        (x[(x["#tax_id"] == 9606)]) for x in dchunks
    ])
    # move the accession file to it's proper directory.
    os.system("mv gene_group.gz ../../data/NCBI/")
    # return
    return groups

# %% [code]
gene_groups = load_or_execute_csv(path+"gene_groups.csv",
                                     download_and_preprocess_human_genegroup)

# %% [code]
def download_and_preprocess_human_geneinfo():
    if not os.path.isfile("gene_info.gz"):
        # download file using bash
        _ = os.system("wget ftp://ftp.ncbi.nlm.nih.gov/gene/DATA/gene_info.gz")
    # now open this file and preprocess
    dchunks = pd.read_csv("gene_info.gz", sep="\t", iterator=True, 
                          chunksize=100000, na_values=["-", "?"], low_memory=False)
    groups = pd.concat([
        (x[(x["#tax_id"] == 9606)]) for x in dchunks
    ])
    # move the accession file to it's proper directory.
    os.system("mv gene_info.gz ../../data/NCBI/")
    # return
    return groups

# %% [code]
gene_info = load_or_execute_csv(path+"gene_info.csv",
                                     download_and_preprocess_human_geneinfo)

# %% [code]
def download_and_preprocess_human_geneneighbors():
    if not os.path.isfile("gene_neighbors.gz"):
        # download file using bash
        _ = os.system("wget ftp://ftp.ncbi.nlm.nih.gov/gene/DATA/gene_neighbors.gz")
    # now open this file and preprocess
    dchunks = pd.read_csv("gene_neighbors.gz", sep="\t", iterator=True, 
                          chunksize=200000, na_values=["-", "?"], low_memory=False)
    groups = pd.concat([
        (x[(x["#tax_id"] == 9606)]) for x in dchunks
    ])
    # move the accession file to it's proper directory.
    os.system("mv gene_neighbors.gz ../../data/NCBI/")
    # return
    return groups

# %% [code]
gene_neighbors = load_or_execute_csv(path+"gene_neighbors.csv",
                                     download_and_preprocess_human_geneneighbors)

# %% [markdown]
## Download RNA sequences (FASTA) from NCBI

We follow this sequence of events:

1. Download files
2. Move to directory
3. Unzip
4. Extract genes that are not experimental (i.e beginning with XM/XR)

**NCBI** uses a distinct nomenclature with naming their RNA files with a number of beginnings:
- **NM_**: mRNA protein-coding transcripts (curated)
- **NR_**: non-protein coding transcripts
- **XM_**: predicted model protein-coding
- **XR_**: predicted model non-protein coding

We will eliminated any predicted-model transcripts as we want to work only with solid curated data.

# %% [code]
%%bash
# fetch
wget ftp://ftp.ncbi.nlm.nih.gov/refseq/H_sapiens/mRNA_Prot/human.*.rna.fna.gz && \
mv human.*.rna.fna.gz ../../data/NCBI/genbank/ && \
gunzip ../../data/NCBI/genbank/human.*.rna.fna.gz && \
cat ../../data/NCBI/genbank/human.*.rna.fna | tr -d '\n' | tr '>' '\n' | grep -iv "X._" > ../../data/NCBI/genbank/ncbi_fasta.fna && \
rm ../../data/NCBI/genbank/human.*.rna.fna

# %% [markdown]
## Download RNA sequences (Genbank) from NCBI

# %% [code]
%%bash
wget ftp://ftp.ncbi.nlm.nih.gov/refseq/H_sapiens/mRNA_Prot/human.*.rna.gbff.gz && \
mv human.*.rna.gbff.gz ../../data/NCBI/genbank/

# %% [markdown]
## Preprocessing into tabular format

Now that we have filtered for **XR_** and **XM_** terms, we want to rearrange each GenBank file by mRNA transcript (rows), this means quantifying the feature table into relevant count data etc.

We need to think about the features we care about from this data. Generic features include:
- The total number of features in the sequence feature table

### Table 1: FEATURE COUNT MATRIX

$$
F=\begin{bmatrix}
f_{11} & f_{12} & \dots & f_{1m} \\
f_{21} & f_{22} & \dots & f_{2m} \\
\vdots & \vdots & \ddots & \vdots \\
f_{n1} & f_{n2} & \dots & f_{nm}
\end{bmatrix}, n=\text{transcripts}, m=\text{features}
$$

Extracting the unique labels in the feature matrix found in all of the curated RNA sets.

Here we go through every mRNA transcript, and count the occurences of feature names and collate this into a matrix, whereby rows represent unique feature names, and columns represent mRNA transcripts.

# %% [code]
# read in curated rna - extract records
def extract_feature_matrix(filename):
    
    print("[Processing file {}]".format(filename))
    # extract records once from file. expensive.
    records = [record for record in SeqIO.parse(open(filename,"r"), "genbank")]
    # extract value_counts.
    value_counts = [(pd.Series([record.features[i].type for i in range(len(record.features))])
                       .value_counts()) for record in records]
    # concatenate all the genes in this set into a 'frequency table' to be appended to list.
    freq_table = pd.concat(value_counts, axis=1, sort=False)
    # append names to the frequency table columns.
    freq_table.columns = [record.name for record in records]
    # delete records list.
    del records
    del value_counts
    
    return freq_table

# 1. extract the unique features names, and stick them into a massive dataframe
freq_frame = pd.concat([extract_feature_matrix(f) for f in files_and_path], axis=1, sort=False).T

# %% [code]
def process_and_remove_experimental(files):
    for f in files:
        print_line = "[File: {}".format(f)
        # using itertools, filter the generator without creating the list
        seq_file = SeqIO.parse(open(f,'r'), 'genbank')
        reduced_records = it.filterfalse(lambda r: (r.name.startswith('XM') or r.name.startswith('XR')), seq_file)
        # if records length > 0, save them to a file
        # print_line += ", n={}".format(len(reduced_records))
        # create a new file name
        regex_num = re.findall("human.([0-9]+).rna.gbff", f)
        if len(regex_num) > 0:
            new_file = "../data/NCBI/curated_{}.gbff".format(regex_num[0])
            print_line += ", renamed: '{}']".format(new_file)
            with open(new_file,"w") as nf:
                SeqIO.write(reduced_records, nf, 'genbank')
        print(print_line)
        # remove that file
        os.remove(f)

# %% [code]
files_in_dir = [path+p for p in os.listdir(path) if p.startswith("human")]
files_in_dir

# %% [code]
process_and_remove_experimental(files_in_dir)

# %% [markdown]
## Re-selecting filtered files

# %% [code]
files_and_path = [path+p for p in os.listdir(path) if p.startswith('curate')]
files_and_path

# %% [code]
freq_frame.head()

# %% [code]
ids = [d.split(" ")[0].split("_")[0] for d in data]

# %% [markdown]
### Saving The Feature Matrix in ../data/NCBI:

# %% [code]
data[:10]

# %% [code]
pd.Series(ids).value_counts()

# %% [code]
freq_frame.to_csv(path+"rna_feature_matrix.csv", sep="\t", na_rep=np.nan)

# %% [markdown]
## Table 2: SEQUENCE TABLE

Here we generate a table that only has the accession, mRNA sequence and description. This is because the text file will be large and hence we wish to reduce the size of the table.

$$
S=\begin{bmatrix}
s_1 \\ s_2 \\ \vdots \\ s_n
\end{bmatrix}
$$

Where $s$ refers to the primary information of use in the table; the primary mRNA transcript. $n$ is the number of transcripts.

For protein-coding sequences, we extract features from the CDS such as the start and end range.

This will create a massive file - be warned!

# %% [code]
def extract_sequence_table(filename):
    
    print("Processing file {}".format(filename))
    # save generator into object.
    records = [r for r in SeqIO.parse(open(filename,"r"), "genbank")]
    rnas = pd.concat([pd.Series([r.id, r.description, str(r.seq)], index=["id","desc","seq"]) 
                      for r in records], axis=1).T.set_index("id")
    cdss = (pd.concat(list(itertools.chain.from_iterable(
        [[pd.DataFrame.from_dict(f.qualifiers, orient="index").T.assign(id=r.id) for f in r.features if f.type=="CDS"]
         for r in records]
    )), axis=0, sort=False).dropna(thresh=3)
        .drop(["product","note","gene_synonym","gene","db_xref"], axis=1).set_index("id"))
    start_end = pd.DataFrame(list(itertools.chain.from_iterable(
        [[[r.id, f.location.start.position,f.location.end.position,f.location.strand] 
          for f in r.features if f.type == "CDS"] 
             for r in records])), columns=["id","cds_start", "cds_end", "strand"]).set_index("id")

    # merge together
    return rnas.join(cdss).join(start_end)

# %% [code]
# 2. save sequence data into a table and concatenate them all together.
seq_table = pd.concat([extract_sequence_table(f) for f in files_and_path], axis=0, sort=False)
seq_table.head(3)

# %% [code]
# save to file
seq_table.to_csv(path+"rna_sequence_table.csv", sep="\t", na_rep=np.nan)

# %% [markdown]
## 3. Coding sequences as FASTA file

There are numerous inputs for proprietary software that desires the coding sequence (CDS) of an mRNA transcript to function. Thus we will save every coding sequence ONLY in FASTA;

# %% [code]
def generate_cds_fasta(filename, output_file):
    
    print("Processing CDS seq: {}".format(filename))
    
    records = [record for record in SeqIO.parse(open(filename,"r"), "genbank") \
                   if (record.name.startswith("NM_") and record.id != "")]
    
    # collect features
    features = [[[f.location.start.position, f.location.end.position] 
             for f in record.features if f.type == "CDS"] for record in records]
    # collect sequences and cut to CDS
    sequences = [record.seq[features[i][0][0]:features[i][0][1]] for i,record in enumerate(records)]
    # check that all sequences are appropriate length and begin with ATG
    try:
        # assert that all of the sequences are modded into 3.
        assert all([len(s)%3==0 for s in sequences])
        # check to make sure each one begins with ATG - methionine
        assert all([s[:3]=="ATG" for s in sequences])
    except AssertionError:
        print("\t[{}/{} seq no mod into 3;{}/{} seq no begin ATG.]".format(
            sum([len(s)%3!=0 for s in sequences]), len(sequences),
            sum([s[:3]!="ATG" for s in sequences]), len(sequences)
        ))
    new_records = [SeqRecord(sequences[i], id=records[i].id, name=records[i].name, description=records[i].description) \
                   for i,record in enumerate(records)]
    # free up space
    del features
    del sequences
    del records
    
    return new_records

# %% [code]
# run and write to disk all
fasta_records = [generate_cds_fasta(f,"cds_cache.fasta") for f in files_and_path]

# write to disk
with open(path+"cds_cache.fasta","a+") as h:
    for fasta in fasta_records:
        SeqIO.write(fasta, h, "fasta")

# %% [markdown]
### FINALLY

Compress any genbank files so as to save disk space. Wrap them together in a `.tar.gz` compartment.

# %% [code]
%%bash
cd ../../data/NCBI/
tar -zcvf refseq_archive.tar.gz *.gbff

# %% [code]
print("complete")

# %% [code]
!rm {path}*.gbff
```
