## Supplementary material for "Predicting Relative Protein Abundance via Sequence-Based Information": S3 Data: Uniprot-Access.ipynb

```
# %% [markdown]
# Downloading Uniprot

We go to https://www.uniprot.org/, type in `Homo sapiens` into the search bar. We filter by `Swissprot`, which is the manually curated Uniprot sequences, and filter by `Homo sapiens`, and by Proteomes `UP000005640`. The columns we select to download are:

- [Entry]
- Proteomes
- HGNC
- CCDS
- Refseq
- BioGrid
- Ensembl
- STRING
- PDB
- Features
- Keywords
- Interacts with
- Length
- Mass
- Sequence

Download the file as `Tab-separated`, move to `../data/Uniprot/`, rename to `uniprot_set` and un-gz.

# %% [code]
_pwd = %pwd
_root = "/home/gparkes/Documents/Research Data"
_local_r = "".join(["../" for i in range(_pwd.count("/") - _root.count("/"))])

# %% [markdown]
### Collect File

We download the file in tab-delimited format to make it easy to parse. The entire file is 80MB in size, which in memory translates to roughly 150MB, which is hefty, but this includes ALL of the possible protein information we could care about.

We compress the original file back into .tar.gz to save space. This can be found in the `/data/Uniprot/` folder.

# %% [code]
import pandas as pd
prot = pd.read_table(_local_r + "data/Uniprot/uniprot_set.tab", sep="\t")

# %% [code]
print(prot.shape)
print("{:0.2f}MB".format(prot.memory_usage(deep=True).sum() /1000000))
prot.head(3)

# %% [markdown]
## Use Regexx for column-name preprocessing

Includes renaming cross-reference, and column names.

# %% [code]
import sys
sys.path.insert(1, _local_r+"scripts/")
import regexx

# %% [code]
new_cols = regexx.replace_chain(prot.columns, [("Cross-reference \(", ""), ("\)","_ID"), ("Entry","Uniprot_ID"),
                                    (" transcript", "_ID"), ("s with", "ions"), ("Sequence","Seq")])

# %% [markdown]
## Simplify Column Names

Firstly, we will replace spaces with underscores to make programming with it easier, then replace cross-references with shorthand DB for database.

# %% [code]
# replace column names with the dictionary that maps old col names to new col names
prot.rename(columns=dict(zip(prot.columns, new_cols)), inplace=True)
prot.columns

# %% [markdown]
### Save File as initial dataset to be cleaned


# %% [code]
import numpy as np
prot.to_csv(_local_r + "data/Uniprot/uniprot_all.csv", sep="\t", index=False, na_rep=np.nan)
```
