## Supplementary material for "Predicting Relative Protein Abundance via Sequence-Based Information": S3 Data: environment.yml

```
name: cycle
channels:
  - defaults
  - conda-forge
dependencies:
  - python>=3.7
  - numpy>=1.11
  - scipy>=1.3
  - joblib>=0.14.1
  - jupyter>=1.0.0
  - pandas>=0.25
  - matplotlib>=3.1.0
  - scikit-learn>=0.22
  - pip>=19.0.3
  - seaborn>=0.11.0
  - xlrd>=1.1.0
  - biopython>=1.7
  - numba>=0.48.0
  - networkx>=2.0
  - bokeh>=1.4.0
  - statsmodels
  - pip:
    - geoparse==1.2.0
    - matplotlib-venn==0.11.5
    - goatools==1.0.6
    - pingouin
    - pyitlib
    - toolz
    - turbopanda>=0.2.8
prefix: /home/gparkes/anaconda3
```
