## Supplementary material for "Predicting Relative Protein Abundance via Sequence-Based Information": S3 Data: Aviner-Microarray.ipynb

```
# %% [markdown]
# Microarray Dataset

We are going to be working with the microarray data that Aviner uses in his 2015 paper. However we are drawing directly from the source instead.

What's particularly exciting is the fact that this microarray data has steps at **0h, 2h, 4h, 6h, 8h** and **12h**. It describes the 0-state as *blocked*. The data already comes as *RMA-normalized*, hence we don't have to worry about performing our own transformations on the data itself.

$$
\text{G1 -> 2h, S -> 8h, G2/M -> 12h}
$$

To do this, we need to be able to parse the results from Gene Expression Omnibus (GEO); GSE26922. We will be using:
    
    GEOparse
    
Which is a python library which exists solely for this purpose. We use `pip install GEOparse` to obtain it. We follow the guide from https://geoparse.readthedocs.io/en/latest/usage.html.

# %% [code]
import GEOparse
from pathlib import Path
import pandas as pd
import numpy as np
import matplotlib.pyplot as plt
%matplotlib inline

import sys
sys.path.insert(0, "../../scripts/")
sys.path.insert(0, "../../../turbopanda/")
import missingno as msno
import turbopanda as turb

import unique_labels as privID

# %% [markdown]
### Downloading and saving the dataset

GEOparse lets us import directly from the website with an internet connection.

We will save it in the data folder.

# %% [code]
def fetch_microarray(file_name):
    # given a file name, either read in the file from local or download.
    # assume file name does not include compressed extension.
    micro_p = Path(file_name+".gz")
    # assumes is a SOFT file
    if micro_p.is_file():
        gse = GEOparse.get_GEO(filepath=file_name+".gz")
    else:
        # download from GEO
        gse = GEOparse.get_GEO("GSE26922", destdir="../data/microarray/")
        gse.to_soft(file_name)
        !gzip {file_name}
    # return the in-memory gse
    return gse

micro_file = "../../data/microarray/GSE26922_family.soft"
gse = fetch_microarray(micro_file)

# %% [markdown]
### Merging together expression levels

We will merge the 18 samples into one larger dataframe, which should be easy.

# %% [code]
# create a list of table results
tables = [gse.gsms[x].table for x in gse.gsms.keys()]
# initialise merging
mergeset = pd.merge(tables[0], tables[1], on="ID_REF", how="outer", validate="one_to_one")
for i in range(2, len(tables)):
    # iterate over mini tables, and merge on id, keeping all records through UNION.
    mergeset = pd.merge(mergeset, tables[i], on="ID_REF", how="outer", validate="one_to_one")
# change column names by updating with meta-data
mergeset.columns = ["ID_REF"] + [gse.gsms[x].metadata["description"][0] for x in gse.gsms.keys()]

# %% [code]
print(mergeset.shape)
print("{:0.2f}MB".format(mergeset.memory_usage(deep=True).sum() /1000000))
mergeset.tail()

# %% [markdown]
### Merging GPL data (metadata) in with expression

We need the labels from GPL so that later on we can map the Ensembl, Genbank or Refseq labels to something in our combined aviner set.

Here we map together the metadata provided in **GPL** with the expression set data found in each **GSM** which we have already merged together.

For the GPL each column represents as follows:

- 'GB_LIST': 'GenBank and RefSeq Accessions from mrna_assignment column LINK_PRE:"http://www.ncbi.nlm.nih.gov/nuccore/?term=" DELIMIT:","',
- 'ID': 'transcript_cluster_id LINK_PRE:"https://www.affymetrix.com/LinkServlet?array=HuGene-1_0-st-v1&probeset="',
- 'RANGE_GB': 'NCBI RefSeq for chromosome of current build',
- 'RANGE_START': 'start  (integer). Coordinates are standard 1-based (length=stop-start+1).',
- 'RANGE_STOP': 'stop  (integer). Coordinates are standard 1-based (length=stop-start+1).',
- 'RANGE_STRAND': 'strand  (+|-)',
- 'SPOT_ID': 'genomic location of the transcript cluster in the version of the genome assembly used at annotation time. Coordinates are standard 1-based (length=stop-start+1).',
- 'category': 'Array design category of the transcript cluster',
- 'gene_assignment': 'Gene information for each assigned mRNA for mRNAs that corresponds to known genes (multipart).',
- 'mrna_assignment': 'Description of the public mRNAs that should be detected by the sets within this transcript cluster based on sequence alignment (multipart).',
- 'seqname': 'chromosome number',
- 'total_probes': 'Total number of probes contained by this transcript cluster.'}}

# %% [code]
# when we merge, we only accept a column if the ID is in both, otherwise it is useless to us!
full_set = pd.merge(gse.gpls["GPL6244"].table.rename(columns={"ID":"ID_REF"}), 
         mergeset,
         on="ID_REF",
        # must be in both!
         how="inner", 
         validate="one_to_one")
print("{:0.2f}MB".format(full_set.memory_usage(deep=True).sum() /1000000))
print(full_set.shape)
full_set.head(3)

# %% [markdown]
### Dropping unnecessary columns

We do not need information regarding ranges, probes nor SPOT_ID.

# %% [code]
full_set.drop([
    "SPOT_ID", "seqname","RANGE_GB","RANGE_STRAND","RANGE_START","RANGE_STOP","total_probes","category"
], axis=1, inplace=True)

# %% [code]
print("{:0.2f}MB".format(full_set.memory_usage(deep=True).sum() /1000000))
print(full_set.shape)
full_set.head(3)

# %% [markdown]
Gene names follow the pattern:

$$
4p+1
$$

# %% [code]
exp_set = full_set.set_index("ID_REF")["gene_assignment"].str.split(" // ", expand=True)

# %% [code]
gene_names = exp_set[list(range(1, exp_set.shape[1], 4))]

# %% [code]
gene_names_uniq = gene_names.apply(lambda x: ";".join(x.dropna().unique()), axis=1).replace("",np.nan)

# %% [code]
gene_names_uniq.head()

# %% [markdown]
### Extracting labels from assignment groups

We have three columns that contain label ID data:

- GB_LIST
- mrna_assignment
- gene_assignment

Extract the transcript labels from each one, and remove duplicated columns.

# %% [code]
genes_ass = full_set.set_index("ID_REF").gene_assignment.str.extractall("((?:ENST|NM_|NR_)[0-9]+)")
mrna_ass = full_set.set_index("ID_REF").mrna_assignment.str.extractall("((?:ENST|NR_|NM_)[0-9]+)")
gb_ass = full_set.set_index("ID_REF").GB_LIST.str.extractall("((?:NC|NR|NM|NG)_?[0-9]+)")

# %% [code]
label_concat = pd.concat([gb_ass, mrna_ass, genes_ass], axis=1)
label_concat.columns = ["GB_LIST","mRNA","genes"]
print(label_concat.shape)
label_concat.head()

# %% [markdown]
### Drop duplicates

# %% [code]
label_concat.drop_duplicates(inplace=True)

# %% [markdown]
### Extract unique labels per mRNA

We create a dictionary mapping **ID_REF** to the unique terms.

# %% [code]
stacked_lbls = {x:";".join(label_concat.loc[x].stack().unique()) for x in label_concat.index.get_level_values(0).unique()}

# %% [code]
full_set = full_set.merge(pd.Series(stacked_lbls).to_frame("mRNA_ID"), 
                          left_on="ID_REF", right_index=True, how="left")
print(full_set.shape)
full_set.head(3)

# %% [markdown]
### Include isolated gene names

# %% [code]
full_set = full_set.merge(gene_names_uniq.to_frame("gene_names"), on="ID_REF", how="left")

# %% [markdown]
### Drop existing GB_LISTs

# %% [code]
full_set.drop(["GB_LIST","gene_assignment","mrna_assignment","ID_REF"], axis=1, inplace=True)

# %% [markdown]
### Drop rows with no ID available

If we haven't extracted a label from either GB_LIST, mRNA_assignment or gene_assignment we drop it.

# %% [code]
full_set.dropna(subset=["mRNA_ID"], inplace=True)

# %% [code]
print(full_set.shape)
print("{:0.2f}MB".format(full_set.memory_usage(deep=True).sum() /1000000))
full_set.head(3)

# %% [markdown]
### Saving microarray to dataset

We will save it as: `../datasets/microarray_proc.csv`.

# %% [code]
full_set.to_csv("../../datasets/microarray_proc.csv")

# %% [markdown]
## Further preprocessing

To make this machine-learning ready and easy to integrate with other datasets, we'll convert this microarray file into a longer form.

# %% [code]
full_set

# %% [code]
rna_pre = turb.MetaPanda(full_set, name="aviner_rna")

# %% [code]
rna_pre

# %% [markdown]
#### Expand out mRNA ID

# %% [code]
rna_pre.expand("mRNA_ID",";")

# %% [code]
rna_pre.df_.head()

# %% [code]
rna_pre.write("../../datasets/clean/mRNA_aviner_adv.csv",with_meta=False)

# %% [markdown]
#### Melt columns and re-assign

# %% [code]
# melt no longer exists. 
rna_pre.apply('melt', id_vars=["mRNA_ID","gene_names"],
              var_name="col_name",value_name="mRNA_level")


# %% [markdown]
#### Generate categories

# %% [code]
rna_pre.split_categories("col_name", "_", ("rlabel","ts","repl"))

# %% [markdown]
#### Pivot table

# %% [code]
rna_pre.apply("pivot_table", "mRNA_level", ["counter","mRNA_ID","ts"], "repl")
rna_pre.apply("mean", axis=1)
rna_pre.apply("unstack","ts")

# %% [code]
rna_pre.apply("pipe", turb.pipe.add_prefix, "mRNA_")

# %% [code]
rna_pre.apply("reset_index")

# %% [code]
rna_pre.head()

# %% [markdown]
#### Convert ts to `timeseries` format

# %% [code]
rna_pre.write("../../datasets/clean/mRNA_aviner_means.csv",with_meta=False)
```
