## Supplementary material for "Predicting Relative Protein Abundance via Sequence-Based Information": S3 Data: HGNC-Preprocessing.ipynb

```
# %% [markdown]
# HGNC Preprocessing

We need to apply some basic modifications to the HGNC files we receive. This is such that we have a consistent *current standard* of nomenclature for the Gene names we work with.

# %% [code]
import sys
import numpy as np
import json
import pandas as pd
sys.path.insert(0, "../../../turbopanda/")
# our main import
import turbopanda as turb

# %% [code]
hgnc = turb.read("../../data/HGNC/HGNC_gene.csv", name="HGNC", sep="\t")
hgnc

# %% [code]
hgnc.head()

# %% [markdown]
## Drop irrelevant HGNC columns

# %% [code]
hgnc.drop("date","vega","mgd","orpha","iupha","homeo",
          "pubmed","rgd","cosm","^location$","omim")

# %% [markdown]
## Simplify column names

# %% [code]
hgnc.rename_axis([("hgnc_",""),("_sortable",""),("_symbol",""), ("gene_",""),
             ("accession","id"), ("ids","id")])

# %% [markdown]
## Create a dictionary map to replace alias/previous symbols with current

The `alias` column contains significantly more information about the name of a gene. We can create a dictionary that maps:

    alias_i : current
    current : current
    
Such that when we apply `.map()` to any Gene Symbol ID column, we will get the fresh/current HGNC-approved gene name for any given dataset.

# %% [code]
temp_map = hgnc.df_.set_index("symbol")["alias"].dropna().str.split("|").explode()
temp_map.head()

# %% [markdown]
Now append on the *current gene name* as `current : current` mapping:

# %% [markdown]
Now we swap them around by creating a new Series where the index are the values, and values are the index:

# %% [code]
# swap them around and turn it into a dict
alias_to_current = pd.concat([
    pd.Series(temp_map.index, index=temp_map.values),
    pd.Series(data=hgnc.df_["symbol"].values, index=hgnc.df_["symbol"].values)
]).to_dict()

# %% [markdown]
## Store as a JSON file

# %% [code]
# store alias as json file
with open("../../data/HGNC/hgnc_alias.json", "w") as f:
    json.dump(alias_to_current, f, allow_nan=False, sort_keys=True)

# %% [markdown]
## Store HGNC dataset as *clean* version

# %% [code]
hgnc.write("../../data/HGNC/HGNC_clean.csv", with_meta=False)
```
