## Supplementary material for "Predicting Relative Protein Abundance via Sequence-Based Information": S3 Data: Lundberg-RNA-Protein.ipynb

```
# %% [markdown]
## RNA-Protein Preprocessing with Lundberg dataset

In a paper entitled: **Defining the transcriptome and proteome in three functionally different human cell lines**, which can be found [here](https://www.ncbi.nlm.nih.gov/pubmed/21179022):

### Abstract

An essential question in human biology is how cells and tissues differ in gene and protein expression and how these differences delineate specific biological function. Here, we have performed a global analysis of both mRNA and protein levels based on sequence-based transcriptome analysis (RNA-seq), SILAC-based mass spectrometry analysis and antibody-based confocal microscopy. The study was performed in three functionally different human cell lines and based on the global analysis, we estimated the fractions of mRNA and protein that are cell specific or expressed at similar/different levels in the cell lines. A highly ubiquitous RNA expression was found with >60% of the gene products detected in all cells. The changes of mRNA and protein levels in the cell lines using SILAC and RNA ratios show high correlations, even though the genome-wide dynamic range is substantially higher for the proteins as compared with the transcripts. Large general differences in abundance for proteins from various functional classes are observed and, in general, the cell-type specific proteins are low abundant and highly enriched for cell-surface proteins. Thus, this study shows a path to characterize the transcriptome and proteome in human cells from different origins.

The following datasets are extracted from the supplementary materials of this paper.

### Different Cell Lines

- **U2OS**: Bone tissue, moderately differentiated sarcoma
- **U251MG**: Human Brain, malignant glioblastoma tumour
- **A431**: Human Skin (epidermis), epidermoid carcinoma

# %% [code]
import numpy as np
import pandas as pd
import matplotlib.pyplot as plt
import sys
sys.path.insert(0,"../../../turbopanda")
import turbopanda as turb
%matplotlib inline

# %% [markdown]
### Lundberg MS (Protein)

This file has been slightly modification since we found `datetime` elements mixed in.

These values are identified by the appearance of `-` and `,`, which we select for.

# %% [code]
tds = turb.read("../../data/lundberg_2010_MS.xls",
                name="Lundberg_MS",
                usecols=['ENSG-ID', 'INTENSITY U2OS', 'INTENSITY U251MG', 'INTENSITY A431'],
                na_values=['NA', 0])

# %% [markdown]
#### Removing Datetime stamps from the blocks, and replacing with NA

Otherwise MetaPanda breaks... apparently

# %% [code]
scrub_dates = turb.Pipe(['transform', lambda x: np.nan if type(x)==str else x, 
                        'selector=is_mixed_type','method=applymap'],
                        ['apply', 'transform', pd.to_numeric, 'errors=ignore', 'downcast=unsigned'])

# %% [code]
tds.compute(scrub_dates, inplace=True, update_meta=True)

# %% [markdown]
#### Preprocess

# %% [code]
preproc_pipe = turb.Pipe(['apply','dropna','how=all'],
                         ['transform', np.log2, 'selector=INTENSITY', 'whole=True'])

# %% [code]
preproc_pipe.p

# %% [code]
tds.compute(preproc_pipe, inplace=True)

# %% [markdown]
#### Saving to file

# %% [code]
tds.write("../../datasets/lundberg_prot.csv")

# %% [markdown]
### Lundberg RNA-Seq (RPKM)

This file has been slightly modification since we found `datetime` elements mixed in. We removed these manually.

# %% [code]
rds = turb.read("../../data/lundberg_2010_RNAseq.xls",
                name='Lundberg_RPKM',
                usecols=['ENSG-ID', 'RPKM U2OS', 'RPKM U251MG', 'RPKM A431'],
                na_values=[0])

# %% [markdown]
#### Removing Datetime stamps from the blocks, and replacing with NA

# %% [code]
rds.compute(scrub_dates, inplace=True, update_meta=True)

# %% [markdown]
#### Preprocess

# %% [code]
preproc_pipe = turb.Pipe(['apply','dropna','how=all'],
                         ['transform', np.log2, 'selector=RPKM', 'whole=True'])
preproc_pipe.p

# %% [code]
rds.compute(preproc_pipe, inplace=True)

# %% [markdown]
#### Saving to file

# %% [code]
rds.write("../../datasets/lundberg_rna.csv")

# %% [markdown]
## Merge together and preprocess

# %% [code]
lundberg = turb.merge([tds, rds])
lundberg

# %% [code]
lundberg.write("../../datasets/lundberg_all.csv")

# %% [markdown]
## Sanity and Interest Plots

# %% [code]
fig, axes = plt.subplots(ncols=3, figsize=(16,4))
data_pairs = [("RPKM_U2OS","INTENSITY_U2OS", axes[0]),
              ("RPKM_U251MG", "INTENSITY_U251MG", axes[1]),
              ("RPKM_A431", "INTENSITY_A431", axes[2])]

for rpkm, ms, ax in data_pairs:
    x = lundberg[rpkm]
    y = lundberg[ms]
    corr = turb.correlate(x, y, method='pearson')
    ax.scatter(x, y, marker='x', label="pearson: {:0.3f}, n={}".format(corr['r'], corr['n']))
    ax.legend()
    ax.set_xlabel(r"$log_2$ RPKM")
    ax.set_ylabel(r"$log_2$ Intensity (MS)")
    ax.set_title(rpkm.split(" ")[-1].upper())
```
