## Supplementary material for "Predicting Relative Protein Abundance via Sequence-Based Information": S3 Data: NCBI-Preprocessing.ipynb

```
# %% [markdown]
# NCBI Preprocessing

In this notebook we are going to derive the sequence-based features from the mRNA sequence, the mRNA coding region and the translation amino acid sequence provided to us by RefSeq.

There are a whole plethora of features we may wish to extract from the underlying mRNA sequence. Of particular interest will be what we can make from upstream 5'-UTR and 3'-UTR features.

According to *NCBI-Access*, we generated a number of different file types that will be useful:
1. A feature matrix which has counted the occurance of every feature.
2. The raw mRNA sequences from each sample.
3. The variations within the mRNA sequence (SNP).
4. The coding region for such mRNAs in FASTA.
5. mRNA source information which contains interesting metadata.

# %% [code]
import pandas as pd
import numpy as np
import matplotlib.pyplot as plt
%matplotlib inline
import os
import re
import time
from itertools import repeat as my_rep

import Bio
from Bio import SeqIO, Seq
from Bio.SeqRecord import SeqRecord
from Bio.SeqUtils import GC

import sys
sys.path.insert(0, "../../scripts/")
sys.path.insert(0, "../../../turbopanda/")
import turbopanda as turb
from caching import cache, load_or_execute_csv
import misc_functions as misc

path = "../../data/NCBI/"

# %% [markdown]
## Filtering out descriptive features

This is an important and crucial step, we want to extract as many features as possible from both **coding** and **non-coding RNAs**.

# %% [code]
super_seq = turb.read(path+"rna_sequence_table.csv",sep="\t")
seqs = super_seq.df_

# %% [code]
turb.missing(seqs)
plt.show()

# %% [markdown]
### Drop GO components, exception, slippage

# %% [code]
super_seq.drop("GO","except","riboso","EC")

# %% [markdown]
### Eliminate duplicate rows

1. Create Refseq ID by chopping off the end index number.
2. Drop duplicates in Refseq ID
3. Re-merge back into dataset using 'inner'

# %% [code]
seqs.drop_duplicates(subset=["id"], keep="first", inplace=True)

# %% [code]
seqs["Refseq"] = seqs["id"].str.extract("(.+).[1-9]").apply(lambda x: x.str.strip())

# %% [code]
super_seq.drop("^id$")

# %% [markdown]
### Sort IDs by coding, noncoding

# %% [code]
seqs["is_coding"] = seqs["protein_id"].notnull()

# %% [markdown]
## Calculating mononucleotide counts

There are a number of full-mRNA features we may be interested in, such as the frequency of each base with respect to it's length.

- Counting the number of A, T, C and G
- Length of the full mRNA sequence

# %% [code]
seqs.head(2)

# %% [code]
for s in "GACT":
    seqs["%s_mrna" % s] = seqs.seq.str.count(s)

seqs["length_mrna"] = seqs.seq.str.len()
seqs["GC_content_mrna"] = (seqs.seq.str.count("G") + seqs.seq.str.count("C")) / seqs["length_mrna"]

# %% [code]
seqs.head(2)

# %% [markdown]
## Break down sequence into 3-UTR, CDS and 5-UTR

For our genes that begin with **NM\_** i.e the coding genes, we are interested in breaking down the sequence into three parts:
- 5'-UTR, the region preceding the coding region.
- CDS, the coding sequence
- 3'-UTR, the region following the coding region.

We are going to fetch directly from refseq genbank file, since this provides rich information including a translation encoding to match it to.

# %% [code]
# we extract coding regions, and drop rows where we have no CDS start, end etc.
coding = seqs.query("is_coding").dropna().set_index("Refseq")
# convert cds start/end to ints
coding.cds_start = coding.cds_start.astype(np.int)
coding.cds_end = coding.cds_end.astype(np.int)
print(coding.shape)
print("{:0.2f}MB".format(coding.memory_usage(deep=True).sum() / 1000000))
coding.head(3)

# %% [code]
def extract_CDS_sequence(row):
    return row.seq[row.cds_start:row.cds_end]

def extract_3p_utr(row):
    return row.seq[row.cds_end:]

def extract_5p_utr(row):
    return row.seq[:row.cds_start]

coding["CDS"] = coding.apply(extract_CDS_sequence, axis=1)
coding["5_UTR"] = coding.apply(extract_5p_utr, axis=1)
coding["3_UTR"] = coding.apply(extract_3p_utr, axis=1)
# drop full sequence
# coding.drop("seq", axis=1, inplace=True)
print(coding.shape)
coding.head(3)

# %% [markdown]
### Checks and balances

We want to check a few things to make sure our data is reliable:
- Making sure CDS begins with M (ATG)
- Making sure CDS ends with a STOP codon (TAA, TAG, TGA)
- Seeing whether a translated CDS matches the translation provided by Refseq
- Making sure the CDS is divisible by 3.

We will eliminate any records that DO NOT start with a methionine.

# %% [code]
def check_translation(row):
    mseq = Bio.Seq.Seq(row.CDS, Bio.Alphabet.IUPAC.unambiguous_dna)
    try:
        aaseq = mseq.transcribe().translate(cds=True)
        return aaseq == row.translation
    except Bio.Data.CodonTable.TranslationError:
        return False

is_match = coding.apply(check_translation, axis=1)

# %% [code]
# 139 records have a translation that doesn't match what's on record.
is_match.shape[0] - is_match.sum()

# %% [code]
# removing non-methionine starters and matches to translation coding
coding.where(coding.CDS.str.startswith("ATG") & is_match, inplace=True)
# keeps shape, but fills rows with NaNs which we will drop
coding.dropna(subset=["CDS"], inplace=True)

# %% [code]
coding.shape

# %% [markdown]
## Extracting CDS, 3'-UTR and 5'-UTR features

From the paper:

    Sequence signatures and mRNA concentration can explain two-thirds of protein abundance variation in a human cell line, 2010, Vogel et al.
    
We draw heavy inspiration for features from this paper.
    
We see that length of each of these features corresponds to negative correlation to protein abundance.

Extracting:
- Length of CDS, 5'-UTR and 3'-UTR
- AUG (M) frequency in 5'-UTR
- uORFs (translation initiation) frequency in 5'-UTR
- Local secondary structures in 3'end of 5'-UTR (window size 40 or 60)
- mRNA decay rate ?
- miRNAs per 3'-UTR
- Codon bias index (CAI)
- tRNA adaptation index (tAI)
- G+C (total in CDS)
- G+C (third codon position only)
- Polyadenylation sites

Stability of the secondary structures in the 5′UTR is defined as the difference between the lowest free energy calculated for a segment of the real RNA sequence and the average of the lowest free energies of a large number of randomized segments with the same base composition and the same size divided by the s.d. of the free energies from the random sample.

### Frequency of dinucleotides

The frequency of dinucleotides in Vogel is recorded and may contribute a small factor to informative prediction.

### Kozak Sequence

The *Kozak consensus sequence* refers to theory that claims that eukaryotic mRNA has a sequence that plays a major role in translation initiation. It's presense therefore within the 5'-UTR should indicate higher translation concentration.

    gccRccATGG
    
where lowercase refers to the highest probable, but ATCG can be filled. ATGG (upper) is ubiquitous, and R refers to a must purine base (A or G).

The Kozak sequence stretches over the 5'-UTR and CDS region, whereby base 1, 2 and 3 is ATG respectively, but base -3 is usually an A/G and base 4 is a G/A.

### uORF

An upstream Open Reading Frame is an ORF within the 5'-UTR of mRNA. uORFs can regulate eukaryotic gene expression. Translation of uORF typically inhibits downstream expression of the primary ORF (CDS). 50% of genes in humans contain uORFs, and cause reductions in protein expression. An open reading frame is a part of the reading frame that has teh ability to be translated. An ORF begins with AUG and has a STOP codon (UAA, UAG, UGA). 

Read

    Gene Expression Regulation by Upstream Open Reading Frames and Human Disease, Barbosa et al, 2013.
    
### RNA secondary structure prediction

We are using the Vienna suite (RNAfold) algorithm (http://rna.tbi.univie.ac.at/), which we import in Ubuntu using:

    sudo apt-add-repository ppa:j-4/vienna-rna
    sudo apt-get update
    sudo apt-get install vienna-rna

The primary uses of this will be to analyse the 5'-UTR, to calculate the **minimum free energy (MFE)** i.e the most stable version of predicted folding, as an indication of the translation rate.

We will also generate the MFE using *windowing*, i.e the closest 10 bases to CDS, 20, 30.. and so on. This is because we believe that secondary structures closest to the beginning of the coding region are more likely to interfere with translation.

### Polyadenylation tails

Poly(A) tails within the 3'-UTR region of a mRNA sequence which contain regulatory regions thought of as stabilizing the mRNA molecule.

We are using a tool from the paper found at ( http://www.cbrc.kaust.edu.sa/omnipolya/ ):

    Omni-PolyA: a method and tool for accurate recognition of Poly(A) signals in human genomic DNA, Arturo Magana-Mora, Manal Kalkatawi, and Vladimir B. Bajic, 2017.    

# %% [code]
# extract codons backwards
def extract_codons_backstep(seq):
    # given an mRNA sequence
    return [seq[i:i+3] for i in range(len(seq), 0, -3)][1:]

def extract_dinucleotides(seq):
    return [seq[i:i+2] for i in range(0, len(seq), 2)]

def extract_dinucleotides_backstep(seq):
    return [seq[i:i+2] for i in range(len(seq), 0, -2)][1:]

def count_M(L):
    # given a list of codons
    if len(L) > 0:
        return sum([((c=="ATG") or (c=="AUG")) for c in L])
    else:
        return 0

def is_stop_codon(c):
    return ((c == "TAA") or (c == "TGA") or (c == "TAG"))

def extract_dinucleotide_count(seq, N, backward=True):
    if len(seq) > 2:
        if backward:
            dinucls = extract_dinucleotides_backstep(seq)
        else:
            dinucls = extract_dinucleotides(seq)
        return dinucls.count(N)
    else:
        return 0

def uORF_count_5_UTR(seq):
    n_uORFS = 0
    codons = extract_codons_backstep(seq)[::-1]
    # if we don't have an ATG start codon and ONE of the stop codons, return 0
    if (codons.count("ATG") > 0) and (codons.count("TAG") > 0 or codons.count("TAA") > 0 or codons.count("TGA") > 0):
        # create a series
        my_series = pd.Series(codons)
        # stack together only relevant codons in sequence
        sequence = my_series.where((my_series=="ATG") | (my_series=="TAA") | (my_series=="TAG") | (my_series=="TGA")).dropna()
        # find ATG, and if it is followed immediately by a STOP, mark that as an uORF
        for i,base in enumerate(sequence.values):
            if (base == "ATG") and (i < len(sequence)-1) and is_stop_codon(sequence.iloc[i+1]):
                n_uORFS += 1
                
    return n_uORFS

def repeats(s,n):
    return ''.join(list(my_rep(s,n)))

def add_spaces(string, n_len=10):
    if len(string) < n_len:
        return repeats(" ",n_len-len(string)) + string
    else:
        return string

from numba import jit

@jit
def scale_matrix(vec):
    D = np.empty((len(vec), len(vec)))
    for i in range(len(vec)):
        D[i,:] = vec[i] / vec
    return D

def prod_scale(row, column_names):
    D = scale_matrix(row.values)
    return pd.Series(np.prod(D, axis=1), index=column_names)

# %% [code]
def extract_5utr_free_energy(df, filepath="../../data/RNAfold/", fileout_e="energy_values.csv", 
                             fileout_s="energy_structures.csv", no_struct=False):
    
    """
    We are using the Vienna RNAfold package software with in-built command line functions
    along the way.
    
    Given a dataframe of two columns, where:
        1. NM_******** (ID)
        2. [AUGC]+ (mRNA sequence)
    
    Return a pandas.series(s) of the structure and free energy.
    
    This is saved in ../data/RNAfold/
    """
    # dictionary of energy values.
    energy_E = {}
    # a dictionary of k:v where k = NM_ accession and v: string representing structure
    if not no_struct:
        struct_D = {}
    
    t1 = time.clock()
    for i,x in df.iterrows():
         # print progress.
        if i % 1000 == 0:
            if not no_struct:
                # move .ps file to data folder
                !mv *_ss.ps {filepath}
        # check whether we have a sequence, and skip over if not
        if len(x.sequence) > 0:
            # create 'out file'.
            with open("out.txt", "w") as f:
                f.write(">"+x.Refseq_ID+"\n"+x.sequence)
            # call RNAfold, output into aptly named file
            if no_struct:
                !RNAfold -o --noPS < out.txt
            else:
                !RNAfold -o < out.txt

            # extract .fold file', the third element contains the output.
            with open(x.Refseq_ID+".fold","r") as f:
                output = f.readlines()
            # find the energy value
            energy = re.findall("(-*[0-9]+.[0-9]+)", output[2])
            # assign energy to dictionary if it's the only one
            if len(energy) == 1:
                energy_E[x.Refseq_ID] = energy[0]
            del energy

            if not no_struct:
                struct = re.findall("([.\(\)]+) ", output)
                if len(struct) == 1:
                    struct_D[x.Refseq_ID] = struct[0]
                del struct

            del output
            # delete output file
            !rm {x.Refseq_ID}".fold"

    # moving or deleting all files
    # if delete, do it, else move
    if not no_struct:
        # move .ps file to data folder
        !mv *_ss.ps {filepath}
        print("[Zipping together .ps files..]")
        # and zip, -m means not to keep the original ps files, output null
        !zip -m {filepath}ps_structures.zip {filepath}*_ss.ps > NULL
    
    !rm out.txt
    # once we have iterated over all the energies and structures, convert to series
    # and save in files in ..data/RNAfold/
    E = pd.Series(energy_E)
    # save to file
    E.to_csv(filepath+fileout_e)
    
    if not no_struct:
        D = pd.Series(struct_D)
        D.to_csv(filepath+fileout_s)

    print("[Tasks complete]")
    
    if not no_struct:
        return (E, D)
    else:
        return E

# %% [markdown]
### Coding sequence (CDS) features

We are NOT calculating amino-acid derived features; these will be done within the **UNIPROT** dataset we have obtained.

Features include:
- CDS Length
- CDS Length Proportion (CDS / mRNA length)
- Base counts (A, T, C and G)
- Dinucleotide count
- (strong kozak sequences)


# %% [code]
# dinucleotide counts
DNs = ["AA", "AC", "AG", "AT", "CA", "CC", "CG", "CT", "GA", "GC", "GG", "GT", "TA", "TC", "TG", "TT"]

# %% [code]
@cache("../../data/NCBI/cds_features.csv")
def calculate_cds_features():
    # create sub.
    cds_sub = coding.CDS.to_frame()
    cds_sub["length_cds"] = cds_sub.CDS.str.len()
    cds_sub["length_prop_cds"] = cds_sub.CDS.str.len() / coding["length_mrna"]
    # bases counts in CDS
    for b in ["A", "C", "T", "G"]:
        cds_sub["{}_cds".format(b)] = cds_sub.CDS.str.count(b)
    for dn in DNs:
        cds_sub["{}_cds".format(dn)] = cds_sub.CDS.apply(extract_dinucleotide_count, N=dn, backward=False)

    # extract consensus sequence identification - kozak
    kozak = (coding["5_UTR"].fillna("")
        .apply(add_spaces, n_len=8).str[-8:].add(coding["CDS"].str[:6]))
    cds_sub["kozak_cds"] = kozak.str.findall("([AG][ATCG]{2}ATGG)").apply(lambda x: len(x))
    # save to local file
    cds_sub.drop("CDS", axis=1, inplace=True)
    return cds_sub

# %% [code]
cds_sub = calculate_cds_features()
cds_sub.head(3)

# %% [code]
coding = coding.join(cds_sub, how="left")

# %% [markdown]
### 5'-UTR features

Features include:
- Length
- GC content
- The frequency of ATG in the region (correlated to translation)
- Base counts (A, T, C and G)
- Count of uORFs
- Dinucleotide counts
- The minimum free energy (MFE) within the 5'-UTR as an estimate for secondary structure

# %% [code]
@cache("../../data/NCBI/utr5_features.csv")
def calculate_5utr_features():
    utr5_sub = coding["5_UTR"].to_frame()
    utr5_sub["length_utr5"] = utr5_sub["5_UTR"].str.len()
    utr5_sub["length_prop_utr5"] = utr5_sub["5_UTR"].str.len() / coding["length_mrna"]
    # bases counts in 5'-UTR
    for b in ["A", "C", "T", "G"]:
        utr5_sub["{}_utr5".format(b)] = utr5_sub["5_UTR"].str.count(b)
    # count the number of uORFs in 5-'uTR
    utr5_sub["uORF_utr5"] = utr5_sub["5_UTR"].apply(uORF_count_5_UTR)
    for dn in DNs:
        utr5_sub["{}_utr5".format(dn)] = utr5_sub["5_UTR"].apply(extract_dinucleotide_count, N=dn)
    # AUG/ATG frequency in 5'-UTR
    utr5_sub["M_utr5"] = utr5_sub["5_UTR"].apply(extract_codons_backstep).apply(count_M)
    # save changes to file
    utr5_sub.drop("5_UTR", axis=1, inplace=True)
    return utr5_sub

# %% [code]
utr5_sub = calculate_5utr_features()
utr5_sub.head(3)

# %% [code]
coding = coding.join(utr5_sub, how="left")

# %% [markdown]
*Mininum Free Energy Calculations*

# %% [code]
if os.path.isfile("../../data/RNAfold/5_UTR_energy_values.csv"):
    rnafold = pd.read_csv("../../data/RNAfold/5_UTR_energy_values.csv", header=None,
                          names=["Refseq_ID","5_UTR_MFE"], index_col=0)
    coding = coding.join(rnafold, how="outer")
else:
    # preparing sequences ready for RNAfold
    intro_seqs = coding.reset_index()[["id","5_UTR"]].dropna()
    # replace thymine with uracil
    intro_seqs["5_UTR"] = intro_seqs["5_UTR"].str.replace("T","U")
    intro_seqs.rename(columns={"5_UTR":"sequence"}, inplace=True)
    # call function - expensive!
    E, D = extract_5utr_free_energy(intro_seqs, filepath="../../data/RNAfold/", 
                                    fileout_e="5_UTR_energy_values.csv",
                                    no_struct=True)
    # add energy values to merge.
    coding["5_UTR_MFE"] = E

# %% [code]
# windowed approach. we will do window sizes of 10, 20, 40, and 80 for 5'-UTR, such that
# the window is closes to the 3' end (CDS)
windows = [10, 20, 30, 40, 60, 80]
if os.path.isfile("../../data/RNAfold/window80/5_UTR_energy_values.csv"):
    for w in windows:
        rnafold = pd.read_csv("../../data/RNAfold/window{}/5_UTR_energy_values.csv".format(w), header=None,
                             names=["Refseq_ID","5_UTR_MFE_win{}".format(w)], index_col=0)
        coding = coding.join(rnafold, how="outer")
else:
    intro_seqs = coding.reset_index()[["id","5_UTR"]].dropna()
    intro_seqs["5_UTR"] = intro_seqs["5_UTR"].str.replace("T","U")
    for w in windows:
        print("[Beginning Window %d]" % w)
        # extract window
        intro_seqs["sequence"] = intro_seqs["5_UTR"].str[-w:]
        # call function - expensive!
        E = extract_5utr_free_energy(intro_seqs.drop("5_UTR", axis=1),
                                     filepath="../../data/RNAfold/window{}/".format(w),
                                     fileout_e="5_UTR_energy_values.csv",
                                     no_struct=True)
        coding["5_UTR_MFE_win{}".format(w)] = E

# %% [markdown]
#### Renaming MFE features

# %% [code]
coding.columns=coding.columns.str.replace("5_UTR_", "")

# %% [code]
coding.count().values

# %% [code]
coding.dropna(inplace=True)

# %% [markdown]
### 3'-UTR features

Features include:
- Length
- Dinucleotide counts
- Proportion of 3'-UTR length with respect to entire sequence

# %% [code]
@cache("../../data/NCBI/utr3_features.csv")
def calculate_3utr_features():
    # fill missing values
    utr3_sub = coding["3_UTR"].to_frame()
    # fill NA
    utr3_sub["3_UTR"].fillna("", inplace=True)
    utr3_sub["length_utr3"] = utr3_sub["3_UTR"].str.len()
    utr3_sub["length_prop_utr3"] = utr3_sub["3_UTR"].str.len() / coding["length_mrna"]
    # bases counts in 3'-UTR
    for b in ["A", "C", "T", "G"]:
        utr3_sub["{}_utr3".format(b)] = utr3_sub["3_UTR"].str.count(b)
    # bases dinucleotide
    for dn in DNs:
        utr3_sub["{}_utr3".format(dn)] = utr3_sub["3_UTR"].apply(extract_dinucleotide_count, N=dn, backward=False)
    # save changes to file
    utr3_sub.drop("3_UTR", axis=1, inplace=True)
    return utr3_sub

# %% [code]
utr3_sub = calculate_3utr_features()
utr3_sub.head()

# %% [code]
coding = coding.join(utr3_sub, how="left")

# %% [code]
for dn in DNs:
    coding["{}_mrna".format(dn)] = coding["seq"].apply(extract_dinucleotide_count, N=dn)

# %% [markdown]
### Polyadenylated Tails

PAS.txt is a tab-delimited file containing all annotated PASs based on information from RefSeq, Ensembl, and FANTOM5 (human only).

Columns:

1. __PAS_ID__: ID of each PAS, shown as Chromosome:Position:Strand
2. __Chromosome__: chromosome ID of PAS
3. __Position__: genomic position of PAS
4. __Strand__: strand information of PAS
5. __Mean RPM__: Average reads per million of PAS reads across all samples
6. __Intron/exon location__: PAS location relative to splicing configuration, including 5'-most exon, internal exon, 3'-most exon, and intron. If a gene has only one exon, it is called single exon. 
7. __Ensembl Gene ID__: Ensembl gene ID of the gene containing the PAS
8. __RefSeq Gene ID__: RefSeq gene ID of the gene containing the PAS
9. __Gene Symbol__: Gene symbol from RefSeq
10. __Gene Name__: Gene name from RefSeq
11. __Extension__: Whether PAS is located on an extended 3' end region beyond RefSeq/Ensembl annotations, Yes/No
12. __PAS type__: PAS location in annotated genes, including 5'UTR, CDS, 3'UTR and intergenic. For PASs in 3'UTRs, they are further divided into First (F), Middle (M), and Last (L). If there is only one PAS in 3'UTR, it is called S. 
13. __PSE__: Percentage of samples with detected expression of all samples.
14. __PAS Signal__: The PAS signal is located within 40-nt upstream from the PAS, including AAUAAA, AUUAAA, Other (including AGTAAA, TATAAA, CATAAA, GATAAA, AATATA, AATACA, AATAGA, AAAAAG, and ACTAAA), A-rich (AAAAAA), and None. 
15. __Conservation__: Whether the PAS in conserved in at least two mammals (human, mouse and rat).
16. __FAMTOM ID__ (human only): FAMTOM lncRNA ID of the gene containing the human PAS
17. __FANTOM_Category__ (human only): LncRNA category defined by the FANTOM5 
18. __Intergenic_TE__: Whether the intergenic PAS is located on an ±24 bp region of an annoated transposable element, Yes/No

# %% [code]
polyas = pd.read_table("../../data/polya_db/human_PAS.txt", na_values=["na","NO"])
polyas["PSE"] = polyas["PSE"].str[:-1].astype(np.float)

# %% [markdown]
### CHECK: Drop samples with chronically low PSE score (i.e detected expression samples as a proportion less than 10%)

# %% [code]
polya_drop = polyas.query("PSE > 10")

# %% [code]
polyas_group = (polya_drop.groupby(["Gene Symbol"])["Mean RPM"].agg(["mean","std"])
    .rename(columns={"mean":"signal_polya","std":"signal_polya_std"}).reset_index())

# %% [code]
polyas_group.head(3)

# %% [code]
polyas_group.shape

# %% [code]
polyas_group.to_csv("../../data/polya_db/polya.csv", na_rep=np.nan, index=None)

# %% [markdown]
#### Integrate PolyAs Gene Symbol with HGNC to get Refseq - then merge

Using `turbopanda`:

# %% [code]
hgnc = turb.read("../../data/HGNC/HGNC_gene.csv",sep="\t")
hgnc.drop("date","vega","mgd","orpha","iupha","homeo","pubmed","rgd","cosm","^location$","omim")
hgnc.rename([("hgnc_",""),("_sortable",""),("_symbol",""), ("gene_",""),
             ("accession","id"), ("ids","id")], apply=True)

# %% [code]
temp_map = hgnc.df_.set_index("symbol")["alias"].dropna().str.split("|").explode()
# swap them around and turn it into a dict
alias_to_current = pd.Series(temp_map.index, index=temp_map.values).to_dict()
# replace aliases in PolyA dataset with current HGNC best.
polyas_group["Gene Symbol"].replace(alias_to_current, inplace=True)

# %% [markdown]
Merge together PolyA data with the HGNC dataset.

# %% [code]
pa_merge = pd.merge(hgnc.df_, polyas_group, left_on="symbol", right_on="Gene Symbol", how="inner")[["refseq_id","signal_polya"]].set_index("refseq_id")

# %% [code]
pa_merge.head(3)

# %% [code]
coding = coding.join(pa_merge, how="left")

# %% [code]
coding.head(3)

# %% [markdown]
### Codon Adaptation Index (CAI)

Here we estimate CAI following from paper:
- **The codon adaptation index - a measure of directional synonymos codon usage bias, and its potential applications, 1987, Paul M. Sharp**

Firstly we construct a reference table of relative synonymous codon usage values from highly expressed genes of the organism. 

$$
\text{RSCU}_{ij}=\frac{X_{ij}}{\frac{1}{n}\sum_{j=1}^{n_i}X_{ij}}
$$

where $X_{ij}$ is the number of occurances of the jth codon for the ith amino acid, and n is the number of alternative codons for the ith amino acid. The relative adaptiveness of a codon, $w_{ij}$, is then the frequency of use of that odon compared to the frequency of the optimal codon for that amino acid.

$$
w_{ij}=\frac{\text{RSCU}_{ij}}{\text{RSCU}_{imax}} - \frac{X_{ij}}{X_{imax}}
$$

where $RSCU_{imax}$ and $X_{imax}$ are the RSCU and X values for the most frequently used codon for the ith amino acid. The CAI for a gene is calculated as the geometric mean of the RSCU values corresponding to each of the codons used in that gene, divided by the maximum possible CAI for a gene of the same amino acid composition.

$$
\text{CAI}=\left(\prod_{k=1}^L w_k\right)^\frac{1}{L}
$$

We made use of an online tool called CAIcal:
- http://genomes.urv.es/CAIcal/

We entered RNA sequence calculations using the Human Codon Usage table as frequencies per thousand:
- http://genomes.urv.es/CAIcal/CU_human_nature

Results were tabulated into cai-output flat files - we mash these into our CAI-Nc dataset, with Primary RNA Accession as an index. As input, we use the Coding sequence (CDS) FASTA file cache that was developed in NCBI-Access.

We execute the __perl__ script (using version 5) in `../tools/CAIcal/`:

    perl CAIcal_v1.4.pl -e cai_and_expected -f ../data/NCBI/cds_cache.fasta -h human -g 1 -c 95 -n 100000 -l 10000 -m markov -o1 ../data/CAI/main_out.txt -o2 ../data/CAI/rand_seqs.txt -o3 ../data/CAI/expected.txt
    
Outputted files are placed in `../data/CAI/`.

Input parameters are:

- -e (cai|expected|cai_and_expected)
- -f file FASTA input
- -g codon reference table (we use default human)
- -c confidence intervals
- -n number of samples to expect
- -l length of genes to expect
- -m method type, we use markov default
- -o1 output file 1
- -o2 output file 2
- -o3 output file 3

# %% [code]
if os.path.isfile("../../data/CAI/main.txt"):
    cai = pd.read_table("../../data/CAI/main.txt", sep="\t").drop("Warning", axis=1)
else:
    # run the script!
    !perl ../tools/CAIcal/CAIcal_ECAI_v1.4.pl -e cai_and_expected -f ../data/NCBI/cds_cache.fasta -h human -g 1 -c 95 -n 100000 -l 10000 -m markov -o1 ../data/CAI/main.txt -o2 ../data/CAI/rand_seqs.txt -o3 ../data/CAI/expected.txt
    # load in file
    cai = pd.read_table("../../data/CAI/main.txt", sep="\t").drop("Warning", axis=1)
    
cai["Refseq_ID"] = cai.NAME.str.extract(">(?P<Refseq_ID>NM_[0-9]+)", expand=False)
cai.drop("NAME", axis=1, inplace=True)
cai.set_index("Refseq_ID", inplace=True)
# remove random error '#Error# The length of this sequence is not divided by three.'
# convert to numeric and coerce 'errors' to nan
cai.CAI = pd.to_numeric(cai.CAI, errors="coerce")

# %% [code]
cai.head(3)

# %% [code]
coding = coding.join(cai, how="left")

# %% [code]
coding.head(3)

# %% [markdown]
### tRNA Adaptation Index (tAI)

We can calculate relatively accurate tRNA-adaptation from sequence data using stAIcalc, from the paper:
- **stAIcalc: tRNA adaptation index calculator based on species-specific weights. Renana Sabi et al. 2016**

There is no direct method where we can extract this metric for each transcript. A website exists to calculate tAI given a gene coding sequence however Homo sapiens is not yet available for this:
- http://tau-tai.azurewebsites.net/

The tool, however has an offline version which I downloaded for Linux Ubuntu. I then extracted Homo sapiens gene-copy number data (as the original paper suggests) from:
- http://gtrnadb.ucsc.edu/genomes/eukaryota/Hsapi19/

which has a complete tRNA genome for Homo sapiens, with gene copy numbers for each anticodon present at the 'Summary' section of the first page on Homo sapiens (hg19 - NCBI build 37.1 Feb 2009).

This information was written into a file like the following:
- *Anticodon* (tab) **tRNA gene copy number** \n

In addition to this new file, we compiled our list of coding sequence transcripts as FASTA files and executed the offline program.

To the run the program on Linux Ubuntu, we used MATLAB 2016a x64 bit. Once set up, we called: 

    ./run_stAIcalc.sh <directory to matlab>

We set the hill climbing stringency to medium.

# %% [code]
records = list(SeqIO.parse("../../data/NCBI/cds_cache.fasta","fasta"))

# %% [code]
# in steps of 1000, we will create fasta records that divide and conquer.
indices = np.empty(round(len(records) / 1000. + 1), dtype=np.int)
indices[:-1] = np.arange(0, 52000, 1000)
indices[-1] = len(records)

# %% [code]
for i in range(1, len(indices)):
    with open("../../data/tRNA/tAI_chunks/{}_{}tAI_slice.fasta".format(indices[i-1],indices[i]-1),"w") as h:
        SeqIO.write(records[indices[i-1]:indices[i]], h, "fasta")

# %% [code]
indices

# %% [markdown]
failed_pairs = [(1000, 2000), (5000, 6000), (6000, 7000), (7000, 8000), (42000, 43000), (44000, 45000)]
for p1, p2 in failed_pairs:
    length = int((p2 - p1) / 2.)
    with open("../../data/tRNA/tAI_chunks/{}_{}tAI_slice.fasta".format(p1, p1+length-1), "w") as f:
        SeqIO.write(records[p1:p1+length], f, "fasta")
    with open("../../data/tRNA/tAI_chunks/{}_{}tAI_slice.fasta".format(p1+length, p2-1), "w") as f:
        SeqIO.write(records[p1+length:p2], f, "fasta")

# %% [markdown]
#### Here we run the tAI GUI....

### merge together tAI

# %% [code]
tai_files = !ls ../../data/tRNA/*_tai.txt

# %% [code]
tai = pd.concat([pd.read_csv(f, sep="\t") for f in tai_files], axis=0)
tai["Refseq_ID"] = tai.genes_names.str.extract("(?P<Refseq_ID>NM_[0-9]+)", expand=False)
tai.drop("genes_names", axis=1, inplace=True)
tai.set_index("Refseq_ID", inplace=True)
tai.head(3)

# %% [code]
print(tai.shape)

# %% [code]
coding = coding.join(tai, how="left")

# %% [code]
coding.head(2)

# %% [markdown]
### Relative Codon Usage Bias (RCBS)

Taken from paper **'Relative Codon Adaptation: A Generic Codon Bias Index for Prediction of Gene Expression, 2010, Jesse M. Fox'**

Similar to CAI which uses a subset of putative highly expressed genes. RCBS attempts to estimate Codon bias without a reference set:

$$
d_{xyz}=\frac{f(x,y,z)}{f_1(x).f_2(y).f_3(z)} \\
RCBS=\left(\prod_{k=1}^L d_{xyz}(k)\right)^\frac{1}{L}-1
$$

where $L$ is the length in codons of the gene, $f(x,y,z)$ the observed frequency of codon xyz and $f_1(x)$, $f_2(y)$, and $f_3(z)$ are the observed frequencies of bases $x$, $y$, $z$ at respectively, codon positions 1, 2 and 3. $d_{xyz}$ is counted on a per-gene basis, and hence all of the frequencies of codons and bases are not global but by gene. Thus it is important when calculating RCBS to substitute $d_{xyz}$ into the list of codons for each gene before calculating the product, as $L$ is by the length of codons, not the length of *unique* codons. Frequencies are calculated as counts scaled by the length of codons. $f_1(x)$, $f_2(y)$, and $f_3(z)$ is the frequency of either A, C, G or T.

It's worth mentioning that due to quickly reaching infinity, computationally we calculate the product as the sum of it's logarithms, as this is considerably more stable from a numerical perspective:

$$
\prod_{k=1}^L d_{xyz}(k) \approx \exp {\left(\sum_{k=1}^L \log (d_{xyz}(k))\right)}
$$

However, Fox provides a pseudo-correction to RCBS, since RCBS is heavily affected by the length of the gene. Fox finds that shorter sequences are significantly biased, compared to longer sequences ($\gt1000$). They estimate the frequencies as:

$$
\phi_{xyz} = \frac{f(x,y,z)L + g(x,y,z)S}{L+S} \\
\phi_i(j) = \frac{f_i(j)L + g_i(j).S}{L+S}
$$

where $L$ is the length in codons of the sampled sequence, $g(x,y,z)$ the observed frequency of codon $xyz$ among genomic-coding regions, $g_n(m)$ the observed frequency of base $m$ at codon position $n$, also in genomic-coding regions, and $S$ the weight on pseudo-counts (priors), also measured in codons. Fox uses $S=300$ to compute $RCBS^{PC}$. We refer to this corrected version as $RCBS^{PC}$, is then computed as above, by substituting $d_{xyz}$ with $\phi_{xyz}$ and $f_i(j)$ with $\phi_i(j)$.

We implement this here:

# %% [code]
def calculate_rcbs(cds_seq):
    """
    Given an mRNA sequence of DNA characters which is the coding sequence,
    return the Relative CUB as denoted by Fox et al., 2010. originally by Roymondal et al., 2009
    
    Can either use Roymondal method or Fox method. I prefer Fox.
    
    Returns a scalar value, usually in the range of 0.2-0.6 (check distribution on hist)
    """
    # split our CDS into codons
    codons = pd.Series([cds_seq[i:i+3] for i in range(0, len(cds_seq), 3)])
    L = len(codons)
    # calculate frequency of every codon f(x,y,z)
    fxyz = codons.value_counts() / L
    # calculate frequency per codon-base position f(x), f(y) and f(z)
    fx = codons.str[0].value_counts() / L
    fy = codons.str[1].value_counts() / L
    fz = codons.str[2].value_counts() / L
    # calculate d(x,y,z) for each codon
    dxyz = pd.Series([freq / (fx[cod[0]]*fy[cod[1]]*fz[cod[2]])
                      for cod, freq in fxyz.iteritems()],
            index=fxyz.index)
    # calculate RCBS
    #  We 'map' dxyz values into each codon slot, then calculate the product, and scale by 1/length in codons - 1
    return np.exp(codons.map(dxyz.to_dict()).map(np.log).sum()) ** (1./L) - 1.

def calculate_rcbs_corrected(cds_seq, g_xyz, gx, gy, gz, S=300.):
    """
    Given an mRNA sequence of DNA characters which is the coding sequence,
    return the Relative CUB as denoted by Fox et al., 2010.
    
    Also pass in the global frequency of codons g(x,y,z) and g1(x), g2(y) and g3(z).
    
    This builds on Roymondal by providing a correction, see above.
    
    g_xyz is a correction to f_xyz, which we do not directly calculate.
    gx, gy, and gz are corrections to fx, fy, fz, which we simplify
    
    Returns a scalar value, usually in the range of 0.2-0.6 (check distribution on hist)
    """
    # split our CDS into codons
    codons = pd.Series([cds_seq[i:i+3] for i in range(0, len(cds_seq), 3)])
    L = len(codons)
    # calculate phi_xyz, where codons.value_counts() = f_xyz/L*L
    phi_xyz = (codons.value_counts() + g_xyz*S) / (L+S)
    # calculate phi_i(j) = f_i(j)*L + g_i(j)*S / L+S
    phi_x = (codons.str[0].value_counts() + gx*S) / (L+S)
    phi_y = (codons.str[1].value_counts() + gy*S) / (L+S)
    phi_z = (codons.str[2].value_counts() + gz*S) / (L+S)
    # calculate RCB^PC
    dxyz = pd.Series([freq / (phi_x[cod[0]]*phi_y[cod[1]]*phi_z[cod[2]])
                      for cod, freq in phi_xyz.iteritems()],
                index=phi_xyz.index)
    
    return np.exp(codons.map(dxyz.to_dict()).map(np.log).sum()) ** (1./L) - 1.

# %% [code]
if os.path.isfile("../../data/CAI/RCBS.csv"):
    rcbs = pd.read_csv("../../data/CAI/RCBS.csv", sep="\t", index_col=0)
    # add to coding
    coding = coding.join(rcbs, how="left")
else:
    # run codes
    # extract all of the bases in every coding sequence (genomic-coding sequence)
    global_bases = coding.CDS.str.extractall("([ATCG]{3})")
    # calculate codon frequency
    g_xyz = global_bases[0].value_counts() / global_bases.shape[0]
    # calculate codon position frequency
    gx = global_bases[0].str[0].value_counts() / global_bases.shape[0]
    gy = global_bases[0].str[1].value_counts() / global_bases.shape[0]
    gz = global_bases[0].str[2].value_counts() / global_bases.shape[0]
    # we will use the fox implementation
    coding["RCBS"] = coding.CDS.dropna().apply(calculate_rcbs)
    coding["RCBS_PC"] = coding.CDS.dropna().apply(calculate_rcbs_corrected, g_xyz=g_xyz, gx=gx, gy=gy, gz=gz, S=300.)
    # save to file
    coding[["RCBS","RCBS_PC"]].to_csv("../../data/CAI/RCBS.csv", sep="\t")

# %% [code]
coding.RCBS_PC.replace([np.inf, -np.inf], np.nan).dropna().hist(bins=20)
plt.show()

# %% [code]
# replacing infs with np.nans
coding.RCBS.replace([np.inf, -np.inf], np.nan, inplace=True)
coding.RCBS_PC.replace([np.inf, -np.inf], np.nan, inplace=True)

# %% [markdown]
## Associate with Feature Counts

We will *now* merge in our feature counts that we calculated in NCBI-Access.

We will also drop some of the feature counts that are IRRELEVANT, such as CDS (they all have...), gene etc.

# %% [code]
feature_counts = turb.read(path+"rna_feature_matrix.csv", sep="\t", index_col=0)

# %% [code]
feature_counts

# %% [markdown]
#### Step 1. Drop useless feature counts such as CDS, gene, source etc.

# %% [code]
# drop useless columns
feature_counts.drop("^CDS","^gene","_base","^source","polyA_")

# %% [markdown]
#### Step 2. Drop features with less than 200 values

# %% [code]
feature_counts.drop(lambda df: df.count()<200)

# %% [markdown]
#### Step 3. Convert features with only one/two options into a boolean switch

# %% [code]
# cache
feature_counts.cache("bool_switch", lambda df: df.value_counts().shape[0]<=2)
# transform nans to 0
feature_counts.transform("bool_switch", np.nan_to_num)
# transform x > 0
feature_counts.transform("bool_switch", lambda x: x > 0)

# %% [markdown]
#### Step 4. Fill missing count features with 0

# %% [code]
feature_counts.apply('fillna', 0)

# %% [code]
feature_counts.head(3)

# %% [markdown]
#### Step 5. Add suffix

# %% [code]
feature_counts.apply("pipe", turb.pipe.add_suffix, "_bs", bool)
feature_counts.apply("pipe", turb.pipe.add_suffix, "_fc", float)

# %% [code]
coding = coding.join(feature_counts.df_, how="left")
print(coding.shape)
coding.head(3)

# %% [markdown]
### Saving NM_ files to dataset

Firstly, we will drop CDS, 5'-UTR and 3'-UTR sequences to save space. These are already extracted in `..data/NCBI/`.

# %% [code]
m_coding = turb.MetaPanda(coding)

# %% [code]
m_coding.drop(object, "codon","cds_","^strand", "^is_coding")

# %% [code]
m_coding.write("../../datasets/refseq_mRNA_unscaled.csv")

# %% [markdown]
## Normalization Strategy

**For every count metric, we will scale it by the respective length of the element.** For example, the number of C bases in the mRNA sequence will be scaled by the *length* of the mRNA sequence.

Whereas for CDS features, we will scale by the length of the CDS sequence, same for 5'-UTR and 3'-UTR.

We have associated column name to normalization within `MetaDF`; so we need to associate group with a numerical value, array or function.

1. **Length-based features**: Scale by $log_2(x + 1)$. This normalizes them.
2. **Mononucleotide frequency**: Divide by the length of the count feature. For instance, if we have A_cds, we divide A_cds by the CDS length. Feature $F_{norm}=\frac{F}{F_{len}}$
3. **Dinucleotide frequency**: Divide by the length of the count feature by 2. For instance, if we have AA_cds, we divide AA_cds by the CDS length divided by 2. Feature $F_{norm}=\frac{F}{F_{len}/2}$
4. **Feature count**: Since this is over the entire NCBI Refseq mRNA, we do by total mRNA length.
5. **Minimum Free energy**: This measurement $\Delta G$ decreases negatively as the amount of secondary structure increases. The window (WIN) determines how many steps back from ORF that is estimated for secondary structure. So we scale by this size to reveal the proportion of free energy taking size into account. With MFE and no window, the entire 5'UTR is measured for secondary structure. $\Delta G_{mod}=-\frac{\Delta G}{n}$


# %% [markdown]
#### Cache names

# %% [code]
m_coding.cache_k(length="length_(?!prop)",
                     mono_mrna="^[ATCG]_mrna",
                     mono_cds="^[ATCG]_cds",
                     mono_utr5="^[ATCG]_utr5",
                     mono_utr3="^[ATCG]_utr3",
                     di_mrna="^[ATCG]{2}_mrna",
                     di_cds="^[ATCG]{2}_cds",
                     di_utr5="^[ATCG]{2}_utr5",
                     di_utr3="^[ATCG]{2}_utr3",
                     kozak="^kozak",
                     feat_count="_fc$",
                     free_e="^MFE$",
                     methio="^M_utr5",
                     **{"MFE%d" % d: "^MFE_win%d" % d for d in [10, 20, 30, 40, 60, 80]})

# %% [markdown]
#### Multi transform

# %% [code]
m_coding.transform_k([
    *[("MFE%d" % d, lambda x: -(x / d)) for d in [10,20,30,40,60,80]],
    ("mono_mrna", lambda x: x / m_coding.df_["length_mrna"]),
    ("di_mrna", lambda x: x / (m_coding.df_["length_mrna"] / 2.)),
    ("mono_cds", lambda x: x / m_coding.df_["length_cds"]),
    ("di_cds", lambda x: x / (m_coding.df_["length_cds"] / 2.)),
    ("mono_utr5", lambda x: x / m_coding.df_["length_utr5"]),
    ("di_utr5", lambda x: x / (m_coding.df_["length_utr5"] / 2.)),
    ("mono_utr3", lambda x: x / m_coding.df_["length_utr3"]),
    ("di_utr3", lambda x: x / (m_coding.df_["length_utr3"] / 2.)),
    ("kozak", lambda x: x / coding["length_cds"]),
    ("feat_count", lambda x: x / coding["length_mrna"]),
    ("free_e", lambda x: -(x / coding["length_utr5"])),
    ("length", lambda x: np.log2(x+1)),
    ("methio", lambda x: x / (coding["length_utr5"] / 3))
])

# %% [code]
m_coding.head()

# %% [markdown]
### Save scaled dataset also

# %% [code]
m_coding.write("../../datasets/refseq_mRNA_scaled.csv")
```
