## Supplementary material for "Predicting Relative Protein Abundance via Sequence-Based Information": S3 Data: Uniprot-Preprocessing.ipynb

```
# %% [markdown]
# Uniprot Preprocessing

In this notebook we are going to derive the sequence-based features from the amino acid sequence, whether that's the translated amino acids or the codons that form it.

There are a whole plethora of features we may wish to extract from the underlying amino acid sequence. Many of them involve physical and chemical attributes we have about each amino acid and their cumulative impact.

We will also access our uniprot data from `Uniprot-Access` and preprocess that file to extract useful things from it.

Our main sources of data are:

- Information found in Uniprot, including sequence-derived features.
- PEST regions
- Post-translational modifications

# %% [code]
import re
import os
import pandas as pd
import numpy as np
import matplotlib.pyplot as plt
import seaborn as sns
import itertools as it

import Bio
from Bio import SeqIO, Seq
from Bio.SeqRecord import SeqRecord
from Bio.SeqUtils import ProtParam

import sys
sys.path.insert(0, "../../scripts/")
import misc_functions as misc
import myfig as mf
import regexx
%matplotlib inline

path = "../../data/Uniprot/"

# %% [markdown]
## The Uniprot dataset

Using the refined Uniprot dataset iterablein `../datasets/Uniprot/` we can extract additional useful features.

# %% [code]
uniprot = pd.read_csv(path+"uniprot_all.csv", sep="\t", index_col=0).drop("Length", axis=1)

print(uniprot.shape)
print(uniprot.columns)
uniprot.head(3)

# %% [markdown]
## Feature Table

In addition to the amino acid sequence, Uniprot provides a database of 'Sequence annotations' with position-specific annotation with respect to the amino acid sequence. Taken from the [Uniprot website](https://www.uniprot.org/help/sequence_annotation):

*Molecular processing features*

* Initiator methionine: Cleavage of the initiator methionine
* Signal peptide: Sequence targeting proteins to the secretory pathway or periplasmic space
* Transit peptide: Extent of a transit peptide for organelle targeting
* Propeptide: Part of a protein that is cleaved during maturation or activation
* Chain: Extent of a polypeptide chain in the mature protein
* Peptide: Extent of an active peptide in the mature protein

*Regions*

* Topological domain: Location of non-membrane regions of membrane-spanning proteins
* Transmembrane: Extent of a membrane-spanning region
* Intramembrane: Extent of a region located in a membrane without crossing it
* Domain: Position and type of each modular protein domain
* Repeat: Positions of repeated sequence motifs or repeated domains
* Calcium binding: Position(s) of calcium binding region(s) within the protein
* Zinc finger: Position(s) and type(s) of zinc fingers within the protein
* DNA binding: Position and type of a DNA-binding domain
* Nucleotide binding: Nucleotide phosphate binding region
* Region: Region of interest in the sequence
* Coiled coil: Positions of regions of coiled coil within the protein
* Motif: Short (up to 20 amino acids) sequence motif of biological interest
* Compositional bias: Region of compositional bias in the protein

*Sites*

* Active site: Amino acid(s) directly involved in the activity of an enzyme
* Metal binding: Binding site for a metal ion
* Binding site: Binding site for any chemical group (co-enzyme, prosthetic group, etc.) 
* Site: Any interesting single amino acid site on the sequence

*Amino acid modifications*

* non-standard residue: Occurence of non-standard amino acids (selenocysteine and pyrrolysine) in the protein sequence
* modified residue: Modified residues excluding lipids, glycans and protein cross-links
* Lipidation: Covalently attached lipid group(s)
* Glycosylation: Covalently attached glycan group(s)
* Disulfide bond: Cysteine residues participating in disulfide bonds
* Cross-link: Residues participating in covalent linkage(s) between proteins

*Natural variations*

* Alternative sequence: Amino acid change(s) producing alternate protein isoforms
* Natural variant: Description of a natural variant of the protein

*Secondary structure*

* Helix: Helical regions within the experimentally determined protein structure
* Turn: Turns within the experimentally determined protein structure
* Beta strand: Beta strand regions within the experimentally determined protein structure

# %% [code]
feature_split = uniprot["Features"].str.split(";").explode().str.extract("(?P<feature_name>[A-Za-z\s]+)\((?:[0-9]+)\)")
feature_split["feature_name"] = feature_split["feature_name"].str.strip()

# %% [code]
feature_table = (uniprot["Features"].str.split(";")
    .explode()
    .str.extract("([a-z\s]+)", re.IGNORECASE, expand=False)
    .str.strip()
    .groupby("Uniprot_ID")
    .apply(lambda x: x.str.cat(sep=";"))
    .str.get_dummies(";"))

# %% [code]
feature_table.head(2)

# %% [markdown]
#### using `regexx` to modify column names in feature table

# %% [code]
feature_table.columns = regexx.replace(feature_table.columns.str.lower(),
                                       [("erroneous","err"), ("compositional","comp"), ("sequence","seq"), ("binding","bind"), (" ", "-")])

# %% [markdown]
### Convert Mass to float

# %% [code]
uniprot.Mass = pd.to_numeric(
    uniprot.Mass.str.extractall("([0-9]+)")
    .unstack("match")
    .apply(lambda x: x.str.cat(), axis=1)
)

# %% [markdown]
## Create workspace subset

# %% [code]
unip_sub = uniprot.Seq.to_frame()
unip_sub.head(2)

# %% [markdown]
### Calculate amino acid length

# %% [code]
unip_sub["length_aa"] = unip_sub.Seq.str.len()

# %% [markdown]
### Calculating counts of every amino acid

# %% [code]
aas = ["M", "F", "L", "I","V", "S", "P", "T", "A", "Y", "H", "Q", "N", "K", "D", "E", "C",
       "W", "R", "G"]
for aa in aas:
    unip_sub["{}_aa".format(aa)] = uniprot.Seq.str.count(aa)

# %% [code]
unip_sub.head(3)

# %% [markdown]
## Biophysical Properties

Most of these we obtain from the ProtParam package associated with `biopython`.

### Isoelectric point

Taken from paper:
- **Bjellqvist, B.,Hughes, G.J., Pasquali, Ch., Paquet, N., Ravier, F., Sanchez, J.-Ch., Frutiger, S. & Hochstrasser, D.F. The focusing positions of polypeptides in immobilized pH gradients can be predicted from their amino acid sequences. Electrophoresis 1993, 14, 1023-1031.**

*'The isoelectric point (pI, pH(I), IEP), is the pH at which a particular molecule carries no net electrical charge in the statistical mean. The standard nomenclature to represent the isoelectric point is pH(I), although pI is also commonly seen, and is used in this article for brevity. The net charge on the molecule is affected by pH of its surrounding environment and can become more positively or negatively charged due to the gain or loss, respectively, of protons (H+).

Surfaces naturally charge to form a double layer. In the common case when the surface charge-determining ions are H+/OH−, the net surface charge is affected by the pH of the liquid in which the solid is submerged.'* From Wiki.

# %% [code]
# some sequences contain amino acid U (stop) or X, so we will 
# remove those before performing ProteinAnalysis.
# perform analysis on proteins, then extract.
analysed_aas = (unip_sub.Seq[~(unip_sub.Seq.str.contains("B") |
                                  unip_sub.Seq.str.contains("U"))]
    .apply(lambda x: ProtParam.ProteinAnalysis(x)))

# %% [code]
unip_sub["pH_Isoelectric_point_bp"] = analysed_aas.apply(lambda x: x.isoelectric_point())

# %% [code]
sns.distplot(unip_sub["pH_Isoelectric_point_bp"].dropna())
plt.show()

# %% [markdown]
### Aromaticity (Protein)

This is a simple metric for averaging over amino acids which contain an aromatic ring within them. Particular amino acids such as Cysteine, Phenylalanine etc.

# %% [code]
unip_sub["Aromaticity_bp"] = analysed_aas.apply(lambda x: x.aromaticity())

# %% [code]
sns.distplot(unip_sub["Aromaticity_bp"].dropna())
plt.show()

# %% [markdown]
### Grand Average of Hydropathicity (GRAVY)

The GRAVY value for a protein or a peptide is calculated by adding the hydropathy values (Kyte, J. and Doolittle, R.F., 1982) of each amino acid residues and dividing by the number of residues in the sequence or length of the sequence. Increasing positive score indicates a greater hydrophobicity.

# %% [code]
unip_sub["GRAVY_bp"] = analysed_aas.apply(lambda x: x.gravy())

# %% [code]
sns.distplot(unip_sub["GRAVY_bp"].dropna())
plt.show()

# %% [markdown]
### Instability Index

The instability index provides an estimate of the stability of your protein in a test tube. A protein whose instability index is smaller than 40 is predicted as stable, a value above 40 predicts that the protein may be unstable.

# %% [code]
unip_sub["Instability_index_bp"] = analysed_aas.apply(lambda x: x.instability_index())

# %% [code]
sns.distplot(unip_sub["Instability_index_bp"].dropna())
plt.show()

# %% [code]
unip_sub.head(3)

# %% [markdown]
### Identifying PEST regions

A PEST sequence is a peptide sequence that is rich in proline (P), glutamic acid (E), serine (S), and threonine (T). This sequence is associated with proteins that have a short intracellular half-life; therefore, it is hypothesized that the PEST sequence acts as a signal peptide for protein degradation.

This protein degradation may be mediated via the proteasome or calpain.

Other signals thought to identify proteins for degradation include cyclin destruction boxes, which are amino acid sequences that mark cell-cycle proteins for destruction. 

We will do this on the fly, since PEST regions can only be generated one at a time.

In order to use `epestfind`, you must install the `emboss` suite as:

    sudo apt install emboss
    
Then we will call epestfind with a number of parameters:

    !epestfind -sequence {pest_file}test.sw -window 10 -order score -outfile {pest_file}test.epestfind -graph none -filter -nomap -sid1 Q16827
    
whereby:

- window: window size of search , default 10
- sequence: input amino acid sequence
- order: how to order results
- outfile: output file to analyse.
- graph: we dont want to draw a graph so set to none
- filter: filter out stdout
- nomap: we dont want the mapping in the output file
- sid1: set the name to our uniprot ID

# %% [code]
import numpy as np

def calculate_external_PEST(uniprot, output_dir="../../data/PEST/"):
    result_list = []
    i = 0
    # iterate over rows
    for unip_id, row in uniprot.iterrows():
        if i % 1000 == 0:
            print("[Progress: {}/{}]".format(i,len(uniprot)))
        # create an input file
        with open(output_dir+"input.sw","w") as f:
            f.write(row.Seq)
        # call epestfind
        !epestfind -sequence {output_dir}input.sw -window 10 -order score -outfile {output_dir}output.epestfind -graph none -filter -nomap -sid1 {unip_id}
        # remove input file
        !rm {output_dir}input.sw
        # analyse output file
        with open(output_dir+"output.epestfind","r") as f:
            outfile = pd.Series(f.readlines())
        # get total number of PESTs
        n_PESTs = re.findall("^\s+([0-9]+) PEST motif", outfile[2])
        # if we have a result, then select it, else set to 0
        if len(n_PESTs) > 0:
            n_PESTs = n_PESTs[0]
            # find poor and potential PEST rows.
            pest_rows = outfile.str.findall("^(?P<PEST_cat>Poor|Potential) PEST motif with (?P<PEST_aa>[0-9]+)")
            pest_scores = outfile.str.findall("\s+PEST score: (?P<PEST_score>[0-9\-.]+)")
            # preprocess away junk
            refined_row = pest_rows[pest_rows.apply(lambda x: len(x)>0)].apply(lambda x: x[0])
            refined_score = pest_scores[pest_scores.apply(lambda x: len(x)>0)].apply(lambda x: x[0]).astype(np.float)
            # merge together
            newset = refined_row.apply(lambda x: pd.Series(x, index=["type","len"])).join(
                pd.Series(refined_score.values, index=refined_row.index, name="score")
            )
            # get counts of each type
            val_counts = newset.type.value_counts()
            # they may not exist in the value counts, so we protect with try/catch
            try:
                n_potent = val_counts.loc["Potential"]
                avg_potent = newset.groupby("type").score.mean().Potential
            except KeyError:
                n_potent = np.nan
                avg_potent = np.nan
            try:
                n_poor = val_counts.loc["Poor"]
            except KeyError:
                n_poor = np.nan
            # append a pd.Series with uniprot id, total pest, potential and poor, with average potential PEST score.
            result_list.append(pd.Series((unip_id, n_PESTs, n_potent, n_poor, avg_potent),
                             index=["Uniprot_ID","PEST_total","PEST_potential","PEST_poor","PEST_avg_potential_score"]))

            # free up memory and disk
            !rm {output_dir}output.epestfind
            del pest_rows
            del pest_scores
            del refined_row
            del refined_score
            del newset
            del outfile
        else:
            result_list.append(
                pd.Series((unip_id, np.nan, np.nan, np.nan, np.nan),
                         index=["Uniprot_ID","PEST_total","PEST_potential",
                                "PEST_poor","PEST_avg_potential_score"])
            )
        i += 1
        
    return result_list

# %% [code]
if not os.path.isfile("../../data/PEST/pests.csv"):
    # create pest list
    pest_list = calculate_external_PEST(uniprot)
    # create DF.
    pests = pd.concat(pest_list, axis=1).T.set_index("Uniprot_ID")
    del pest_list
    pests.to_csv("../../data/PEST/pests.csv")
else:
    pests = pd.read_csv("../../data/PEST/pests.csv", index_col=0)

# replace 0-pests with np.nan, we don't want 0s to skew our distribution.
pests = pests.mask(pests.eq(0))

pests.head(2)

# %% [code]
pests = pests.drop(["PEST_avg_potential_score","PEST_potential","PEST_poor"], axis=1)

# %% [code]
unip_sub = unip_sub.join(pests.add_suffix("_ptmc"), how="left")

# %% [code]
unip_sub.shape

# %% [markdown]
### Post-Translational Modifications

We will make use of *PhosphoSitePlus*, a database containing phosphorylation, methylation, ubiquitination and other forms of modifications.

We must reference this paper to use PTM data:

    “Hornbeck PV, Zhang B, Murray B, Kornhauser JM, Latham V, Skrzypek E PhosphoSitePlus, 2014: mutations, PTMs and recalibrations. Nucleic Acids Res. 2015 43:D512-20.”
    


# %% [code]
phospho_link = "../../data/PhosphoSitePlus/"

# %% [code]
# unzip all files.
!gunzip -k {phospho_link}*

# %% [code]
# get file names.
filenames = !ls {phospho_link}*_site_dataset
filenames

# %% [code]
ptms = [pd.read_csv(f, sep="\t", header=2) for f in filenames]

# %% [markdown]
### PTM counts for each gene, per set.

We can see that phosphorylation is dominant, with ubiquitination then acetylation coming third.

# %% [code]
ptm_counts = pd.concat([ptm.query("ORGANISM == 'human'").groupby("ACC_ID").GENE.count() for ptm in ptms],
                   axis=1, sort=True)
ptm_counts.columns = [re.findall("../../data/PhosphoSitePlus/([A-Za-z\-]+)\_site\_dataset", f)[0] for f in filenames]

# %% [markdown]
Fill missing values with 0

# %% [code]
ptm_counts.fillna(0, inplace=True)

# %% [code]
print(ptm_counts.shape)
ptm_counts.notnull().sum()

# %% [code]
# remove files
!rm {phospho_link}*_site_dataset

# %% [code]
unip_sub = unip_sub.join(ptm_counts.add_suffix("_ptmc"), how="left")

# %% [code]
unip_sub.head(2)

# %% [markdown]
#### adding 'boolean counter amino acid' (bcaa) flag

# %% [code]
unip_sub = unip_sub.join(feature_table.add_suffix("_bcaa"), how="left")

# %% [markdown]
### Clean and remove sequence, Save the count-set to file

# %% [code]
unip_sub.drop(["Seq"], axis=1, inplace=True)
unip_sub.to_csv("../../datasets/uniprot_unscaled.csv", sep="\t")

# %% [markdown]
### Plotting the entire dataset

# %% [code]
_ = mf.dataset_hist(unip_sub, normaltest=False)

# %% [markdown]
## Normalization strategy

1. **Length based features**: Scale by $\log_2(x+1)$.
2. **Amino acid counts**: Scale by amino acid length
3. **Post-translational modifcation counts (including PEST)**: Scale by amino acid length
4. **Isoelectric point**: Centered to zero by 7.
5. **Instability Index**: Centered to zero by 40. 40 is the threshold at which above indicates instable.

# %% [code]
string_groups = ["^length_", "_aa$", "pH_Isoelectric_point", 
                 "Instability_index", "_ptmc$"]

operations = [lambda x: np.log2(x+1), lambda x: x / unip_sub["length_aa"],
              lambda x: x - 7., lambda x: x - 40., lambda x: x / unip_sub["length_aa"]]

unip_normed = misc.fchain_args(unip_sub, 
                            [misc.apply_column_function]*len(string_groups), 
                            list(zip(string_groups, operations)))

# %% [code]
unip_normed.head()

# %% [code]
_ = mf.dataset_hist(unip_normed, normaltest=False)

# %% [code]
unip_normed.to_csv("../../datasets/uniprot_proc_freq.csv", sep="\t")
```
