## Supplementary material for "Predicting Relative Protein Abundance via Sequence-Based Information": S3 Data: Vogel-RNA-Protein.ipynb

```
# %% [markdown]
## RNA-Protein Preprocessing with Christine Vogel's dataset

In a paper entitled: **Sequence signatures and mRNA concentration can explain two-thirds of protein abundance variation in a human cell line**, which can be found [here](https://www.ncbi.nlm.nih.gov/pmc/articles/PMC2947365/):

### Abstract

Transcription, mRNA decay, translation and protein degradation are essential processes during eukaryotic gene expression, but their relative global contributions to steady-state protein concentrations in multi-cellular eukaryotes are largely unknown. Using measurements of absolute protein and mRNA abundances in cellular lysate from the human Daoy medulloblastoma cell line, we quantitatively evaluate the impact of mRNA concentration and sequence features implicated in translation and protein degradation on protein expression. Sequence features related to translation and protein degradation have an impact similar to that of mRNA abundance, and their combined contribution explains two-thirds of protein abundance variation. mRNA sequence lengths, amino-acid properties, upstream open reading frames and secondary structures in the 5′ untranslated region (UTR) were the strongest individual correlates of protein concentrations. In a combined model, characteristics of the coding region and the 3′UTR explained a larger proportion of protein abundance variation than characteristics of the 5′UTR. The absolute protein and mRNA concentration measurements for >1000 human genes described here represent one of the largest datasets currently available, and reveal both general trends and specific examples of post-transcriptional regulation.

The following datasets are extracted from the supplementary materials of this paper.

### Cell line

- **Daoy**: medulloblastoma

# %% [code]
import numpy as np
import pandas as pd
import matplotlib.pyplot as plt
import sys
sys.path.insert(0,"../../../turbopanda")
import turbopanda as turb
%matplotlib inline

# %% [code]
ds = turb.read("../../data/vogel_dataset_rna-prot.xls", name='Vogel', header=9,sheet_name="Data")
ds

# %% [code]
cleaner = turb.Pipe(
    ['keep', "Ensembl", "Protein_Expression", "mRNA_Expression"],
    ['transform', np.log2, ("Protein_", 'mRNA_')],
    ['rename_axis', [("Expression","expr"),('_(Levels|levels)',''),
                ('molecules','mol'),('Ensembl','Ens'), ("Protein",'prot')]],
    ['rename_axis', [('prot_expr_', 'prot_'),
                  ('mRNA_expr_', 'mRNA_'),
                  ('\(mol/cell\)', 'Daoy'),
                  ('\(arbitrary_units\)', 'Daoy'),
                  ('(', ''), (')','')]],
)

# %% [code]
ds.compute(cleaner, inplace=True)
ds.head()

# %% [code]
ds.write("../../datasets/vogel_all.csv")

# %% [markdown]
## Sanity and Interest Plots

# %% [code]
plt.scatter(ds["mRNA_"], ds['prot_'],marker='x')
plt.xlabel(r"$log_2$ mRNA expression (arbitrary units)")
plt.ylabel(r"$log_2$ Protein expression (molecules/cell)")
plt.show()

# %% [code]
```
