## Supplementary material for "Predicting Relative Protein Abundance via Sequence-Based Information": S3 Data: README.md

[Owned by Gregory Parkes, Ph.D, University of Southampton, UK, 2016+] 
[Last Access: 10 Dec 2020]
[All Rights Reserved]

# Supplementary Material 3: Data and Code

We deploy a series of jupyter notebooks to handle the downloading, preprocessing
 and plotting of data, codes and so on. Scripts associated will be contained in Supplementary Material 4.

See https://github.com/gregparkes/sdf_paper for details on how to work with Jupyter notebooks. See https://anaconda.org/ for installing and working with the Python programming language.

Note that Jupyter notebooks can be converted into HTML and PDF formats if so desired. However this leads them into an un-executable state, so we have kept them in their original .ipynb format (JSON) for maximum accessibility. 

See below for what is contained in each subfolder:

### Data Access

Here we handle notebooks concerned with NCBI and Uniprot access.

- **NCBI-Access**: This notebook is concerned with retrieval and basic preprocessing 
such as dropping experimental transcript variants of NCBI mRNA transcript data 
from genbank format. We make extensive use of `biopython` package in order to handle our 
data. This is then outputted into a folder in `../data/NCBI/` into a variety of 
files, including feature counts.
- **Uniprot-Access**: This notebook is concerned with the retrieval of Uniprot
 data using a custom URL to select appropriate columns from the website. 
 See the [Uniprot website](https://www.uniprot.org/). We perform minimal processing on
  the file, including dropping an irrelevant column and changing some of the 
  column names. This is then dumped in `../datasets/Uniprot/` subfolder ready 
  to be preprocessed and integrated.

### Preprocessing

This subset is concerned with handling sequence data from NCBI and 
Uniprot, alongside datasets we require and preprocessing them into an 
intelligable form.

- **Aviner-Microarray**: This notebook is concerned with preprocessing and loading 
the original microarray dataset that *Aviner et al.* uses rather than the curated
 one he provides in his 2015 paper. This original set has a whopping 32K samples,
  with 7 timesteps and 3 replicates per timestep, totalling 21 samples. This
   notebook downloads and/or inputs from `../data/microarray/GSE26922_family.soft.gz`
    and the output of this notebook can be found in `../datasets/microarray_proc.csv.gz`. 
  You can access this yourself by downloading GSE26922 from GEO website.
- **NCBI-Preprocessing**: This notebook handles the preprocessing NCBI genbank 
files containing the mRNA sequence and coding sequence (CDS) of every known 
curated transcript. The primary ID is REFSEQ IDs. We extract CDS, 5'-UTR and
 3'-UTR features from their respective sequences in a long format, alongside 
 combining our feature count matrix data into a curated dataset to be placed 
 in `..datasets/`. 
- **Uniprot-Preprocessing**: This notebook handles the preprocessing of 
Uniprot amino acid sequence files by extract amino-acid related features
 from the AA sequence of every known transcript. The primary ID is 
 UNIPROT IDs. Here we integrate PEST data, Post-translational modifications
  (PTMs) and amino-acid features known directly by Uniprot. 
- **HGNC-Preprocessing**: This notebook handles basic preprocessing of 
the HGNC labels which help to map across multiple datasets. This is
 essential in SDF formation between Refseq/Uniprot sources.
- **Vogel-RNA-Protein**: This notebook handles the preprocessing of
 the *Vogel et al.* 2010 paper data source of Daoy cell line data.
  This dataset also includes a SDF-like processed dataset benchmark to 
  compare/contrast against.
- **Lundberg-RNA-Protein**: This notebook handles the preprocessing of the 
*Lundberg et al.* 2010 paper data source of U2)S, UMG512 and A431 cell 
lines dataset. They contain many mRNA and protein pair samples.

### Sequence-derived Features (SDF)

Here we take preprocessed sequence-derived features from NCBI and Uniprot and perform analyses on them for the SDF paper/dataset.

- **SDF-Create-SDF**: This notebook is concerned with integrating the results
 from `NCBI-Preprocessing` and `Uniprot-Preprocessing` to create a fully-fledged 
 sequence-derived feature dataset. This involves cleaning, eliminating useless 
 features, converting features by scaling, and mapping against HGNC. We also 
 sort the features by their logical groups.
- **SDF-Sanity**: A notebook for drawing sanity plots, including normalization strategies and distribution fitting between SDF features.
- **SDF-Official[n]**: Each of these notebooks is responsible for at least 
one major Figure to be displayed in the paper, along with potential 
supplementary material related to the figure.
- **SDF-Base-Frequency**: Measures of the profiles for mononucleotide and dinucleotide bases
along the gene length and the analysis of these profiles as supplementary material for 
the SDF paper.
***

Ensure that any use of this material is appropriately referenced and in compliance 
with the license.

Gregory Parkes, All Rights Reserved, 2019-.
