## Supplementary material for "Predicting Relative Protein Abundance via Sequence-Based Information": S3 Data: SDF-Create-DF.ipynb

```
# %% [markdown]
# Creating the Dataset

This notebook is purely designed to integrate together the datasets that come out of `NCBI-Preprocessing` and `Uniprot-Preprocessing` notebooks and create a sequence-derived feature dataset.

Files generated: `refseq_mRNA_scaled.csv` and `uniprot_proc_freq.csv` from each respective preprocessing file. We rely on `hgnc_labels.csv` to map together the refseq and uniprot labels.

1. [Import and clean mRNA dataset](#s1)
2. [Eliminating features in mRNA based on criteria](#s2)
3. [Import and clean Uniprot dataset](#s3)
4. [Eliminating features in Uniprot based on missing data](#s4)
7. [Integration with HGNC](#s5)
8. [Sort out the order of features by groups](#s6)
9. [Associating colors to metadata](#s7)
9. [Saving File](#s8)

# %% [code]
import numpy as np
import pandas as pd
import sys
sys.path.insert(0, "../../../turbopanda/")
import turbopanda as turb

# %% [markdown]
## <a id='s1'> Stage 1. Import and clean the mRNA (scaled by length) dataset. </a>

# %% [code]
rna_unsc = turb.read("../../datasets/refseq_mRNA_unscaled.csv",name="Refseq_un")
rna_unsc.rename_axis([('Unnamed: 0', "Refseq_ID")])
rna_unsc.apply("drop_duplicates")

# %% [markdown]
## <a id='s2'> Stage 2. Eliminate features based on criteria </a>

1. Features with fewer than 50% actual data. (50% missing)
2. Features with cronically low variance (i.e all 0), $\sigma^2 < 10^{-7}$

# %% [code]
missing_50 = lambda df: (df.count() / df.shape[0]) < 0.5
low_var = lambda df: df.var() < 1e-7 if df.dtype != object else False

# %% [code]
# perform drop
rna_unsc.drop(missing_50, low_var)

# %% [markdown]
## <a id='s3'> Stage 3. Import and clean the Uniprot dataset. </a>

# %% [code]
prot_unsc = turb.read("../../datasets/uniprot_unscaled.csv","Uniprot_un",sep="\t")
prot_unsc.apply("drop_duplicates", "Uniprot_ID")

# %% [markdown]
## <a id='s4'> Stage 4. Eliminate uniprot features based on criteria </a>

1. Features with fewer than 50% actual data. (50% missing) But there are none!

# %% [code]
prot_unsc.view(missing_50)

# %% [markdown]
## <a id='s5'> Stage 5. Integration via HGNC </a>

In order to merge together the labels, we are going to bring in HGNC as a unifying database.

We have some additional steps to take:

1. Import the dataset
2. Clean the column labels
3. Expand out Uniprot and Refseq datasets into long-form
4. Merge using 'inner', keeping rows that only exist in both.

# %% [code]
import os

# %% [code]
hgnc = turb.read("../../data/HGNC/HGNC_clean.csv", name="HGNC",
                 index_col=0)
hgnc

# %% [markdown]
### Merging HGNC, Uniprot and Refseq

# %% [code]
sdf = turb.merge([hgnc, rna_unsc, prot_unsc], name="SDF_unsc", verbose=2)

# %% [markdown]
## <a id='s6'> Stage 6. Sorting out the groupings of features </a>

There are 138 features, including labels. They break down into these categories:

1. HGNC (labels): 11
2. Refseq (mRNA-like features): 93
3. Uniprot (amino-acid-like features): 34

Refseq features also break down into 3'UTR, 5'UTR and CDS regions, mostly.

They also break down into types:

1. ID feature (label)
1. Length
2. Mononucleotide frequency
3. Dinucleotide frequency
4. Free energy
5. GC content
6. Codon bias (RCBS, tAI, CAI)
7. Feature counts
8. Amino acid properties
9. post-translational modifications

# %% [markdown]
### Regular expressions to group features

`turbopanda` caching associations first.

object types are now converted to string to enable encoding by JSON.

# %% [code]
caches = dict(
    aa_freq="^[A-Z]_aa$", 
    id_col=(object, "_ID$", "_id$"),
    mononuc_freq="^[AGCT]{1}_(?:mrna|cds|utr5|utr3)$",
    dinuc_freq="^[AGCT]{2}_(?:cds|utr3|utr5|mrna)$",
    spec_freq="GC_content",
    length="length_(?!prop)",
    l_prop="length_prop_(?:cds|utr5|utr3)",
    free_energy="MFE_?",
    codon_bias="tAI|CAI|RCB|uORF",
    translation_freq="^kozak|M_(?!aa)",
    rna_misc="fc|bs$",
    feat_count="_fc$",
    ptm="_ptmc$",
    aminoacid_meta="_bcaa$",
    biophysical="_bp$",
    hgnc="id_col",
    di_mrna="^[AGCT]{2}_mrna$",
    di_cds="^[ATCG]{2}_cds$",
    di_utr5="^[ATCG]{2}_utr5$",
    di_utr3="^[ATGC]{2}_utr3$",
    mono_mrna="^[AGCT]{1}_mrna$",
    mono_cds="^[ATCG]{1}_cds$",
    mono_utr5="^[ATCG]{1}_utr5$",
    mono_utr3="^[ATGC]{1}_utr3$",
    uniprot=("_bcaa$", "_aa$", "_bp$"),
    refseq=("_utr5$","_utr3$","_cds$","_mrna$","^MFE",
              "tAI|CAI|RCB|uORF","_fc$","_bs$"),
    ptm_source="ptm",
    mrna=("_mrna$", "_fc$"),
    cds=("_cds$", "CAI|tAI|RCB", "ccds"),
    utr5=("_utr5$","^MFE"),
    utr3="_utr3$",
    protein=("_aa$", "_bp$", "_bcaa$", "_ptmc$"),
    mrna_f="_mrna$",
    cds_f="_cds$",
    utr5_f="^(?!uORF).*_utr5$",
    aa_f="_aa$",
    exc_gc="^[CG]{1,2}_(?:mrna|cds|utr5|utr3)$",
    exc_at="^[AT]{1,2}_(?:mrna|cds|utr5|utr3)$",
    base_mix="^([AT][CG]|[CG][AT])_(?:mrna|cds|utr5|utr3)$"
)

# %% [code]
sdf.cache_k(**caches)

# %% [markdown]
#### The ordering of Central Dogma and Data Groups...

The order goes as mRNA -> UTR3 -> UTR5 (free energy) -> CDS -> Amino acid -> PTM

# %% [code]
order_dogma = ("mrna_f", "rna_misc", "utr3",
    "utr5_f", "free_energy", "cds_f", "codon_bias", 
    "aa_f", "aminoacid_meta", "biophysical", "ptm_source")

# re order data_groups - which kind of represents the 'type' of feature we have also.
order_data_groups = (
    "spec_freq","mononuc_freq","dinuc_freq","length","l_prop","translation_freq","free_energy",
    "rna_misc","codon_bias","aa_freq","aminoacid_meta","biophysical","ptm"
)

base_pairing_order = ("exc_at", "exc_gc", "base_mix")

# %% [markdown]
#### Mapping these caches into meta_ features


# %% [code]
sdf.meta_map("data_region", ("mrna","cds","utr5","utr3","protein"))
sdf.meta_map("data_source", ("hgnc","refseq","uniprot","ptm_source"))
sdf.meta_map("data_groups", order_data_groups)
sdf.meta_map("dogma_order", order_dogma)
sdf.meta_map("base_pairing", base_pairing_order)

# %% [markdown]
#### Cleaning pipeline...

# %% [code]
rn_corr = lambda axis: ["rename_axis", [("pH_",""), ("_prop"," rel."),
        ("_cds"," CDS"),("_utr5"," 5'UTR"), ("_utr3"," 3'UTR"),
        ("_win"," (win"), ("_total",""), ("_bp",""), ("_bcaa",""),
        ("mfe","free energy"), ("_mrna"," mRNA"), ("_aa", " (amino acid)"),
        ("_ptmc", " PTM"), ("_fc",""),("_pc"," (corrected)"), ("_"," ")],
      "axis={}".format(axis)]

clean_and_rename = turb.Pipe(
    ["apply_columns", "lower"],
    rn_corr(1)
)

# %% [markdown]
## Designing 'Pretty' Table

..using the meta data provided!

# %% [code]
cleaned = sdf.copy()

# %% [code]
cleaned.apply_columns("lower")
cleaned.rename_axis([("pH_",""), ("_prop"," rel."),
        ("_cds"," CDS"),("_utr5"," 5'UTR"), ("_utr3"," 3'UTR"),
        ("_win"," (win"), ("_total",""), ("_bp",""), ("_bcaa",""),
        ("mfe","free energy"), ("_mrna"," mRNA"), ("_aa", " (amino acid)"),
        ("_ptmc", " PTM"), ("_fc",""),("_pc"," (corrected)"), ("_"," ")])

# %% [code]
sdf.meta_['Column Clean Name'] = cleaned.meta_.index

# %% [code]
sdf.meta_.sort_index(axis=1).reset_index().to_csv("../../datasets/FULL_SDF_META.csv", index=None)

# %% [code]
sdf.drop("^counter", "Unnamed")

# %% [markdown]
## Transformation of Features

| **Feature** | **Normalization 1** | **Normalization 2** |
| ------------- | --------------------------- | --------------------- |
| *Free energy* | Length of window | Length of window |
| *Dinucleotide count* | See equation 1 | See equation 2 |
| *Mononucleotide count* | See equation 1 | See equation 2 |
| *Amino acid count* | See equation 1 | See equation 2 |
| *Isoelectric point* | $x - 7$ | $x-7$ |
| *Instability Index* | $x-40$ | $x-40$ |
| *kozak* | CDS length | CDS length |
| *mRNA feature counts* | mRNA length | mRNA length |
| *PTM* | AA length | AA length |
| *Length* | $\log(x+1)$ | $\log(x+1)$ |

### Version 1

We normalize the features by their **respective length**, for example, MFE features are normalized by sliding window, base counts normalized by the length of the gene, etc.

$$
\begin{equation}
\hat{f}_i=\frac{f_i}{L_i}
\label{eq:sc1}
\tag{1}
\end{equation}
$$

# %% [code]
sdf_unsc1 = sdf.copy()
sdf_unsc1.transform_k([
        *[(lambda x: -(x / d), "MFE_win%d" % d) for d in [10,20,30,40,60,80]],
        (lambda x: x / sdf.df_["length_mrna"], ("mono_mrna","feat_count")),
        (lambda x: x / (sdf.df_["length_mrna"] / 2.), "di_mrna"),
        (lambda x: x / sdf.df_["length_cds"], ("mono_cds","kozak")),
        (lambda x: x / (sdf.df_["length_cds"] / 2.), "di_cds"),
        (lambda x: x / sdf.df_["length_utr5"], ("mono_utr5", "uORF_utr5")),
        (lambda x: x / (sdf.df_["length_utr5"] / 2.), "di_utr5"),
        (lambda x: x / sdf.df_["length_utr3"], "mono_utr3"),
        (lambda x: x / (sdf.df_["length_utr3"] / 2.), "di_utr3"),
        (lambda x: -(x / sdf.df_["length_utr5"]), "MFE"),
        (lambda x: x / (sdf.df_["length_utr5"] / 3), "M_utr5"),
        (lambda x: x / sdf.df_["length_aa"], ("aa_freq","ptm_source")),
        (lambda x: x - 40, "Instability"),
        (lambda x: x - 7, "Isoelectric"),
        (lambda x: np.log2(x+1), "length"),
        ])

sdf_unsc1.update_meta()

# %% [markdown]
### Version 2

If each frequency $f_{ij}$ belongs to an amino acid $j \in [1, 20]$ for sequence $i$, then each amino acid $j$ has an expected, unbiased frequency:

$$
\mathbb{E}_j^{\text{AA}}=\frac{1}{20}
$$

likewise for mRNA base $k \in [1, 4]$, the expectation is $\mathbb{E}_k^{\text{mRNA}}=1/4$, and for dinucleotide pair $m \in [1, 16]$, we have $\mathbb{E}_m^{\text{mRNA}}=1/32$. Thus to normalize *base counts* by the **relative frequency** from the expected value, for example an amino acid frequency. So given count $f$, we would calculate an updated value $\hat{f}$ as:

$$\begin{equation}
\hat{f}_{ij}=\frac{f_{ij}}{\mathbb{E}_j^{\text{AA}} L_i} - 1
\label{eq:sc2}
\tag{2}
\end{equation}$$

where $\mathbb{E}_j^{\text{AA}}$ is the unbiased frequency of that amino acid drawn uniformly from all other amino acids. Since there are 20 amino acids, this is $\frac{1}{20}$, factoring in gene/mRNA length $L$. This means that values of $\hat{f} \equiv 0$ are where there is no deviation from $\mathbb{E}^{\text{AA}} L_i$, positive values suggest above-expectation and negative values suggest below-expectation. Because it is centered around 0 we would expect normal(ish) distributions...

Values to scale the length by:

1. Mononucleotides: $\frac{1}{4}=0.25$
2. Dinucleotides: $\frac{1}{32}=0.0625$
3. Amino acids: This is a special case as we have to factor in the coding table, i.e certain amino acids have multiple trinucleotides encoding it.

In the dinucleotide case, because our *count data* counts *pairs*, we have to factor that into our length calculation.

This is nice because we can see that we get smoother distributions by combining this *expected value*.

In addition, with counts following a data generation process, we will try square-root/Freeman-tukey transformation?

# %% [code]
codon_chart = {'phe': ['uuu', 'uuc'], 'leu': ['uua', 'uug', 'cuu', 'cuc', 'cua', 'cug'],
               'ile': ['auu', 'auc', 'aua'], 'val': ['guu', 'guc', 'gua', 'gug'],
               'met': ['aug'], 'ser': ['ucu', 'ucc', 'uca', 'ucg', 'agu', 'agc'],
               'pro': ['ccu', 'ccc', 'cca', 'ccg'], 'thr': ['acu', 'acc', 'aca', 'acg'], 
               'ala': ['gcu', 'gcc', 'gca', 'gcg'], 'tyr': ['uau', 'uac'], 
               'stop': ['uaa', 'uag', 'uga'], 'his': ['cau', 'cac'],
               'gln': ['caa', 'cag'], 'asn': ['aau', 'aac'], 'lys': ['aaa', 'aag'],
               'asp': ['gau', 'gac'], 'glu': ['gaa', 'gag'], 'cys': ['ugu',' ugc'],
               'trp': ['ugg'], 'arg': ['cgu', 'cgc', 'cga', 'cgg', 'aga', 'agg'],
               'gly': ['ggu', 'ggc', 'gga', 'ggg']
               }

# %% [code]
aa_abbr = {'A': 'ala', 'C': 'cys', 'D': 'asp', 'E': 'glu', 'F': 'phe', 'G': 'gly', 'H': 'his', 
           'I': 'ile', 'K': 'lys', 'L': 'leu', 'M': 'met', 'N': 'asn', 'P': 'pro', 'Q': 'gln',
           'R': 'arg', 'S': 'ser', 'T': 'thr', 'V': 'val', 'W': 'trp', 'Y': 'tyr'}

n_synonomous = pd.Series({a +"_aa": len(codon_chart[b]) for a, b in aa_abbr.items()})


# %% [markdown]
Expected values for each amino acid are as follows then
 the uniform distribution across all trinucleotides.

# %% [code]
expect_synon = n_synonomous/64

# %% [markdown]
Scaler 2 here

# %% [code]
scaler_2 = [
    ('transform_k',
      ([
        *[(lambda x: -(x / d), "MFE_win%d" % d) for d in [10,20,30,40,60,80]],
        (lambda x: (x / (0.25*sdf.df_["length_mrna"])) - 1, "mono_mrna"),
        (lambda x: (x / (0.0625*0.5*sdf.df_["length_mrna"])) - 1, "di_mrna"),
        (lambda x: (x / (0.25*sdf.df_["length_cds"])) - 1, "mono_cds"),
        (lambda x: (x / (0.0625*0.5*sdf.df_["length_cds"])) - 1, "di_cds"),
        (lambda x: (x / (0.25*sdf.df_["length_utr5"])) - 1, "mono_utr5"),
        (lambda x: (x / (0.0625*0.5*sdf.df_["length_utr5"])) - 1, "di_utr5"),
        (lambda x: (x / (0.25*sdf.df_["length_utr3"])) - 1, "mono_utr3"),
        (lambda x: (x / (0.0625*0.5*sdf.df_["length_utr3"])) - 1, "di_utr3"),
        (lambda x: x / sdf.df_["length_cds"], "kozak"),
        (lambda x: x / sdf.df_["length_mrna"], "feat_count"),
        (lambda x: -(x / sdf.df_["length_utr5"]), "MFE"),
        (lambda x: (x / ((1/64)*(1/3)*sdf.df_["length_utr5"])) - 1, "M_utr5"),
        (lambda x: x / sdf.df_['length_utr5'], "uORF_utr5"),
        # scaling factor subject to amino acid expected value type.
        (lambda x: (x / (expect_synon[x.name]*sdf.df_["length_aa"])) - 1, "aa_freq"),
        (lambda x: x - 40, "Instability"),
        (lambda x: x - 7, "Isoelectric"),
        (lambda x: x / sdf.df_["length_aa"], "ptm_source"),
        (lambda x: np.log2(x+1), "length"),
        ],
      ),
      {}),
]

# %% [code]
sdf_unsc2 = sdf.copy()
sdf_unsc2.transform_k([
        *[(lambda x: -(x / d), "MFE_win%d" % d) for d in [10,20,30,40,60,80]],
        (lambda x: (x / (0.25*sdf.df_["length_mrna"])) - 1, "mono_mrna"),
        (lambda x: (x / (0.0625*0.5*sdf.df_["length_mrna"])) - 1, "di_mrna"),
        (lambda x: (x / (0.25*sdf.df_["length_cds"])) - 1, "mono_cds"),
        (lambda x: (x / (0.0625*0.5*sdf.df_["length_cds"])) - 1, "di_cds"),
        (lambda x: (x / (0.25*sdf.df_["length_utr5"])) - 1, "mono_utr5"),
        (lambda x: (x / (0.0625*0.5*sdf.df_["length_utr5"])) - 1, "di_utr5"),
        (lambda x: (x / (0.25*sdf.df_["length_utr3"])) - 1, "mono_utr3"),
        (lambda x: (x / (0.0625*0.5*sdf.df_["length_utr3"])) - 1, "di_utr3"),
        (lambda x: x / sdf.df_["length_cds"], "kozak"),
        (lambda x: x / sdf.df_["length_mrna"], "feat_count"),
        (lambda x: -(x / sdf.df_["length_utr5"]), "MFE"),
        (lambda x: (x / ((1/64)*(1/3)*sdf.df_["length_utr5"])) - 1, "M_utr5"),
        (lambda x: x / sdf.df_['length_utr5'], "uORF_utr5"),
        # scaling factor subject to amino acid expected value type.
        (lambda x: (x / (expect_synon[x.name]*sdf.df_["length_aa"])) - 1, "aa_freq"),
        (lambda x: x - 40, "Instability"),
        (lambda x: x - 7, "Isoelectric"),
        (lambda x: x / sdf.df_["length_aa"], "ptm_source"),
        (lambda x: np.log2(x+1), "length"),
        ])
sdf_unsc2.update_meta()

# %% [markdown]
### The size of the dataset: (19198, 206)

# %% [markdown]
## Construction of the Supplementary 1 Table:

Here the columns we want are:

* Name of the feature
* Shorthand name of the feature
* Groups it belongs to
* Normalization to it

# %% [code]
supp1 = (sdf_unsc2.meta_
    .copy()
    .drop(["dogma_order","base_pairing","is_mixed_type","true_type","is_unique_id"], axis=1)
    .reset_index()
    .rename(columns={'index': 'codename'}))

# %% [code]
group_to_full = {
    "hgnc": "",
    "id_col": "ID Columns",
    "ptm": "Post-Translational Modification",
    "codon_bias": "Codon Bias",
    "dinuc_freq": "Dinucleotide Frequency",
    "l_prop": "Proportional Length",
    "length": "Sequence Length",
    "mononuc_freq": "Mononucleotide Frequency",
    "rna_misc": "RNA Miscallaenous Feature",
    "spec_freq": "Special Sequence Frequency",
    "aa_freq": "Amino Acid Frequency",
    "biophysical": "Amino Acid Biophysical Property",
    "translation_freq": "Translation Frequency",
    "free_energy": "Minimum 5'UTR Free Energy",
    "aminoacid_meta": "Amino Acid Text-Mined Feature"
}

source_to_full = {
    "refseq": "Refseq/Ensembl",
    "uniprot": "Uniprot/Swissprot",
    "hgnc": "HGNC",
    "ptm_source": "PhosphoSitePlus" 
}

col_renames = {'name': 'Feature', 'codename': 'Feature (Code)', 
               'data_region': 'Feature Type', 'data_source': 'Source', 
               'data_groups': 'Labelled Group'}

pretty_names = [("pH_",""), ("_prop"," rel."),
        ("_cds"," CDS"),("_utr5"," 5'UTR"), ("_utr3"," 3'UTR"),
        ("_win"," (win"), ("_total",""), ("_bp",""), ("_bcaa",""),
        ("mfe","free energy"), ("_mrna"," mRNA"), ("_aa", " (amino acid)"),
        ("_ptmc", " PTM"), ("_fc",""),("_pc"," (corrected)"), ("_"," ")]

# %% [markdown]
#### Make changes here using our dictionaries

# %% [code]
supp1['name'] = turb.str.string_replace(supp1['codename'], pretty_names)
supp1['data_source'] = supp1['data_source'].replace(source_to_full)
supp1['data_groups'] = supp1['data_groups'].replace(group_to_full)
supp1['data_region'] = supp1['data_region'].str.capitalize()
# reorder columns
supp1 = supp1[['name','codename','data_source','data_groups','data_region']]
# rename columns
supp1 = supp1.rename(columns=col_renames)

# %% [markdown]
#### Write out

# %% [code]
supp1.to_excel("../../datasets/features.xlsx", index=None)

# %% [markdown]
## <a id='s8'> Writing to file </a>

We will save the `.csv` file and a `.json` file.

# %% [code]
sdf.write("../../datasets/SDF_unscaled.json")

# %% [markdown]
#### Associate transformation pipes to unsc?

# %% [code]
sdf_unsc1.downcast()
sdf_unsc1.write("../../datasets/SDF_scaled1.json")

# %% [code]
sdf_unsc2.downcast()
sdf_unsc2.write("../../datasets/SDF_scaled2.json")

# %% [markdown]
### Creation of excel spreadsheet of scaled features

# %% [code]
sdf_unsc2.df_.to_excel("../../datasets/full_sdf.xlsx", index=None)

# %% [code]
```
