## Supplementary material for "Predicting Relative Protein Abundance via Sequence-Based Information": S3 Data: SDF-Official1Pie.ipynb

```
# %% [code]
import matplotlib.pyplot as plt
import numpy as np
import pandas as pd
import itertools as it
from matplotlib import cm
%matplotlib inline

import sys
sys.path.insert(0, "../../scripts/")
sys.path.insert(0, "../../../turbopanda/")
import myfig as mf
import version
import turbopanda as turb

# %% [markdown]
## Current Version

# %% [code]
version.current()
print("turbopanda: %s" % turb.__version__)

# %% [markdown]
# SDF Pie

### This Notebook is responsible for generating the PIE chart for Figure 1

## Figure 1: SDF Figure of Pie Charts with SDF breakdown

1. Load in SDF dataset
2. Generate 2 groups
    1. group 1: every single data grouping such as codon bias, length etc.
    2. group 2: groups of the grouping, i.e the data source
3. Generate palette for data source, data groups
4. Map clean names
5. Plot pie chart with extras

#### Import SDF

# %% [code]
sdf = turb.read("../../datasets/SDF_scaled2.json")
sdf

# %% [code]
sdf.head(2)

# %% [code]
sdf.meta_['data_groups'].cat.add_categories("hgnc", inplace=True)
sdf.meta_['data_groups'].fillna("hgnc", inplace=True)

# %% [markdown]
#### Form color groups

# %% [code]
# generate groups based on the METADATA in SDF.
# calculate data group counts
grp1 = (sdf.meta_.groupby(["data_source","data_groups"], as_index=False)["true_type"].count()).dropna()
# calculate counts for the cmap
grp2 = grp1.groupby("data_source")['true_type'].agg(['sum','count'])
# set HGNC: blue, PTM: purple, Refseq: green, Uniprot: red
grp2["cm"] = pd.Series({'hgnc': 'Blues', 'refseq': 'Greens', 'uniprot': 'Reds', 'ptm_source': 'Purples'})
grp2["count_scaled"] = grp2["count"] / grp2['count'].sum() * 100.
# determine palette
grp2['palette'] = pd.Series(map(lambda text: cm.get_cmap(text)(0.5), grp2['cm']), index=grp2.index)
# assign palette for subgroups
grp1['palette'] = pd.Series(
    it.chain.from_iterable(
        map(lambda c, text: cm.get_cmap(text)(np.linspace(0.8, 0.2, c)), grp2['count'], grp2['cm'])),
    index=grp1.index
)

# %% [code]
grp1

# %% [code]
grp2

# %% [code]
import seaborn as sns

# %% [code]
sns.palplot(grp2['palette'])

# %% [code]
sns.palplot(grp1['palette'])

# %% [markdown]
#### Plot the PIE chart

# %% [code]
def plot_pie_chart():
    # mappables
    clean_sources = {
        "refseq": "Refseq/\nEnsembl",
        "uniprot": "Uniprot/\nSwissprot",
        "hgnc": "HGNC",
        "ptm_source": "PhosphoSitePlus" 
    }

    clean_groups = {
        "hgnc": "",
        "id_col": "ID Columns",
        "ptm": "Post-Translational\nModification",
        "codon_bias": "Codon Bias",
        "dinuc_freq": "Dinucleotide\nFrequency",
        "l_prop": "Proportional\nLength",
        "length": "Sequence Length",
        "mononuc_freq": "Mononucleotide\nFrequency",
        "rna_misc": "RNA Miscallaenous\nFeature",
        "spec_freq": "Special Sequence\nFrequency",
        "aa_freq": "Amino Acid Frequency",
        "biophysical": "Amino Acid\nBiophysical Property",
        "translation_freq": "Translation Frequency",
        "free_energy": "Minimum 5'UTR\nFree Energy",
        "aminoacid_meta": "Amino Acid\nText-Mined Feature"
    }
    
    # function to autopct
    def label_func(pct, allvals):
        absolute = int(np.round(pct/100.*np.nansum(allvals)))
        return "{:.1f}% ({:d})".format(pct, absolute)
    
    # create figure
    fig, ax = plt.subplots(figsize=(5,5), dpi=360)
    
    # create pie plots
    p1, tex1, autotex1 = ax.pie(grp1["true_type"], 
                      labels=grp1["data_groups"].map(clean_groups),
                      colors=grp1['palette'], radius=3,
                      startangle=60,
                      shadow=False,
                      autopct=lambda pct: label_func(pct, grp1["true_type"]),
                      pctdistance=.8,
                      labeldistance=1.05,
                      rotatelabels=True,
                      textprops=dict(fontsize=15),
                      wedgeprops=dict(linewidth=1.5, edgecolor="w", width=1.1))

    p2, tex2 = ax.pie(grp2["sum"], 
                      labels=grp2.index.map(clean_sources),
                      colors=grp2['palette'], startangle=60,
                      labeldistance=0.3,
                      radius=2, 
                      shadow=False,
                      rotatelabels=True,
                      textprops=dict(fontsize=17),
                      wedgeprops=dict(linewidth=1.5, edgecolor="w", width=1.5)
    )
    
    # adjust rotation for autopct
    for t, auto in zip(tex1, autotex1):
        auto.set_rotation(t.get_rotation())
    
    _ = turb.plot.save(fig, 
                plot_type="pie", 
                name="SDF_Features",
                save_types=('pdf', 'svg', 'png', 'eps'),
                fp="../../Figures/SDF/",
                savemode="update",
                dpi=1200)
    
    plt.show()
    

# %% [markdown]
## Description

### The Breakdown of Sequence-Derived Features and their sources.

   Pie chart detailing the number and proportion of features by database source (inner ring) and broad data-grouping (outer ring). mRNA/Ensembl/Refseq-like features (54\%) are denoted in shades of green, amino-acid/Uniprot/Swissprot-like features (33\%) are denoted in shades of red. Post-translational modification (4\%) features are extracted separately (in purple). HGNC labels (7\%) are not included in the machine-learning algorithms but are shown here for completeness (in blue).

# %% [code]
plot_pie_chart()

# %% [code]
```
