## Supplementary material for "Predicting Relative Protein Abundance via Sequence-Based Information": S3 Data: SDF-Official2Corr.ipynb

```
# %% [markdown]
# Figure 2

What is contained within this notebook:

To navigate the notebook, click the links below or scroll:

1. **[Initialize and import](#init)**
2. **[Sorting based on the central dogma](#dogma)**
3. **[The official plot](#offic)**
4. **[Plot Description](#desc)**
5. **[Interactive Bokeh Plots](#bokeh)**
6. **[Supplementary Material 1: Determining Prominence of Correlations](#corrmap)**
7. **[Supplementary Material 2: Comparison to Vogels' SDF dataset](#vogel)**
7. **[Supplementary Material 3: Mutual Information Matrix between SDF features for nonlinearity](#MI)**

## Objectives

We describe a correlation matrix between random variables in vectors $\mathbf{X}_i$ as:

$$
\begin{equation}
\text{corr}[\mathbf{X}]=\mathbf{\Omega}_{\mathbf{X}}=\left(\text{diag} \left( \mathbf{\Sigma}_{\mathbf{X}} \right) \right)^{-\frac{1}{2}} \mathbf{\Sigma}_{\mathbf{X}} \left( \text{diag}\left( \mathbf{\Sigma}_{\mathbf{X}} \right) \right)^{-\frac{1}{2}}
\label{eq:corr1} \tag{1}
\end{equation}
$$

where $\mathbf{\Sigma}_X$ is the covariance matrix. Equation $\eqref{eq:corr1}$ defines the official correlation matrix (mostly).

# %% [code]
import matplotlib.pyplot as plt
import numpy as np
import seaborn as sns
import pandas as pd
from matplotlib import cm
import matplotlib as mpl
from scipy.cluster.hierarchy import linkage, dendrogram
import itertools as it
%matplotlib inline

import sys
sys.path.insert(0, "../../scripts/")
sys.path.insert(0, "../../../turbopanda/")
import misc_functions as misc
import version
import myfig as mf
import turbopanda as turb

from matplotlib.gridspec import GridSpec, GridSpecFromSubplotSpec
from matplotlib.lines import Line2D

# %% [markdown]
## Version

# %% [code]
version.current()
print("turbopanda: %s" % turb.__version__)

# %% [code]
from turbopanda import cache

# %% [markdown]
## <a id='init'> SDF Official Spearman-rank correlation matrix of SDF features </a>

#### Import SDF

# %% [code]
sdf = turb.read("../../datasets/SDF_scaled2.json")
sdf

# %% [markdown]
### Determining 'distance' according to the Central Dogma of Molecular Biology

    mRNA features -> misc features -> 3'UTR -> 5'UTR misc features -> Free energy -> CDS f -> codon bias -> amino acids -> meta amino acid -> biophysical protein -> PROTEIN -> ptm features
    
#### Subgroups within distance groups

Within mRNA features, for example, we will group nucleotide groups as follows:

1. Features that exclusively use $G$ and/or $C$
2. Features that exclusively use $A$ and/or $T$
3. Features that mix $G/C$ and $A/T$

We apply this across mRNA, CDS, 5'UTR and 3'UTR.

This is because we see in the correlation matrix that the correlation between fellow $G/C$ counts are **positive**, but correlations between $G/C$ and $A/T$ features are **negative**.

# %% [markdown]
#### Calculate correlation

We cache this function because it's moderately expensive to keep re-calculating.

# %% [code]
def row_to_matrix(df, y="r"):
    # pivot
    square = df.pivot("column1","column2",y)
    # fill missing with 0, no correlation
    square.fillna(0., inplace=True)
    # tranpose+
    square += square.T
    # remove 1 from the diagonal elements
    square.values[np.diag_indices_from(square.values)] -= 1.
    return square

# %% [markdown]
### Convert `bool` features to `uint8`

# %% [code]
pre_corr = sdf.copy().drop("object", ".*id$", 
                           ".*ID$", "^ID.*", "^id.*",
                           lambda z: z.nunique() == 1)

# %% [code]
pre_corr.downcast()

# %% [code]
@cache(filename="../../_intermediate/corr_matrix_sdf.csv", return_as="pandas")
def calculate_correlation(sdf):
    return turb.correlate(data=sdf)

# %% [code]
corr_long = calculate_correlation(pre_corr)

# %% [markdown]
#### Convert and copy across attributes

# %% [code]
corr = turb.corr.row_to_matrix(corr_long)

# %% [code]
pre_corr

# %% [markdown]
#### Hierarchical Clustering of Correlations


# %% [code]
Z=linkage(corr)
Y=dendrogram(Z,get_leaves=True, no_plot=True)
clust=corr.columns[Y["leaves"]]

# %% [code]
clust

# %% [markdown]
### Sorting ORDER:

1. Central dogma order
2. Data groups (dinucleotide, mononucleotide, free energy)
3. Base pairing (within dinucleotide, mononucleotide groups)
4. Hierarchical clustering of ($r_s$, $r_{bs}$) correlates

# %% [code]
# incorporate hierarchical clustering data as final layer
pre_corr.meta_["hierarchical"] = pd.Series(np.arange(len(clust)), index=clust)
# create a sorted order
sorted_order = pre_corr.meta_.sort_values(by=["dogma_order", 'data_groups', "base_pairing", "hierarchical"]).index

# %% [code]
# apply sorted order to our matrix
corr_sorted = corr.reindex(sorted_order, axis=0).reindex(sorted_order, axis=1)
# apply sorting back to the 'pre-matrix' meta
meta_info = pre_corr.meta_.reindex(corr_sorted.index)

# %% [markdown]
### Sorting out colours - a nightmare

- Group 1: `g1`: data groups (such as length, length proportio, biophysical etc)
- Group 2: `g2`: data sources (hgnc, refseq, uniprot)

# %% [code]
g1 = meta_info.groupby(["data_source","data_groups"], as_index=False)["true_type"].count().dropna()
g2 = g1.groupby("data_source", as_index=False)["true_type"].agg(["sum","count"])
g2["cm"] = pd.Series({"ptm_source":"Purples", "refseq":"Greens", "uniprot":"Reds", "hgnc":"Blues"})

# %% [code]
g1

# %% [code]
g2

# %% [markdown]
Here we assign a palette color to each *data group*, such that Greens are Refseq/RNA sources, Reds are Uniprot/Protein sources, and Purples are Post-translational modification sources

# %% [code]
# assign palettes
g1["palette"]=pd.Series(list(it.chain.from_iterable(
    [cm.get_cmap(r["cm"])(np.linspace(0.8, .1, r["count"])) for i, r in g2.iterrows()]
)), index=g1.index)
g2["palette"] = pd.Series([cm.get_cmap(r["cm"])(0.5) for i, r in g2.iterrows()], index=g2.index)
# create 'color litmus ID'
g1['color_litmus'] = turb.str.reformat("{data_source}__{data_groups}", g1)
meta_info["color_litmus"] = turb.str.reformat("{data_source}__{data_groups}", meta_info)

# map the colors on
mapping_group = meta_info["color_litmus"].map(g1.set_index(["color_litmus"])["palette"].to_dict())
mapping_source = meta_info["data_source"].astype(object).map(g2["palette"].to_dict())

# %% [code]
sns.palplot(g2['palette'])

# %% [code]
sns.palplot(g1['palette'])

# %% [markdown]
Here we mix the colors such that Red and Purples overlap for the central dogma mapping

# %% [code]
def mix_colors(c1="Reds_r", c2="Purples", n=20):
    """ In this case, we disproportionately mix red and purple to get a different color ending. """
    m1 = cm.get_cmap(c1)(np.linspace(0.1, 3, n))
    m2 = cm.get_cmap(c2)(np.linspace(0.1, 1.5, n))
    return np.clip(m1*m2, 0, 1)

# %% [code]
sns.palplot(mix_colors(n=int(g2.loc["ptm_source","sum"])))

# %% [markdown]
Here we create a longer color scale for the central dogma mapping -> all the way from green to purple.

# %% [code]
dogma_stretch = np.vstack((
    cm.RdYlGn_r(np.linspace(0,.33,int(g2.loc["refseq","sum"]))),
    cm.RdYlGn_r(np.linspace(.66,1.,int(g2.loc["uniprot","sum"]))),
    mix_colors(n=int(g2.loc["ptm_source","sum"]))
))

mapping_dogma = pd.Series(dict(zip(mapping_source.index, dogma_stretch)))

# %% [code]
sns.palplot(dogma_stretch)

# %% [markdown]
### Figuring out which x/yticklabels to display...

1. First off, we clean the labels using a mapping dictionary
2. We create a category of 'htm labels' combining data region and groups
3. We rename
4. We calculate the correct ticks to show up on the plot, with the appropriate label
5. We drop excess ticks based on text size

# %% [code]
def dict_to_tuple(d):
    return list((a, b) for a, b in d.items())

# %% [code]
clean_groups_abbriev = {
    'hgnc': '',
    "id_col": "ID Columns",
    "ptm": "PTM",
    "codon_bias": "Codon Bias",
    "dinuc_freq": "DM Frequency",
    "l_prop": "Proportional\nLength",
    "length": "Sequence Length",
    "mononuc_freq": "MN Frequency",
    "rna_misc": "Misc. Feature",
    "spec_freq": "Special Frequency",
    "aa_freq": "AA Frequency",
    "biophysical": "AA Biophysical Property",
    "translation_freq": "Trans. Property",
    "free_energy": "MFE",
    "aminoacid_meta": "AA Text-\nMined Feature"
}

# %% [markdown]
Here we add updated labels to our meta information which include region and group.

# %% [code]
region_clean = {'mrna': '(mRNA)', 'protein':'(Protein)',"utr3":"(3'UTR)",
                'utr5':"(5'UTR)", 'cds':'(CDS)'}


# %% [code]
meta_info["htm_labels"] = list(map(lambda x, y: str(x) + "__" + str(y), 
    meta_info["data_region"].map(region_clean), 
    meta_info['data_groups'].map(clean_groups_abbriev)))

# %% [code]
htm_renaming3 = [
    ("refseq__", "(Refseq) "),
    ("uniprot__", "(UniProt) "),
    ("ptm_source__", "(Phospho) ")
]

# %% [code]
import re

# %% [code]
# fetch unique labels
unique_group_labels = np.asarray(list(map(lambda x: x.replace("__", " "), meta_info['htm_labels'].unique())))
# filter out labels that include sequence length, special frequency.
filtered_out_names = "Sequence Length|Special Frequency|Codon Bias|nan|Proportional"

# %% [code]
# extract start and end indices
reduced_uniq_group_labels = meta_info['htm_labels'].unique()
start_ind = meta_info.reset_index()['htm_labels'].drop_duplicates(keep='first').index
end_ind = meta_info.reset_index()['htm_labels'].drop_duplicates(keep='last').index

# %% [code]
median_inds = np.asarray(list(map(lambda a, b: int(a + (b - a) / 2), start_ind, end_ind)))
median_inds

# %% [markdown]
#### Dropping indices that we don't want to display

# %% [code]
unique_group_filter = ~np.array([re.search(filtered_out_names, x) for x in unique_group_labels], dtype=bool)

# %% [markdown]
### Generating (and cleaning) Legend

# %% [code]
g1_ordered = g1.set_index("color_litmus").reindex(meta_info["color_litmus"].drop_duplicates().values)
new_legend = [Line2D([0], [0], color=c, lw=4) for c in g1_ordered["palette"].values]
legend_labels = turb.str.string_replace(
    turb.str.string_replace(g1_ordered.index,htm_renaming3), dict_to_tuple(clean_groups_abbriev))

# %% [markdown]
## <a id='offic'> Static Heatmap with Matplotlib </a>

Abbrievations being used...:

1. Mononucleotide (MN)
2. Dinucleotide (DN)
3. Miscallaenous (misc.)
4. Amino acid (AA)
5. Post-translational modification (PTM)
6. Minimum Free Energy (MFE)

# %% [code]
def plot_heatmap():
    fig = plt.figure(figsize=(16,16), dpi=360)

    gs0 = GridSpec(32, 32)
    ax = fig.add_subplot(gs0[2:-1,1:-1])
    ax_dogma = fig.add_subplot(gs0[1, 1:-1])
    ax_dogma_label = fig.add_subplot(gs0[0, 1:-1])
    ax_group = fig.add_subplot(gs0[2:-1, 0])
    ax_cbar = fig.add_subplot(gs0[2:-1,-1])

    cbar_range =  np.linspace(-1, 1, 21)
    norm = mpl.colors.BoundaryNorm(cbar_range, 20)
    label = r"correlation ($r_s$ | $r_{bs}$)"

    cbar_args = dict(label=label, cmap="seismic",
                     ticks=cbar_range, spacing="proportional",
                     boundaries=cbar_range, format="%0.1f")

    sns.heatmap(corr_sorted, 
                cbar_kws=cbar_args, cmap="seismic", cbar_ax=ax_cbar, vmin=-1., vmax=1., ax=ax)

    # plot colorbars for groups.

    ax_group.barh(np.arange(corr_sorted.shape[0]), width=1., height=1., align="edge",
                 color=mapping_group[::-1])

    # set x and y ticks for the heatmap
    ax.tick_params("x", length=10, labelrotation=45, labelsize=13)

    ax.set_xticks(median_inds[unique_group_filter])
    ax.set_xticklabels(unique_group_labels[unique_group_filter])
    for tick in ax.get_xticklabels():
        tick.set_horizontalalignment("right")


    # set x and y ticks for the heatmap
    ax_group.tick_params("y", length=10, labelsize=12)

    ax_group.set_yticks(meta_info.shape[0] - median_inds[unique_group_filter])
    ax_group.set_yticklabels(unique_group_labels[unique_group_filter])


    ax_dogma.bar(np.arange(corr_sorted.shape[0]), width=1., height=1., align="edge",
                 color=mapping_dogma)

    bbox_props=dict(boxstyle="rarrow,pad=0.3", fc="#ffeeee", ec="k", lw=1)
    ax_dogma_label.text(0.5, .8, "Direction of Central Dogma [DNA - mRNA - Protein - PTM]", 
                        ha="center", va="center", rotation=0, size=20, bbox=bbox_props)

    ax_group.legend(new_legend, legend_labels.tolist(),
                    bbox_to_anchor=(-6.5, 0.8),shadow=True,fancybox=True,
                    ncol=1, title="Data Groups", fontsize=15)

    # set bar plot margins
    ax_group.margins(x=0, y=0)
    ax_dogma.margins(x=0, y=0)

    # detick and despine
    mf.detick(ax, axis="y")
    mf.detick(ax_dogma)
    mf.detick(ax_dogma_label)
    mf.detick(ax_group, axis="x")
    mf.despine(ax_group)
    mf.despine(ax_dogma)
    mf.despine(ax_dogma_label)

    # adjust sizes
    ax_cbar.tick_params(labelsize=15)
    ax_cbar.set_ylabel(label, fontsize=15)

    # adjust global labels
    ax.set_ylabel("")
    ax.set_xlabel("Feature (x)")

    _ = turb.plot.save(fig, 
                plot_type="heatmap",
                save_types=('png','pdf','svg', 'eps'),
                name="SDF_IntraCorr",
                savemode="first",
                fp="../../Figures/SDF/", 
                dpi=1200)

    plt.show()
    
    return fig, ax
    

# %% [markdown]
## <a id='desc'> Description </a>

### Large-scale interdependencies between SDFs reveal source dependency.

    Spearman-rank ($r_s$)/Biserial correlation ($r_{bs}$)-mixed correlation matrix between sequence-derived features (SDFs). See Methods for details on correlation method. Both axes indicate the direction of molecular biology (from DNA to post-protein). mRNA/RNA features are denoted in green shades, amino-acid features are denoted in red shades, PTM features are denoted in purple. Off-diagonal elements indicate a correlation ($r_s$|$r_{bs}$) between two features (red: positive, blue: negative).

# %% [code]
f, a = plot_heatmap()

# %% [markdown]
## <a id='bokeh'> Interactive Heatmap with Bokeh </a>

# %% [code]
from bokeh.plotting import figure, show, ColumnDataSource
from bokeh.models import LinearColorMapper, ColorBar, BasicTicker, HoverTool, Div, Paragraph
from bokeh.io import output_notebook, output_file
from bokeh import transform
from bokeh.layouts import row, column, grid
output_notebook()

# %% [markdown]
#### Make column1, column2 categoricals in order to sort them.

# %% [code]
# pretty names
pretty_names = [("pH_",""), ("_prop"," rel."),
        ("_cds"," CDS"),("_utr5"," 5'UTR"), ("_utr3"," 3'UTR"),
        ("_win"," (win"), ("_total",""), ("_bp",""), ("_bcaa",""),
        ("mfe","free energy"), ("_mrna"," mRNA"), ("_aa", " (amino acid)"),
        ("_ptmc", " PTM"), ("_fc",""),("_pc"," (corrected)"), ("_"," ")]

# %% [code]
# define categorical as a way of ordering them.
cats = pd.CategoricalDtype(categories=turb.str.string_replace(corr_sorted.columns, pretty_names), ordered=True)

# %% [code]
def mirror_long(df, x='column1', y='column2'):
    df = df.df_ if isinstance(df, turb.MetaPanda) else df
    df_mirror = df.copy(deep=True)
    df_mirror[x] = df[y]
    df_mirror[y] = df[x]
    return pd.concat([df, df_mirror], axis=0, sort=False)

# %% [code]
corr_long['clean_x'] = turb.str.string_replace(corr_long['x'], pretty_names)
corr_long['clean_y'] = turb.str.string_replace(corr_long['y'], pretty_names)

# %% [markdown]
#### Clean the column, index names

# %% [code]
# save memory
corr_long_2 = mirror_long(corr_long, x='clean_x', y='clean_y').round(3)
corr_long_2.columns = turb.str.string_replace(corr_long_2.columns, [("p-val", "p_val")])
bok_dogma = ColumnDataSource(corr_long_2)

# %% [code]
def ndarray_to_hex(arr):
    return ["#%02x%02x%02x" % (int(r*255), int(g*255), int(b*255)) for r, g, b, _ in arr]

# %% [code]
def plot_bokeh_corr(dataset, 
                    axis_order,
                    filename="../../Figures/SDF/_interactive/Fig2.html",
                    x='x',
                    y='y',
                    method='method',
                    title=None,
                    desc=None):
    
    if title is None:
        title = """Fig 2. Large-scale interdependencies between SDFs reveal source dependency."""
    if desc is None:
        desc = """Spearman-rank ($r_s$)/Biserial correlation ($r_{bs}$)-mixed correlation matrix
        between sequence-derived features (SDFs). See Methods for details on correlation method. Both axes 
        indicate the direction of molecular biology (from DNA to post-protein). mRNA/RNA features are denoted 
        in green shades, amino-acid features are denoted in red shades, PTM features are denoted in purple.
        Off-diagonal elements indicate a correlation ($r_s$|$r_{bs}$) between two features 
        (red: positive, blue: negative)."""
    
    # generate output
    output_file(filename, title=title)
    
    # create the colormap
    palette = ndarray_to_hex(cm.get_cmap("seismic")(np.linspace(0, 1, 20)))
    # map it into the color palette
    mapper = LinearColorMapper(palette=palette, low=-1., high=1.)

    # specify the ordering of the axes as our category type
    axis0 = axis_order.categories.tolist()
    axis1 = axis_order.categories.tolist()
    # tooltips
    tt = [("Feature 1", "@%s" % x),
          ("Feature 2", "@%s" % y),
          ("correlation", "@r{1.111}"),
          ("p-value", "@p_val{1.11}"),
          ("n", "@n"),
          ("method", "@%s" % method)]
    
    # figure main
    p = figure(
        title=title,
        plot_width=1000, plot_height=900,
        x_axis_label="Feature (x)",
        y_axis_label="Feature (y)",
        x_range=axis0,
        y_range=axis1,
        tooltips=tt,
        toolbar_location="above"
    )
    
    # draw rectangles
    p.rect(x=x, y=y, 
           width=1, height=1, 
           source=dataset,
           line_color=None, 
           fill_color=transform.transform("r", mapper))
    
    p.xaxis.major_label_orientation = "vertical"
    
    dd_title = Div(text="<b>{}</b>".format(title), width=1000, height=20)
    dd_desc = Paragraph(text=desc, width=1000, height=100)
    
    # add colorbar
    cbar = ColorBar(color_mapper=mapper, location=(0, 0),
                    ticker=BasicTicker(desired_num_ticks=len(palette)))
    p.add_layout(cbar, "right")
    show(column(dd_title, dd_desc, p))

# %% [code]
plot_bokeh_corr(bok_dogma, axis_order=cats, x='clean_x', y='clean_y', method="method")

# %% [markdown]
# <a id='corrmap'> Supplementary Information (2) </a>

Which features are the most correlated, on average, with other features?

We calculate the $\ell_1$-norm for each row of correlations $r_i$:

$$
||r_i||_1 = \sum_j \left|r_{ij} \right|
$$

Features with extremely high $||r_i||_1$ may indicate heavy redundancy, whereas features with low $||r_i||_1$ may not be valuable as features, have a strictly nonlinear relationship or only come into play when other intercorrelations are removed.

# %% [code]
def bokeh_supp_inf1(matrix, clean_columns, data_groups):
    
    title="""Supplementary 2. Which features are the most or least correlated, on average, with each other?"""
    
    desc="""
    Which features are the most correlated, on average, with other features? We calculate the L1-norm for each row of correlations r.

    Features with extremely high L1 may indicate heavy redundancy, whereas features with low L1
    may not be valuable as features, have a strictly nonlinear relationship or only come into play when other
    intercorrelations are removed. One factor that jumps out is the reduced L1 for dinucleotide features that have one CG and one AT, i.e AG/CT.
    """
    
    output_file("../../Figures/SDF/_interactive/supp1_fig2.html", title=title)
    
    supp_ds = ColumnDataSource({"cols": matrix.columns,
                                "cols_clean": clean_columns,
                                "l1": matrix.aggregate(np.linalg.norm, ord=1, axis=0),
                                "colors": ndarray_to_hex(mapping_group),
                                "groups": data_groups})
    hvt = HoverTool(
        tooltips = [
            ("SDF", "@cols_clean"),
            ("l1-norm", "@l1"),
            ("Data Group", "@groups")
        ],
        mode="vline"
    )
    # figure main
    supp1_p = figure(
        title="",
        plot_width=1600, plot_height=500,
        x_axis_label="Sequence-derived Features",
        y_axis_label=r"||r||",
        x_range=supp_ds.data['cols_clean'].tolist(),
        tools=[hvt],
        toolbar_location="above"
    )
    supp1_p.xaxis.major_label_orientation=np.pi/2.5
    
    dd_title = Div(text="<b>{}</b>".format(title), width=1600, height=20)
    dd_desc = Paragraph(text=desc, width=1600, height=70)
    
    r = supp1_p.vbar(x="cols_clean", top="l1", source=supp_ds, color="colors", width=.7, line_color="#444444")
    show(column(dd_title, dd_desc, supp1_p))

# %% [code]
bokeh_supp_inf1(corr_sorted, 
                turb.str.string_replace(corr_sorted.columns, pretty_names),
                meta_info['htm_labels'].str.replace("__", " "))

# %% [markdown]
# <a id='vogel'> Supplementary Material (3) </a>

Intercorrelations between Vogels' 2010 sequence-derived feature dataset.. see if it's different?


# %% [code]
vogel = turb.read("../../data/vogel_dataset_rna-prot.xls", name='Vogel', header=5,sheet_name="MARS_data")
vogel

# %% [markdown]
### Calculate correlations here...

# %% [code]
C = turb.correlate(vogel[float])

# %% [code]
# save memory
corr_long_3 = mirror_long(C, x='x', y='y').round(3)
# use the clean_map to rename stuff
bok_dogma2 = ColumnDataSource(corr_long_3.rename(columns={'p-val':'pval'}))

# %% [code]
plot_bokeh_corr(bok_dogma2, 
                pd.CategoricalDtype(vogel.view(float)),
                x='x', y='y',
                method='method',
                filename="../../Figures/SDF/_interactive/supp2_fig2.html",
                title="Supplementary 1. Interdependencies between SDFs from Vogel et al. 2010 Dataset.",
                desc="""Spearman-rank correlation between SDFs as derived from Vogel et al. See Methods for details on correlation method. 
                The data is not sorted in any particular way and is as presented in the order of the Supplementary file. """)

# %% [markdown]
## Supplementary Material (4)

For each data group, what is the intra-inter magnitude of correlations. We want to demonstrate here that the relationships between each feature in the group are in general stronger than with other features and hence a future dimensionality reduction along these lines seems sensible.

# %% [code]
name_to_group_map = pre_corr.meta_['data_groups'].to_dict()

# %% [code]
# using corr_long, associate data group data to that
corr_long['data_group_x'] = corr_long['x'].map(name_to_group_map)
corr_long['data_group_y'] = corr_long['y'].map(name_to_group_map)

# %% [code]
corr_long['data_groups_same'] = ((corr_long['data_group_x'] == corr_long['data_group_y'])
    .astype("category")
    .cat.rename_categories({True: 'Same group', False: 'Different groups'}))

# %% [code]
remove_duplicates_corrlong = corr_long['x'] != corr_long['y']
unique_groups = turb.utils.union(corr_long['data_group_x'], corr_long['data_group_y'])

# %% [code]
def supp4_select_subset_and_groupby(u):
    return (corr_long.loc[remove_duplicates_corrlong & (corr_long['data_group_x']==u)|(corr_long['data_group_y']==u)]
                .groupby("data_groups_same")['adj_r2'].mean())

# %% [code]
bar_plot_data = pd.concat(list(map(supp4_select_subset_and_groupby, unique_groups)), axis=1).T
bar_plot_data.index = unique_groups

# %% [code]
(bar_plot_data.index, )

# %% [code]
def widebar(data):
    fig, ax = plt.subplots(figsize=(8, 6))
    n, p = data.shape
    pretty_names = turb.str.string_replace(data.index, 
                                           turb.utils.dict_to_tuple(clean_groups_abbriev))
    ticks = np.arange(n)
    prop_tick_add = 1. / p - 0.1
    pal = turb.plot.color_qualitative(p)
    order = np.argsort(bar_plot_data.mean(axis=1))
    
    for d, c in enumerate(pal):
        ax.barh(ticks + (d * prop_tick_add), data.iloc[:, d], 
               height=prop_tick_add, color=c, label=data.columns[d])
    ax.set_yticks(ticks + .25)
    ax.set_yticklabels(pretty_names)
    ax.legend(loc='best')
    ax.set_xlabel(r"Mean Adjusted $R^2$")
    ax.set_ylabel("Data Groups")
    ax.set_title("Relationship between Intra/Inter Data Group Correlates.")
    
    # save as figure
    _ = turb.plot.save(fig, 
                plot_type="bar",
                save_types=('png','pdf','svg'),
                name="SDF_supp4",
                savemode="first",
                filepath="../../Figures/SDF/", 
                transparent=True,
                dpi=360)

# %% [markdown]
## Plot of the Mean Adjusted $R^2$ for each data group

All features show a stronger intra-group correlation than with features outside of it's data group.

# %% [code]
widebar(bar_plot_data)

# %% [code]
```
