## Supplementary material for "Predicting Relative Protein Abundance via Sequence-Based Information": S3 Data: SDF-Official3Corr.ipynb

```
# %% [markdown]
# Figure 3

What is contained within this notebook:

To navigate the notebook, click the links below or scroll:

1. **[Initialize and import](#init)**
2. **[Calculate the partial correlations, fixing mRNA level in-place](#partial_calc)**
3. **[Defining the Official Plot](#offic_plot)**
3. **[Plot Description](#desc)**
4. **[Interactive Bokeh Plots](#bokeh)**
5. **[Supplementary Material 1 - Conditional on mRNA/Length](#supp1)**
6. **[Supplementary Material 2 - Canonical Correlation Analysis (CCA) between Vogel and us](#supp2)**


## Objectives

We describe a correlation matrix between random variables in vectors $\mathbf{X}_i$ as:

$$
\begin{align}
\text{corr}[\mathbf{X}]=\mathbf{\Omega}_{\mathbf{X}}=\left(\text{diag} \left( \mathbf{\Sigma}_{\mathbf{X}} \right) \right)^{-\frac{1}{2}} \mathbf{\Sigma}_{\mathbf{X}} \left( \text{diag}\left( \mathbf{\Sigma}_{\mathbf{X}} \right) \right)^{-\frac{1}{2}}
\label{eq:corr1} \tag{1}
\end{align}
$$

where $\mathbf{\Sigma}_X$ is the covariance matrix. Equation $\eqref{eq:corr1}$ defines Figure 2. In Figure 3 we draw the scatter of correlations:

$$
\begin{align}
\rho=\begin{cases}
r_s & X_{\text{SDF},k} \in \mathbb{R} \\
r_{bs} & X_{\text{SDF},k} \in \mathbb{Z} 
\end{cases}
\end{align}
$$

where $r_s$ is the Spearman-rank correlation coefficient, $r_{bs}$ is the biserial correlation coefficient, $\mathbb{R},\mathbb{Z}$ describe whether the variable is continuous or boolean-like, respectively. Thus we draw;

$$
\text{LEFT}=\begin{equation}
\left( \rho_k \left(X_{\text{SDF},k}, Y_{\text{mRNA}} \right), \rho_k \left(X_{\text{SDF},k}, Y_{\text{protein}} \right) \right) \quad \forall k \label{eq:corr2} \tag{2} \end{equation} \\
$$

$$
\text{RIGHT}=\begin{equation}
\left( \rho_k \left(X_{\text{SDF},k}, Y_{\text{mRNA}} \right), \rho_k \left(X_{\text{SDF},k}, Y_{\text{protein}}  | Z_{\text{mRNA}}\right) \right) \quad \forall k \label{eq:corr3} \tag{3}
\end{equation}
$$

Equations $\eqref{eq:corr2},\eqref{eq:corr3}$ describe the Official figure which looks at eliminating the bias of mRNA in protein prediction. The Supplementary material looks at instead:

$$
\text{LEFT}=\begin{equation}
\left( \rho_k \left(X_{\text{SDF},k}, Y_{\text{mRNA}} \right), \rho_k \left(X_{\text{SDF},k}, Y_{\text{protein}} \right) \right) \quad \forall k \label{eq:corr4} \tag{4} \end{equation} \\
$$

$$
\text{RIGHT}=\begin{equation}
\left( \rho_k \left(X_{\text{SDF},k}, Y_{\text{mRNA}} \right), \rho_k \left(X_{\text{SDF},k}, Y_{\text{protein}}  | Z_{\text{Length}}\right) \right) \quad \forall k \label{eq:corr5} \tag{5}
\end{equation}
$$

Alternatively, to look at the changes directly, we could take the difference of the partial w.r.t the associated data expression value:

$$
\begin{equation}
\Delta \rho_k=\left(\rho_k \left(X_{\text{SDF},k}, Y_{\text{mRNA}}  | Z_m\right)- \rho_k \left(X_{\text{SDF},k}, Y_{\text{mRNA}} \right), \rho_k \left(X_{\text{SDF},k}, Y_{\text{protein}} | Z_m\right)- \rho_k \left(X_{\text{SDF},k}, Y_{\text{protein}} \right) \right) \label{eq:corr6} \tag{6} \end{equation} \quad \forall k, \, k \neq m
$$

where $m$ could represent any SDF-controlling variable(s), such as sequence length, codon bias etc. This then represents the relative change in correlation that inducing the partial-correlation has on the feature. Large changes in this value indicate that $Z$ is highly controlling/related to this feature. In particular when $\Delta \rho_k = 0$, the variable has no effect.

# %% [code]
import os
import matplotlib.pyplot as plt
import numpy as np
import seaborn as sns
import pandas as pd
from matplotlib import cm
import matplotlib as mpl
from scipy.cluster.hierarchy import linkage, dendrogram
from scipy import stats
import itertools as it
import string
%matplotlib inline

from matplotlib import patches as mpatches
from matplotlib.gridspec import GridSpec, GridSpecFromSubplotSpec
from matplotlib.lines import Line2D

import sys
sys.path.insert(0, "../../scripts/")
sys.path.insert(0, "../../../turbopanda/")
# scripts
import version
import myfig as mf
# turbopanda
import turbopanda as turb
from turbopanda import cache

# %% [markdown]
## Version

# %% [code]
version.current()
print("turbopanda: %s" % turb.__version__)
print(version.last_run())

# %% [markdown]
## <a id='init'> SDF Official Scatter matrices of SDF features against mRNA/Protein </a>

#### Import SDF

# %% [code]
sdf = turb.read("../../datasets/SDF_scaled2.json", 
                name="SDF", with_clean=True)
sdf

# %% [markdown]
### Loading RNA/Protein Expression Datasets

1. Aviner (2015), HeLa cell line, time-series dataset, includes translation
2. Vogel (2010), Daoy cell line
3. Lundberg (2010), U2OS, U251MG and A431 cell lines

Each of these datasets undergoes three main stages: *Load, Merge and Correlate*.

#### Stage 1. Load

# %% [code]
aviner_rna, aviner_prot, aviner_tr = turb.read("../../datasets/clean/*_aviner_adv.csv", 
                    name=("mRNA_aviner", "prot_aviner", "tr_aviner"), index_col=0)
lund_all, lund_prot, lund_rna = turb.read("../../datasets/lundberg_*.csv")

# %% [markdown]
#### Stage 2. Merge with SDF

Pairwise-merges with our sequence-derived features to maximise number of samples.

# %% [code]
av_prot_sdf = turb.merge([sdf, aviner_prot])
av_tr_sdf = turb.merge([sdf, aviner_tr])
av_rna_sdf = turb.merge([sdf, aviner_rna])

lund_rna_sdf = turb.merge([sdf, lund_rna])
lund_prot_sdf = turb.merge([sdf, lund_prot])

vogel_rnaprot_sdf = turb.merge([sdf, "../../datasets/vogel_all.csv"])

# %% [markdown]
#### Stage 3. Generate correlations (spearman-rank, biserial correlation)

# %% [code]
def correlate_i(ds, y, x_add=None, metric="spearman"):
    _not = ("_id$", "^counter", object)
    _y = ds.view(y)
    _x = ds.view_not(y, *_not) if x_add is None else ds.view_not(y, x_add, *_not)
    return turb.correlate(data=ds.compute(turb.Pipe.no_id()), x=_x, y=_y)

@cache(filename="../../_intermediate/corr_datasets_fig3.csv")
def generate_super_corr():
    correlates = (
        correlate_i(vogel_rnaprot_sdf, x_add="prot_Da|mRNA_", y="prot_Da"),
        correlate_i(vogel_rnaprot_sdf, x_add="mRNA_|prot_Da", y="mRNA_"),
        correlate_i(av_prot_sdf, y="prot_[G12SM]"),
        correlate_i(av_tr_sdf, y="translation_[G12SM]"),
        correlate_i(av_rna_sdf, y="HeLa_"),
        correlate_i(lund_prot_sdf, y="INTENSITY_"),
        correlate_i(lund_rna_sdf, y="RPKM_")
    )
    return pd.concat(correlates,axis=0,sort=False)

# %% [code]
supers = generate_super_corr()

# %% [code]
supers.head(2)

# %% [code]
supers.n_, supers.p_

# %% [markdown]
### <a id='partial_calc'> Calculating Partial Correlations (to protein), Fixing mRNA </a>

Some of these functions are mildly expensive, so we cache the result, so that if you re-run it, you won't have to wait minutes.

# %% [markdown]
#### Declaring wrapping functions (to compute or cache)

# %% [code]
def correlate_j(ds, y, covar, exclude_extra=None, metric="spearman", verbose=0):
    """ Determine X, Y and Z groups in order to compute partial correlation."""
    # extract y and Z covar columns
    _Y = ds.view(y)
    _Z = ds.view(covar)
    # X is by default everything usable that's not y or a covar.
    if exclude_extra is None:
        _X = ds.view_not(y, covar, object, "_id$", "^counter", lambda z: z.nunique() == 1)
    else:
        _X = ds.view_not(y, covar, exclude_extra, object, "_id$", "^counter", lambda z: z.nunique() == 1)
    # compute changes using correlate
    return turb.correlate(data=ds.compute(turb.Pipe.no_id()),
                          x=_X, y=_Y, 
                          covar=_Z,
                          method=metric,
                          verbose=verbose)

@cache(filename="../../_intermediate/pcorr_datasets_fig3.csv")
def generate_partial_corr():
    """Partials with respect to mRNA length."""
    correlates = (
        correlate_j(aviner_sdf, "prot_[G12SM]", "HeLa_"),
        correlate_j(lund_sdf, "INTENSITY_", "RPKM_"),
        correlate_j(vogel_rnaprot_sdf, "prot_Daoy", "mRNA_Daoy"),
    )
    return pd.concat(correlates, axis=0, sort=False)

# %% [markdown]
#### Additional datasets (for partials)

# %% [code]
aviner_sdf = turb.merge([sdf, aviner_prot, aviner_rna])
lund_sdf = turb.merge([sdf, lund_all])

# %% [markdown]
#### Computation

# %% [code]
partials_prot = generate_partial_corr()

# %% [markdown]
### Creating Meta-Information (such as colormaps)

We use the meta data stored in SDF to group our data by colors that make sense for their group.

This gives us a color to plot the data scatter points with that may reveal further interesting trends.

In this case:
- Purple: PTM
- Red: Uniprot/Protein
- Green: Refseq/mRNA

# %% [code]
# convert colors to 255
grp1 = sdf.meta_.groupby(['data_source','data_groups'],as_index=False)['true_type'].count().dropna()
# calculate the subgroup of data sources
grp2 = grp1.groupby("data_source",as_index=False)["true_type"].agg(['sum','count']).drop("hgnc")
grp2['cm']=pd.Series({'ptm_source':"Purples",'refseq':"Greens",'uniprot':"Reds"})
# create color palettes
grp1['palette'] = pd.Series(
    it.chain.from_iterable(
       map(lambda c, text: cm.get_cmap(text)(np.linspace(.8, .2, c)), grp2['count'], grp2['cm'])
), index=grp1.index)

grp1['chex'] = turb.plot._palette._colormap_to_hex(grp1['palette'])

sdf.meta_['hex'] = sdf.meta_['data_groups'].cat.rename_categories(
    dict(zip(grp1['data_groups'].tolist(), grp1['chex'].tolist()))
)
sdf.meta_['source_str_color'] = sdf.meta_['data_source'].cat.rename_categories(grp2['cm'].to_dict())

# %% [markdown]
#### Defining pretty column name for legend

# %% [code]
clean_groups_abbr = {
    'hgnc': '', "id_col": "ID Columns",
    "ptm": "PTM", "codon_bias": "Codon Bias", "dinuc_freq": "DM Frequency",
    "l_prop": "Proportional\nLength", "length": "Sequence Length",
    "mononuc_freq": "MN Frequency", "rna_misc": "Misc. Feature",
    "spec_freq": "Special Frequency", "aa_freq": "AA Frequency",
    "biophysical": "AA Biophysical\nProperty","translation_freq": "Trans. Property",
    "free_energy": "MFE","aminoacid_meta": "AA Text-\nMined Feature"
}

clean_source_abbr = {"refseq": "Refseq", "ptm_source":"Phospho", "uniprot":"Uniprot"}

# %% [code]
# rebrand categories
grp1['data_groups_clean'] = grp1['data_groups'].cat.rename_categories(clean_groups_abbr)
grp1['data_source_clean'] = grp1['data_source'].cat.rename_categories(clean_source_abbr)
# reformat to make pretty name
grp1['pretty_group'] = turb.str.reformat("{data_groups_clean} ({data_source_clean})", grp1)

# %% [markdown]
### Pipeline for pivoting, aggregating `aviner` source data.

# %% [markdown]
In this version, we do not aggregate over all of the timesteps for HeLa, but keep G1 (2h), S (8h) and G2/M (12h) steps for mRNA and protein G1 (2h), S (8.5h) and G2/M (14h).


# %% [code]
cell_cycle_phases = ['G1',"S","G2M"]

# %% [code]
super_piv = (supers.copy()
                # pivot into a matrix
                .apply("pivot", "x", "y", "r")
                .update_meta()
                .drop("translation_")
                # aggregate together replicates and take the mean for mRNA, protein
                .aggregate_k("mean", 
                             names=["mrna_HeLa_"+x for x in cell_cycle_phases],
                             selectors=['HeLa_2hR_', 'HeLa_8hR_', 'HeLa_12hR_'])
                .aggregate_k("mean",
                             names=["prot_HeLa_"+x for x in cell_cycle_phases],
                             selectors=['prot_%s_r[0-9]' % x for x in cell_cycle_phases])
                # drop excess data
                .drop("_[1-3]$")
                # rename intensity, rpkm as mRNA/prot respectively
                .rename_axis([('INTENSITY','prot'), ('RPKM','mrna'), ("mRNA", "mrna")])
            )

# %% [markdown]
### Pipeline for pivoting, aggregating partial data.

Essentially repeating as above, with some small differences

# %% [code]
partial_piv = (partials_prot.copy()
                    .apply("pivot","x","y","r")
                    .update_meta()
                    .aggregate_k("mean", 
                                 names=["part_HeLa_"+x for x in cell_cycle_phases],
                                 selectors=['prot_%s_r[0-9]' % x for x in cell_cycle_phases],
                                 keep=False)
                   # rename intensity, rpkm as above
                    .rename_axis([('INTENSITY','part'), ('prot','part')])
                    .update_meta())

# %% [markdown]
### Merge together on index.

# %% [code]
# update sdf meta source so the bloody labels align - sigh
sdf.meta_.index = turb.str.string_replace(sdf.meta_.index, [("-","")])

# %% [code]
piv_join = turb.merge([super_piv, partial_piv], name="super_corr")
# extra crap here.
piv_join.name_ = "super_corr"
piv_join.df_['colors'] = sdf.meta_['hex']
piv_join.df_['data_source'] = sdf.meta_['data_source']
piv_join.df_['source_color'] = sdf.meta_['source_str_color']

# %% [markdown]
### Grouping expression values by source, type

- source: as in cell line
- type: as in expression type (mRNA, protein)

# %% [code]
cell_line = ('A431', "Daoy", "U251MG", "U2OS", "HeLa")
exprs = ('protein', 'mrna', 'partial')

# %% [code]
CACHES_ = dict(
    protein=("INTENSITY_","prot_"), 
    mrna=("RPKM_","mrna_","mRNA_"),
    partial="part_",
    aviner=("HeLa_G1", "HeLa_S", "HeLa_G2M"),
    lundberg=("_A431", "_U251MG", "_U2OS"),
    vogel="_Daoy"
)

# %% [code]
piv_join.cache_k(**CACHES_)
piv_join.meta_map("expr_type", exprs)
piv_join.meta_map("cell_line", cell_line)
piv_join.meta_map("cell_source", ['aviner', 'lundberg', 'vogel'])

# %% [code]
# write out the correlation matrix to file
piv_join.write('../../datasets/supercorr_fig3.json')

# %% [markdown]
### Scatterpoints of interest to mark

# %% [code]
# write some code here to annotate interesting points.

# %% [markdown]
## <a id='offic_plot'> Static Plot Version </a>

The version that will appear in the paper.

# %% [code]
def plot_matplotlib(ds):
    """A function to encapsulate the 'official' plot for Figure 3."""
    # figure object.
    section_size_vert = (8, 15, 15)
    section_size_cum = np.cumsum(section_size_vert)
    section_size_hoz = (2, 8, 2, 8)

    fig = plt.figure(figsize=(13, 16))
    
    # convenience function to draw some lines.
    def add_crosslines(a, bounds=.5):
        a.vlines([0], ymin=-bounds, ymax=bounds, linestyle=":")
        a.hlines([0], xmin=-bounds, xmax=bounds, linestyle=':')
        a.plot([-bounds, bounds], [-bounds, bounds], 'k--')
        a.grid()

    # specify our grid layout
    gs0 = GridSpec(sum(section_size_vert)+2, 21)
    # top two figures.
    ax_top1 = fig.add_subplot(gs0[:section_size_vert[0], 2:10])
    ax_top2 = fig.add_subplot(gs0[:section_size_vert[0], 12:20])
    # middle section
    gs00 = GridSpecFromSubplotSpec(8, 2, subplot_spec=gs0[section_size_vert[0]+2:section_size_cum[1], 2:10])
    ax_midleft1 = fig.add_subplot(gs00[:-4, :])
    ax_midleft2 = fig.add_subplot(gs00[-3:, 0])
    ax_midleft3 = fig.add_subplot(gs00[-3:, 1])
    gs01 = GridSpecFromSubplotSpec(8, 2, subplot_spec=gs0[section_size_vert[0]+2:section_size_cum[1], 12:20])
    ax_midright1 = fig.add_subplot(gs01[:-4, :])
    ax_midright2 = fig.add_subplot(gs01[-3:, 0])
    ax_midright3 = fig.add_subplot(gs01[-3:, 1])
    # bottom section
    gs02 = GridSpecFromSubplotSpec(8, 2, subplot_spec=gs0[section_size_cum[1]+2:section_size_cum[2], 2:10])
    ax_botleft1 = fig.add_subplot(gs02[:-4, :])
    ax_botleft2 = fig.add_subplot(gs02[-3:, 0])
    ax_botleft3 = fig.add_subplot(gs02[-3:, 1])
    gs03 = GridSpecFromSubplotSpec(8, 2, subplot_spec=gs0[section_size_cum[1]+2:section_size_cum[2], 12:20])
    ax_botright1 = fig.add_subplot(gs03[:-4, :])
    ax_botright2 = fig.add_subplot(gs03[-3:, 0])
    ax_botright3 = fig.add_subplot(gs03[-3:, 1])
    # text axes
    ax_yl = fig.add_subplot(gs0[:-2,0])
    ax_yr = fig.add_subplot(gs0[:-2,10])
    ax_xb = fig.add_subplot(gs0[-1, 1:])
    # text axes for journal sources
    ax_stop = fig.add_subplot(gs0[:section_size_vert[0], 20])
    ax_smid = fig.add_subplot(gs0[section_size_vert[0]+2:section_size_cum[1], 20])
    ax_sbot = fig.add_subplot(gs0[section_size_cum[1]+2:section_size_cum[2],20])
    
    # group together axes coherently.
    large_ = (ax_top1, ax_top2, ax_midleft1, ax_midright1, ax_botleft1, ax_botright1)
    small_ = (ax_midleft2,ax_midleft3,ax_midright2,ax_midright3,
              ax_botleft2,ax_botleft3,ax_botright2,ax_botright3)
    all_ = (ax_top1,ax_top2,ax_midleft1,ax_midleft2,ax_midleft3,
            ax_midright1,ax_midright2,ax_midright3,ax_botleft1,ax_botleft2,
            ax_botleft3, ax_botright1, ax_botright2, ax_botright3)
    text_ = (ax_yl, ax_yr, ax_xb, ax_stop, ax_smid, ax_sbot)
    big_labelled_ = (ax_top1, ax_midleft1, ax_botleft1)
    journal_ = (ax_stop, ax_smid, ax_sbot)
    
    # map together which cell line to each group of plots.
    pairings = [
        ("Daoy", ax_top1, ax_top2),
        ("U2OS", ax_midleft1, ax_midright1),
        ("A431", ax_midleft2, ax_midright2),
        ("U251MG", ax_midleft3, ax_midright3),
        ("HeLa_G1", ax_botleft1, ax_botright1),
        ("HeLa_S", ax_botleft2, ax_botright2),
        ("HeLa_G2M", ax_botleft3, ax_botright3),
    ]
    journal_tag = ('Vogel et al.(2010)', 'Lundberg et al.(2010)', 'Aviner et al.(2015)')
    
    # iterate over the pairings and plot the data.
    for cl, ax1, ax2 in pairings:
        # extract correct column.
        x_rna = ds.select("{} & mrna_".format(cl))[0]
        y_prot = ds.select("{} & prot_".format(cl))[0]
        y_part = ds.select("{} & part_".format(cl))[0]
        
        # draw points.
        ax1.scatter(ds[x_rna], ds[y_prot], c=ds['colors'], marker='o')
        ax2.scatter(ds[x_rna], ds[y_part], c=ds['colors'], marker='o')
            
        # add in cross lines
        add_crosslines(ax1)
        add_crosslines(ax2)
        # stick the cell line name at the top
        ax1.set_title(cl.replace("_"," "), fontsize=15)
        ax2.set_title(cl.replace("_"," "), fontsize=15)
    
    # some differences between the smaller and larger plots, like text size, locators.
    for a in large_:
        a.locator_params(axis="both", nbins=5)
        a.tick_params(axis="both", labelsize=12)
    for a in small_:
        a.locator_params(axis="both", nbins=3)
        a.tick_params(axis="both", labelsize=9)
    
    # plot text.
    text_kwargs = dict(ha="center", va="center", fontsize=15)
    ax_xb.text(.5, .5, r"SDF-to-mRNA level correlation ($r_s$|$r_{{bs}}$)", **text_kwargs)
    ax_yl.text(.5, .5, "SDF-to-protein level correlation ($r_s$|$r_{bs}$)", rotation=90, **text_kwargs)
    ax_yr.text(.5, .5, "SDF-to-protein level partial correlation, fixing mRNA level ($r_s$|$r_{bs}$)",
               rotation=90, **text_kwargs)
    
    # iterate over journal source labels and add
    for a, text in zip(journal_, journal_tag):
        a.text(.5, .5, text, **text_kwargs, rotation=-90)
    
    # prepare text ticks and despining for labels.
    for a in text_:
        mf.despine(a)
        a.tick_params(axis="both", which="both", bottom=False, right=False, 
                      left=False, top=False, labelbottom=False, labelleft=False,
                      labelright=False, labeltop=False)
    
    
    # prep legend
    new_legend = [Line2D([0], [0], color=c, lw=4) for c in grp1['chex'].values]
    # add legend
    ax_midright1.legend(new_legend, grp1['pretty_group'].values.tolist(),
                       bbox_to_anchor=(1.15, 0.9),shadow=True,fancybox=True,
                    ncol=1, title="Data Groups", fontsize=11)

    # add big label
    for i, a in enumerate(big_labelled_):
        a.text(-0.05, 1.05, string.ascii_uppercase[i], transform=a.transAxes, size=25, weight="bold")
    
    # save figure
    _ = turb.plot.save(fig, 
                plot_type="scatter", 
                save_types=('png','pdf','svg', 'eps'),
                name="SDF_corrcorr",
                savemode="update",
                fp="../../Figures/SDF/",
                dpi=1200)
    
    plt.show()

# %% [markdown]
## <a id='desc'> Description </a>

### Variations in *a priori* experimental dataset assumptions considerably alter SDF impactfulness.

    Scatterplots of Spearman-rank ($r_s$)/Biserial correlation ($r_{bs}$)-mixed correlation between sequence-derived features (SDFs) and mRNA (x-axis), protein (y-axis, left) and partial protein fixing mRNA (y-axis, right). From top; (A) Daoy (Vogel et al.), (B) U2OS;A431;U251MG (Lundberg et al.), and (C) HeLa (Aviner et al.) from different mRNA-protein expression datasets. Each point represents a correlation between an SDF and an expression dataset. See Methods for details on correlation method. mRNA/RNA features are denoted in green shades, amino-acid features are denoted in red shades, PTM features are denoted in purple.

# %% [code]
plot_matplotlib(piv_join)

# %% [markdown]
## <a id='bokeh'> Interactive (Bokeh) Plot Versions </a>
#### Imports

# %% [code]
# bokeh
from bokeh.plotting import figure, output_file, show, ColumnDataSource
from bokeh.models import HoverTool, BoxZoomTool, ResetTool, BoxSelectTool, LinearColorMapper, \
    ColorBar, BasicTicker, Div
from bokeh.models.widgets import CheckboxButtonGroup
from bokeh.io import output_notebook
from bokeh.layouts import row, column, gridplot
from bokeh import transform
from bokeh.palettes import mpl
output_notebook()

# %% [markdown]
#### Preprocessing

# %% [code]
# shifting meta information into the updated MetaPandas.
sdf.meta_['data_groups_clean'] = sdf.meta_['data_groups'].cat.rename_categories(clean_groups_abbr)
piv_join.df_['data_groups_clean'] = sdf.meta_['data_groups_clean']

# %% [code]
cell_line_long = ('A431 (Lundberg et al.)', "Daoy (Vogel et al.)",
                  "U251MG (Lundberg et al.)", "U2OS (Lundberg et al.)",
                  "HeLa (Aviner et al.)", "HeLa (Aviner et al.)", "HeLa (Aviner et al.)")

# %% [markdown]
#### renaming pipe

# %% [code]
index_clean_map = [['pH_', ''],
    ['_prop', ' rel.'],
    ['_cds', ' CDS'],
    ['_utr5', " 5'UTR"],
    ['_utr3', " 3'UTR"],
    ['_win', ' (win'],
    ['_total', ''],
    ['_bp', ''],
    ['_bcaa', ''],
    ['mfe', 'free energy'],
    ['_mrna', ' mRNA'],
    ['_aa', ' (amino acid)'],
    ['_ptmc', ' PTM'],
    ['_fc', ''],
    ['_pc', ' (corrected)'],
    ['_', ' ']]

# %% [markdown]
#### Pair together cell line and expression data

# %% [code]
pairings = it.product(cell_line,exprs)

# %% [markdown]
#### Create dataset and plot

# %% [code]
def plot_bokeh_main(ds, dataset, filename="../../Figures/SDF/_interactive/Fig3.html"):
    def_fig_width = 350
    def_fig_height = 250
    
    title="""Fig 3. Variations in a priori experimental dataset assumptions considerably alter SDF impactfulness."""
    
    desc="""Scatterplots of Spearman-rank ($r_s$)/Biserial correlation ($r_{bs}$)-mixed correlation between sequence-derived features (SDFs) and mRNA (x-axis), 
    protein (y-axis, left) and partial protein fixing mRNA (y-axis, right). From top; (A) Daoy (Vogel et al.), (B) U2OS;A431;U251MG (Lundberg et al.), 
    and (C) HeLa (Aviner et al.) from different mRNA-protein expression datasets. Each point represents a correlation between an SDF and an expression dataset.
    See Methods for details on correlation method.
    mRNA/RNA features are denoted in green shades, amino-acid features are denoted in red shades, PTM features are denoted in purple.
    """
    
    output_file(filename, title=title)

    def lines(p):
        p.line([-.5, .5], [-.5, .5], color="black")
        p.line([0, 0], [-.5, .5], line_dash="dotted", color="black")
        p.line([-.5, .5], [0, 0],line_dash="dotted", color="black")

    column_sets = []

    for cl, cll in zip(cell_line, cell_line_long):
        mRNA_f = ds.select("{} & mrna".format(cl))[0]
        prot_f = ds.select("{} & protein".format(cl))[0]
        part_f = ds.select("{} & partial".format(cl))[0]

        p1 = figure(plot_width=def_fig_width, plot_height=def_fig_height,
                    title=cll,
                    x_axis_label="Correlation to mRNA level",
                    y_axis_label="Correlation to protein level")
        p2 = figure(plot_width=def_fig_width, plot_height=def_fig_height,
                    title=cll,
                    x_axis_label="Correlation to mRNA level",
                    y_axis_label="Partial Correlation to protein level")
        lines(p1)
        lines(p2)
        circ1 = p1.circle(mRNA_f, prot_f, size=7,
                          fill_color="colors", source=dataset, alpha=.8)
        circ2 = p2.circle(mRNA_f, part_f, size=7,
                          fill_color="colors", source=dataset, alpha=.8)
        
        # hovertool
        hover1 = HoverTool(tooltips=[
            ("Feature", "@counter"),
            ("Group", "@data_groups_clean"),
            ("corr-to-mRNA", "@%s{1.111}" % mRNA_f),
            ("corr-to-protein", "@%s{1.111}" % prot_f)
        ], renderers=[circ1])
        hover2 = HoverTool(tooltips=[
            ("Feature", "@counter"),
            ("Group", "@data_groups_clean"),
            ("corr-to-mRNA", "@%s{1.111}" % mRNA_f),
            ("corr-to-partial", "@%s{1.111}" % part_f)
        ], renderers=[circ2])

        p1.add_tools(hover1)
        p2.add_tools(hover2)
        
        # add to column sets
        column_sets.append(row(p1, p2))
    
    # create divs for title and description
    dd_title = Div(text="<b>{}</b>".format(title), width=def_fig_width*2, height=20)
    dd_desc = Div(text=desc, width=def_fig_width*2, height=150)
    
    show(column(dd_title, dd_desc, column(column_sets)))

# %% [code]
piv_join_clean = piv_join.copy().apply_index("lower").rename_axis(index_clean_map, axis=0)

# %% [code]
piv_join_clean.select("HeLa & protein")[0]

# %% [code]
piv_join_clean.select("HeLa & part")[0]

# %% [code]
piv_join_clean.select("HeLa & mrna")[0]

# %% [code]
dset = ColumnDataSource(piv_join_clean.df_)
_ = plot_bokeh_main(piv_join_clean, dset, filename="../../Figures/SDF/_interactive/Fig3.html") #

# %% [markdown]
# <a id='supp1'> Supplementary Material (1) </a>

Relationship for SDF variables *controlling for LENGTH* across the datasets.

Previously we calculate the partial correlation *controlling for mRNA/RNA level*.

* mRNA objects: control for mRNA length
* CDS objects: CDS length control
* 3'UTR objects: 3'UTR length control
* 5'UTR objects: 5'UTR length control
* AA objects: amino acid length control

#### Mapping region to a length column.

# %% [code]
region_length = {"mrna_obj": "length_mrna", "cds_obj": "length_cds",
                 "protein_obj": "length_aa", "utr5_obj": "length_utr5", 
                 "utr3_obj": "length_utr3"}

# %% [markdown]
#### Compute partial correlation using respective length for each block.


$\rho(X, Y | Z)$, where $Z$ is appended at the end of each section to identify which length variable was used.

# %% [code]
@turb.cache(filename="../../_intermediate/pcorr_datasets_supp_length_fig3.csv")
def generate_partial_length_corr(verbose=0):
    """Partials with respect to mRNA/amino acid length, where appropriate."""
    correlates = []
    # for each subregion, compute correlations and stack
    for region, length_feature in region_length.items():
        correlates.append(correlate_j(av_prot_sdf, 
                                      "prot_[G12SM]",
                                      length_feature, verbose=verbose).assign(Z=length_feature))
        correlates.append(correlate_j(lund_sdf, "INTENSITY_", 
                                      length_feature, verbose=verbose).assign(Z=length_feature))
        correlates.append(correlate_j(vogel_rnaprot_sdf, "prot_Daoy", 
                                      length_feature, verbose=verbose).assign(Z=length_feature))
    return pd.concat(correlates,axis=0,sort=False)

# %% [markdown]
### Pivot, add metainformation

# %% [code]
av_prot_sdf.downcast()
lund_sdf.downcast()
vogel_rnaprot_sdf.downcast()

# %% [code]
part_length = generate_partial_length_corr()

# %% [code]
length_piv = (part_length.copy()
     .apply("pivot_table", index='x', columns='y', values='r')
     .update_meta()
     .aggregate("mean", name="prot_HeLa", selector="prot_[G12SM]", keep=False)
     .rename_axis([('INTENSITY','part'), ('prot','part')])
     .update_meta())

# %% [code]
length_piv_join = turb.merge([super_piv, length_piv], name="length_corr")
length_piv_join.df_['colors'] = sdf.meta_['hex']
length_piv_join.df_['data_groups_clean'] = sdf.meta_['data_groups_clean']

length_piv_join.cache_k(**CACHES_)

length_piv_join.meta_map("expr_type", exprs)
length_piv_join.meta_map("cell_line", cell_line)
length_piv_join.meta_map("cell_source", ['aviner', 'lundberg', 'vogel'])

# %% [markdown]
### Define Difference Function

# %% [code]
length_piv_clean = length_piv_join.copy().apply_index("lower").rename_axis(index_clean_map, axis=0)

# %% [code]
dset_l = ColumnDataSource(length_piv_clean.df_)

# %% [code]
def plot_bokeh_diff(ds, dataset, filename=None, Z="Length", title=None, desc=None):
    def_fig_width = 300
    def_fig_height = 300
    
    if title is None:
        title="Supplementary 3. Correlation changes when partials are factored in, factoring %s." % Z
    if desc is None:
        desc="""Scatterplots of the change in correlation from partial-correlation to protein factoring %s to mRNA (x) against 
        partial-correlation to protein factoring %s to protein (y). Large delta changes indicate a significant impact of %s factor
        in correcting the correlation, indicating potential feature redundancy, or features that need heavy correcting for.
        Useful in the knowledge of which features can aid in appropriate mRNA proxying for protein abundance.""" % (Z, Z, Z)
    
    output_file(filename=filename, title=title)
    
    def grouper(n, iterab):
        args = [iter(iterab)]*n
        return it.zip_longest(*args)
    
    
    def lines(p):
        p.line([0, 0], [-.5, .5], line_dash="dotted", color="black")
        p.line([-.5, .5], [0, 0],line_dash="dotted", color="black")

    column_sets = []

    for cl, cll in zip(cell_line, cell_line_long):
        mRNA_f = ds.select(cl + "& mrna")[0]
        prot_f = ds.select(cl + "& protein")[0]
        part_f = ds.select(cl + "& partial")[0]
        # compute differences
        dataset.data['diff_prot_%s' % cl] = ds[part_f] - ds[prot_f]
        dataset.data['diff_mRNA_%s' % cl] = ds[part_f] - ds[mRNA_f]
        
        p1 = figure(plot_width=def_fig_width, plot_height=def_fig_height,
                    title=cll,
                    x_axis_label="Correlation change Partial-mRNA",
                    y_axis_label="Correlation change Partial-Prot")
        lines(p1)
        circ1 = p1.circle('diff_mRNA_%s' % cl, 'diff_prot_%s' % cl, size=7,
                          fill_color="colors", source=dataset, alpha=.8)
        
        # hovertool
        hover1 = HoverTool(tooltips=[
            ("Feature", "@counter"),
            ("Group", "@data_groups_clean"),
            ("corr-to-mRNA", "@%s{1.111}" % mRNA_f),
            ("corr-to-protein", "@%s{1.111}" % prot_f),
            ("corr-to-partial", "@%s{1.111}" % part_f),
            ("part-mRNA-diff", "@%s{1.111}" % ('diff_mRNA_%s' % cl)),
            ("part-prot-diff", "@%s{1.111}" % ('diff_prot_%s' % cl))
        ], renderers=[circ1])

        p1.add_tools(hover1)
        
        # add to column sets
        column_sets.append(p1)
    
    column_sets = tuple(grouper(3, column_sets))
    
    # create divs for title and description
    dd_title = Div(text="<b>{}</b>".format(title), width=def_fig_width*3, height=20)
    dd_desc = Div(text=desc, width=def_fig_width*3, height=100)

    show(column(dd_title, dd_desc, gridplot(column_sets)))

# %% [markdown]
## Plot: Correlation Changes w.r.t mRNA level

where $Z=\text{mRNA}$

# %% [code]
_ = plot_bokeh_diff(length_piv_clean, dset_l, filename="../../Figures/SDF/_interactive/supp1_Fig3.html", Z="mRNA")

# %% [markdown]
## Plot: Correlation changes w.r.t Sequence Length

where $Z=L_{\text{source}}$

# %% [code]
_ = plot_bokeh_diff(length_piv_clean, dset_l, filename="../../Figures/SDF/_interactive/supp2_Fig3.html", Z="Length")

# %% [code]
```
