## Supplementary material for "Predicting Relative Protein Abundance via Sequence-Based Information": S3 Data: SDF-Official3Supp2.ipynb

```
# %% [markdown]
# <a id='supp2'> Supplementary Material (2): CCA </a>

Canonical Correlation Analysis will allow us to analyse the correlation between features in our SDF matrix and the Vogel SDF matrix used.


# %% [code]
import os
import matplotlib.pyplot as plt
import numpy as np
import seaborn as sns
import pandas as pd
from matplotlib import cm
import matplotlib as mpl
from scipy.cluster.hierarchy import linkage, dendrogram
from scipy import stats
import itertools as it
import string
%matplotlib inline

from matplotlib import patches as mpatches
from matplotlib.gridspec import GridSpec, GridSpecFromSubplotSpec
from matplotlib.lines import Line2D

import sys
sys.path.insert(0, "../../scripts/")
sys.path.insert(0, "../../../turbopanda/")
# scripts
import misc_functions as misc
import version
import myfig as mf
# turbopanda
import turbopanda as turb

# %% [code]
from sklearn.cross_decomposition import PLSRegression

# %% [markdown]
### Preparing vogel dataset

# %% [code]
vogel = turb.read("../../data/vogel_dataset_rna-prot.xls", name='Vogel', header=5,sheet_name="MARS_data")
vogel

# %% [markdown]
### Our dataset


# %% [code]
sdf = turb.read("../../datasets/SDF_scaled2.json", 
                name="SDF", with_clean=False)
sdf

# %% [code]
# sdf_cl = sdf.compute(turb.Pipe.no_id())

# %% [markdown]
### Merge

# %% [code]
cca_ds = turb.merge([sdf, vogel])
cca_ds.meta_split_category("datasets")

# %% [markdown]
### Prep Model

# %% [code]
cca = PLSRegression(2, scale=True, max_iter=1000)

# %% [markdown]
#### Fetch subset columns

# %% [code]
sdf_v = turb.utils.intersect(cca_ds.view("is_SDF"), 
                             cca_ds.view_not(object, "_id$", lambda z: z.nunique()==1))
vog_v = turb.utils.intersect(cca_ds.view("is_Vogel"), 
                                         cca_ds.view_not(object, "_id$", lambda z: z.nunique()==1))

# %% [markdown]
#### Drop missing rows

# %% [code]
cca_ds.apply("dropna", subset=turb.utils.join(sdf_v, vog_v))

# %% [markdown]
**NOTE**: Only 334 genes actually map across the two datasets, which is quite shocking.

# %% [code]
cca_ds.n_, cca_ds.p_

# %% [markdown]
### Fit

# %% [code]
cca.fit(cca_ds[sdf_v], cca_ds[vog_v])

# %% [code]
cca_ds.meta_['uniqable'] = cca_ds.df_.nunique()>1

# %% [markdown]
## Plots

What is most odd is the fact that barely any of my features correlate strongly with Vogels' dataset.

# %% [code]
fig, ax = plt.subplots(figsize=(16,13))
sns.heatmap(cca_ds[cca_ds.select("~object & ~_id$ & uniqable")].corr(), cmap="seismic", center=0, ax=ax)
plt.show()

# %% [markdown]
A couple of the features appear to heavily correlate with a large selection of mine, but the vast majority do not.. hmm.

# %% [code]
betas = pd.DataFrame(cca.coef_, index=sdf_v, columns=vog_v)
betas['Nc'].sort_values()

# %% [code]
sns.clustermap(betas, cmap="seismic", center=0)

# %% [code]
X_c, Y_c = cca.transform(cca_ds[sdf_v], cca_ds[vog_v])

# %% [markdown]
#### Compressed into 2D

Stronger correlation in reduced space along the diagonal.

# %% [code]
fig,ax=plt.subplots(ncols=2, figsize=(11,4))
ax[0].scatter(X_c[:, 0], Y_c[:, 0])
ax[1].scatter(X_c[:, 1], Y_c[:, 1])
print("Dim1: {:0.3f}\nDim2: {:0.3f}".format(
    np.corrcoef(X_c[:, 0], Y_c[:, 0])[0, 1],
    np.corrcoef(X_c[:, 1], Y_c[:, 1])[0, 1]
))

# %% [markdown]
... substantially no correlation further apart between dimensions.

# %% [code]
fig,ax=plt.subplots(ncols=2, figsize=(11,4))
ax[0].scatter(X_c[:, 0], X_c[:, 1])
ax[1].scatter(Y_c[:, 0], Y_c[:, 1])
print("Dim1: {:0.3f}\nDim2: {:0.3f}".format(
    np.corrcoef(X_c[:, 0], X_c[:, 1])[0, 1],
    np.corrcoef(Y_c[:, 0], Y_c[:, 1])[0, 1]
))
```
