## Supplementary material for "Predicting Relative Protein Abundance via Sequence-Based Information": S3 Data: SDF-Official4ML.ipynb

```
# %% [markdown]
# Figure 4

What is contained within this notebook:

To navigate the notebook, click the links below or scroll:

### Computation

1. **[Initialize and import](#init)**
2. **[Computing mRNA vs. protein relationship models](#lm)**
2. **[Computing SDF vs. protein relationship models](#sdflm)**
3. **[Determining weights for each feature using whole SDF (best models) vs. protein relationships](#fi)**
3. **[Comparison to Vogels' sequence-derived features](#compvogel)**
5. **[Computing Reduced SDF vs. protein relationship models](#pcared)**
6. **[Computing comparisons mRNA vs. protein, SDF vs. protein and combinations](#compare)**

### Visualization

1. **[Defining the Official Plot](#officplot)**
2. **[Plot Description](#desc)**


## Objectives

To calculate and plot the results of a machine learning analysis on the full and reduced input data subsets, both on our dataset and comparing to vogels' SDF features.

#### Product Table of Variables

| Cell Line | Model type | Input matrix type | Includes mRNA as input |
| -- | -- | -- | -- |
| Daoy; HeLa (multiple); ... | OLS; Ridge; Lasso; ... | PCA (full); FA; ... | Yes/No |

--- 
* __Cell Lines__: Daoy; HeLa (3); U521MG; A431; U2OS
* __Model type__: OLS; Ridge; Elastic Net; SVM; Random Forest; Decision Tree; AdaBoost; GradientBoostedTree; KNN
* __Input design matrix__: Whole data; PCA; PCA (by data group); Factor Analysis (FA)
* __Input sources__: Us; Vogel et al., 2010

Further to this, we compare between models with/without mRNA expression corresponding to the protein level in question.

# %% [code]
import matplotlib.pyplot as plt
import numpy as np
import itertools as it
import seaborn as sns
import pandas as pd
%matplotlib inline

from matplotlib import patches as mpatches
from matplotlib.gridspec import GridSpec, GridSpecFromSubplotSpec

import sys
sys.path.insert(0, "../../../turbopanda/")
sys.path.insert(0, "../../scripts/")
# scripts
import version

# turbopanda
import turbopanda as turb
from turbopanda import cache

# %% [markdown]
## Version

# %% [code]
version.current()
print("turbopanda: %s" % turb.__version__)
print(version.last_run())

# %% [code]
from turbopanda.utils import join, umap, retuple
from turbopanda.str import patproduct, string_replace
from turbopanda import melt

# %% [markdown]
## <a id='init'> SDF Official Machine Learning Models of SDF against protein (and mRNA) levels </a>

### Import the various datasets

#### 1. Original SDF dataset

# %% [code]
sdf = turb.read("../../datasets/SDF_scaled2.json", 
                name="SDF")
sdf

# %% [markdown]
#### 2. Preprocessed PCA/Factor Analysis datasets

# %% [code]
sdf_pcas = turb.read("../../datasets/pca_reduced/*.csv", name=('FA', 'PCA_DG', 'PCA_W'), index_col=0)
sdf_pcas

# %% [markdown]
#### 3. Expression datasets (Aviner et al., Lundberg et al., Vogel et al.)

# %% [code]
aviner_rna, aviner_prot, aviner_tr = turb.read("../../datasets/clean/*_aviner_adv.csv", 
                    name=("mRNA_aviner", "prot_aviner", "tr_aviner"), index_col=0)

# %% [code]
lund_all, lund_prot, lund_rna = turb.read("../../datasets/lundberg_*.csv", 
                                          name=('Lundberg', 'Lundberg_Prot', 'Lundberg_RNA'))

# %% [code]
vogel_expr = turb.read("../../datasets/vogel_all.csv")

# %% [markdown]
#### 4. Vogels' SDF dataset

# %% [code]
vogel_sdf = turb.read("../../data/vogel_dataset_rna-prot.xls", name='Vogel', 
                      header=5, sheet_name="MARS_data")
vogel_sdf

# %% [markdown]
### Minor preprocessing of expression datasets

Aviners' dataset needs to aggregate over the replicates, we ignore the translation data.

# %% [code]
aviner_rna.aggregate_k("mean", 
                        names=["mRNA_HeLa_%dh" % i for i in [0, 2, 4, 6, 8, 12]], 
                        selectors=["_%s_" % i for i in ['0hB','2hR', '4hR', '6hR', '8hR', '12hR']], 
                        keep=False)

aviner_prot.aggregate_k("mean", 
                        names=["prot_HeLa_%s" % i for i in ["G1", "S", "G2M"]], 
                        selectors=["_%s_" % i for i in ["G1", "S", "G2M"]], 
                        keep=False)

# %% [code]
aviner_rna.update_meta()
aviner_prot.update_meta()

# %% [markdown]
Now for aviner we need to join together the mRNA and protein for combined datasets for where we need like-for-like between corresponding mRNA and protein levels for various models.

For this we need to merge them together, but bare in mind that the labels for this dataset are duplicated.

# %% [code]
aviner_rna_prot = (turb.merge([aviner_rna, aviner_prot], name="Aviner", verbose=2)
                    .apply("drop_duplicates", subset='gene_names'))
aviner_rna_prot

# %% [markdown]
### Label dataset in HGNC

# %% [code]
hgnc = turb.read("../../datasets/global_id.csv")
# make gene id ensembl unique
hgnc.apply("drop_duplicates", subset=['ensembl_gene_id'])
hgnc.apply("dropna", subset=['ensembl_gene_id'])

# %% [markdown]
### Unify naming conventions

Different datasets use different naming conventions to describe mRNA or protein level data.

# %% [code]
aviner_rna_prot.rename_axis([('_2h',"_G1"), ('_8h',"_S"), ('_12h','_G2M')])

# %% [code]
lund_all.rename_axis([('RPKM','mRNA'), ('INTENSITY','prot')])
lund_prot.rename_axis([('INTENSITY','prot')])
lund_rna.rename_axis([('RPKM','mRNA')])

# %% [markdown]
## <a id='lm'> Computing basic mRNA - protein relationships </a>

In this step, we'll see the relationship between mRNA and protein with machine learning models and see how much mRNA can infer on protein concentration without including extra features.

We undergo the following steps:

1. Preprocess to get input $X$ and target $y$
2. Perform linear regression models (OLS) with mRNA as input and protein as target, with 5-fold cross validation scoring using $R^2$, the coefficient of determination

# %% [code]
from sklearn.model_selection import cross_val_score, cross_validate
from sklearn.linear_model import LinearRegression

# %% [code]
def generate_basic_linear_model_scores(dataset, x: str, y: str, preproc=True):
    if preproc:
        X, y = turb.ml.preprocess_continuous_X_y(dataset.df_, x, y)
    else:
        X, y = np.asarray(dataset[x]).reshape(-1, 1), np.asarray(dataset[y])
    return cross_val_score(LinearRegression(), X, y, scoring="neg_root_mean_squared_error", cv=5)

# %% [code]
cell_line = ('U2OS','A431','U251MG','Daoy','HeLa G1','HeLa S','HeLa G2M')
cell_line_dirty = tuple(f.replace(" ", "_") for f in cell_line)

# %% [code]
lmcomb = (
    [lund_all]*3 + [vogel_expr] + [aviner_rna_prot]*3,
    patproduct("mRNA_%s", cell_line_dirty),
    patproduct("prot_%s", cell_line_dirty)
)

# %% [markdown]
### Generating basic linear model plots

We generate a boxplot of the resulting scores of those models.

Here we see that, with a basic linear model, we get RMSE scores between 0.9 to 0.92 across multiple different cell lines, whereby the aim is to minimize this error.

Here error refers to 5-fold cross-validation.

# %% [code]
mrna_protein_results = pd.DataFrame(
    map(generate_basic_linear_model_scores, *lmcomb),
    index=pd.Index(cell_line, name="Cell_Line"),
    columns=patproduct("split%d_test_score", range(5))
)

# %% [code]
fig, ax1 = plt.subplots(figsize=(8, 5))
turb.plot.widebox(-mrna_protein_results.T, ax=ax1,
                        vert=True, measured=r"RMSE (OLS)", outliers=False)
ax1.set_xlabel("Cell Line")
ax1.set_title("mRNA vs protein level models across different cell lines")
plt.show()

# %% [markdown]
## <a id='sdflm'> Computing SDF-to-protein relationships </a>

In this use case we use all SDF-compatible features into a SDF-expression matrix to predict protein and/or mRNA. We break this into the following steps:

1. Merge SDF dataset into expression dataset.
2. Perform primitive feature selection to drop useless SDF features, if they exist.
3. Design a sklearn pipeline with grid search over hyperparameters for the best ML model
4. Select best model and use scores for comparison.

Here is our product table of competing variables

| Cell Line | Model type | Input matrix type | Includes mRNA as input |
| -- | -- | -- | -- |
| Daoy; HeLa (multiple); ... | OLS; Ridge; Lasso; ... | PCA (full); FA; ... | Yes/No |

# %% [markdown]
### Defining pre-filtering functions

# %% [code]
# perform some pre-filtering.
# missing value filter
def mvf(df, thresh=.3):
    return df.drop(df.columns[(1. - (df.count() / df.shape[0])) > thresh], axis=1)
# low variance filter
def lvf(df, thresh=1e-6):
    fcols = df.select_dtypes(float)
    return df.drop(fcols.columns[fcols.var() < thresh], axis=1)

# %% [markdown]
### Defining our expression to SDF groups

# %% [code]
sdf_to_protein_groups = list(zip(
    [sdf]*7, 
    [lund_prot]*3 + [vogel_expr] + [aviner_prot]*3,
    ["SDF_%s_prot" % (f.replace(" ","_")) for f in cell_line],
    ["prot_%s" % (f.replace(" ","_")) for f in cell_line],
    ["~hgnc & ~object & ~^prot & ~^mRNA & ~^HeLa_prot"]*7
))

# %% [code]
sdf_to_protein_groups_t = retuple(sdf_to_protein_groups)

# %% [markdown]
### Defining Pipeline and GridSearch area

Here we search over a selection of different models:

**LINEAR**:

1. Ordinary Least Squares (LinearRegression)
2. Ridge
3. Elastic Net (A combination of Ridge and Lasso regularization)

**NON-LINEAR**:

1. Support Vector Machine (SVM)
2. Decision Tree
3. Random Forest (of Decision Trees)
4. AdaBoost
5. Gradient Boosted Decision Trees
6. K-Nearest-Neighbors

# %% [code]
from sklearn.pipeline import Pipeline
from sklearn.model_selection import GridSearchCV
from sklearn.linear_model import LinearRegression, Ridge, ElasticNet
from sklearn.svm import SVR
from sklearn.tree import DecisionTreeRegressor
from sklearn.ensemble import RandomForestRegressor, AdaBoostRegressor, GradientBoostingRegressor
from sklearn.neighbors import KNeighborsRegressor

ntree_steps = sorted([10, 15, 20, 25, 50, 100, 125, 150, 200])

mypipe = Pipeline([('model', LinearRegression())])
params = [
    {'model': [LinearRegression()]},
    {
        'model': [Ridge()],
        'model__alpha': np.logspace(-4., 1.5, 50),
    },
    {
        'model': [ElasticNet()],
        'model__alpha': np.logspace(-4., 1.5, 50),
        'model__l1_ratio': [.2, .5, .8],
    },
    {
        'model': [SVR()],
        'model__C': np.logspace(-2., 2., 50),
    },
    {
        'model': [DecisionTreeRegressor()],
        'model__max_depth': [1, 2, 3],
        'model__max_features': ['auto', 'sqrt'],
    },
    {
        'model': [RandomForestRegressor()],
        'model__n_estimators': ntree_steps,
        'model__max_features': ['auto', 'sqrt'],
    },
    {
        'model': [AdaBoostRegressor(), GradientBoostingRegressor()],
        'model__n_estimators': ntree_steps,
        'model__learning_rate': np.logspace(-3, -1, 8),
    },
    {
        'model': [KNeighborsRegressor()],
        'model__n_neighbors': np.arange(3, 20, 3, dtype=np.int),
        'model__weights': ['uniform','distance'],
    }
]

# %% [markdown]
### Define model selection function, with run

The steps are as follows:

1. Merge into the SDF dataset
2. Pipe to prefilter useless columns
3. Select X
4. Preprocess into design matrix X and y target
5. Prepare iterative function to gridsearch on data
6. Fit each dictionary of gridsearch parameters into a mappable chunked
7. Concat results and return

# %% [code]
def computing_sdf_to_protein_model(sdfs, expression, name, ycol, xsel, verbose=2):
    # all of this goes inside the wrapper...
    
    def _initial_grid(param):
        # merge together the sequence-derived features to the expression dataset
        merge_tuple = [sdfs, expression] if not isinstance(expression, (list, tuple)) else join([sdfs], expression)
        sdf_expr = turb.merge(merge_tuple, verbose=0, name=name)
        # pipe in pre-filtering functions
        sdf_pipe = sdf_expr.copy().apply("pipe", mvf).apply("pipe", lvf)
        # get x selected
        xselected = sdf_pipe.select(xsel)
        # preprocess and split dataset into X and y
        X, y = turb.ml.preprocess_continuous_X_y(sdf_pipe, xselected, ycol)
        # use mypipe and params to generate ML models in cached chunks
        gs = GridSearchCV(
            mypipe, param, scoring="neg_root_mean_squared_error",
            n_jobs=-3, cv=5, refit=False, verbose=1
        )
        gs.fit(X, y)
        # returns a dictionary, which is joblib.dump able
        return gs.cv_results_
    
    # run as mappable cached with chunks - this takes a while
    res_map = turb.utils.umapcc(cache_dir+name+".pkl", _initial_grid, params)
    # concat together chunks into df - with model
    res_df = pd.concat(map(lambda x: pd.DataFrame(x), res_map), axis=0, sort=False)
    res_df['model'] = res_df['param_model'].astype(str).str.split("(", expand=True)[0]
    # return result total
    return res_df

# %% [markdown]
### Run Model selection: **WARNING** Will take a long time the first time

This section is running SDF-to-protein machine learning model selection where SDF is the whole input feature set with minimal preprocessing, with NO mRNA input.

# %% [code]
cache_dir = "../../_intermediate/sdf_to_prot/"
expression_set_run = list(map(computing_sdf_to_protein_model, *sdf_to_protein_groups_t))

# %% [markdown]
### Defining a plot method for each of these SDF-to-protein models

Here we take across the grid search, for each algorithm type, which parameters did the best by selecting **the maximum negative root-mean-squared-error metric**

In this plot, we draw a box for each machine learning model, the error 
    is the 5-fold cross-validation for the test score.
    Different axes refer to each cell line dataset.

# %% [code]
def plot_expression_run(data, ax):
    """
    In this plot, we draw a box for each machine learning model, the error 
    is the 5-fold cross-validation for the test score.
    Different axes refer to each cell line dataset.
    """
    indices = data.reset_index(drop=True).groupby("model")['mean_test_score'].idxmax()
    # get data
    best_mo = data.reset_index(drop=True).loc[indices.values, :]
    # rearrange to keep split scores
    best_split = best_mo[turb.str.pattern("split[0-9]+_test_score", best_mo)].T
    best_split.columns = indices.index
    # plot as widebox
    _ = turb.plot.widebox(best_split, measured="Negative RMSE", vert=False, ax=ax, outliers=False)
    return best_split

# %% [markdown]
### Plotting for each expression dataset, each *best model* scenario

For all datasets, a **gradient-boosted tree algorithm** gave the highest accuracy, of around 0.7 to 0.8 RMSE. The only exception to this is the Daoy cell line, where an ElasticNet algorithm only slightly outperforms.

# %% [code]
fig, ax = turb.plot.gridplot(8, arrange="square", ax_size=4)
_ = list(map(plot_expression_run, expression_set_run, ax))
_ = list(map(lambda t, axes: axes.set_title(t), sdf_to_protein_groups_t[2], ax))
ax[-1].set_visible(False)
fig.tight_layout()

# %% [markdown]
## <a id="fi"> Determining the best features from our best model(s) by cell line that are effective against protein level </a>

Firstly we need to **re-run** our models and extract the model with its coefficients.

#### Defining how to extract the 'best model' from our gridsearch

# %% [code]
def best_model(expr):
    # sklearn packages
    from sklearn import linear_model, tree, svm, ensemble, neighbors
    packages = (linear_model, tree, svm, ensemble, neighbors)
    _expr = expr.reset_index(drop=True)
    best_model_indx = _expr.loc[_expr['mean_test_score'].idxmax(), :]
    mod_n = best_model_indx["model"]
    # select the model object
    mod = [getattr(obj, mod_n) for obj in packages if hasattr(obj, mod_n)][0]()
    # fit the model with the parameters
    pname, pval = turb.utils.dictsplit(best_model_indx['params'])
    # remove model from name and truncate
    _pname = string_replace(pname[1:], [("model__", "")])
    # fix parameters with set_params
    mod.set_params(**dict(zip(_pname, pval[1:])))
    return mod


def get_feature_weights(model):
    # check to see which attribute it has
    _options = ('coef_', 'feature_importances_')
    
    if hasattr(model, "feature_importances_"):
        return model.feature_importances_
    elif hasattr(model, "coef_"):
        # convert to fi before returning
        # get a maximum value so we can divide by sum for normalization, should sum == 1 for relative importance
        _abs = np.abs(model.coef_)
        return _abs / np.sum(_abs)
    else:
        raise ValueError("model '{}' doesn't have [coef_, feature_importances_] attribute.".format(model))


def computing_best_model_coefficients(results, sdfs, expression, name, ycol, xsel, verbose=2):
    # all of this goes inside the wrapper...
    
    def _initial_grid(best_model):
        # merge together the sequence-derived features to the expression dataset
        merge_tuple = [sdfs, expression] if not isinstance(expression, (list, tuple)) else join([sdfs], expression)
        sdf_expr = turb.merge(merge_tuple, verbose=0, name=name)
        # pipe in pre-filtering functions
        sdf_pipe = sdf_expr.copy().apply("pipe", mvf).apply("pipe", lvf)
        # get x selected
        xselected = sdf_pipe.select(xsel)
        # preprocess and split dataset into X and y
        X, y = turb.ml.preprocess_continuous_X_y(sdf_pipe, xselected, ycol)
        # cross validate
        est = cross_validate(best_model, X, y, 
                             scoring="neg_root_mean_squared_error",
                             cv=5, n_jobs=5, return_estimator=True,
                             verbose=1)
        # get each estimator and create a table of feature importances...
        fis = pd.DataFrame(dict(zip(
            range(5),
            map(get_feature_weights, est['estimator'])
        )), index=xselected)
        
        return fis
    
    # run as mappable cached with chunks - this takes a while
    res_map = turb.utils.cache(cache_dir+name+".pkl", _initial_grid, best_model(results))
    # return result total
    return res_map

# %% [code]
model_coefficient_groups_t = retuple(list(zip(
    expression_set_run,
    [sdf]*7, 
    [lund_prot]*3 + [vogel_expr] + [aviner_prot]*3,
    ["fi_%s" % (f.replace(" ","_")) for f in cell_line],
    ["prot_%s" % (f.replace(" ","_")) for f in cell_line],
    ["~hgnc & ~object & ~^prot & ~^mRNA & ~^HeLa_prot"]*7
)))

# %% [code]
cache_dir = "../../_intermediate/sdf_to_prot/"
best_coefficients_ = list(map(computing_best_model_coefficients, *model_coefficient_groups_t))

# %% [markdown]
### Plotting the 'steepness' of coefficient dependency for each cell line

# %% [code]
fig, ax = plt.subplots()
for bcf, cl in zip(best_coefficients_, cell_line):
    ax.plot(range(50), 
            np.cumsum(bcf.mean(axis=1).sort_values(ascending=False))[:50],
            '*--',
            label=cl)
ax.legend()
ax.set_xlabel("Sorted Features")
ax.set_ylabel("Feature Importance (%)")
plt.show()

# %% [markdown]
### Selecting the $K$-best features by importance for each cell line

# %% [code]
def select_kbest_features_importance(bcf, cl, K=6):
    # take the mean of the FIs
    bcf_k = bcf.mean(axis=1).nlargest(K)
    # extract best rows and return
    return melt(bcf.loc[bcf_k.index], var_name="splits", value_name="Feature Importance").assign(Cell_Line=cl)

# %% [code]
df_kbest_coefficients = (pd.concat(map(select_kbest_features_importance, 
                                       best_coefficients_, cell_line))
    .reset_index(drop=True))

# %% [markdown]
## <a id='compvog'> Comparison to Vogels' derived feature set </a>

Here we will compare the results of our full SDF-to-protein dataset to the features extracted by Vogel et al., 2010.

### Defining our new expression set (with vogel SDF)

We merge this into HGNC to get a label which Vogel can bind to:

# %% [code]
# here we 'de-duplicate aviner prot for this set due to duplicate labels
decoupled_aviner_prot = turb.merge([hgnc, aviner_prot.copy().apply("drop_duplicates", "gene_name")],verbose=2)

# %% [code]
# group zippables: sdf dataset, expression dataset, name to give, y column, x selector

vogel_to_protein_groups = tuple(zip(
    [vogel_sdf]*7, 
    [lund_prot]*3 + [vogel_expr] + [decoupled_aviner_prot]*3,
    ["vog_%s_prot" % cl for cl in cell_line_dirty],
    ["prot_%s" % cl for cl in cell_line_dirty],
    ["~hgnc & ~object & ~^prot & ~mRNA & ~Protein"]*7
))

# %% [markdown]
### Run Model Selection (using Vogel), *warning takes a while*

This shouldn't take anywhere near as long however, due to the much smaller sample sizes.

# %% [code]
vogel_to_protein_groups_t = retuple(vogel_to_protein_groups)

# %% [code]
cache_dir = "../../_intermediate/sdf_to_prot/"
expression_set_vog = list(map(computing_sdf_to_protein_model, *vogel_to_protein_groups_t))

# %% [markdown]
### Plotting for each expression dataset, each *best model* scenario for Vogels dataset

One of the key differences between our dataset and Vogels is a substantial change in sample size: we're sampling between 30-40% of the human proteome, whereby Vogels' set is a fraction of that.

Further to this, our SDF features seem to perform *better* than Vogels' set, even though there is a smaller sample size.

# %% [code]
fig, ax = turb.plot.gridplot(8, arrange="square", ax_size=4)
_ = list(map(plot_expression_run, expression_set_vog, ax))
_ = list(map(lambda t, axes: axes.set_title(t), vogel_to_protein_groups_t[2], ax))
ax[-1].set_visible(False)
fig.tight_layout()

# %% [markdown]
## Compare sample sizes between our and Vogels dataset (for each expression dataset)

Here we see that our sequence-derived features capture much more of the whole human proteome than any dataset before.

# %% [code]
def computing_dataset_sizes(sdfs, expression, name, ycol, xsel):
    # if expression is one, just list them, else use join
    merge_tuple = [sdfs, expression] if not isinstance(expression, (list, tuple)) else join([sdfs], expression)
    sdf_expr = turb.merge(merge_tuple, verbose=0, name=name)
    sdf_pipe = sdf_expr.copy().apply("pipe", mvf).apply("pipe", lvf)
    xselected = sdf_pipe.select(xsel)
    X, y = turb.ml.preprocess_continuous_X_y(sdf_pipe, xselected, ycol)
    # now return the size of sdf_expr
    return X.shape

# %% [code]
n_sample_size_sdf = list(map(computing_dataset_sizes, *sdf_to_protein_groups_t))
n_sample_size_vog = list(map(computing_dataset_sizes, *vogel_to_protein_groups_t))

# %% [code]
# map them together into dataframe, and plot
N_sample_sizes = pd.DataFrame({
    "Parkes": map(lambda x: x[0], n_sample_size_sdf),
    "Vogel": map(lambda x: x[0], n_sample_size_vog)
}, index=cell_line)

ax_1 = turb.plot.widebar(N_sample_sizes, c=['#444444', '#bbbbbb'], measured=r"Number of genes")
ax_1.set_xlabel("Expression dataset")
plt.show()

# %% [markdown]
## Comparison between best score model for SDF (whole) vs. Vogels SDF sets (factoring in sample size)

There isn't a particularly large difference in score between our features and Vogel-based models. But it is hard to quantify how much of this is because of the smaller dataset on proteins we know more about now.

# %% [code]
import sklearn as skl

# %% [code]
def best_score(expr):
    return -expr['mean_test_score'].max()

def best_cv_score(expr):
    # reorder index
    _expr = expr.reset_index(drop=True)
    # filter for split values, get location and transform by absolute
    return _expr.filter(regex="split*").loc[_expr['mean_test_score'].idxmax(), :].transform(np.abs)

def best_model_name(expr):
    expr_r = expr.reset_index(drop=True)
    return expr_r.loc[expr_r['mean_test_score'].idxmax(), 'model']

# %% [code]
dataset_names = ('Parkes', 'Vogel')

comp_vogel_sdf_plot = pd.DataFrame({
    "Parkes": list(map(best_score, expression_set_run)),
    "Vogel": list(map(best_score, expression_set_vog))
}, index=cell_line)

ax_2 = turb.plot.widebar(comp_vogel_sdf_plot, c=['#444444', '#bbbbbb'], measured="RMSE (Best Model)")
ax_2.set_xlabel("Expression dataset")
plt.show()

# %% [markdown]
## <a id='pcared'> Computing reduced (PCA) SDF-to-protein relationships </a>

Here we repeat what we did as above using our sequence-derived feature dataset, but first we are drawing on the **reduced PCA** models which we calculated in the *SDF-Reduction* iPython notebook as our primary input matrix.

For the majority of these features, often we don't have a direct feature name but a **Factor** on which to draw from in terms of interpretation - but we see modest improvements in the *scalability* and *generalization* capacity of the model.

# %% [code]
reduced_sdf_to_protein_groups = list(zip(
    join(*[[s]*7 for s in sdf_pcas]), 
    join(*[[lund_prot]*3 + [vogel_expr] + [aviner_prot]*3]*3),
    join(*[["red_%s_%s_prot" % (f.replace(" ","_"), t) for f in cell_line] for t in ['FA', 'PCA_DG', 'PCA_W']]),
    ["prot_%s" % (f.replace(" ","_")) for f in cell_line]*3,
    ["~hgnc & ~object & ~^prot & ~^mRNA & ~^HeLa_prot"]*21
))

# %% [code]
reduced_sdf_to_protein_groups_t = retuple(reduced_sdf_to_protein_groups)

# %% [code]
expression_set_redsdf = list(map(computing_sdf_to_protein_model, *reduced_sdf_to_protein_groups_t))

# %% [markdown]
### Factor Analysis Results

# %% [code]
fig, ax = turb.plot.gridplot(8, arrange="square", ax_size=4)
_ = list(map(plot_expression_run, expression_set_redsdf[:7], ax))
_ = list(map(lambda t, axes: axes.set_title(t), vogel_to_protein_groups_t[2], ax))
ax[-1].set_visible(False)
fig.tight_layout()

# %% [markdown]
### PCA (Data groups)

# %% [code]
fig, ax = turb.plot.gridplot(8, arrange="square", ax_size=4)
_ = list(map(plot_expression_run, expression_set_redsdf[7:14], ax))
_ = list(map(lambda t, axes: axes.set_title(t), vogel_to_protein_groups_t[2], ax))
ax[-1].set_visible(False)
fig.tight_layout()

# %% [markdown]
### PCA (Whole)

# %% [code]
fig, ax = turb.plot.gridplot(8, arrange="square", ax_size=4)
_ = list(map(plot_expression_run, expression_set_redsdf[14:], ax))
_ = list(map(lambda t, axes: axes.set_title(t), vogel_to_protein_groups_t[2], ax))
ax[-1].set_visible(False)
fig.tight_layout()

# %% [markdown]
## Plot between best score model for SDF (various) vs. Vogels SDF sets (factoring in sample size)

Basically not a huge difference, although the normal SDF dataset does seem to do a bit better, this is always a tradeoff between model performance (in this example) and generalizability to other datasets.

# %% [code]
dataset_names2 = ('SDF (Norm)', "SDF (FA)", "SDF (PCA_DG)", "SDF (PCA_W)", 'Vogel')
datasets_5 = (expression_set_run, expression_set_vog, expression_set_redsdf[:7], 
              expression_set_redsdf[7:14], expression_set_redsdf[14:])

comp_vogel_sdf_plot2 = pd.DataFrame(
{m: list(map(best_score, ds)) for m, ds in zip(dataset_names2, datasets_5)}
, index=cell_line)

ax = turb.plot.widebar(comp_vogel_sdf_plot2, measured="RMSE (Best Model)", cmap="Set3")
ax.set_title("Comparison of best scores between ours and Vogels datasets - without mRNA")
ax.set_xlabel("Expression dataset")
ax.legend(loc='lower right')
plt.show()

# %% [markdown]
## <a id='compare'> Comparions between mRNA-to-protein, SDF-to-protein and SDF+mRNA-to-protein </a>

First off we need to run models that **factor in mRNA** as a input to the model.

# %% [code]
def chunks(lst, n):
    """Yield successive n-sized chunks from lst."""
    for i in range(0, len(lst), n):
        yield lst[i:i + n]

# %% [code]
# here we 'de-duplicate aviner-rna-prot for this set due to duplicate labels
decoupled_aviner_rna_prot = turb.merge([hgnc, aviner_rna_prot],verbose=2).apply("drop_duplicates", "ensembl_gene_id")

# %% [code]
expr_7set = [lund_all]*3 + [vogel_expr] + [aviner_rna_prot]*3
vogexpr_7set = [lund_all]*3 + [vogel_expr] + [decoupled_aviner_rna_prot]*3

yc7 = patproduct("prot_%s", cell_line_dirty)
xsel_w_rna = "~hgnc & ~object & ~id & ~^prot & ~Protein & ~^HeLa_prot"
# first cell line, X second model input type (PCA, FA, whole, Vogel, etc)
fn_tmplt = "%s_%s_plus_mrna_to_prot"

dataset_names3 = ('SDF_Norm', "SDF_FA", "SDF_PCA_DG", "SDF_PCA_W", 'Vogel')

sdfsets = [[s]*7 for s in (sdf, sdf_pcas[0], sdf_pcas[1], sdf_pcas[2], vogel_sdf)]
exprsets = [expr_7set]*4 + [vogexpr_7set]
namesets = list(chunks(patproduct(fn_tmplt, dataset_names3, cell_line_dirty), 7))
ynsets = [yc7]*5
xselset = [patproduct("mRNA_%s | "+xsel_w_rna, cell_line_dirty)]*5
# iterate on the dataset_names
fullset_args = tuple(zip(join(*sdfsets), join(*exprsets), join(*namesets), join(*ynsets), join(*xselset)))

fullset_args_t = retuple(fullset_args)

# %% [markdown]
### Run Methods here

# %% [code]
def computing_sdf_withrna_to_protein_model(sdfs, expression, name, ycol, xsel, verbose=2):
    # all of this goes inside the wrapper...
    
    def _initial_grid(param):
        # merge together the sequence-derived features to the expression dataset
        merge_tuple = [sdfs, expression] if not isinstance(expression, (list, tuple)) else join([sdfs], expression)
        sdf_expr = turb.merge(merge_tuple, verbose=0, name=name)
        # pipe in pre-filtering functions
        sdf_pipe = sdf_expr.copy().apply("pipe", mvf).apply("pipe", lvf)
        # get x selected
        xselected = sdf_pipe.select(xsel).tolist() + ['mRNA_%s' % ycol.split("_", 1)[-1]]
        # preprocess and split dataset into X and y
        X, y = turb.ml.preprocess_continuous_X_y(sdf_pipe, xselected, ycol)
        # use mypipe and params to generate ML models in cached chunks
        gs = GridSearchCV(
            mypipe, param, scoring="neg_root_mean_squared_error",
            n_jobs=-3, cv=5, refit=False, verbose=1
        )
        gs.fit(X, y)
        # returns a dictionary, which is joblib.dump able
        return gs.cv_results_
    
    # run as mappable cached with chunks - this takes a while
    res_map = turb.utils.umapcc(cache_dir+name+".pkl", _initial_grid, params)
    # concat together chunks into df - with model
    res_df = pd.concat(map(lambda x: pd.DataFrame(x), res_map), axis=0, sort=False)
    res_df['model'] = res_df['param_model'].astype(str).str.split("(", expand=True)[0]
    # return result total
    return res_df

# %% [code]
cache_dir = "../../_intermediate/sdf_with_mrna_to_prot/"
sdf_plus_mrna_to_protein_groups = list(map(computing_sdf_withrna_to_protein_model, *fullset_args_t))

# %% [markdown]
### Map cell line, X input to our result data

# %% [code]
clxin = tuple(it.product(cell_line, dataset_names3))
assign_clxin = lambda df, clarg: df.assign(Cell_Line=clarg[0], X_input=clarg[1])

# %% [code]
sdf_plus_mrna__results = pd.concat(
    tuple(map(assign_clxin, sdf_plus_mrna_to_protein_groups, clxin))
, axis=0, sort=False).assign(With_Expression="SDF plus mRNA")

# %% [markdown]
### Do the same for the w/out mRNA group + add label

# %% [code]
sdf_sin_mrna__results = pd.concat(tuple(map(assign_clxin, join(*datasets_5), clxin))
    , axis=0, sort=False).assign(With_Expression="SDF (alone)")

# %% [markdown]
### Concatenate together

# %% [code]
sdf_model__results = (pd.concat(
    [sdf_sin_mrna__results, sdf_plus_mrna__results]
, axis=0, sort=False).reset_index(drop=True)
[['model', 'Cell_Line', 'X_input', 'With_Expression', 'mean_test_score'] + patproduct("split%d_test_score", range(5))])


# %% [markdown]
### Filter for the best model for each X_input / Cell line / W-out mRNA combinations

# %% [code]
best_indices = (sdf_model__results.groupby(['X_input', 'Cell_Line', 'With_Expression'])
['mean_test_score'].idxmax().values)

# %% [code]
filtered_sdf_model__results = sdf_model__results.loc[best_indices]

# %% [markdown]
### Add column to differentiate between SDF (our) and Vogel source

# %% [code]
filtered_sdf_model__results['source'] = ((filtered_sdf_model__results['X_input'] == 'Vogel')
                                             .map({False: "SDF", True: "Vogel"}))

# %% [markdown]
### Melt down splits

# %% [code]
from turbopanda import melt
from turbopanda.str import pattern

# %% [markdown]
## Plotting RMSE by cell line including our mRNA-protein data

#### 1. Appending the mRNA-protein data into the same table

# %% [code]
sdf_mrna_model__results = pd.concat([
    filtered_sdf_model__results, 
    (mrna_protein_results
     .reset_index()
     .pipe(lambda df: df.assign(mean_test_score=df[pattern("split*", filtered_sdf_model__results)].mean(axis=1)))
     .assign(source="mRNA", X_input='mRNA', With_Expression="mRNA (alone)",model="OLS"))
], sort=False).reset_index(drop=True)

# %% [markdown]
#### 2. Groupby and select the best X input for each cell line

# %% [code]
choose_best_xinput = sdf_mrna_model__results.groupby(['Cell_Line','With_Expression'])['mean_test_score'].idxmax()

# %% [markdown]
#### 2. Melt down RMSE scores into one col

# %% [code]
cl_by_split_expression = melt(
    # input
    sdf_mrna_model__results.loc[
        choose_best_xinput
    ].drop(['mean_test_score'], axis=1),
    # args
    value_vars="split*", var_name="splits",
    value_name="RMSE", include_index=False
).pipe(turb.pipe.absolute, "RMSE")

# %% [code]
def sdf_only_filtered(df):
    fdf = df.query("With_Expression == 'SDF (alone)'")
    idx = fdf.groupby(['Cell_Line', 'X_input'])['mean_test_score'].idxmax().values
    melt_fdf = melt(
        fdf.loc[idx].drop("mean_test_score", axis=1),
        value_vars="split*", include_index=False, var_name="splits", value_name="RMSE"
    ).pipe(turb.pipe.absolute, "RMSE")
    # clean the names within a column
    melt_fdf['X_input_clean'] = string_replace(melt_fdf['X_input'], [('DG', 'Data Groups'), ('_W', ' Full'), ("_",' ')])
    return melt_fdf

# %% [code]
from turbopanda.pipe import replace
hues2 = ('SDF_FA', 'SDF_Norm', 'SDF_PCA_DG', 'SDF_PCA_W', 'Vogel')

# %% [markdown]
## <a id='officplot'>Official Plot</a>

# %% [code]
import pylab

# %% [markdown]
#### Prepare axes 3 data...

Here for each cell line, we keep the K largest features to display... as a graph

# %% [code]
# pretty names
pretty_names = [("pH_",""), ("_prop"," rel."),
        ("_cds"," CDS"),("_utr5"," 5'UTR"), ("_utr3"," 3'UTR"),
        ("_win"," (win"), ("_total",""), ("_bp",""), ("_bcaa",""),
        ("mfe","free energy"), ("_mrna"," mRNA"), ("_aa", " (amino acid)"),
        ("_ptmc", " PTM"), ("_fc",""),("_pc"," (corrected)"), ("_"," ")]

df_kbest_coefficients['colnames_pretty'] = string_replace(df_kbest_coefficients['colnames'], pretty_names)
df_kbest_coefficients['Feature Importance'] *= 100.

# %% [code]
df_kbest_coefficients.head()

# %% [code]
sns.boxplot(data=df_kbest_coefficients, x="Feature Importance", y="colnames_pretty",
            hue="Cell_Line")

# %% [markdown]
#### Matplotlib defaults

# %% [code]
params = {'legend.fontsize': 12,
          'legend.handlelength': 2,
          'legend.labelspacing':0.25}
pylab.rcParams.update(params)

# %% [markdown]
#### Prepare axes 1 data...

# %% [code]
comp_vogel_sdf_plot__split = pd.concat(map(
    lambda ds, n: melt(pd.DataFrame(map(best_cv_score, ds), index=cell_line),
                    var_name="splits", value_name="RMSE").assign(Source=n),
    (expression_set_run, expression_set_vog), dataset_names)
).rename(columns={"index":"Cell Line"})

# %% [markdown]
#### Pre-look at axes 1

# %% [code]
sns.barplot(data=comp_vogel_sdf_plot__split, x="Cell Line", 
            y="RMSE", hue="Source", order=cell_line,
            palette=['#444444', '#bbbbbb'])

# %% [code]
import string

# %% [code]
def official_plot():
    fig = plt.figure(figsize=(13, 16))
    gs0 = GridSpec(9, 2)
    ax1 = fig.add_subplot(gs0[:3, 0])
    ax2 = fig.add_subplot(gs0[:3, 1])
    ax3 = fig.add_subplot(gs0[3:6, :])
    ax4 = fig.add_subplot(gs0[6:, :])
    axes = [ax1, ax2, ax3, ax4]

    # fig, axes = turb.plot.gridplot(4, 'square', (5, 4))

    first_two_hue_color = ("#fa9fb5", "#43a2ca")

    # 1. Comparison between ours and Vogels SDF feature scores.

    sns.barplot(data=comp_vogel_sdf_plot__split, x="Cell Line", 
                y="RMSE", hue="Source", order=cell_line, ax=axes[0], 
                capsize=.12, palette=first_two_hue_color)
    axes[0].set_ylim(.5, 1.)
    axes[0].set_yticks([.5, .6, .7, .8, .9, 1.])
    axes[0].tick_params("x", rotation=30)
    axes[0].set_xlabel("Cell Line", fontsize=14)
    axes[0].set_ylabel(r"RMSE ($\pm$ 5-fold CV)", fontsize=14)
    axes[0].legend(loc='lower left', title="Source", prop={"size": 13})
    axes[0].grid(axis="y")
    # 2. Comparison between ours and Vogels data retention
    turb.plot.widebar(N_sample_sizes, ax=axes[1], c=first_two_hue_color,
                      vert=True, measured=r"Number of genes",
                      label_rotation=30)
    axes[1].legend(loc="lower left", title="Source", prop={"size": 13})
    axes[1].set_xlabel("Cell Line", fontsize=14)
    axes[1].set_ylabel("Number of genes", fontsize=14)
    axes[1].set_yscale("log")
    axes[1].set_yticks([10, 100, 1000, 10000])
    axes[1].grid(axis="y")
    # 3. Relative Feature Importance across Cell Lines
    kb_order = (df_kbest_coefficients.groupby("colnames_pretty")['Feature Importance']
                    .mean().sort_values().index)
    ax2pal = ('#1669f7', '#3d7ab8', '#30accf', '#46e302', '#e69832', '#e3732d', '#fa817a')
    sns.barplot(data=df_kbest_coefficients, x="colnames_pretty", hue="Cell_Line",
                y="Feature Importance", ax=axes[2], 
                order=kb_order[::-1], hue_order=cell_line,
                palette=ax2pal, capsize=.05)
    axes[2].set_ylabel("Feature Importance %\n""($\pm$ 5-fold CV)", fontsize=14)
    axes[2].set_xlabel("Input feature", fontsize=14)
    axes[2].legend(loc="upper center", title="Cell Line", ncol=2, prop={"size": 13})
    axes[2].grid(axis="y")
    # 4. Comparison between mRNA alone, SDF alone and SDF + mRNA models
    ax3pal = ('#8eba43', '#eb8a44', '#7d4427')
    sns.barplot(data=cl_by_split_expression, x="Cell_Line", 
                  y="RMSE", hue="With_Expression",
                  order=comp_vogel_sdf_plot.index, 
                  palette=ax3pal, capsize=.12,
                  hue_order=('mRNA (alone)', 'SDF (alone)', 'SDF plus mRNA'),
                  ax=axes[3])
    axes[3].set_ylim(.5, 1.)
    axes[3].set_yticks([.5, .6, .7, .8, .9, 1.])
    axes[3].set_ylabel(r"RMSE ($\pm$ 5-fold CV)", fontsize=14)
    axes[3].set_xlabel("Cell Line", fontsize=14)
    axes[3].tick_params("x", rotation=30, size=12)
    axes[3].legend(loc='lower center', title="ML Input Source", ncol=3, prop={"size": 13})
    axes[3].grid(axis="y")
    
    for i, a in enumerate(axes):
        a.text(-0.05, 1.05, string.ascii_uppercase[i],
               transform=a.transAxes, size=25, weight="bold")
    
    fig.tight_layout()
    
    # save figure
    _ = turb.plot.save(fig, 
                plot_type="bar", 
                save_types=('png','pdf','svg', 'eps'),
                name="SDF_ML",
                savemode="update",
                fp="../../Figures/SDF/",
                dpi=1200)
    plt.show()

# %% [markdown]
## <a id='desc'>Description</a>

### Substantial improvement in SDF inference capacity with respect to protein concentration to mRNA

    Barplots of (A) RMSE estimations ($\pm$ 5-fold CV) as a result of ML regression models across different cell lines for whole input sources, (B) Sample sizes for each resultant cell line ML model by input source, (C) Feature weights as a relative importance by the 6 most important features by cell line, (D) Whole genome RMSE estimations ($\pm$ 5-fold CV) by cell line depending on input features (using just mRNA expression, just SDFs, or both). See Methods and Supplementary Material for details on model and feature preprocessing/selection.

# %% [code]
official_plot()
```
