## Supplementary material for "Predicting Relative Protein Abundance via Sequence-Based Information": S3 Data: SDF-Official5GO.ipynb

```
# %% [markdown]
# Figure 5

What is contained within this notebook:

To navigate the notebook, click the links below or scroll:

### Initialization and Computation

1. **[Initialize and import](#init)**
2. **[Import best ML models from previous chapter](#mlmodels)**
3. **[Computing SDF vs. protein models by Gene Ontology](#sdflm)**
4. **[Preparing Results](#result)**

### Visualization

1. **[Pre-plot sanity checks](#sanity)**
2. **[Defining the Official Plot](#offic_fig)**
3. **[Plot Description](#desc)**


## Objectives

To calculate the scores of machine learning models by specific biological process, such as cell cycle, transcription factor, and so on. We can do this on our different cell lines.

#### Product Table of Variables

--- 
* __Cell Lines__: Daoy; HeLa (3); U521MG; A431; U2OS
* __Model type__: OLS; Ridge; Elastic Net; SVM; Random Forest; Decision Tree; AdaBoost; GradientBoostedTree; KNN
* __Input design matrix__: Whole data; PCA; PCA (by data group); Factor Analysis (FA)
* __Input sources__: Us; Vogel et al., 2010

# %% [code]
import matplotlib.pyplot as plt
import numpy as np
import re
import seaborn as sns
import pandas as pd
import networkx as nx
%matplotlib inline

from matplotlib import patches as mpatches

import sys
sys.path.insert(0, "../../../turbopanda/")
sys.path.insert(0, "../../scripts/")
# scripts
import version

# turbopanda
import turbopanda as turb

# %% [markdown]
## Version

# %% [code]
version.current()
print("turbopanda: %s" % turb.__version__)
print(version.last_run())

ds_dir = "../../_intermediate/sdfds_half/"

# %% [code]
from turbopanda.utils import umap, retuple, union, absdifference, join, zipe, cache
from turbopanda.str import patproduct, string_replace, pattern
from turbopanda import melt

from turbopanda import cache as cachedec

# %% [markdown]
## <a id='init'> SDF Official Machine Learning Models for Gene Ontology Comparison </a>

### Import the various datasets

#### 1. Original SDF dataset

# %% [code]
sdf = turb.read("../../datasets/SDF_scaled2.json", 
                name="SDF")
sdf

# %% [markdown]
#### 2. Expression datasets (Aviner et al., Lundberg et al., Vogel et al.)

# %% [code]
aviner_rna, aviner_prot, aviner_tr = turb.read("../../datasets/clean/*_aviner_adv.csv", 
                    name=("mRNA_aviner", "prot_aviner", "tr_aviner"), index_col=0)
lund_all, lund_prot, lund_rna = turb.read("../../datasets/lundberg_*.csv", 
                                          name=('Lundberg', 'Lundberg_Prot', 'Lundberg_RNA'))
vogel_expr = turb.read("../../datasets/vogel_all.csv", name="Vogel_EXPR")

# %% [markdown]
#### 3. Vogels' SDF dataset

# %% [code]
vogel_sdf = turb.read("../../data/vogel_dataset_rna-prot.xls", name='Vogel_SDF', 
                      header=5, sheet_name="MARS_data")
vogel_sdf

# %% [markdown]
#### 4. HGNC dataset

# %% [code]
hgnc = turb.read("../../datasets/global_id.csv", name="HGNC")
# make gene id ensembl unique
hgnc.apply("drop_duplicates", subset=['ensembl_gene_id'])
hgnc.apply("dropna", subset=['ensembl_gene_id'])

# %% [markdown]
### 5. GO dataset

Head to [Gene Ontology](http://geneontology.org/docs/go-annotation-file-gaf-format-2.1/) website to see how the .obo files are oriented. There are currently no **easy** way to cleanly access this filetype. I find the GOATOOLS package for python absolutely disgusting and a hideous maze of cryptic functions and objects.

# %% [code]
def read_obo_file(fn):
    # find all terms
    with open(fn) as f:
        lines = f.read().splitlines()
    # get line counts of where each term starts
    term_starts = np.asarray([i for i, line in enumerate(lines) if re.match("\[Term\]", line)]) + 1
    end_point = [i for i, line in enumerate(lines[term_starts[-1]:]) if line == ""][0]
    term_ends = np.hstack((term_starts[1:]-2, np.array([end_point])))
    terms = []
    for t_start, t_end in zip(term_starts, term_ends):
        # dictionary ify the subject matter inside this
        try:
            d = dict([l.split(": ", 1) for l in lines[t_start:t_end]])
            terms.append(d)
        except Exception:
            print(t_start, t_end)
    # convert to pandas.dataframe
    return pd.DataFrame(terms)

# %% [markdown]
#### GO Reference

GODB.id is a unique column, so we can use this for nodes.

# %% [code]
namespaces = ('biological_process', 'molecular_function', 'cellular_component')

# %% [code]
go_db = (read_obo_file("../../data/GO/go-basic.obo")
             .query("is_obsolete != 'true'")
             .drop(['replaced_by', 'is_obsolete', 'comment', 'consider', 'subset', 'xref'], axis=1)
             .dropna(how="all")
             .pipe(turb.pipe.replace, "name", [("_", " ")]))

# %% [code]
go_db.head(3)

# %% [markdown]
### Create a network from the GO terms to gain information on term depth

In order to 'prioritise' higher up terms, we need to determine the depth by creating a Graph of the terms, to do this we will use the `networkx` package and build the graph manually using nodes and edges.

Using NX, we also know the **direction** of the graph, using the `is_a` attribute, children go to the parent node. Hence we can use a `nx.DiGraph` object.

For Biological Process, Cellular Component and Molecular Function, we have a *separate* DiGraph, as these are defined as separate within the Gene Ontology documentation.

Firstly we define the 'parent' node from the relationship var.

# %% [code]
def fetch_go_id(n):
    return go_db.loc[go_db['name'] == n, 'id'].iloc[0]

# %% [markdown]
#### Make an edge list by determining parent id

# %% [code]
go_db['parent'] = go_db['is_a'].str.split(" ! ", expand=True)[0]

# %% [code]
fetch_go_id('biological process')

# %% [code]
fetch_go_id('molecular function')

# %% [code]
fetch_go_id('cellular component')

# %% [markdown]
#### Instantiate and fill nodes, edges

# %% [code]
def fetch_edgelist(nms):
    return (go_db.query("namespace == @nms")
            .dropna(subset=['is_a'])
            [['parent', 'id']]
            .values.tolist())

def fetch_nodelist(nms):
    return (go_db.query("namespace == @nms")
           ['id'].values.tolist())

# %% [code]
# instantiate graphs
G_bp = nx.DiGraph()
G_cc = nx.DiGraph()
G_mf = nx.DiGraph()
# define root nodes.
G_bp.add_nodes_from(fetch_nodelist("biological_process"))
G_mf.add_nodes_from(fetch_nodelist("molecular_function"))
G_cc.add_nodes_from(fetch_nodelist("cellular_component"))
# add in all nodes via edge list
G_bp.add_edges_from(fetch_edgelist("biological_process"))
G_cc.add_edges_from(fetch_edgelist("cellular_component"))
G_mf.add_edges_from(fetch_edgelist("molecular_function"))

# %% [markdown]
#### Terms that are one-level down from 'biological process'

# %% [code]
termsbp_d1 = go_db.loc[go_db['id'].isin(G_bp.neighbors("GO:0008150")),'name'].values

# %% [markdown]
### GO-to-ensembl mapping

# %% [code]
go_ensembl = (pd.read_csv("../../data/biomart/biomart_go.csv", index_col=0)
    # drop any row which has no GO_ID
    .dropna(subset=['go_id'])
    .drop(['ensembl_peptide_id','ensembl_transcript_id','name_1006'], axis=1)
    .drop_duplicates())

# %% [markdown]
#### map this to GO_DB to get namespace and other useful attrs

# %% [code]
go_detailed = turb.merge([go_db.drop(['def', 'alt_id', 'synonym', 'relationship'], axis=1),
                          go_ensembl], verbose=2).drop("index__").df_

# %% [markdown]
#### split datasets into GOBP, GOCC and GOMF GO IDs

# %% [code]
def concat_strings_per_go(df):
    # firstly lets do some cleaning of the columns to prevent NaN spam
    df['name'] = df['name'].str.strip()
    df['namespace'] = df['namespace'].str.strip()
    df['ensembl_gene_id'] = df['ensembl_gene_id'].str.strip()

    return pd.concat([
        df.query("namespace == @nms").groupby("ensembl_gene_id")['name'].apply(lambda x: x.str.cat(sep=";")).to_frame(nms)
    for nms in namespaces], axis=1, sort=False)

# %% [code]
go_collapsed = concat_strings_per_go(go_detailed)

# %% [code]
print(go_collapsed.shape)
go_collapsed.head(3)

# %% [markdown]
#### Append an 'all' column to GO

# %% [code]
go_collapsed['go_all'] = go_collapsed['biological_process'].fillna("") +\
                                ";" + go_collapsed['molecular_function'].fillna("") + \
                                ";" + go_collapsed['cellular_component'].fillna("")

# %% [markdown]
### Sparse format for GO terms

# %% [code]
gobp, gomf, gocc = [
    pd.get_dummies(go_collapsed[t].str.split(";").explode().dropna(), sparse=True) for t in namespaces
]

# %% [markdown]
### Join together SDF and GO

# %% [code]
sdf_go = turb.merge([sdf, go_collapsed], verbose=2)
# rename the dataset
sdf_go.name_ = "SDF_GO"

# %% [markdown]
### Minor preprocessing of expression datasets

Aviners' dataset needs to aggregate over the replicates, we ignore the translation data.

# %% [code]
aviner_rna.aggregate_k("mean", 
                        names=["mRNA_HeLa_%dh" % i for i in [0, 2, 4, 6, 8, 12]], 
                        selectors=["_%s_" % i for i in ['0hB','2hR', '4hR', '6hR', '8hR', '12hR']], 
                        keep=False)

aviner_prot.aggregate_k("mean", 
                        names=["prot_HeLa_%s" % i for i in ["G1", "S", "G2M"]], 
                        selectors=["_%s_" % i for i in ["G1", "S", "G2M"]], 
                        keep=False)

aviner_rna.update_meta()
aviner_prot.update_meta()

# %% [code]
aviner_rna_prot = (turb.merge([aviner_rna, aviner_prot], name="Aviner", verbose=2)
                    .apply("drop_duplicates", subset='gene_names'))
aviner_rna_prot.name_ = "Aviner_RNA_Prot"

# %% [markdown]
### Unify naming conventions

Different datasets use different naming conventions to describe mRNA or protein level data.

# %% [code]
aviner_rna_prot.rename_axis([('_2h',"_G1"), ('_8h',"_S"), ('_12h','_G2M')])
lund_all.rename_axis([('RPKM','mRNA'), ('INTENSITY','prot')])
lund_prot.rename_axis([('INTENSITY','prot')])
lund_rna.rename_axis([('RPKM','mRNA')])

# %% [markdown]
### Couple Aviners' combine set with a label from HGNC so GO can bind to it (later...)

# %% [code]
coupled_aviner_rna_prot = turb.merge([hgnc, aviner_rna_prot.copy().apply("drop_duplicates", "gene_names")],verbose=2)
coupled_aviner_rna_prot.name_ = "Aviner_wHGNC"

# %% [markdown]
## <a id='mlmodels'> Import the best ML models of SDF-protein from previous chapter </a>

# %% [code]
cell_line = ('U2OS','A431','U251MG','Daoy','HeLa G1','HeLa S','HeLa G2M')
cell_line_dirty = tuple(f.replace(" ", "_") for f in cell_line)

# %% [code]
path_to_models = "../../_intermediate/sdf_to_prot/"

def associate_data(df, cl):
    res_df = pd.concat(map(pd.DataFrame, df), axis=0, sort=False)
    res_df['model'] = res_df['param_model'].astype(str).str.split("(", expand=True)[0]
    # add cell line
    res_df['Cell_Line'] = cl
    return res_df

# %% [markdown]
#### Importing SDF-protein model results

# %% [code]
fs = list(map(associate_data, turb.read(path_to_models + "SDF_*_prot.pkl"), sorted(cell_line)))

# %% [markdown]
#### Code to select the best sklearn model from this set

# %% [code]
def best_model(expr):
    # sklearn packages
    from sklearn import linear_model, tree, svm, ensemble, neighbors
    packages = (linear_model, tree, svm, ensemble, neighbors)
    _expr = expr.reset_index(drop=True)
    best_model_indx = _expr.loc[_expr['mean_test_score'].idxmax(), :]
    mod_n = best_model_indx["model"]
    # select the model object
    mod = [getattr(obj, mod_n) for obj in packages if hasattr(obj, mod_n)][0]()
    # fit the model with the parameters
    pname, pval = turb.utils.dictsplit(best_model_indx['params'])
    # remove model from name and truncate
    _pname = string_replace(pname[1:], ("model__", ""))
    # fix parameters with set_params
    mod.set_params(**dict(zip(_pname, pval[1:])))
    return mod

# %% [markdown]
## Defining and Computing GO runs

**First off, keep all GO terms that have at least 50 genes associated with them.**

# %% [code]
gocols_red = [
    ds.columns[ds.sum() >= 50] for ds in (gobp, gomf, gocc)
]

# %% [markdown]
### Now collapse such that one row is one gene

This takes a while, hence why we cache the result into a pickle file.

# %% [code]
go_red = (pd.concat(
    turb.utils.umapc("../../data/GO/go_by_gene.pkl", 
                    lambda df, cols: df.loc[:, cols].reset_index().groupby(['index']).sum(),
                    (gobp, gomf, gocc), 
                     gocols_red),
    axis=1, sort=False)
    .fillna(0)
    .astype(np.uint8)
    .drop(list(namespaces), axis=1))

# %% [markdown]
### Pre-defined prep functions

# %% [code]
from turbopanda.pipe import impute_missing

# perform some pre-filtering.
# missing value filter
def mvf(df, thresh=.3):
    return df.drop(df.columns[(1. - (df.count() / df.shape[0])) > thresh], axis=1)

# low variance filter
def lvf(df, thresh=1e-6):
    fcols = df.select_dtypes(float)
    return df.drop(fcols.columns[fcols.var() < thresh], axis=1)

# root mean squared error function
def scoring_rmse(y, yp):
    return np.sqrt(np.mean((y - yp)**2))

# selecting the appropriate columns from data - i.e filtering out text columns (keeping only numeric, bool)
def select_ml_approp_cols(df, pat):
    return absdifference(
        df.columns,
        union(pattern(pat, df.columns), df.select_dtypes(include=['object']).columns)
    ).tolist()

# provide a function to find the number of accepted GO terms
def find_true_number_goterms(df, terms, col="go_all", cutoff=50):
    nterms = pd.Series(map(lambda t: df[col].str.contains(t).sum(), terms), index=terms)
    return nterms[nterms >= cutoff].index.tolist()

# %% [markdown]
#### Caching for merged intermediate steps

# %% [code]
def _cache_merge_sdf(sdf, expr, verbose=0):
    # merge together
    sdf_expr = turb.merge([sdf, expr], verbose=verbose)
    # apply pre-filtering functions
    _sdf_pipe = sdf_expr.copy().apply("pipe", mvf).apply("pipe", lvf).apply("pipe", impute_missing)
    return _sdf_pipe


def _cache_merge_mrna(expr, go, verbose=0):
    # merge together and drop subset
    return turb.merge([expr, go], verbose=verbose)

# %% [markdown]
#### Intermediate SDF-expr step

# %% [code]
def generate_sdf_expr_dataset(sdf, expr, cl, with_xmrna=False, verbose=0):
    """
    - sdf: the sequence derived feature matrix SDF
    - expression: the expr dataset with mRNA/protein
    - cl: the cell line name
    - with_xmrna: whether to include 'mRNA_%s' cell line in input or not
    """
    _subdir = "{}{}__{}.pkl".format(ds_dir, sdf.name_, expr.name_)
    # should save time substantially
    sdf_pipe = cache(_subdir, _cache_merge_sdf, sdf, expr, verbose)
    # determine selected cols
    if with_xmrna:
        xsel = select_ml_approp_cols(sdf_pipe.df_, "prot_ | mRNA_ | HeLa_prot | _?id$") + ["mRNA_%s" % cl]
    else:
        xsel = select_ml_approp_cols(sdf_pipe.df_, "prot_ | mRNA_ | HeLa_prot | _?id$")
    
    # define X and y ml arrays
    X, y = turb.ml.preprocess_continuous_X_y(sdf_pipe, xsel, "prot_%s" % cl)
    return sdf_pipe, xsel, X, y

# %% [markdown]
#### Specify reduced GO terms

# %% [code]
go_terms = go_red.columns.tolist()

# %% [markdown]
# <a id='sdflm'>Computation of the models by Gene Ontology</a>

We have three broad groups of computation:

1. SDF-to-protein GO models
2. mRNA-to-protein GO models
3. SDF-plus-mRNA-to-protein GO models

# %% [code]
from sklearn.linear_model import LinearRegression
from copy import deepcopy

# %% [markdown]
#### 1. SDF-to-protein GO models

# %% [code]
# here in this function, we expect:
def computing_sdf_to_protein_with_go(sdf, expression, model, name, cl, terms):
    """
    In this function we expect:
    
    - sdf: the sequence derived feature matrix SDF
    - expression: the expr dataset with mRNA/protein
    - model: the 'best model' as determined by grid search from official4ml
    - name: a name to use for the temp file(s)
    - cl: the name of the cell line, to derive mRNA and protein data from.
    - terms: a list of GO terms to get results for.
    """
    sdf_pipe, xsel, X, y = generate_sdf_expr_dataset(sdf, expression, cl)
    # select new subset of GO terms using our dataset
    newgos = find_true_number_goterms(sdf_pipe.df_, terms)
    
    # now we define an inner function to calculate our train-test-splits for each GO term
    def _inner_train_test_model(term):
        # local model obj
        mod = deepcopy(model)
        # we find all rows containing term, any NaNs are false entries
        t_rows = sdf_pipe['go_all'].str.contains(term).fillna(False)
        # ensure that there is at least 1 element in t_rows
        # model is ready to be fit - we train on every row that is NOT in the GO term
        mod.fit(X[~t_rows.values], y[~t_rows.values])
        # and score the model based on every row IN the GO term
        return scoring_rmse(y[t_rows.values], mod.predict(X[t_rows.values]))
    
    # now iterate over this list using something like umapc
    res_scores = turb.utils.umappcc(name+".pkl", _inner_train_test_model, newgos)
    
    return res_scores

# %% [markdown]
#### 2. mRNA-to-protein GO models

# %% [code]
def computing_mrna_to_protein_with_go(expr, go, cl, terms):
    # generate mrna and prot strings
    xcol = "mRNA_" + cl
    ycol = "prot_" + cl
    # combine together expr and go
    expr_go = (cache("{}{}__go.pkl".format(ds_dir, expr.name_), _cache_merge_mrna, expr, go)
               .apply("dropna", subset=[xcol, ycol]))
    # select new subset of GO terms using our dataset
    newgos = find_true_number_goterms(expr_go.df_, terms)
    # split into x and y
    X, y = turb.ml.preprocess_continuous_X_y(expr_go, xcol, ycol)
    # get split terms
    def _inner_train_test_model(term):
        mod = LinearRegression()
        # we find all rows containing term, any NaNs are false entries
        t_rows = expr_go['go_all'].str.contains(term).fillna(False)
        # ensure that there is at least 1 element in t_rows
        # model is ready to be fit - we train on every row that is NOT in the GO term
        mod.fit(X[~t_rows.values], y[~t_rows.values])
        # and score the model based on every row IN the GO term
        return scoring_rmse(y[t_rows.values], mod.predict(X[t_rows.values]))
    
    # now iterate over the go term list using umap, making OLS models and fitting them
    res_scores = umap(_inner_train_test_model, newgos)
    return res_scores

# %% [markdown]
#### 3. SDF-plus-mRNA-to-protein GO models

# %% [code]
def computing_sdf_plus_mrna_to_protein_with_go(sdf, expression, model, name, cl, terms):
    # generate our data subsets
    sdf_pipe, xsel, X, y = generate_sdf_expr_dataset(sdf, expression, cl, with_xmrna=True)
    # select new subset of GO terms using our dataset
    newgos = find_true_number_goterms(sdf_pipe.df_, terms)
    # now we define an inner function to calculate our train-test-splits for each GO term
    def _inner_train_test_model(term):
        # local model obj
        mod = deepcopy(model)
        # we find all rows containing term, any NaNs are false entries
        t_rows = sdf_pipe['go_all'].str.contains(term).fillna(False)
        # ensure that there is at least 1 element in t_rows
        # model is ready to be fit - we train on every row that is NOT in the GO term
        mod.fit(X[~t_rows.values], y[~t_rows.values])
        # and score the model based on every row IN the GO term
        return scoring_rmse(y[t_rows.values], mod.predict(X[t_rows.values]))
    
    # now iterate over this list using something like umapc
    res_scores = turb.utils.umappcc(name+".pkl", _inner_train_test_model, newgos)
    
    return res_scores

# %% [markdown]
#### Prep to stack the GO terms, sort the cell line names

# %% [code]
go_stack = [go_terms]*7
cl_sorted = sorted(cell_line_dirty)
bmodels = list(map(best_model, fs))

# %% [markdown]
### Model Composer: Composing the Inputs for each of the three GO model sets

#### What outputs we expect for each GO term:

* The RMSE score for the out-of-sample test, (by cell line), for SDF-to-protein
* The number of samples for the out-of-sample, to ensure > 50 for ML reliability.

#### 1. Scores for composed 1

# %% [code]
cache_dir="../../_intermediate/sdf_to_prot_go/"

eorder1 = [lund_prot] + [vogel_expr] + [aviner_prot]*3 + [lund_prot]*2

igroups1 = retuple(zipe(
    sdf_go, 
    eorder1,
    bmodels,
    patproduct(cache_dir+"SDF_%s_GOprot", cl_sorted),
    cl_sorted,
    go_stack
))

# %% [markdown]
#### 2. Scores for composed 2

# %% [code]
igroups2 = retuple(zipe(
    [lund_all] + [vogel_expr] + [coupled_aviner_rna_prot]*3 + [lund_all]*2,
    go_collapsed,
    cl_sorted,
    go_stack
))

# %% [markdown]
#### 3. Scores for composed 3

# %% [code]
cache_dir="../../_intermediate/sdf_to_prot_go/"

eorder3 = [lund_all] + [vogel_expr] + [aviner_rna_prot]*3 + [lund_all]*2

igroups3 = retuple(zipe(
    sdf_go, 
    eorder3,
    bmodels,
    patproduct(cache_dir+"SDF_%s_GOmrnaprot", cl_sorted),
    cl_sorted,
    go_stack
))

# %% [markdown]
# <a id='result'>Prepare results dataset</a>

We need to 're-extract' the selected GO terms from each of our models such that we have an index for each column.

Mappings:

* sdf-to-protein: cycle_go_run
* mrna-to-protein: cycle_go_run2
* sdf-plus-mrna-to-protein: cycle_go_run3

Stages:

1. Extracting correct GO labels for each of the 300+ values for each cell line
2. Extract sample size for each score (so we know how reliable it is - how many GO terms in test)

#### Prep functions

# %% [code]
def select_go_label_mrna(expr, go, cl, terms):
    # combine together expr and go
    expr_go = (cache("{}{}__go.pkl".format(ds_dir, expr.name_), _cache_merge_mrna, expr, go)
                .apply("dropna", subset=["mRNA_"+cl, "prot_"+cl]))
    # select new subset of GO terms using our dataset
    newgos = find_true_number_goterms(expr_go.df_, terms)
    return newgos

def select_go_label_sdf(sdf, expr, terms):
    # prefilter and impute missing values - this is important so the row filtering 'lines up'
    sdf_pipe = cache("{}{}__{}.pkl".format(ds_dir, sdf.name_, expr.name_), _cache_merge_sdf, sdf, expr)
    # select new subset of GO terms using our dataset
    newgos = find_true_number_goterms(sdf_pipe.df_, terms)
    return newgos

# here in this function, we expect:
def computing_sdf_to_protein_sizes(sdf, expr, cl, terms):
    # load in pipe
    sdf_pipe = cache("{}{}__{}.pkl".format(ds_dir, sdf.name_, expr.name_), _cache_merge_sdf, sdf, expr)
    # select new subset of GO terms using our dataset
    newgos = find_true_number_goterms(sdf_pipe.df_, terms)
    # now we define an inner function to calculate our train-test-splits for each GO term
    def _inner_train_test_size(term):
        return sdf_pipe['go_all'].str.contains(term).sum()
    
    return turb.utils.umap(_inner_train_test_size, newgos)


def computing_mrna_to_protein_sizes(expr, go, cl, terms):
    expr_go = (cache("{}{}__go.pkl".format(ds_dir, expr.name_), _cache_merge_mrna, expr, go)
                .apply("dropna", subset=["mRNA_"+cl, "prot_"+cl]))
    # select new subset of GO terms using our dataset
    newgos = find_true_number_goterms(expr_go.df_, terms)
    # get split terms
    def _inner_train_test_size(term):
        return expr_go['go_all'].str.contains(term).sum()
    
    return turb.utils.umap(_inner_train_test_size, newgos)

# %% [markdown]
### Collating and caching the result function

# %% [code]
@cachedec(filename="../../datasets/GO_analysis/go_results_fig5.csv", return_as="pandas")
def collate_results():
    
    cycle_go_run = list(map(computing_sdf_to_protein_with_go, *igroups1))
    cycle_go_run2 = list(map(computing_mrna_to_protein_with_go, *igroups2))
    cycle_go_run3 = list(map(computing_sdf_plus_mrna_to_protein_with_go, *igroups3))
    
    cycle_go_label1 = list(map(select_go_label_sdf, *retuple(zipe(sdf_go, eorder1, go_stack))))
    cycle_go_label2 = list(map(select_go_label_mrna, *igroups2))
    cycle_go_label3 = list(map(select_go_label_sdf, *retuple(zipe(sdf_go, eorder3, go_stack))))
    
    cycle_go_n1 = list(map(computing_sdf_to_protein_sizes, *retuple(zipe(sdf_go, eorder1, cl_sorted, go_stack))))
    cycle_go_n2 = list(map(computing_mrna_to_protein_sizes, *igroups2))
    cycle_go_n3 = list(map(computing_sdf_to_protein_sizes, *retuple(zipe(sdf_go, eorder3, cl_sorted, go_stack))))
    
    # compile together and cache
    go_results = pd.concat(map(
        lambda go_rmse, go_n, go_lab, dgroup: pd.DataFrame({
            "GO term": join(*go_lab),
            "RMSE": join(*go_rmse),
            "N": join(*go_n),
            "Cell Line": np.repeat(sorted(cell_line), list(map(len, go_lab)))
        }).assign(Input_Source=dgroup),
        (cycle_go_run, cycle_go_run2, cycle_go_run3), 
        (cycle_go_n1, cycle_go_n2, cycle_go_n3),
        (cycle_go_label1, cycle_go_label2, cycle_go_label3),
        ('SDF (alone)', 'mRNA (alone)', 'SDF plus mRNA')
    )).reset_index(drop=True)
    
    return go_results

# %% [markdown]
## Compiling Data together into results

Sincec we have so much data for each GO term, we'll be selecting the K most and least predictable terms to glean some useful information about the predictability of such genes.

# %% [code]
go_results = collate_results()

# %% [markdown]
## <a id='sanity'>Sanity check: Relationship between RMSE and N should be fairly random? </a>

# %% [code]
goterm_grouped = go_results.groupby(['GO term', 'Input_Source'], as_index=False).mean()
goterm_grouped2 = go_results.groupby(['GO term'], as_index=False).mean()
goterm_grouped3 = go_results.groupby(['GO term', 'Cell Line'], as_index=False).mean()

# %% [code]
fig, axes = turb.plot.gridplot(4, ax_size=(6, 4))
turb.plot.scatter(go_results['N'], go_results['RMSE'], c=go_results['Input_Source'],
                  s=1.5, ax=axes[0], title="All terms by source")
turb.plot.scatter(goterm_grouped['N'], goterm_grouped['RMSE'], c=goterm_grouped['Input_Source'],
                  x_scale="log", ax=axes[1], s=5., title="Grouped GO terms by source",
                  fit_line=True)
turb.plot.scatter(goterm_grouped3['N'], goterm_grouped3['RMSE'], c=goterm_grouped3['Cell Line'],
                  x_scale="log", ax=axes[2], s=5., title="Grouped GO terms by Cell Line",
                  fit_line=True, cmap="magma")
turb.plot.scatter(goterm_grouped2['N'], goterm_grouped2['RMSE'],
                  x_scale="log", ax=axes[3], s=3., title="Grouped GO terms",
                  fit_line=True, fit_line_degree=1, c='g')
_ = list(map(lambda ax: ax.set_ylim(.5, 1.7), axes))
fig.tight_layout()
plt.show()

# %% [markdown]
### Highest and lowest GO term RMSE

# %% [code]
def krange(df, col, k):
    return pd.concat([df.nlargest(k, col).assign(Order="Bottom RMSE"),
                      df.nsmallest(k, col).assign(Order="Top RMSE")])

def filter_min_nwords(df, col, n):
    return df[df[col].str.split("[ -]").apply(len) <= n]


go_kbest = krange(goterm_grouped2.pipe(filter_min_nwords, "GO term", 3), "RMSE", 10)

# %% [code]
ax = turb.plot.bar1d(go_kbest['GO term'], go_kbest['RMSE'],
                c=go_kbest['Order'], vert=False, cmap="viridis", legend=True)
ax.set_xlabel("RMSE")
plt.show()

# %% [markdown]
### GO terms with the highest and lowest variance between cell lines and X inputs

# %% [code]
go_var = go_results.groupby(['GO term'], as_index=False)['RMSE'].var()

# %% [code]
go_kbest2 = krange(go_var.pipe(filter_min_nwords, "GO term", 3), "RMSE", 7)
ax = turb.plot.bar1d(go_kbest2['GO term'], go_kbest2['RMSE'],
                c=go_kbest2['Order'], vert=False,
                cmap="viridis", legend=True)
ax.set_xlabel("Var(RMSE)")
ax.set_xscale('log')
plt.show()

# %% [markdown]
### GO terms that 'improve the most' in RMSE from mRNA to SDF to SDF+mRNA

# %% [code]
# cell lines are aggregated over.
go_diff_source = go_results.pivot_table(values="RMSE", columns="Input_Source", index="GO term").reset_index()
# assign an 'improvement column'
go_diff_source['error_loss'] = go_diff_source['SDF plus mRNA'] - go_diff_source['mRNA (alone)']

# %% [markdown]
### GO terms that 'vary mostly' across different cell lines

# %% [code]
dataname_order = ('mRNA (alone)', 'SDF (alone)', 'SDF plus mRNA')

# %% [code]
def plot_figure_test(subterms, ax=None, *args, **kwargs):
    ax3pal = ('#8eba43', '#eb8a44', '#7d4427')
    gosub = go_results[go_results['GO term'].isin(subterms)]
    if ax is None:
        ax = sns.barplot(data=gosub, x="RMSE", y="GO term", hue="Input_Source",
                    order=subterms,
                    hue_order=dataname_order,
                    palette=ax3pal, *args, **kwargs)
    else:
        sns.barplot(data=gosub, x="RMSE", y="GO term", hue="Input_Source",
                    order=subterms,
                    hue_order=dataname_order,
                    palette=ax3pal, ax=ax, *args, **kwargs)
    ax.set_xlim(.4, 1.8)
    ax.get_legend().remove()
    ax.set_xlabel(r"RMSE ($\pm$ Cell Line)", fontsize=14)
    ax.set_ylabel("GO term", fontsize=14)
    ax.grid(axis='x')
    
    # replace names with clean ones?
    
    # optionally return the ax if it started as none

# %% [code]
goterm_piv = goterm_grouped.pivot('GO term', 'Input_Source', 'RMSE').dropna()
goterm_piv['SDF improvement'] = -(goterm_piv['SDF plus mRNA'] - goterm_piv['mRNA (alone)'])

# %% [markdown]
#### Display test figures

# %% [code]
plot_figure_test(goterm_piv['SDF plus mRNA'].nlargest(10).index)

# %% [code]
plot_figure_test(goterm_piv['SDF plus mRNA'].nsmallest(10).index)

# %% [code]
plot_figure_test(goterm_piv['SDF improvement'].nlargest(10).index)

# %% [markdown]
# <a id='offic_fig'>Official Figure</a>

#### Converting greek words into characters

# %% [code]
basic_greek_alphabet = ('alpha', 'beta', 'gamma', 'phi', 'pi', 'zeta', 'rho')
latin_to_greek = list(turb.utils.dict_to_tuple({x:r"$\{}$-".format(x) for x in basic_greek_alphabet}))

# %% [markdown]
#### Other text modifications to GO names

# %% [code]
text_replace_d = list(turb.utils.dict_to_tuple({'growth factor': 'GF', 'nonsense-mediated decay': '(nonsense)', 
                  'transforming ': '', 'from RNA polymerase II promoter': '(RNA poly.II promoter)',
                  ' ?pathway ?': '', 'nuclear-transcribed': 'N-T',
                  'membrane': 'ER', 'cotranslational': 'cotrans.', 'dependent': 'dep.',
                  'elongation': 'elong.'}))

# %% [markdown]
#### GO term selections here

# %% [code]
# figure 2 - TOP GO terms
fig5b = goterm_piv['SDF plus mRNA'].nsmallest(10).index
# figure 3 - BOTTOM GO terms
fig5c = goterm_piv['SDF plus mRNA'].nlargest(10).index
# figure 4 - BEST IMPROVED GO terms
fig5d = goterm_piv['SDF improvement'].nlargest(10).index

# %% [code]
def replace_indices(cols):
    return string_replace(cols, join(text_replace_d, latin_to_greek))

# %% [code]
ax3pal = ('#8eba43', '#eb8a44', '#7d4427')

# %% [code]
import string

def official_figure5():
    fig, ax = turb.plot.gridplot(4, ax_size=(7, 5), arrange='square')
    # figure 1 - all of the GO terms by group
    
    sns.kdeplot(x="N", y="RMSE", hue="Input_Source", ax=ax[0], 
                legend=False, data=goterm_grouped,
                clip=((50, None), (None, None)), 
                hue_order=dataname_order, palette=ax3pal)
    
    # end kdes
    ax[0].set_xlabel(r'$N$ genes (in GO term)', fontsize=14)
    ax[0].set_ylabel(r'RMSE', fontsize=14)
    ax[0].set_xlim(0, 2000)
    ax[0].set_xticks([500, 1000, 1500, 2000])
    ax[0].set_xticklabels(["0.5k", "1k", "1.5k", "2k"])
    ax[0].grid()
    # figure 2 - TOP GO terms 
    plot_figure_test(fig5b, ax=ax[1])
    ax[1].set_yticklabels(replace_indices(fig5b), fontsize=12)
    # figure 3 - BOTTOM GO terms
    plot_figure_test(fig5c, ax=ax[2])
    ax[2].set_yticklabels(replace_indices(fig5c), fontsize=12)
    # figure 4 - BEST IMPROVED GO terms
    plot_figure_test(fig5d, ax=ax[3])
    ax[3].set_yticklabels(replace_indices(fig5d), fontsize=12)
    ax[3].set_ylabel("")
    for i, a in enumerate(ax):
        a.text(-0.05, 1.05, string.ascii_uppercase[i],
               transform=a.transAxes, size=25, weight="bold")
    
    # create the custom legend
    boxes = [mpatches.Patch(color=c, label=l) for l, c in zip(dataname_order, ax3pal)]
    fig.legend(boxes, dataname_order, loc='lower center', 
               borderaxespad=.2, title="ML Input Source(s)",
               ncol=1, bbox_to_anchor=(1.07, .5), prop={"size": 13})
    fig.tight_layout()

    # save the figure
    
    _ = turb.plot.save(fig, 
                plot_type="bar", 
                save_types=('png','pdf','svg', 'eps'),
                name="SDF_GO",
                savemode="update",
                fp="../../Figures/SDF/",
                dpi=1200)
    
    
    plt.show()

# %% [markdown]
# <a id='desc'>Description</a>

### Representation of translation-oriented GO terms predominant in best SDF-based models

    Out-of-sample root mean squared error (RMSE) scores of Gradient-boosted regression trees (GBRT) by Gene Ontology (GO) terms. (A) Bivariate kernel density estimation of the number of genes by RMSE for each GO term model by data input source. Barplots of the 10 (B) lowest, (C) highest and (D) most improved (difference between SDF plus mRNA and mRNA alone) RMSE-scoring GO term models by data input sources. Within GO; Biological Process, Cellular Component and Molecular Function terms are included. See Methods and Supplementary Material for details on model and feature preprocessing/selection.

# %% [code]
official_figure5()

# %% [code]
```
