## Supplementary material for "Predicting Relative Protein Abundance via Sequence-Based Information": S3 Data: SDF-Reduction.ipynb

```
# %% [markdown]
# SDF Reduction Techniques

Here we explore potential methods of reducing the dimensionality of the SDF dataset and to produce more meaningful data.

What is contained within this notebook:

To navigate the notebook, click the links below or scroll:

### Computation

1. **[Initialize and import](#init)**
2. **[Dimensionality Reduction and Feature Selection Technique Summary](#summ)**
3. **[Recomputing selected methods](#selected)**
4. **[Saving outputs to file](#save)**


## Objectives

To explore each of the feature selection and dimensionality reduction techniques, see how much they compress the data, how well they preserve variance, and find a good combination to use pre-machine learning.

# %% [markdown]
### Imports

# %% [code]
import matplotlib.pyplot as plt
import numpy as np
import seaborn as sns
import pandas as pd
%matplotlib inline

import sys
sys.path.insert(0, "../../../turbopanda/")
sys.path.insert(0, "../../scripts/")
import turbopanda as turb
import version

# shortened as *PARAMETERIZE*
from turbopanda import Param as PRMT
from turbopanda import cache
from turbopanda.dev import Bleep

# %% [markdown]
## Version

# %% [code]
version.current()
print("turbopanda: %s" % turb.__version__)
print(version.last_run())

# %% [markdown]
## <a id='init'>Importing the data</a>

# %% [code]
sdf2 = turb.read("../../datasets/SDF_scaled2.json", name="SDF")
sdf2

# %% [code]
sdf2.head(3)

# %% [code]
sdf2.source_

# %% [markdown]
## Handling missing values

- fillna on the PTMC and BCAA columns

# %% [code]
sdf2.transform(pd.DataFrame.fillna, selector="_ptmc$", whole=True, value=0)

# %% [markdown]
## Aggregation by data columns

# %% [code]
group_names = "is_" + sdf2.meta_['data_groups'].cat.categories

# %% [markdown]
#### Split `data_groups` meta into k-boolean selectors

# %% [code]
sdf2.meta_split_category("data_groups")

# %% [markdown]
#### Drop columns with only <1 unique value

# %% [code]
sdf2.view(lambda z: z.nunique()<=1)

# %% [code]
sdf2.drop(lambda z: z.nunique()<=1)

# %% [markdown]
Downcast to avoid any problems with column typing.

# %% [code]
sdf2.downcast()

# %% [markdown]
# <a id='summ'>Table of Feature Selection and Dimensionality Reduction Techniques:</a>

There are many ways to skin a cat, below is a table for comparison, including pros and cons (taken from [here](https://www.analyticsvidhya.com/blog/2018/08/dimensionality-reduction-techniques-python/)):

| Method Name | Technique | Description | Pros | Cons | Target? |
| ------------ | :---------: |-------- | :------------: | :----------------------------------: | :-----------: |
| **Missing Value Filter** | Feature <br> Selection | Drops columns that have missing values above <br> threshold percentage $\phi$ | Simple | Can drop a lot of rows, leading <br> to ill-conditioning | No |
| **Low Variance Filter** | Feature <br> Selection | Drops columns with variance below some <br> threshold $\phi$ | Very intuitive with ML | May have no effect on predictability | No |
| **High Correlation Filter** | Feature <br> Selection |  Drops columns that only correlate <br> very strongly, except columns <br> that correlate to target well | Reduces multi-collinearity, <br> improves model stability | Only works with linear relationships | Yes/No |
| **Random Forest** | Feature <br> Selection |  A complex tree-like algorithm which contains <br> in-built feature selection | Handles non-linear relationships | Models can be too complex, <br> do not generalize well | Yes |
| **Reverse Feature <br> Elimination (RFE)** | Feature <br> Selection |  Iteratively trains a model and drops a feature at <br> each step until model changes are <br> negligible | Thorough process, informs the <br> impact of each feature | 1. Assumes each underlying model fits well <br> 2. Computationally expensive | Yes |
| **Forward Feature <br> Selection** | Feature <br> Selection |  Similar to RFE, trains an additive model <br> using F-score by adding 1 feature <br> at a time | Technique well established <br> in biology, known as ANOVA | 1. Only works with linear relationships <br> 2. Computationally expensive | Yes |
| **Factor Analysis** | Dimensionality <br> Reduction |  Variables are grouped by their correlations, <br> and are known as factors | 1. Factors can directly relate to <br> real-world phenomena <br> 2. Target variable not needed | 1. Number of factors must be known beforehand <br>2. Factors can be difficult to observe | No |
| **Principle Component <br> Analysis (PCA)** | Dimensionality <br> Reduction | Extracts a subset of uncorrelated <br> *principle components* from a large set <br> of variables through SVD | 1. Computationally cheap <br> 2. Provides *explained variance* | Only produces linear combinations | No |
| **Independent Component <br> Analysis (ICA)** | Dimensionality <br> Reduction | Reduces variable set using *information theory* <br> by maximising the kurtosis of projected <br> values | Modelled close to PCA in theory, <br> hence intuitive | 1. Assumes variables are linear mixtures <br> 2. Assumes latent variables are <br> mutually independent | No |
| **Manifold Isomap** | Dimensionality <br> Reduction | Projects many points onto a <br> lower dimensional space by finding <br> manifolds | Handles strongly non-linear <br> relationships | 1. Assumes manifold is continuous <br> and differentiable <br> 2. Very computationally expensive | No |
| **t-Distributed Stochastic <br> Neighbor Embedding <br> (t-SNE)** | Dimensionality <br> Reduction | Uses neighbouring algorithms to <br> provide low-dimensional representation | 1. Retains both local and <br> global data structure <br> 2. Extremely good visualizations | 1. Computationally expensive <br> 2. Large loss of information | No |
| **UMAP** | Dimensionality <br> Reduction | Similar to t-SNE, but helps to preserve <br> more information, and larger datasets | Shorter runtime to t-SNE | "^" | No |
| **Nonnegative Matrix <br> Factorization (NMF)** | Dimensionality <br> Reduction | Factorizes the data matrix <br> to two approximate matrices. | 1. Improved interpretability of models over PCA<br> 2. Provides automatic regularization | Requires all nonnegative elements | No | 

# %% [code]
labels = sdf2.select("~object & ~_id$ & ~_ID$")

# %% [code]
keep_rows = turb.ml.preprocess_continuous_X(sdf2, labels).index

# %% [markdown]
# Missing Value Filter

- $+$ Simple and effective
- $-$ Selection of drop threshold arbitrary

Drops columns that have more than some proportion of missing values.

# %% [code]
mvf = lambda df, thresh=0.3: df.drop(df.columns[(1 - (df.count() / df.shape[0])) > thresh], axis=1)
# mvf(sdf2.df_)
res1 = mvf(sdf2.df_)
res1.shape

# %% [markdown]
# Low variance filter

- $+$ Simple
- $-$ Value of threshold can be arbitrary
- $-$ Variance may not impact on predictability

Drops columns that have less than some value of variance. Only applies to numerical columns.

# %% [code]
def lvf(df, thresh=1e-6):
    fcols = df.select_dtypes(float)
    return df.drop(fcols.columns[fcols.var() < thresh], axis=1)
#lvf(sdf2.df_)
res2 = lvf(sdf2.df_)
res2.shape

# %% [markdown]
# High correlation filter

- $+$ Improves model stability
- $+$ Reduces multicollinearity
- $-$ Selection of threshold arbitrary

# %% [code]
def hcf(df, thresh=.6):
    fcols = df.select_dtypes(float)
    return df.drop(
        fcols.columns[np.mean(np.square(fcols.corr()) - np.eye(fcols.shape[1])) > (thresh**2)]
    , axis=1)
# hcf(sdf2.df_)
res3 = hcf(sdf2.df_)
res3.shape

# %% [markdown]
# Random Forest

- $+$ Non-linear effects captured
- $+$ Reasonably quick
- $-$ Relies on target variable
- $-$ May not generalize well, can overfit

# %% [markdown]
### Incorporate using Vogel's dataset for target

# %% [code]
vogel = turb.merge([sdf2, "../../datasets/vogel_all.csv"])
vogel.downcast()

# %% [code]
# selection for X
vog_x = vogel.select("~object & ~_id$ & ~_ID$ & ~_Daoy")

# %% [markdown]
#### Perform small grid search to check number of trees

Cross-validation is indicated by the error bars for each $N$ given.

# %% [code]
forest_grid = {'RandomForestRegressor': {'n_estimators': [50, 100, 200]}}

# %% [code]
turb.ml.fit.grid(vogel, "prot_Daoy", vog_x, forest_grid, plot=True)

# %% [markdown]
#### Fit a direct model knowing that cross-validation variance is low

# %% [code]
from sklearn.ensemble import RandomForestRegressor
from sklearn.pipeline import Pipeline
from sklearn.feature_selection import SelectFromModel

# %% [code]
_x, _y = turb.ml.preprocess_continuous_X_y(vogel, vog_x, "prot_Daoy")
rf = SelectFromModel(RandomForestRegressor(200, oob_score=True)).fit(_x, _y)

# %% [code]
fi = pd.Series(rf.estimator_.feature_importances_, index=vog_x).sort_values()
groups = sdf2.meta_.loc[fi.index, 'data_groups'].values.tolist()

ax = turb.plot.scatter(np.arange(len(fi)), fi.values, c=groups)
ax.hlines(rf.threshold_, 0, len(fi), linestyle=":")
plt.show()

# %% [markdown]
#### Most and least important features

# %% [code]
mlif = pd.concat([fi[:5], fi[-5:]])

# %% [code]
plt.barh(mlif.index, mlif.values)
plt.xscale("log")
plt.show()

# %% [code]
res4 = rf.transform(_x)
res4.shape

# %% [markdown]
# Recursive Feature Elimination

Using RFE to iteratively remove useless features could be interesting. We use support vector machines as they are relatively cheap and provide reasonably good score.

The issue with this method however is that it relies on the underlying estimator fitting the model well, and producing coherent coefficients.

- $+$ Good coverage of model space
- $-$ Expensive
- $-$ Relies on underlying model being correct
- $-$ Not scalable

# %% [code]
_ = turb.ml.fit.grid(vogel, "prot_Daoy", vog_x, ['SVR'], plot=True)

# %% [code]
from sklearn.feature_selection import RFECV
from sklearn.svm import SVR

# %% [code]
@cache(filename="../../_intermediate/sdf_red_rfe.pkl")
def rfe_method():
    rf_mod = SVR(kernel="linear")
    rfe = RFECV(rf_mod, step=1, min_features_to_select=10, cv=3,
                scoring="neg_root_mean_squared_error", n_jobs=-2)
    rfe.fit(_x, _y)
    
    return rfe

# %% [code]
rfe = rfe_method()

# %% [markdown]
Further to this, we see that RFE seems to actively punish most features for some reason, leaving only 13 significant features...?

# %% [code]
plt.plot(np.arange(10, _x.shape[1]+1, 1, np.int), -rfe.grid_scores_)
plt.xlabel("N")
plt.ylabel("CV score")
plt.show()

# %% [code]
_xc[rfe.support_]

# %% [code]
res5 = rfe.transform(_x)
res5.shape

# %% [markdown]
# Forward Greedy Additive Approach (F-Test)

Here we use a cutoff of $p > 10^{-2}$ or 1\%.

There is a big problem with this technique however:

- $+$ simple, intuitive
- $-$ contains many statistical assumptions (variance homoscedasticity, normally distributed)
- $-$ cannot handle categorical inputs

As seen with the output of this method, it finds all of the categorical variables as non-normal/significant.

# %% [code]
from sklearn.feature_selection import f_regression, f_classif, SelectFpr

# %% [code]
f_test, p_test = f_regression(_x, _y, center=False)

# %% [code]
fgaa = SelectFpr(f_regression, alpha=0.05).fit(_x, _y)

# %% [code]
res6 = fgaa.transform(_x)
res6.shape

# %% [markdown]
# Factor Analysis

- $+$: Able to use whole SDF set
- $+$: Not bound to target variable
- $-$: Slow/expensive

# %% [code]
from sklearn.decomposition import FactorAnalysis
from sklearn.model_selection import cross_val_score
from scipy.optimize import minimize_scalar
from scipy import stats

# %% [code]
sdf_x_select = sdf2.select("~object & ~_id$ & ~_ID$")
_X = turb.ml.preprocess_continuous_X(sdf2, sdf_x_select)

# %% [code]
_X.shape[1]

# %% [markdown]
#### Firstly, determine the best `n_components` using cross-val

# %% [code]
fa = FactorAnalysis(svd_method='randomized')

# %% [markdown]
#### Using minimized scalar to find best initial guess

# %% [code]
def min_cross_val_factor(n):
    return -np.mean(cross_val_score(fa.set_params(n_components=int(n)), _X, cv=3))

# %% [code]
@cache(filename="../../_intermediate/sdf_red_fa.pkl")
def fa_method():
    return minimize_scalar(min_cross_val_factor, bounds=(1., float(_X.shape[1]-1)), tol=1e-1)

# %% [code]
fa_min = fa_method()
fa_min

# %% [markdown]
#### Create a range of search values around the best crude *minimize_scalar*

# %% [code]
@cache(filename="../../_intermediate/sdf_red_fa_scores.pkl")
def fa2_method():
    fa_xrange = np.arange(int(fa_min.x)-10, int(fa_min.x)+10, 1, np.int)
    # fa_scores = [np.mean(cross_val_score(fa.set_params(n_components=n), _x)) for n in fa_xrange]
    fa_scores = turb.utils.lparallel(min_cross_val_factor, fa_xrange.tolist())
    return fa_xrange, fa_scores

# %% [code]
fa_xrange, fa_scores = fa2_method()

# %% [code]
plt.plot(fa_xrange, fa_scores, 'x-')
fa_best_n = fa_xrange[np.asarray(fa_scores).argmin()]
plt.vlines(fa_best_n, min(fa_scores), max(fa_scores), linestyle=":")
plt.xlabel("N")
plt.ylabel("CV Scores")
plt.show()

# %% [markdown]
#### Fit FA Model

# %% [code]
fa.set_params(n_components=fa_best_n).fit(_X)

# %% [code]
fa.components_.shape[0]

# %% [markdown]
#### Results of fitted FA model

# %% [code]
res7 = pd.DataFrame(fa.transform(_X), columns=turb.str.patproduct("Factor%d", range(1, fa.n_components+1)))
res7.shape

# %% [code]
factor_matrix = (pd.DataFrame(fa.components_, 
                             columns=sdf_x_select, 
                             index=turb.str.patproduct("Factor%d", range(1,fa.n_components+1))).T
                 .assign(Data_groups=sdf2.meta_.loc[sdf_x_select,"data_groups"]))

# %% [markdown]
#### The effect of each 'data group' in the first Factor

# %% [code]
_fm_groups_bar = factor_matrix.groupby("Data_groups")['Factor1'].mean()
turb.plot.bar1d(_fm_groups_bar, c=turb.plot._palette.palette_cmap(13, "Blues"), vert=False)

# %% [markdown]
#### Distribution of components for the first two factors

# %% [code]
turb.plot.scatter(factor_matrix["Factor1"].values, 
                factor_matrix["Factor2"].values,
                sdf2.meta_.loc[sdf_x_select,"data_groups"].tolist(),
                legend_outside=True, marker='8',
                x_label="Factor 1", y_label="Factor 2",
                s=30, alpha=.7)
plt.show()

# %% [markdown]
# PCA

## Strategy 1: Global PCA on dataset

# %% [code]

strat1 = turb.ml.fit.pca(sdf2, labels, plot=True)
# generate datasubset

# %% [code]
from sklearn.preprocessing import scale
from sklearn.decomposition import PCA

# %% [code]
s1_xt = pd.DataFrame(
    PCA(np.where(np.cumsum(strat1.explained_variance_ratio_) > .9)[0][0] + 1).fit_transform(turb.ml.preprocess_continuous_X(sdf2, labels))
)

# %% [markdown]
## Strategy 2: PCA on continous and discrete features

# %% [code]
_select_float = "float & ~object & ~_id$ & ~_ID$"
_select_uints = "uint8 & ~object & ~_id$ & ~_ID$"

# %% [code]
strat2 = turb.ml.fit.pca(sdf2, PRMT(sdf2.select(_select_float),sdf2.select(_select_uints)), 
                    plot=True, refit=True, with_transform=True, plot_kwargs=dict(ax_size=3))

# %% [code]
s2_models, s2_xt = turb.ml.fit.stratified_pca(sdf2, {"is_float":_select_float,"is_uint":_select_uints})

# %% [markdown]
## Strategy 3: PCA on each `data_group` found in meta attribution.

# %% [markdown]
## Group 1: Mononucleotide frequencies

We will check which aggregation method seems to work the best.

#### Methods:

* PCA

Split by:

- region (mRNA, cds, 5'-utr)..
- T, A, C, G


# %% [code]
_rna_sources = ('mrna', 'cds', 'utr5', 'utr3')
_mononuc_examples = PRMT(*["^[AGCT]_%s" % i for i in _rna_sources] + ['mononuc_freq'])

# %% [code]
_ = turb.ml.fit.pca(sdf2, _mononuc_examples, plot=True, plot_kwargs=dict(ax_size=3))

# %% [markdown]
#### Differences in Individual Bases

# %% [code]
_indiv_bases = PRMT(*["^%s_(?!aa)" % i for i in ['T','C','G','A']] + ["mononuc_freq"])

# %% [code]
_ = turb.ml.fit.pca(sdf2, _indiv_bases, plot=True, plot_kwargs=dict(ax_size=3))

# %% [markdown]
## Group 2: Dinucleotide frequencies

Split into:

- by region (mrna, cds, 5'-utr)
- by AT-rich/GC-rich or mixed-base pairings

# %% [code]
_dinuc_bases = PRMT(*["[ATGC]{2}_%s" % i for i in _rna_sources])

# %% [code]
_ = turb.ml.fit.pca(sdf2, _dinuc_bases, plot=True, plot_kwargs=dict(ax_size=3))

# %% [code]
_ = turb.ml.fit.pca(sdf2, PRMT("exc_gc","exc_at","base_mix","dinuc_freq")
                    , plot=True, plot_kwargs=dict(ax_size=3))

# %% [markdown]
## Group 3: Length proportion

Very compressable.

# %% [code]
_ = turb.ml.fit.pca(sdf2, 'l_prop', plot=True, plot_kwargs=dict(ax_size=3))

# %% [markdown]
## Group 4: Gene length

Not particularly compressable...?

# %% [code]
_ = turb.ml.fit.pca(sdf2, 'length_(?!prop)', plot=True,
                    plot_kwargs=dict(ax_size=3))

# %% [markdown]
## Group 5: Free energy

Issues as the group displays heterogeneous distributions as the window size increases.

# %% [code]
_ = turb.ml.fit.pca(sdf2, 'free_energy', plot=True,
                    plot_kwargs=dict(ax_size=3))

# %% [markdown]
## Group 6: RNA misc features

PCA seems to struggle with this group due to nonlinearity: maybe let's try NMF?

# %% [code]
_ = turb.ml.fit.pca(sdf2, 'rna_misc', plot=True,
                    plot_kwargs=dict(ax_size=3, n_samples_annotate=3))

# %% [markdown]
## Group 7: Codon biases

# %% [code]
_ = turb.ml.fit.pca(sdf2, 'codon_bias', plot=True,
                    plot_kwargs=dict(ax_size=3, n_samples_annotate=3))

# %% [markdown]
## Group 8: Amino acid frequency

# %% [code]
_ = turb.ml.fit.pca(sdf2, 'aa_freq', plot=True,
                    plot_kwargs=dict(ax_size=3, n_samples_annotate=3))

# %% [markdown]
## Group 9: Amino acid meta features

# %% [code]
_ = turb.ml.fit.pca(sdf2, 'aminoacid_meta', plot=True,
                    plot_kwargs=dict(ax_size=3, n_samples_annotate=3))

# %% [markdown]
## Group 10: Post-translational modifications

# %% [code]
_ = turb.ml.fit.pca(sdf2, 'ptm', plot=True,
                    plot_kwargs=dict(ax_size=3, n_samples_annotate=3))

# %% [markdown]
## Group 11: Biophysical properties

# %% [code]
_ = turb.ml.fit.pca(sdf2, 'biophysical', plot=True,
                    plot_kwargs=dict(ax_size=3, n_samples_annotate=3))

# %% [code]
# strat3 = turb.ml.fit.pca(sdf2, PRMT(*sdf2.meta_['data_groups'].cat.categories[1:]), refit=True)
strat3_models, s3_xt = turb.ml.fit.stratified_pca(sdf2, sdf2.meta_['data_groups'].cat.categories[1:].tolist())

# %% [markdown]
## Strategy 4: Grouping on Central Dogma Order

# %% [code]
# strat3 = turb.ml.fit.pca(sdf2, PRMT(*sdf2.meta_['data_groups'].cat.categories[1:]), refit=True)
strat4_models, s4_xt = turb.ml.fit.stratified_pca(sdf2, sdf2.meta_['dogma_order'].cat.categories.tolist())

# %% [markdown]
## Strategy 5: `PCA` on data sources

# %% [code]
strat5_models, s5_xt = turb.ml.fit.stratified_pca(sdf2, ['refseq', 'uniprot', 'ptm_source'])

# %% [markdown]
# ICA (Independent Component Analysis)

- $-$ Failure to converge properly with data

# %% [code]
from sklearn.decomposition import FastICA

# %% [code]
ica = FastICA(max_iter=500, tol=1e-3).fit(_X)

# %% [code]
_X.shape

# %% [code]
ica_component_matrix = (pd.DataFrame(ica.components_, 
                             columns=sdf_x_select, 
                             index=sdf_x_select))

# %% [code]
mag_component = pd.Series(np.linalg.norm(ica.components_[:, :2], axis=1), index=sdf_x_select, name="magnitude")

# %% [code]
fig, ax = plt.subplots()
turb.plot.scatter(ica.components_[:, 0], 
                  ica.components_[:, 1], 
                  c=mag_component,
                  ax=ax,
                  x_label="IC1", y_label="IC2")
turb.plot.annotate(ica.components_[:, 0], 
                ica.components_[:, 1],
                sdf_x_select,
                np.where(mag_component > 0.15)[0],
                ax=ax)
plt.show()

# %% [code]
res9 = ica.transform(_X)

# %% [code]
res9.shape

# %% [markdown]
# t-SNE

- $+$ Reasonably fast computation
- $+$ Performs a non-linear embedding
- $+$ Easy to visualize
- $-$ Arbitrary selection of $n$
- $-$ Substantial loss of interpretability

Since there is no clear and objective way of selecting n components, we're not going to use this technique

# %% [code]
from sklearn.manifold import TSNE

# %% [code]
tsne = TSNE(2, n_iter=300, init="pca", n_jobs=-2).fit(_X)

# %% [code]
turb.plot.scatter(tsne.embedding_[:, 0], tsne.embedding_[:, 1], 
                  marker='x', alpha=.5, dense=False,
                  x_label="embedding 1", y_label="embedding 2")
plt.show()

# %% [markdown]
# Comparison of Methods

### A: Reduction in N size

# %% [code]
result_sets = (res1, res2, res3, res4, res5, res6, res7, s3_xt, res9)
names_of_techniques = ('Missing value filter', 'Low variance filter', 'High correlation filter', 
                       'Random Forest', 'RFE', 'ANOVA', 'Factor Analysis', 'PCA', 'ICA')
method_type = ['Feature Selection']*6 + ['Dimensionality Reduction']*3
# shapes

__data = pd.Series(map(lambda x: x.shape[1], result_sets), 
                   index=names_of_techniques, name="N components remaining")
ax = turb.plot.bar1d(__data, c=method_type, vert=False, annotate=True, legend=True)
plt.show()

# %% [markdown]
### Removal of Correlation between Reduced Subsets of PCA

# %% [code]
fig, ax = plt.subplots(ncols=5, figsize=(20, 4))
for p, dataset in zip(ax, (s1_xt, s2_xt, s3_xt, s4_xt, s5_xt)):
    sns.heatmap(dataset.corr(), cmap="seismic", center=0, ax=p)
fig.tight_layout()

# %% [markdown]
# <a id='selected'>Recomputing chosen methods</a>

We choose the following methods:

* **FULL PCA**
* **PCA by Data groups**
* **FACTOR ANALYSIS**

The pre-dimensionality reduction pipeline includes:

1. Missing value filter (70\%)
2. Low variance filter (<$10^{-6}$)

We save into `datasets/pca_reduced` subfolder.

- The model (as a pkl) using `joblib`
- The $\hat{X}$ reduced matrix

# %% [code]
chosen_method_names = ('fa', 'pca_whole', 'pca_datagroup')
fpa = "../../datasets/pca_reduced/"

# %% [markdown]
### Preprocess dataset with mvf, lvf

# %% [code]
labels = sdf2.select("~object & ~_id$ & ~_ID$")
sdf_preml = (sdf2[labels]#
               .pipe(turb.ml.missing_value_filter, threshold=.7)
               .pipe(turb.ml.low_variance_filter, threshold=1e-6)
               .pipe(turb.ml.preprocess_continuous_X))

# %% [markdown]
#### Features dropped using mvf and lvf (compared to before)

# %% [code]
turb.utils.difference(
    turb.ml.preprocess_continuous_X(sdf2.df_, labels).columns,
    sdf_preml.columns
)

# %% [markdown]
### 1. Whole PCA

# %% [code]
from sklearn.decomposition import PCA

def get_best_pca_n(model, threshold=.9):
    return np.where(np.cumsum(model.explained_variance_ratio_) > threshold)[0][0] + 1

# %% [code]
pca_n = get_best_pca_n(strat1)
pca_n

# %% [code]
Xt_1 = pd.DataFrame(
    PCA(pca_n).fit_transform(sdf_preml.values),
    columns=turb.str.patproduct("PC%d", range(1,pca_n+1)),
    index=keep_rows
)

# %% [markdown]
### 2. PCA by data group

Essentially a group of PCAs joined together.

# %% [code]
from turbopanda.utils import intersect as reduce_to

# %% [code]
# fetches the column groups from each 'data group' tag
column_groups = list(map(lambda x: reduce_to(sdf2.columns[sdf2.meta_['data_groups'] == x], sdf_preml.columns),
                    sdf2.mapper_['data_groups'][1:]))
# fit a PCA model to each group
prefitmodels_2 = [PCA(len(g)).fit(sdf_preml[g]) for g in column_groups]
# get the 'reduced 90% n' for each subgroup
pca_group_n = list(map(get_best_pca_n, prefitmodels_2))
# now fit-transform using a new PCA model to get Xt
Xt_2_parts = [PCA(n).fit_transform(sdf_preml[g]) for n, g in zip(pca_group_n, column_groups)]
# define xt_column names as PC parts
Xt_cols = [turb.str.patproduct("{}_PC%d".format(group), range(1,n+1)) for n, group in zip(pca_group_n, sdf2.mapper_['data_groups'][1:])]
# concatenate together and panderise
Xt_2 = pd.DataFrame(np.concatenate(Xt_2_parts, axis=1), columns=turb.utils.join(*Xt_cols), index=keep_rows)

# %% [markdown]
### 3. Factor Analysis

use `fa_best_n` as calculated above to do Factor Analysis again

# %% [code]
from sklearn.decomposition import FactorAnalysis

# %% [code]
Xt_3 = pd.DataFrame(
    FactorAnalysis(fa_best_n).fit_transform(sdf_preml.values),
    columns=turb.str.patproduct("Factor%d", range(1,fa_best_n+1)),
    index=keep_rows
)

# %% [markdown]
### Tack on labelled data

# %% [code]
id_columns = ('id', 'symbol', 'ensembl_id', 'refseq_id', 'uniprot_id')

# %% [code]
id_metadata = sdf2.df_.loc[keep_rows, id_columns]
id_metadata.head()

# %% [markdown]
## <a id='save'>Saving chosen methods to file</a>

# %% [code]
foutpath = "../../datasets/pca_reduced/"

# %% [code]
pd.concat([id_metadata, Xt_1], axis=1, sort=False).to_csv(foutpath+"whole_pca.csv")
pd.concat([id_metadata, Xt_2], axis=1, sort=False).to_csv(foutpath+"pca_data_group.csv")
pd.concat([id_metadata, Xt_3], axis=1, sort=False).to_csv(foutpath+"factor_analysis.csv")
```
