## Supplementary material for "Predicting Relative Protein Abundance via Sequence-Based Information": S3 Data: SDF-RNA-Prot-Relationships.ipynb

```
# %% [markdown]
# Relationships between mRNA and protein

What is contained within this notebook:

To navigate the notebook, click the links below or scroll:

1. **[Initialize and import](#init)**
2. **[Relationship Plots](#ml)**


# %% [code]
import matplotlib.pyplot as plt
import numpy as np
import seaborn as sns
import pandas as pd
import string
%matplotlib inline

import sys
sys.path.insert(0, "../../scripts/")
sys.path.insert(0, "../../../turbopanda/")
# scripts
import version
import myfig as mf
# turbopanda
import turbopanda as turb

# %% [markdown]
## Version

# %% [code]
version.current()
print("turbopanda: %s" % turb.__version__)

# %% [markdown]
## <a id='init'> Import SDF </a>

# %% [code]
sdf = turb.read("../../datasets/SDF_scaled2.json", 
                name="SDF")
sdf

# %% [code]
sdf.info()

# %% [markdown]
## Loading RNA/Protein Expression Datasets

1. Aviner (2015), HeLa cell line, time-series dataset, includes translation
2. Vogel (2010), Daoy cell line
3. Lundberg (2010), U2OS, U251MG and A431 cell lines

Each of these datasets undergoes three main stages: *Load, Merge and Correlate*.

# %% [code]
# read in files
aviner_rna, aviner_prot, aviner_tr = turb.read("../../datasets/clean/*_aviner_adv.csv", 
                    name=("mRNA_aviner", "prot_aviner", "tr_aviner"), index_col=0)
lund_all, lund_prot, lund_rna = turb.read("../../datasets/lundberg_*.csv")
vogel = turb.read("../../datasets/vogel_all.csv")

# join mRNA, protein and drop duplicates on the gene names.
av_rna_prot = turb.merge([aviner_rna, aviner_prot])
av_rna_prot.apply("drop_duplicates", subset=['gene_names'])

# %% [markdown]
### Aggregating replicates

Aviner's datasets have replicates that we average over.

# %% [code]
av_rna_prot.aggregate_k("mean", 
                        names=["HeLa_mRNA_%dh" % i for i in [0, 2, 4, 6, 8, 12]], 
                        selectors=["_%s_" % i for i in ['0hB','2hR', '4hR', '6hR', '8hR', '12hR']], 
                        keep=False)

av_rna_prot.aggregate_k("mean", 
                        names=["HeLa_prot_%s" % i for i in ["G1", "S", "G2M"]], 
                        selectors=["_%s_" % i for i in ["G1", "S", "G2M"]], 
                        keep=False)

av_rna_prot.update_meta()

# %% [markdown]
## <a id='ml'> Machine Learning Models </a>

There are a number of approaches to developing an effective machine learning model. Many of them include proper analysis of the data residuals and weights. We will begin with a simple Linear Regression model, from `sklearn`.

# %% [code]
import sklearn
from sklearn.linear_model import LinearRegression
from sklearn.model_selection import cross_validate, cross_val_predict, RepeatedKFold, GridSearchCV

# %% [code]
import statsmodels.api as sm
import statsmodels.formula.api as smf
import statsmodels.stats.api as sms
from patsy import dmatrices

# %% [markdown]
## mRNA vs protein relationships.

# %% [code]
sklearn.__version__

# %% [markdown]
### Practice Daoy Cell Line

# %% [code]
cv, yp = turb.ml.fit.basic(vogel, "mRNA_Daoy", "prot_Daoy", plot=True)

# %% [code]
turb.plot.scatter_grid(vogel, "mRNA_", "prot_Daoy")

# %% [code]
turb.plot.histogram(vogel['prot_Daoy'])

# %% [code]
turb.plot.histogram(vogel['mRNA_Daoy'])

# %% [code]
# y, X = dmatrices("prot_Daoy ~ mRNA_Daoy", data=vogel.df_, return_type="dataframe")
lm_daoy = smf.ols("prot_Daoy ~ mRNA_Daoy", data=vogel.df_.set_index("EnsGeneID")).fit()
lm_daoy.summary()

# %% [markdown]
### Practice Lundberg Lines

# %% [code]
cv, yp = turb.ml.fit.basic(lund_all, 'RPKM_U2OS', 'INTENSITY_U2OS', plot=True)

# %% [code]
turb.plot.scatter_grid(lund_all, "RPKM_", "INTENSITY_", arrange="column")

# %% [code]
turb.plot.hist_grid?

# %% [code]
y, X = dmatrices("INTENSITY_U2OS ~ RPKM_U2OS", data=lund_all.df_, return_type="dataframe")
lm = sm.OLS(y, X).fit()
lm.summary()

# %% [code]
cv, yp = turb.ml.fit.basic(lund_all, 'RPKM_U251MG', 'INTENSITY_U251MG', plot=True)

# %% [code]
y, X = dmatrices("INTENSITY_U251MG ~ RPKM_U251MG", data=lund_all.df_, return_type="dataframe")
lm = sm.OLS(y, X).fit()
lm.summary()

# %% [code]
cv, yp = turb.ml.fit.basic(lund_all, 'RPKM_A431', 'INTENSITY_A431', plot=True)

# %% [code]
y, X = dmatrices("INTENSITY_A431 ~ RPKM_A431", data=lund_all.df_, return_type="dataframe")
lm = sm.OLS(y, X).fit()
lm.summary()

# %% [markdown]
### Practice HeLa Lines

# %% [code]
turb.plot.scatter_grid(av_rna_prot, "HeLa_mRNA_", "HeLa_prot_")

# %% [code]
cv, yp = turb.ml.fit.basic(av_rna_prot, "HeLa_mRNA_2h", "HeLa_prot_G1", plot=True)

# %% [code]
cv, yp = turb.ml.fit.basic(av_rna_prot, "HeLa_mRNA_8h", "HeLa_prot_S", plot=True)

# %% [code]
cv, yp = turb.ml.fit.basic(av_rna_prot, "HeLa_mRNA_12h", "HeLa_prot_G2M", plot=True)
```
