## Supplementary material for "Predicting Relative Protein Abundance via Sequence-Based Information": S3 Data: SDF-Sanity.ipynb

```
# %% [markdown]
# Data Sanity Checks for SDF

This notebook is for performing sanity/clean checks on the `SDF` dataset we have.

1. [Normalization Strategies](#norm)
2. [Sanity Plots](#sane)

# %% [code]
import matplotlib.pyplot as plt
import numpy as np
import seaborn as sns
import pandas as pd
import seaborn as sns
%matplotlib inline

import sys
sys.path.insert(0, "../../../turbopanda/")
import turbopanda as turb

# %% [code]
sdf = turb.read("../../datasets/SDF_unscaled.json", name="SDF")
sdf

# %% [markdown]
## <a id='norm'> Normalization Strategies </a>

| **Feature** | **Normalization 1** | **Normalization 2** |
| ------------- | --------------------------- | --------------------- |
| *Free energy* | $-\frac{x}{d}$ | $-\frac{x}{d}$ |
| *Dinucleotide count* | $\frac{x} {0.5L_{r}}$ | See equation 2 |
| *Mononucleotide count* |$\frac{x}{L_{r}}$ | See equation 2 |
| *Amino acid count* | $\frac{x} {L_{aa}}$ | See equation 2 |
| *Isoelectric point* | $x - 7$ | $x-7$ |
| *Instability Index* | $x-40$ | $x-40$ |
| *kozak* | $\frac{x}{L_{cds}}$ | $\frac{x}{L_{cds}}$ |
| *mRNA feature counts* | mRNA length | mRNA length |
| *PTM* | AA length | AA length |
| *Length* | $\log(x+1)$ | $\log(x+1)$ |

# %% [markdown]
### Scaler 1:

We normalize the features by their **respective length**, for example, MFE features are normalized by sliding window, base counts normalized by the length of the gene, etc.

$$
\begin{equation}
\hat{f}_i=\frac{f_i}{L_i}
\label{eq:sc1}
\tag{1}
\end{equation}
$$

# %% [code]
sdf_scaled1 = sdf.copy().transform_k([
        # free energy window is - x / window size
        *[(lambda x: -(x / d), "MFE_win%d" % d) for d in [10,20,30,40,60,80]],
        # feature counts, mononucleotide - x / L
        (lambda x: x / sdf.df_["length_mrna"], ("mono_mrna","feat_count")),
        # dinucleotide count -> x / 0.5L
        (lambda x: x / (sdf.df_["length_mrna"] / 2.), "di_mrna"),
        # mononuc CDS, kozak -> x / L
        (lambda x: x / sdf.df_["length_cds"], ("mono_cds","kozak")),
        # dinuc CDS -> x / 0.5L
        (lambda x: x / (sdf.df_["length_cds"] / 2.), "di_cds"),
        # mononuc 5' -> x / L
        (lambda x: x / sdf.df_["length_utr5"], "mono_utr5"),
        # dinuc 5' -> x / 0.5L
        (lambda x: x / (sdf.df_["length_utr5"] / 2.), "di_utr5"),
        # mononuc 3' -> x / L
        (lambda x: x / sdf.df_["length_utr3"], "mono_utr3"),
        # dinuc 3' -> x / 0.5L
        (lambda x: x / (sdf.df_["length_utr3"] / 2.), "di_utr3"),
        # MFE single, -> -(x / L)
        (lambda x: -(x / sdf.df_["length_utr5"]), "MFE"),
        # methionine f -> x / 1/3 L
        (lambda x: x / (sdf.df_["length_utr5"] / 3), "M_utr5"),
        # aa freq, ptm count -> x / L_aa
        (lambda x: x / sdf.df_["length_aa"], ("aa_freq","ptm_source")),
        # instability -> center on zero -> x - 40
        (lambda x: x - 40, "Instability"),
        # isoelectric -> center on zero -> x - 7
        (lambda x: x - 7, "Isoelectric"),
        # length -> log2(x+1) scaling
        (lambda x: np.log2(x+1), "length"),
        ])

# %% [markdown]
For scaling 2, remember we must calculate the codon frequency tables:

# %% [code]
codon_chart = {'phe': ['uuu', 'uuc'], 'leu': ['uua', 'uug', 'cuu', 'cuc', 'cua', 'cug'],
               'ile': ['auu', 'auc', 'aua'], 'val': ['guu', 'guc', 'gua', 'gug'],
               'met': ['aug'], 'ser': ['ucu', 'ucc', 'uca', 'ucg', 'agu', 'agc'],
               'pro': ['ccu', 'ccc', 'cca', 'ccg'], 'thr': ['acu', 'acc', 'aca', 'acg'], 
               'ala': ['gcu', 'gcc', 'gca', 'gcg'], 'tyr': ['uau', 'uac'], 
               'stop': ['uaa', 'uag', 'uga'], 'his': ['cau', 'cac'],
               'gln': ['caa', 'cag'], 'asn': ['aau', 'aac'], 'lys': ['aaa', 'aag'],
               'asp': ['gau', 'gac'], 'glu': ['gaa', 'gag'], 'cys': ['ugu',' ugc'],
               'trp': ['ugg'], 'arg': ['cgu', 'cgc', 'cga', 'cgg', 'aga', 'agg'],
               'gly': ['ggu', 'ggc', 'gga', 'ggg']
               }

aa_abbr = {'A': 'ala', 'C': 'cys', 'D': 'asp', 'E': 'glu', 'F': 'phe', 'G': 'gly', 'H': 'his', 
           'I': 'ile', 'K': 'lys', 'L': 'leu', 'M': 'met', 'N': 'asn', 'P': 'pro', 'Q': 'gln',
           'R': 'arg', 'S': 'ser', 'T': 'thr', 'V': 'val', 'W': 'trp', 'Y': 'tyr'}

n_synonomous = pd.Series({a +"_aa": len(codon_chart[b]) for a, b in aa_abbr.items()})

expect_synon = n_synonomous/64

# %% [code]
sdf_scaled2 = sdf.copy().transform_k([
        *[(lambda x: -(x / d), "MFE_win%d" % d) for d in [10,20,30,40,60,80]],
        (lambda x: (x / (0.25*sdf.df_["length_mrna"])) - 1, "mono_mrna"),
        (lambda x: (x / (0.0625*0.5*sdf.df_["length_mrna"])) - 1, "di_mrna"),
        (lambda x: (x / (0.25*sdf.df_["length_cds"])) - 1, "mono_cds"),
        (lambda x: (x / (0.0625*0.5*sdf.df_["length_cds"])) - 1, "di_cds"),
        (lambda x: (x / (0.25*sdf.df_["length_utr5"])) - 1, "mono_utr5"),
        (lambda x: (x / (0.0625*0.5*sdf.df_["length_utr5"])) - 1, "di_utr5"),
        (lambda x: (x / (0.25*sdf.df_["length_utr3"])) - 1, "mono_utr3"),
        (lambda x: (x / (0.0625*0.5*sdf.df_["length_utr3"])) - 1, "di_utr3"),
        (lambda x: x / sdf.df_["length_cds"], "kozak"),
        (lambda x: x / sdf.df_["length_mrna"], "feat_count"),
        (lambda x: -(x / sdf.df_["length_utr5"]), "MFE"),
        (lambda x: (x / ((1/64)*(1/3)*sdf.df_["length_utr5"])) - 1, "M_utr5"),
        (lambda x: (x / (expect_synon[x.name]*sdf.df_["length_aa"])) - 1, "aa_freq"),
        (lambda x: x - 40, "Instability"),
        (lambda x: x - 7, "Isoelectric"),
        (lambda x: x / sdf.df_["length_aa"], "ptm_source"),
        (lambda x: np.log2(x+1), "length"),
        ]
)

# %% [markdown]
## <a id='sane'> Sanity Plots </a>

# %% [markdown]
### Gene Length

#### Unnormalized

For unmodified length, we expect it to follow an *expontential distribution* given that it's a count of DNA characters.

Here we see that CDS and amino acid length don't cleanly fit the *exp* distribution as well as the others for some reason. We suspect the length of tail
plays a part in the success of the fit.

#### Normalized

For log-normalized properties; we expect to follow a normal or gamma-like distribution instead.

We see that *norm* distributions fit mRNA, CDS and amino acid here when adjusted, but 5'UTR and 3'UTR have a left-skew bias, which we model using the `gumbel_l` distribution.

# %% [code]
from scipy import stats

# %% [code]
turb.plot.hist_grid(sdf, "length", shared_dist=["expon", 'gamma'])

# %% [code]
turb.plot.hist_grid(sdf_scaled2, "length", shared_dist=["norm", 'gumbel_l', 'gumbel_r'])

# %% [markdown]
### mRNA feature counts

#### Unnormalized

For unmodified mRNA feature counts, we would broadly expect them all to follow an *exponential distribution*, unfortunately there are many categories with extreme outliers which limits the ability to model the distribution effectively.

#### Normalized

When normalized against the length of the mRNA, they tend to perform much better. However `transit_peptide`, `mat_peptide` seem to have such a low frequency that statistically analysing them might simply be pointless - they may instead act as features useful for the *outlier analysis* part of the study.

# %% [code]
turb.plot.hist_grid(sdf, "_fc$", shared_dist=['expon','chi2'], kde_range=1e-2)

# %% [code]
turb.plot.hist_grid(sdf_scaled2, "_fc$", kde_range=1e-2, shared_dist=['expon','norm','gumbel_r','gumbel_l', 't'])

# %% [markdown]
### Codon Bias

The relative frequency of certain codons over others.

#### Unnormalized

For this case, *kozak sequence occurrence* is binary decision, with relatively high frequency, but uORF frequency follows a good *poisson distribution* which is quite interesting. The others broadly follow exponential or normal distributions, but corrected RCBS has a very long tail.

#### Normalized

Corrected kozak and uORF features now follow *chi-squared/gamma* based- distributions more cleanly.


# %% [code]
turb.plot.hist_grid(sdf, "CAI|tAI|RCB|uORF|kozak", "column", 
                    shared_dist=['bernoulli', 'poisson','cauchy', 
                                 'norm', 'gumbel_l', 'gumbel_r'])

# %% [code]
turb.plot.hist_grid(sdf_scaled2, "CAI|tAI|RCB|uORF|kozak", "column", shared_dist=['chi2', 'norm', 'gumbel_r', 'gumbel_l', 'gamma'])

# %% [markdown]
### Mononucleotide frequency

The count of single-base nucleotides in mRNA base features.

#### Unnormalized (Difficulties with CDS-region)

In this case A/T (cds) give $r$-scores of less than 75, indicating very poor fits. In addition G/C (cds) give scores 75-90. These distributions in the CDS range appear to have a large positive skew which needs to be normalized for. The other data sources appear to fit very well within an *exponential distribution*.

#### Normalized

- Strategy 1 includes dividing the base count by the respective gene length, eliminating the length bias.
- Strategy 2 includes scaling the division by the *expected proportion* of that base. We assume that A/C/G/T are uniformly distributed across a sequence so we do $\frac{1}{4}L$ where $L$ is the gene length. In this case values represent a ratio between observed and expected, with $0$ represents no deviation from *expected value*.

It looks as though mRNA samples for scaled1 could belong to a bimodal distribution with two distinct modes. 5'UTR and 3'UTR samples in general seem to have cleaner fits with regards to distribution. The T-distribution is a popular choice for them.

# %% [code]
turb.plot.hist_grid(sdf, "mononuc_freq", "column", 
                    shared_dist=['expon','chi2','poisson'])

# %% [code]
turb.plot.hist_grid(sdf_scaled1, "mononuc_freq", "column", 
                    shared_dist=['norm', 'gumbel_r', 'gumbel_l', 't', 'cauchy'])

# %% [code]
turb.plot.hist_grid(sdf_scaled2, "mononuc_freq", "column", 
                    shared_dist=['norm', 'gumbel_r', 'gumbel_l', 't', 'cauchy'])

# %% [markdown]
#### Seaborn plot of paired G/C-A/T mRNA

Here we see that CDS region has the least overlap between CG/AT pairs, which is not what we'd expect as we would expect these distributions to mirror each other.

# %% [code]
# melt together the data.
xx = sdf_scaled2['mononuc_freq'].melt(value_name="count ratio")
# append expanded region.
xx_exp = xx['colnames'].str.split("_", expand=True)
xx_exp.columns = ['base', 'region']
# calculate base pair using individual bases.
xx_exp['base pair'] = xx_exp['base'].replace({'G': 'GC', 'C': 'GC', 'A': 'AT', 'T': 'AT'})
xx = xx.join(xx_exp)

# %% [code]
# generate the grid
g = sns.FacetGrid(xx, col='region', row='base pair', hue='base')
# map a kde to it.
g.map(sns.distplot, 'count ratio')
g.add_legend()
plt.show()

# %% [markdown]
### Dinucleotide frequency

The count of dual-base nucleotides in mRNA base features.

#### Unnormalized 

The count of DNA bases comes under a count-like distribution and so we'll rely on the discrete *poisson* and/or continuous *exponential* distributions to model them.

In this case it's harder to tell given that the length plays a substantial role in determining how many of these counts can be detected. A larger number of these base counts appear to have significantly skewed distributions further to this.

#### Normalized

A number of 5'UTR dinucleotides have strong right skews, indicating a lack of normality in this subset of the gene sequence. 

# %% [code]
turb.plot.hist_grid(sdf, "dinuc_freq", "column", bins=20, shared_dist=['expon','chi2','poisson'])

# %% [code]
turb.plot.hist_grid(sdf_scaled1, "dinuc_freq", "column", bins=20, shared_dist=['norm', 't', 'gumbel_l', 'gumbel_r', 'expon', 'logistic', 'chi2'])

# %% [code]
turb.plot.hist_grid(sdf_scaled2, "dinuc_freq", "column", bins=20, shared_dist=['norm', 't', 'gumbel_l', 'gumbel_r', 'expon', 'logistic', 'chi2'])

# %% [markdown]
#### Seaborn to map comparisons between same dinucleotide, same pairing or different pairing.

# %% [code]
import re

# %% [code]
# melt together the data.
xx = sdf_scaled2['dinuc_freq'].melt(value_name="count ratio")
# append expanded region.
xx_exp = xx['colnames'].str.split("_", expand=True)
xx_exp.columns = ['base', 'region']

non_pair = [s for s in xx_exp['base'].unique() if re.search("(?:[GC][AT])|(?:[AT][GC])", s)]
pair = [s for s in xx_exp['base'].unique() if re.search("TA|AT|GC|CG", s)]
equal = [s for s in xx_exp['base'].unique() if re.search("(G{2}|A{2}|C{2}|T{2})", s)]
mmap = {**{a: "S" for a in equal}, **{a: "N" for a in non_pair}, **{a: "P" for a in pair}}

xx_exp['base group'] = xx_exp['base'].replace(mmap)
xx = xx.join(xx_exp)

# %% [code]
# generate the grid
g = sns.FacetGrid(xx, col='region', row='base group', hue='base', height=2)
# map a kde to it.
g.map(sns.distplot, 'count ratio')
g.add_legend()
plt.show()

# %% [markdown]
### Length proportion

# %% [code]
turb.plot.hist_grid(sdf, "_prop_", shared_dist=['gamma','beta'])

# %% [markdown]
### Minimum free energy

The minimum free energy expected when attempting to fold the 5'UTR region closest to the starting sequence CDS. Window specifies the number of bases away from this region we check.

#### Unnormalized 

The data follows a negative left skew with higher density as we approach zero. This is to be expected as this value is going to be influenced by the size of the window we are working with. For example a window of 10 means thato only the 10 bases nearest to CDS start are considered for free energy analysis.

#### Normalized

Normalizing by window length, and by inverting to make the score positive, we see the convergence of a normal distribution as the window size increases, which is quite interesting, but also suggests that samples of a small window size contain too much statistical noise to infer an effective distribution from.

# %% [code]
turb.plot.hist_grid(sdf, "^MFE", "column")

# %% [code]
turb.plot.hist_grid(sdf_scaled2, "^MFE", "column")

# %% [markdown]
### Amino acid meta features

These features are binary and hence the useful information contained in them will be low, but the interrelationships between these features could accumulate into a potentially useful metafeature or group of metafeatures.

# %% [code]
from turbopanda.utils import difference, intersect

# %% [code]
aamf = intersect(sdf_scaled2.view("_bcaa$"), sdf_scaled2.view(lambda z: z.nunique()==1))

# %% [code]
turb.plot.hist_grid(sdf_scaled2, difference(sdf_scaled2.view("_bcaa$"), aamf), "column", shared_dist="bernoulli")

# %% [markdown]
### Biophysical properties

These cover biophysical attributes relaying to the constitution of amino acids as determined by tried-and-tested metrics. There is minimal normalization applied to these features.

Isoelectric point clearly follows a bimodal distribution, which two groups forming around 'acidic' and 'basic' amino acids. The other metrics appear to be more balanced and normally distributed.

# %% [code]
turb.plot.hist_grid(sdf_scaled2, "_bp$", "column", shared_dist=['t', 'gumbel_r', 'gumbel_l', 'norm'])

# %% [markdown]
### Amino acid frequency

The frequency of amino acids is tied to the frequency of trinucleotides within the CDS region of the gene. 

#### Unnormalized 

Again, amino acid counts will be heavily biased by the length of the amino acid, so this needs to be corrected for.

#### Normalized

We have option two corrections:

1. Simply scale by amino acid length. 
2. Scale by CDS length, factoring in the number of synonymous codons each amino acid may have. i.e M has 1 trinucleotide it can use, whereas R has 6, so there is a bias simply in the number of trinucleotide selection, which we factor for.

We generally see in the normalized versions a more heavy right skewed-tail.

# %% [code]
turb.plot.hist_grid(sdf, "aa_freq", "column", shared_dist=['expon','chi2'], kde_range=1e-2)

# %% [code]
turb.plot.hist_grid(sdf_scaled1, "aa_freq", "column")

# %% [code]
turb.plot.hist_grid(sdf_scaled2, "aa_freq", "column", shared_dist=['gamma', 'norm', 'cauchy', 't', 'gumbel_l', 'gumbel_r'])

# %% [markdown]
### Translation frequency

The presense of methionine in the 5'UTR can be indicative of faster translation.

# %% [code]
turb.plot.hist_grid(sdf, "translation_freq", shared_dist=['bernoulli', 'poisson'])

# %% [code]
turb.plot.hist_grid(sdf_scaled2, "translation_freq", shared_dist=['expon','chi2'], kde_range=1e-2)

# %% [markdown]
### Post-translational modifications

# %% [code]
turb.plot.hist_grid(sdf, "_ptmc$")

# %% [code]
turb.plot.hist_grid(sdf_scaled2, "_ptmc$")

# %% [code]
```
