## Supplementary material for "Predicting Relative Protein Abundance via Sequence-Based Information": S3 Data: SDF-Supplementary-Base-Frequency.ipynb

```
# %% [markdown]
# Supplementary Material: Base Frequency

This notebook is supplementary material to Figure 2 of Paper.

# %% [code]
import sys
sys.path.insert(0,"../../../turbopanda")
import turbopanda as turb
import seaborn as sns
import numpy as np
import pandas as pd
import itertools as it

import matplotlib.pyplot as plt
%matplotlib inline

# %% [markdown]
# Average Profiles of each Nucleotide Frequency along the gene.

# %% [code]
sseq = turb.read("../../data/NCBI/rna_sequence_table.csv",sep="\t")

# %% [markdown]
## Calculate CDS, 5'UTR, 3'UTR

# %% [code]
coding = sseq.df_[sseq['protein_id'].notnull()].dropna(subset=['seq','cds_start','cds_end'])
# cast to int
coding.loc[:,'cds_start'] = coding['cds_start'].astype(np.int)
coding.loc[:,'cds_end'] = coding['cds_end'].astype(np.int)

# %% [code]
def extract_CDS(row):
    return row['seq'][row['cds_start']:row['cds_end']]
def extract_5p_utr(row):
    return row["seq"][:row["cds_start"]]
def extract_3p_utr(row):
    return row["seq"][row["cds_end"]:]

# %% [code]
coding['CDS'] = coding.apply(extract_CDS, axis=1)
coding["5_UTR"] = coding.apply(extract_5p_utr, axis=1)
coding["3_UTR"] = coding.apply(extract_3p_utr, axis=1)

# %% [markdown]
### Distribution of mRNA Length

Our window size is quite a bit below the bulk of mRNA transcripts.

# %% [code]
w=200

# %% [code]
fig, axes = plt.subplots(ncols=4, figsize=(18, 5))

for ax, LENGTH in zip(axes, ("seq", "CDS", "5_UTR", "3_UTR")):
    # plot
    sns.distplot(coding[LENGTH].str.len().apply(lambda x: np.log2(x+1)), kde=False, ax=ax)
    ax.vlines(np.log2(w), ymin=0, ymax=6000)
    ax.set_xlabel(r"$log_2\, L$ {}".format(LENGTH))
    ax.text(np.log2(w)+.2, 6000, r"window")
    ax.grid(True)
plt.show()

# %% [markdown]
### Profiling code to determine profiles and plot them:

# %% [code]
def chunks(n, k):
    """Yield successive n-sized chunks from lst."""
    for i in range(0, n, k):
        yield i, i+k

def mapped_counts(s, w=500, base="CG"):
    chk = list(chunks(len(s), w))
    return list(map(lambda r: s.count(base, r[0], r[1]), chk))

def determine_base_profile(seqs, sequence='seq', window=500, base="CG"):
    """
    window is the sub-length size in the sequence to count the number of instances of 'base'.
    
    Cases:
        mononucleotide: uniform_factor=4
        dinucleotide: uniform_factor=16
        trinucleotide: uniform_factor=64
        following 4**m where m is the length of 'base'.
    """
    seqs = seqs.df_ if isinstance(seqs, turb.MetaPanda) else seqs
    # generate
    counted_set = seqs[sequence].apply(mapped_counts, w=window, base=base).apply(np.asarray)
    # initialize numpy array - using NUMPY is way faster.
    dim = counted_set.apply(len).max()
    X = np.zeros((seqs.shape[0], dim), dtype=np.float64)
    # factor size
    uniform_factor=4**len(base)
    factor = window/uniform_factor
    # iterate over the rows and set to numpy
    for i, s in enumerate(counted_set):
        # calculate ratio as the NUMBER OF COUNTS IN WINDOW SCALED by MONO/DINUC EXPECTED NUMBER
        X[i, :s.shape[0]] = s / factor
    # convert zeros to np.nans 
    X[X <= 0] = np.nan
    # create pd.dataframe
    pdX = pd.DataFrame(X)
    # rename index
    ind = list(map(lambda x: x[0], list(chunks(dim*window, window))))
    pdX.columns = ind
    return pdX
    
def plot_base_profile(seqs, seq_name="seq", window=500, base=["CG"], k=-1, figsize=(12,6)):
    # turb function for designing the layout. yum.
    fig, axes = turb.visualise._generate_diag_like_grid(len(base), direction="column", ax_size=3)
    
    for b, ax in zip(base, axes):
        # get data
        L = len(b)
        # call our function with the appropriate column, window, base etc.
        pdX = determine_base_profile(seqs, seq_name, window, b)
        
        # first k, if relevant - calculate mean and std
        m1 = np.nanmean(pdX.iloc[:,:k], axis=0)
        std1 = np.nanstd(pdX.iloc[:,:k], axis=0)
        
        # plot
        ax.plot(pdX.columns.values[:k], m1, 'x-')
        ax.plot([0, pdX.columns.values[k]], [1., 1.], 'k--')
        ax.fill_between(pdX.columns.values[:k], m1-std1, m1+std1, alpha=.4)
        ax.set_xlabel("Sequence Index")
        ax.set_ylabel("Observed/Expected {base} count".format(base=b))
        ax.set_title("{} count".format(b))
        ax.grid()
        
    fig.tight_layout()
    plt.show()

# %% [code]
mononucs = ("A","C","T","G")
dinucs = list(it.starmap(lambda x,y:"".join([x,y]), it.product("ATCG",repeat=2)))

# %% [markdown]
## Profiles of aggregate mRNA sequences for each mononucleotide

# %% [code]
plot_base_profile(sseq, base=mononucs, window=200)

# %% [markdown]
## Profiles of aggregate mRNA sequences for each dinucleotide

# %% [code]
plot_base_profile(sseq, base=dinucs, window=200)

# %% [markdown]
## Profiles of aggregate CDS sequences for each mononucleotide

# %% [code]
plot_base_profile(coding, seq_name="CDS", base=mononucs, window=200)

# %% [markdown]
## Profiles of aggregate CDS sequences for each dinucleotide

# %% [code]
plot_base_profile(coding, seq_name="CDS", base=dinucs, window=200)

# %% [markdown]
## Profiles of aggregate 5'UTR sequences for each mononucleotide

# %% [code]
plot_base_profile(coding, seq_name="5_UTR", window=64, base=mononucs, k=-1)

# %% [markdown]
## Profiles of aggregate 5'UTR sequences for each dinucleotide

# %% [code]
plot_base_profile(coding, seq_name="5_UTR", window=64, base=dinucs, k=-1)

# %% [markdown]
## Profiles of aggregate 3'UTR sequences for each mononucleotide

# %% [code]
plot_base_profile(coding, seq_name="3_UTR", window=200, base=mononucs)

# %% [markdown]
## Profiles of aggregate 3'UTR sequences for each dinucleotide

# %% [code]
plot_base_profile(coding, seq_name="3_UTR", window=200, base=dinucs)

# %% [code]
```
