## Supplementary material for "Predicting Relative Protein Abundance via Sequence-Based Information": S4 Data: caching.py

```
#!/usr/bin/env python3
# -*- coding: utf-8 -*-
"""
Created on Thu Aug 15 15:15:24 2019

@author: gparkes
"""
import os
import numpy as np
import pandas as pd
import warnings

from IPython.display import Image
import matplotlib.pyplot as plt


__all__ = ["cache", "cache_img", "cache_numpy", "load_or_execute_csv", "load_or_draw"]


def cache(filename="example1.csv"):
    """
    Provides automatic .csv caching for pandas.DataFrames.

    use as:
    @cache("file.csv")
    def f(x):
        return pd.Series(x)

    f([1, 2, 3])
    """
    if not isinstance(filename, str):
        raise TypeError("'filename' must be of type [str]")
    if not filename.endswith(".csv"):
        raise ValueError("filename '{}' must end with .csv extension".format(filename))

    # define decorator
    def decorator(func):
        # parameters passed to this function.
        def caching_function(*args, **kwargs):
            if os.path.isfile(filename):
                return pd.read_csv(filename, index_col=0)
            else:
                df = func(*args, **kwargs)
                if isinstance(df, pd.DataFrame):
                    df.to_csv(filename)
                    return df
                else:
                    warnings.warn("returned object from func not of type [pd.DataFrame], not saved.", FutureWarning)
                    return df
        return caching_function
    return decorator


def cache_img(filename="example1.png"):
    """
    Provides automatic image caching for plot functions.

    Accepted image types are [png, pdf, svg, jpg]

    use as:
    @cache_img("nice.png")
    def f():
        fig, ax = plt.subplots()
        ax.plot(np.random.randn(100), np.random.randn(100))
        return fig

    f()
    """
    if not isinstance(filename, str):
        raise TypeError("'filename' must be of type [str]")
    ending = filename.split(".")[-1]
    if ending not in ["png", "svg", "pdf", "jpg"]:
        raise TypeError("filename extension does not end in [png, svg, pdf, jpg]")

    # define decorator
    def decorator(func):
        # parameters passed to this function.
        def caching_function(*args, **kwargs):
            if os.path.isfile(filename):
                return Image(filename)
            else:
                I = func(*args, **kwargs)
                if isinstance(I, plt.Figure):
                    I.savefig(filename, format=ending)
                    return
                else:
                    warnings.warn("returned object from func not of type [plt.Figure], not saved.", FutureWarning)
                    return
        return caching_function
    return decorator


def cache_numpy(filename="example1.npy"):
    """
    Provides automatic .npy caching for np.ndarray arrays.

    use as:
    @cache_numpy("file.npy")
    def f(x):
        return np.asarray(x)

    f([1, 2, 3])
    """
    if not isinstance(filename, str):
        raise TypeError("'filename' must be of type [str]")
    if not filename.endswith(".npy"):
        raise ValueError("filename '{}' must end with .csv extension".format(filename))

    # define decorator
    def decorator(func):
        # parameters passed to this function.
        def caching_function(*args, **kwargs):
            if os.path.isfile(filename):
                return np.load(filename, allow_pickle=False, mode="r")
            else:
                array = func(*args, **kwargs)
                if isinstance(array, np.ndarray):
                    # save array to file
                    np.save(filename, array, allow_pickle=False)
                    return array
                else:
                    warnings.warn("returned object from func not of type [np.ndarray], not saved.", FutureWarning)
                    return array
        return caching_function
    return decorator


def load_or_execute_csv(filename, f_exe, exe_args={}, csv_args={}, warning='raise'):
    """
    Loads a csv file in, if it exists, else executes a function passed to it
    with the corresponding keyword arguments.

    Function version of @cache.

    WARNING: This function *will* put files in your directory, or read them.

    Parameters
    --------
    filename : str
        The filepath and filename, 'with extension'. MUST BE a CSV
    f_exe : function
        A function to execute if filename is not present
    exe_args : dict
        Keywords to pass to f_exe
    csv_args : dict
        Keywords to pass to pd.read_csv and pd.to_csv
    warning : str
        ['raise', 'warn'], if raise, it produces an exception, else just warns

    Returns
    -------
    x : pd.DataFrame
        A pandas.DataFrame
    """
    # split csv_args into read/to_csv kws
    to_csv_kws = ["path_or_buf", "sep", "na_rep", "float_format", "columns", "header", "index",
                  "index_label", "compression", "chunksize"]
    read_csv_kws = ["filepath_or_buffer", "sep", "header", "names", "index_col", "usecols",
                    "squeeze", "prefix", "dtype", "engine", "na_values", "true_values",
                    "false_values", "skiprows", "nrows", "parse_dates", "compression",
                    "encoding", "iterator", "chunksize"]
    tcsv_l, rcsv_l = {}, {}

    for arg, val in csv_args.items():
        if arg in to_csv_kws:
            tcsv_l[arg] = val
        if arg in read_csv_kws:
            rcsv_l[arg] = val

    # check if file is present
    if os.path.isfile(filename):
        print("[Loading in from: {}]".format(filename))
        x = pd.read_csv(filename, **rcsv_l)
        if isinstance(x, (pd.DataFrame,pd.Series)):
            return x
        else:
            raise TypeError("'x' must be of type ['pd.Series','pd.DataFrame'] not '{}'".format(type(x)))
    else:
        print("[Executing and dumping to {}]".format(filename))
        x = f_exe(**exe_args)
        if isinstance(x, (pd.Series,pd.DataFrame)):
            x.to_csv(filename, **tcsv_l)
            return x
        else:
            if warning == 'raise':
                raise TypeError("'x' must be of type ['pd.Series','pd.DataFrame'] not '{}'".format(type(x)))
            elif warning == 'warn':
                warnings.warn("Return of function 'f' is not of type ['pd.Series','pd.DataFrame'], no data saved.")
                return x


def load_or_draw(filename, f_exe, exe_kwargs={}, savefig_kwargs={}):
    """
    For use in Jupyter Notebook.

    Loads or executes an algorithm to produce a matplotlib.Figure object to draw. Note that this
    only works with objects like .png, .jpg, .svg etc.

    WARNING: This function *will* put files in your directory, or read them.

    Parameters
    --------
    filename : str
        name of file if present
    f_exe : function
        The executing function to call
    exe_kwargs : dict
        Arguments to pass to f_exe
    savefig_kwargs : dict
        Arguments to pass to plt.savefig

    Returns
    -------
    None
    """
    if os.path.isfile(filename):
        return Image(filename)
    else:
        F = f_exe(**exe_kwargs)
        if isinstance(F, plt.Figure):
            F.savefig(filename, **savefig_kwargs)
        else:
            warnings.warn("Return of function 'f' was not of type 'plt.Figure', no data saved.")
```
