## Supplementary material for "Predicting Relative Protein Abundance via Sequence-Based Information": S4 Data: correlation.py

```
#!/usr/bin/env python3
# -*- coding: utf-8 -*-
"""
Created on Mon Sep  9 16:21:08 2019

@author: gparkes
"""

from numba import jit
import numpy as np
import pingouin as pg
import itertools as it
import pandas as pd
import seaborn as sns
import matplotlib.pyplot as plt

from sklearn import covariance as cov_pkg


def cov_to_corr(cov):
    """
    Given the covariance matrix, compute the correlation matrix
    """
    Dinv = np.diagflat(1. / np.sqrt(np.diag(cov)))
    return np.dot(Dinv, np.dot(cov, Dinv))


@jit(nopython=True)
def pcorr(P):
    """
    Given the precision matrix P, calculate the partial correlation matrix.
    """
    C_inv = np.zeros_like(P)
    # given precision matrix
    N,D = P.shape
    for i in range(N):
        for j in range(D):
            C_inv[i, j] = -(P[i, j] / (np.sqrt(P[i,i] * P[j,j])))
    # set diag to 1
    for i in range(N):
    	C_inv[i, i] = 1.

    return C_inv


@jit(nopython=True)
def r2_adjusted(r2, n, k):
    return 1. - ((n - 1) / (n - k))*(1-r2)


@jit(nopython=True)
def r2_adj_matrix(R):
    # assumes k is the number of dimensions D in R
    R_adj = np.zeros_like(R)
    N, D = R.shape
    for i in range(N):
        for j in range(D):
            R_adj[i, j] = 1. - ((N - 1) / (N - D))*(1-R[i, j])
    return R_adj


def pairwise_partials(df, method="spearman"):
    """
    Calculates the pairwise partial correlations between every feature in the
    dataframe.
    """
    partials = [pg.partial_corr(df, x=c1, y=c2,
                                covar=df.columns.difference([c1, c2]).tolist(),
                                method=method)
                for c1, c2 in it.combinations(df.columns, 2)]

    c1s = [c1 for c1, c2 in it.combinations(df.columns,2)]
    c2s = [c2 for c1, c2 in it.combinations(df.columns,2)]
    return pd.concat(partials, axis=0).assign(feature1=c1s, feature2=c2s)


def pairwise_set_partials(df, y, method="spearman"):
    """
    Given the dataset and 'y' column, calculate pairwise partials for each x with y, taking
    all other x's into account.
    """
    # remove all non-numeric columns
    num_features = df.columns[df.dtypes.ne(object)]
    ndf = df[num_features]
    # calculate partials
    partials = [pg.partial_corr(data=ndf, x=x, y=y, covar=ndf.columns.difference([x, y]).tolist(),\
                                method=method) for x in ndf.drop(y, axis=1).columns]

    return pd.concat(partials,axis=0).assign(feature1=ndf.drop(y,axis=1).columns,
                     feature2=[y]*(ndf.shape[1]-1))


def plot_covariance_models(df, dtype="covariance"):
    """
    Plot different scikit-learn data models in different plots.

    Parameters
    -------
    df : pd.DataFrame
        The dataset - numeric columns extracted
    dtype : str
        Choose which variable to plot. Choose [covariance/cov, correlation/corr,
         precision/prec, partial correlation/pcorr]

    Returns
    -------
    fig
    """
    if not isinstance(df, pd.DataFrame):
        raise TypeError("'df' is not of type [pd.DataFrame]")

    num_features = df.columns[df.dtypes.ne(object)]
    df_num = df[num_features].dropna()

    dtypes = ["covariance", "correlation", "precision", "partial correlation"]
    dtypes_short = ["cov", "corr", "prec", "pcorr"]

    cwrap = lambda x: pd.DataFrame(x, index=df.columns, columns=df.columns)
    functions = [
        lambda x: cwrap(x.covariance_), lambda x: cwrap(cov_to_corr(x.covariance_)),
        lambda x: cwrap(x.precision_), lambda x: cwrap(pcorr(x.precision_))
    ]

    if dtype in dtypes:
        F = dict(zip(dtypes, functions))[dtype]
    elif dtype in dtypes_short:
        F = dict(zip(dtypes_short, functions))[dtype]
    else:
        raise TypeError("'dtype' not recognized")

    gl_alpha = np.logspace(-3, 0, 30).tolist()

    # fit the models
    models = {
        "Empirical": cov_pkg.EmpiricalCovariance().fit(df_num),
        "Ledoit Wolf": cov_pkg.LedoitWolf().fit(df_num),
        "OAS": cov_pkg.OAS().fit(df_num),
        "Graphical Lasso": cov_pkg.GraphicalLassoCV(alphas=gl_alpha, cv=10, mode="cd", n_jobs=2).fit(df_num)
    }

    fig, axes = plt.subplots(ncols=4, figsize=(20, 4))
    # axes = list(it.chain.from_iterable(axes))
    for i, (name, mod) in enumerate(models.items()):
        sns.heatmap(F(mod), cmap="seismic", vmin=-1., vmax=1., ax=axes[i])
        axes[i].set_title(name)

    return fig
```
