## Supplementary material for "Predicting Relative Protein Abundance via Sequence-Based Information": S4 Data: extract_IDs.R

```
# this is a comment

source("http://bioconductor.org/biocLite.R")
biocLite("biomaRt")
# listing all of the databases in Biomart
library(biomaRt)
library(gdata)

# pre this stage - you need to download a list of all the ENSG genes that exist in Biomart!

# file path
path <- "../data/biomart/"
mir_path <- "../data/mirbase/"

# list all of the Ensembl mart availables
listEnsembl()

getwd()

# read in ENSEMBL ENSG labels for everything!
ensg <- read.table(paste(path,"ensg_all.txt",sep=""), header=1, sep="\n")
head(ensg)

# extract mir labels
mirdb <- read.xls(paste(mir_path, "miRNA.xls",sep=""), header=1)
head(mirdb)
# extract mirbase that begin with hsa (homo sapiens)
mir_ids = mirdb[grepl("^hsa.+", mirdb[, "Mature1_ID"]), "Accession"]
head(mir_ids)

# connect to the human dataset of Ensembl GRCh=37.
ensembl = useEnsembl(biomart="ensembl", GRCh=37, dataset="hsapiens_gene_ensembl")
ensembl
head(listFilters(ensembl))
head(listAttributes(ensembl))
listAttributes(ensembl)[0:60,]

# ENSEMBL-HGNC query
hgnc <- getBM(attributes=c('ensembl_gene_id','ensembl_transcript_id','ensembl_peptide_id',
                             'hgnc_id','hgnc_symbol','hgnc_trans_name'),
                filters = "ensembl_gene_id", values=ensg, mart = ensembl)
# check to be sure
head(hgnc)
# write to file
write.csv(hgnc, file = paste(path,"biomart_hgnc.csv",sep=""))
rm(hgnc)

# ENSEMBL-GO query
GO <- getBM(attributes=c(
  "ensembl_gene_id","ensembl_transcript_id","ensembl_peptide_id","go_id","name_1006"
), filters = "ensembl_gene_id", values=ensg, mart = ensembl)
head(GO)
write.csv(GO, file = paste(path, "biomart_go.csv", sep=""))
rm(GO)

# ENSEMBL-Refseq query
refseq <- getBM(attributes=c(
  "ensembl_gene_id","ensembl_transcript_id","ensembl_peptide_id","refseq_mrna","refseq_peptide","refseq_ncrna"
), filters = "ensembl_gene_id", values=ensg, mart = ensembl)
head(refseq)
write.csv(refseq, file = paste(path, "biomart_refseq.csv", sep=""))
rm(refseq)

# ENSEMBL-miRbase query
mirbase <- getBM(attributes = c(
  "ensembl_gene_id","ensembl_transcript_id","ensembl_peptide_id","mirbase_id","mirbase_accession","mirbase_trans_name"
), filters = "mirbase_accession", values=mir_ids, mart = ensembl)
head(mirbase)
write.csv(mirbase, file = paste(path, "biomart_mirbase.csv", sep=""))
rm(mirbase)

# ENSEMBL-Uniprot query
uniprot <- getBM(attributes = c(
  "ensembl_gene_id","ensembl_transcript_id","ensembl_peptide_id","uniprotswissprot","uniprot_gn","uniprotsptrembl"
), filters = "ensembl_gene_id", values=ensg, mart = ensembl)
head(uniprot)
write.csv(uniprot, file = paste(path, "biomart_uniprot.csv", sep=""))
rm(uniprot)

# ENSEMBL-Gene info query
geneinfo <- getBM(attributes = c(
  "ensembl_gene_id","ensembl_transcript_id","ensembl_peptide_id","chromosome_name","start_position","end_position","strand"
), filters = "ensembl_gene_id", values=ensg, mart = ensembl)
head(geneinfo)
write.csv(geneinfo, file = paste(path, "biomart_genechr.csv", sep=""))
rm(geneinfo)
```
