## Supplementary material for "Predicting Relative Protein Abundance via Sequence-Based Information": S4 Data: myfig.py

```
# utf-8
import matplotlib.pyplot as plt
import os
import numpy as np
import pandas as pd
import seaborn as sns
import itertools as it
import warnings

from matplotlib import patches
from matplotlib.collections import PatchCollection
from mpl_toolkits.axes_grid1.axes_divider import make_axes_locatable
from matplotlib import animation, colorbar
from scipy import stats

from typing import Tuple

# local
import misc_functions as misc


def _adjust_xy(xy, w, h):
    """
    Offset x and y by the width and height /2 to centralize the coordinates.
    """
    return xy[0] - (w / 2), xy[1] - (h / 2)


def save(fig: plt.Figure,
         plot_type: str,
         name: str = "example1",
         save_types: Tuple = ("png", "pdf"),
         filepath: str = "../Figures/",
         savemode: str = "first",
         **fig_kws) -> bool:
    """
	Given a matplotlib.Figure object, save appropriate numbers of Figures to the respective
	folders.

	Parameters
	----------
	fig : matplotlib.figure.Figure
		The figure object to save.
	plot_type : str
		The type of plot this is, accepted inputs are:
		{"scatter", "kde", "heatmap", "cluster", "bar", "hist", "kde", "quiver",
        "box", "line", "venn", "multi", "pie"}
	name : str, optional
		The name of the file, this may be added to based on the other parameters
	save_types : list/tuple, optional
		Contains every unique save type to use e.g {"png", "pdf", "svg", "ps", "eps"}
	filepath : str, optional
		The file path to the root directory of saving images
	savemode : str, optional
        ['first', 'update']: if first, only saves if file isn't present, if update,
        overrides saved figure
    fig_kws : dict
    	Additional keyword arguments to pass to plt.savefig()
	
	Raises
	------
	IOError
		When the filepath does not exist
	TypeError
		When `fig` is not of type matplotlib.figure.Figure
	ValueError
		When 'savemode' is not a valid option

	Returns
	-------
	success : bool
		Whether it was successful or not
	"""
    if type(fig) is not plt.Figure:
        raise TypeError("'fig' object must be a matplotlib.figure.Figure")

    accepted_types = (
        "scatter", "kde", "heatmap", "cluster", "bar", "hist", "kde", "quiver",
        "box", "line", "venn", "multi", "pie"
    )
    file_types_supported = ("png", "pdf", "svg", "eps", "ps")
    accepted_savemodes = ('first', 'update')

    if plot_type not in accepted_types:
        raise TypeError("plot_type: [%s] not found in accepted types!" % plot_type)

    for st in save_types:
        if st not in file_types_supported:
            TypeError("save_type: [%s] not supported" % st)
    if savemode not in accepted_savemodes:
        raise ValueError("savemode: '{}' not found in {}".format(savemode, accepted_savemodes))

    # correct to ensure filepath has / at end
    if not filepath.endswith("/"):
        filepath += "/"

    # check whether the filepath exists
    if os.path.exists(filepath):
        for t in save_types:
            # if the directory does not exist, create it!
            if not os.path.isdir(filepath + "_" + t):
                os.mkdir(filepath + "_" + t)
            # check if the figures themselves already exist.
            filename = "{}_{}/{}_{}.{}".format(filepath, t, plot_type, name, t)
            if os.path.isfile(filename):
                warnings.warn("Figure: '{}' already exists: Using savemode: {}".format(filename, savemode), UserWarning)
                if savemode == 'update':
                    fig.savefig(filename, format=t, bbox_inches='tight', **fig_kws)
            else:
                # make the file
                fig.savefig(filename, format=t, bbox_inches="tight", **fig_kws)

    else:
        raise IOError("filepath: [%s] does not exist." % filepath)

    return True


def create_gif(df,
               filename="example_dataset.gif",
               save=True,
               normaltest=True,
               hist_kws=dict(bins=10, color="blue"),
               anim_kws=dict(writer="pillow")):
    """
    From a pandas.DataFrame, save a .gif image of every histogram of each dataset, with a 1sec delay between each histogram.

    Parameters
    -------
    df : pd.DataFrame object.
        The dataset
    filename : str
        The filename to save the gif as
    save : bool
        Whether to save a gif
    normaltest : bool
        Whether to calculate Anderson-Darling tests for normality.
    hist_kws : dict
        arguments to pass to plt.hist()
    anim_kws : dict
        arguments to pass to animation.FuncAnimation.save
    """

    nframes = df.shape[1]

    if normaltest:
        # calculate anderson-darling tests.
        anderson_dists = ("norm", "expon", "logistic", "gumbel")
        # calculate the anderson for each distribtuion type, and select the one with the lowest score.
        best_dists = pd.DataFrame(
            {distribution: df.apply(lambda x: stats.anderson(x, dist=distribution)[0]) for distribution in
             anderson_dists})

    def update_hist(num, data):

        plt.cla()
        if normaltest:
            plt.hist(df.iloc[:, num].dropna(),
                     label="best-score: %.2f,\nbest-dist: %s" % (best_dists.min(axis=1).iloc[num],
                                                                 best_dists.idxmin(axis=1).iloc[num]), **hist_kws)
            plt.legend()
        else:
            plt.hist(df.iloc[:, num].dropna(), **hist_kws)

        plt.xlabel(df.columns[num])

    fig = plt.figure()
    plt.hist(df.iloc[:, 0].dropna(), **hist_kws)
    anim = animation.FuncAnimation(fig, update_hist, nframes, interval=1000, fargs=(df,))
    # save
    if save:
        anim.save(filename, **anim_kws)
        return fig
    else:
        return anim


def create_gif_compare(df1,
                       df2,
                       filename="example_dataset.gif",
                       save=True,
                       normaltest=True,
                       hist1_kws=dict(bins=10, color="blue"),
                       hist2_kws=dict(bins=10, color="red"),
                       anim_kws=dict(writer="pillow")):
    """
    Similar to .create_gif(), except we create two plots
    in order to enable comparison between df columns. Assumes the columns
    between df1 and df2 are the same.

    Parameters
    -------
    df1 : pd.DataFrame object.
        The dataset 1
    df2 : pd.DataFrame object.
        The dataset 2
    filename : str
        The filename to save the gif as
    save : bool
        Whether to save a gif
    normaltest : bool
        Whether to calculate Anderson-Darling tests for normality.
    hist1_kws : dict
        arguments to pass to ax[0].hist()
    hist2_kws : dict
        arguments to pass to ax[1].hist()
    anim_kws : dict
        arguments to pass to animation.FuncAnimation.save
    """

    if not isinstance(df1, pd.DataFrame):
        raise TypeError("df1 must be of type [pd.DataFrame]")
    if not isinstance(df2, pd.DataFrame):
        raise TypeError("df2 must be of type [pd.DataFrame]")

    shared_columns = df1.columns.intersection(df2.columns)
    ndf1 = df1[shared_columns]
    ndf2 = df2[shared_columns]
    nframes = len(shared_columns)

    # do normaltest stuff if true
    if normaltest:
        anderson_dists = ["norm", "expon", "logistic", "gumbel"]
        best_dists_1 = pd.DataFrame(
            {distribution: ndf1.apply(lambda x: stats.anderson(x, dist=distribution)[0]) for distribution in
             anderson_dists})
        best_dists_2 = pd.DataFrame(
            {distribution: ndf2.apply(lambda x: stats.anderson(x, dist=distribution)[0]) for distribution in
             anderson_dists})

    def update_hist(num, data):

        plt.cla()
        if normaltest:
            axes[0].hist(ndf1.iloc[:, num],
                         label="best-score: %.2f,\nbest-dist: %s" % (best_dists_1.min(axis=1).iloc[num],
                                                                     best_dists_1.idxmin(axis=1).iloc[num]),
                         **hist1_kws)
            axes[1].hist(ndf2.iloc[:, num],
                         label="best-score: %.2f,\nbest-dist: %s" % (best_dists_2.min(axis=1).iloc[num],
                                                                     best_dists_2.idxmin(axis=1).iloc[num]),
                         **hist2_kws)
            axes[0].legend()
            axes[1].legend()
        else:
            axes[0].hist(ndf1.iloc[:, num], **hist1_kws)
            axes[1].hist(ndf2.iloc[:, num], **hist2_kws)

        axes[0].set_xlabel(ndf1.columns[num])
        axes[1].set_xlabel(ndf2.columns[num])

    fig, axes = plt.subplots(ncols=2, figsize=(12, 5))
    axes[0].hist(ndf1.iloc[:, 0], **hist1_kws)
    axes[1].hist(ndf2.iloc[:, 0], **hist2_kws)
    anim = animation.FuncAnimation(fig, update_hist, nframes, interval=1000, fargs=(ndf1,))
    # if save
    if save:
        anim.save(filename, **anim_kws)
    else:
        return anim


def dataset_hist(df,
                 normaltest=True,
                 plot_arrange="square",
                 hist_kws=dict(bins=10)):
    """
    Plot every numeric column of the dataframe as a histogram.

    Parameters
    -------
    df : pd.DataFrame
        The dataset
    normaltest : bool
        Whether to calculate Anderson-Darling tests for normality
    plot_arrange : str
        Choose from ['square', 'row', 'column']. Default square. How
        to arrange plots.d
    hist_kws : dict
        Parameters to pass to sns.distplot

    Returns
    -------
    fig : matplotlib.Figure
        the figure.
    """

    # check types
    misc.instance_map([(df, pd.DataFrame), (normaltest, bool),
                       (plot_arrange, str), (hist_kws, dict)])

    df = df.copy()
    # modify the column names to shorten them.
    df.columns = misc.shorten_string(df.columns, n=20)

    # number of numeric features.
    num_features = df.columns[df.dtypes.ne(object)]

    if plot_arrange == "square":
        f1, f2 = misc.nearest_square_factors(len(num_features))
        fig, ax = plt.subplots(ncols=f1, nrows=f2, figsize=(2.5 * f1, 2.5 * f2))
        if len(num_features) > 3:
            # unchain
            ax = list(it.chain.from_iterable(ax))
    elif plot_arrange == "row":
        fig, ax = plt.subplots(ncols=len(num_features), figsize=(2.5 * len(num_features), 4))
    elif plot_arrange == "column":
        fig, ax = plt.subplots(nrows=len(num_features), figsize=(2.5, 2.5 * len(num_features)))
    else:
        raise ValueError("plot_arrange must be in [square, row, column]")

    # perform normaltest
    if normaltest:
        dist_types = ["norm", "expon", "logistic", "gumbel"]
        best_dists = pd.DataFrame(
            {d: df[num_features].apply(lambda x: stats.anderson(x, dist=d)[0]) for d in dist_types}
        )
        dd_templ = "%s: %.2f"
        best_score = best_dists.min(axis=1)
        best_label = best_dists.idxmin(axis=1)
    else:
        dd_templ = "No test"

    # use sns.distplot with kde=false
    for a, f in it.zip_longest(ax, num_features):
        a.set_xticklabels([])
        a.set_yticklabels([])
        a.set_xlabel("")
        if f is None:
            continue
        if normaltest:
            sns.distplot(df[f].dropna(), ax=a, kde=False, norm_hist=True,
                         label=dd_templ % (best_label.loc[f], best_score.loc[f]), **hist_kws)
        else:
            sns.distplot(df[f].dropna(), ax=a, kde=False, norm_hist=True,
                         label=dd_templ, **hist_kws)
        a.set_title(f)
        a.set_xlabel("")
        a.legend()

    fig.tight_layout()
    return fig


def plot_bag(data_bag, plotting_func, *func_args, **func_kwargs):
    """
    Generates a plot for each pandas.DataFrame within a list/bag, using the appropriate
    plotting function, with arguments and keyword arguments.

    Parameters
    -------
    data_bag : list/tuple/dict of pandas.DataFrame
        The group of data to plot in different plots. If dictionary, the key becomes
        the title of the plot.
    plotting_func : function
        Choose from ["hist", "heatmap"]
    func_args : list
        arguments to pass to plt.hist, sns.heatmap
    func_kwargs : dict
        keyword arguments to pass to plotting_func

    Returns
    -------
    None
    """

    f1, f2 = misc.nearest_square_factors(len(data_bag))
    fig, axes = plt.subplots(ncols=f1, nrows=f2, figsize=(4 * f1, 4 * f2))
    if len(data_bag) > 3:
        axes = list(it.chain.from_iterable(axes))

    if isinstance(data_bag, (list, tuple)):
        for i, dg in enumerate(data_bag):
            _ = plotting_func(dg, ax=axes[i], *func_args, **func_kwargs)
    elif isinstance(data_bag, dict):
        for i, (name, dg) in enumerate(data_bag.items()):
            _ = plotting_func(dg, ax=axes[i], *func_args, **func_kwargs)
            axes[i].set_title(name)
    else:
        raise TypeError("data_bag must be of type [list, tuple, dict], not '{}'".format(type(data_bag)))

    fig.tight_layout()
    return


def plot_scatter_grid(df, xs, y):
    """
    Plots a grid of scatter plots comparing each column in xs to y value.

    Parameters
    --------
    df : pd.DataFrame
        The dataset
    xs : list of str
        Column names to plot
    y : str
        The y-response variable to plot

    Returns
    -------
    None
    """
    f1, f2 = misc.nearest_square_factors(len(xs))
    fig, axes = plt.subplots(ncols=f1, nrows=f2, figsize=(3 * f1, 3 * f2))
    if len(xs) > 3:
        axes = list(it.chain.from_iterable(axes))

    for i, x in enumerate(xs):
        axes[i].scatter(df[x], df[y], alpha=.5)
        # corr
        pair_corr = df[[x, y]].corr(method="spearman").iloc[0, 1]
        axes[i].set_title("{}: r={:0.3f}".format(x, pair_corr))
    fig.tight_layout()
    return


def despine(axes, keep=[]):
    """
    Given a matplotlib.ax object, despine the entire plot.

    Parameters
    ---------
    axes : matplotlib.ax or list of matplotlib.ax
        The axes to despine
    keep : list of str (top, bottom, left, right)
        Tell the function which spines to keep if any

    Return
    -------
    None
    """
    orientations = {"top", "right", "left", "bottom"}
    to_remove = list(orientations - set(keep))

    if isinstance(axes, (list, tuple)):
        for ax in axes:
            for r in to_remove:
                ax.spines[r].set_visible(False)
    else:
        for r in to_remove:
            axes.spines[r].set_visible(False)


def detick(axes, axis=["x", "y"]):
    """
    Given a matplotlib.ax object, remove ticks and ticklabels from it.

    Parameters
    -------
    axes : matplotlib.ax or list of matplotlib.ax
        The axes to detick

    Returns
    -------
    None
    """
    if isinstance(axes, (list, tuple)):
        # apply detick to all axes
        for ax in axes:
            if "x" in axis:
                ax.set_xticks([])
                ax.set_xticklabels([])
            if "y" in axis:
                ax.set_yticks([])
                ax.set_yticklabels([])
    else:
        if "x" in axis:
            axes.set_xticks([])
            axes.set_xticklabels([])
        if "y" in axis:
            axes.set_yticks([])
            axes.set_yticklabels([])


def annotate_rects(axes,
                   locations,
                   sizes,
                   descriptions,
                   color="r",
                   textcolor="white",
                   alpha=.3):
    """
    Draws 'n' rectangles on your axes object, given locations, sizes and descriptions
    to place on top of the rectangle.

    Parameters
    -------
    axes : matplotlib.ax
        The axes to draw on
    locations : list of 2-tuples
        Some locations to draw at
    sizes : list of 2-tuples
        The size of the rectangles
    descriptions : list of str
        Text to draw on each rect
    color : str or hex
        Passed directly to draw function
    alpha : float
        The alpha of all rects

    Returns
    ------
    None
    """
    PATCH = [
        patches.Rectangle(
            _adjust_xy(l, s[0], s[1]), s[0], s[1]
        ) for l, s in zip(locations, sizes)
    ]
    p = PatchCollection(PATCH, alpha=alpha, color=color)
    axes.add_collection(p)
    # add text
    for l, d in zip(locations, descriptions):
        axes.annotate(d, xy=l,
                      xytext=l, xycoords="data",
                      ha="center",
                      fontsize=15, color=textcolor)


def annotated_circles(ax, locations, sizes, descriptions, color="r", alpha=.3):
    """
    Draws 'n' circles on your axes object, given locations, sizes and descriptions
    to place on top of the rectangle.

    Parameters
    -------
    axes : matplotlib.ax
        The axes to draw on
    locations : list of 2-tuples
        Some locations to draw at
    sizes : list of 2-tuples
        The size of the circles
    descriptions : list of str
        Text to draw on each circle
    color : str or hex
        Passed directly to draw function
    alpha : float
        The alpha of all circle

    Returns
    ------
    None
    """
    PATCH = [patches.Ellipse(l, s[0], s[1]) for l, s in zip(locations, sizes)]
    p = PatchCollection(PATCH, alpha=alpha, color=color)
    ax.add_collection(p)
    # add text
    for l, d in zip(locations, descriptions):
        ax.annotate(d, xy=l, xytext=l,
                    xycoords="data", ha="center",
                    fontsize=15, color="white")


def custom_colorbar(axes, label="colorbar"):
    """
    Create a custom and attractive colorbar on top of the plot.

    Parameters
    --------
    axes : matplotlib.ax
        The axes to draw on
    label : str
        The label to put on the colorbar

    Returns
    ------
    None
    """
    ax_divider = make_axes_locatable(axes)
    # define size and padding of axes for colorbar
    cax = ax_divider.append_axes('top', size='3%', pad='5%')
    # make colorbar for heatmap.
    # Heatmap returns an axes obj but you need to get a mappable obj (get_children)
    colorbar.colorbar(axes.get_children()[0], cax=cax, orientation='horizontal')
    # locate colorbar ticks
    cax.xaxis.set_ticks_position('top')
    cax.set_xlabel(label)


def rotate_labels(axes, degree=0, axis="x", align="left"):
    """
    Rotate the ticklabels by a given degree on a chosen axis.

    Parameters
    --------
    axes : matplotlib.ax, list or tuple
        The axes to rotate on
    degree : int
        The rotation in degrees
    axis : str
        'x' or 'y', which axes to rotate the labels on
    """
    if not isinstance(axes, (list, tuple)):
        # pluralize
        axes = [axes]

    for ax in axes:
        if axis == 'x':
            for tick in ax.get_xticklabels():
                tick.set_rotation(degree)
                tick.set_horizontalalignment(align)
        elif axis == 'y':
            for tick in ax.get_yticklabels():
                tick.set_rotation(degree)
                tick.set_horizontalalignment(align)
        else:
            raise ValueError("axis must be in ['x', 'y']")


def colormap_to_hex(cm_array):
    """
    Given a colormap array arranged as:
        [[r, g, b, a],
         [r, g, b, a],
         ............,
         [r, g, b, a]]

    Computes the hexadecimal for each row and returns as list
    """
    if isinstance(cm_array, np.ndarray):
        return ["#%02x%02x%02x" % (int(r * 255), int(g * 255), int(b * 255)) for r, g, b, _ in cm_array]
    elif isinstance(cm_array, pd.Series):
        # convert to ndarray
        cm_array = pd.DataFrame(np.vstack(cm_array.values))
    if isinstance(cm_array, pd.DataFrame):
        if cm_array.shape[1] == 4:
            return ["#%02x%02x%02x" % (int(r * 255), int(g * 255), int(b * 255)) for idx, (r, g, b, _) in
                    cm_array.iterrows()]
        elif cm_array.shape[1] == 3:
            return ["#%02x%02x%02x" % (int(r * 255), int(g * 255), int(b * 255)) for idx, (r, g, b) in
                    cm_array.iterrows()]
        else:
            raise ValueError("dimension of 'cm_array' must be 3 or 4, not {}".format(cm_array.shape[1]))
```
